# DNA-encoded Library Screening Uncovers Potent DNMT2 Inhibitors Targeting a Cryptic Allosteric Binding Site

**DOI:** 10.1101/2025.01.15.632061

**Authors:** Ariane F. Frey, Merlin Schwan, Annabelle C. Weldert, Valerie Kadenbach, Jürgen Kopp, Zarina Nidoieva, Robert A. Zimmermann, Lukas Gleue, Collin Zimmer, Marko Jörg, Kristina Friedland, Mark Helm, Irmgard Sinning, Fabian Barthels

## Abstract

The human RNA methyltransferase DNMT2 is thought to be involved in various pathophysiological processes, yet, a major challenge in drug targeting DNMT2 is given by the fact that current SAH-derived inhibitors have poor target selectivity and limited cellular permeability. In this study, we have performed a DNA-encoded library (DEL) screening on DNMT2 yielding five non-SAH-like hit structures, three of which feature a peptidomimetic scaffold. All DEL hits could be validated by orthogonal biophysical and biochemical assays for DNMT2 binding. At the same time, the lead structure did not interact with related methyltransferases from the DNMT and NSUN families highlighting an unmatched DNMT2-targeting selectivity profile. Subsequent crystallographic studies revealed the unique ligand binding mode including an active site loop rearrangement and the formation of a cryptic allosteric binding pocket able to modulate the enzymatic activity by non-covalent DNMT2 dimerization. Based on the crystallographic results, we performed a structure-activity relationship study around the inhibitor lead structure resulting in an optimized DNMT2 inhibitor (*K_D_*=3.04 µM), which was able to reduce m^5^C levels in MOLM-13 tRNA.

**GRAPHICAL ABSTRACT:** 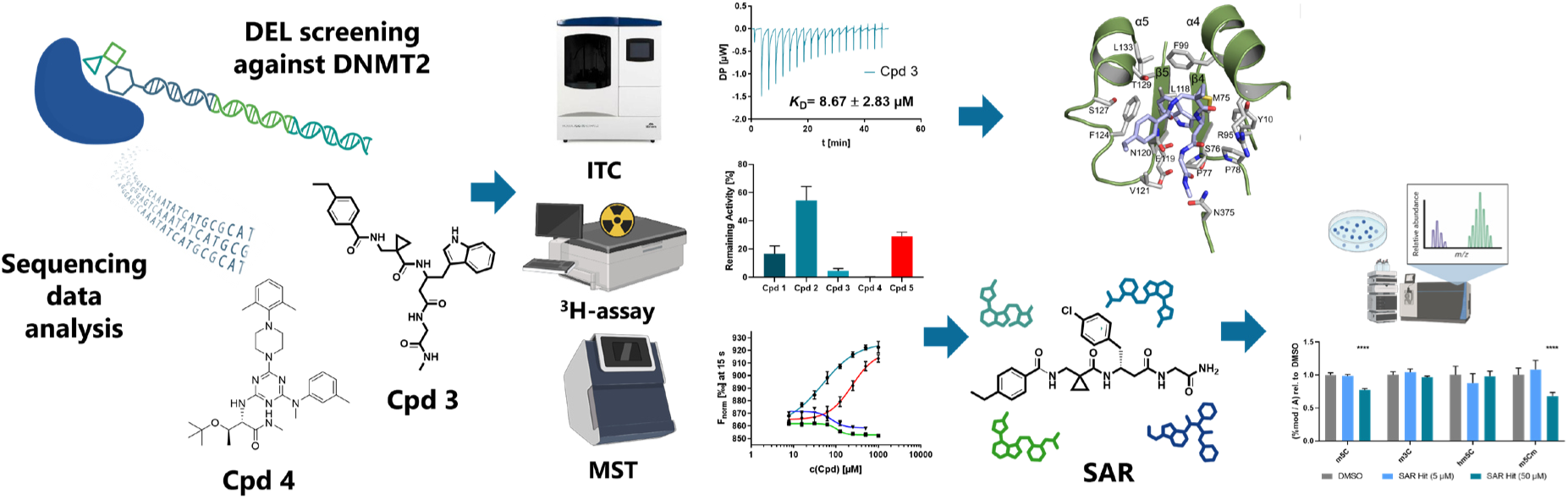

## INTRODUCTION

Targeting RNA methyltransferases (MTases) by active site-competitive small-molecule drugs proves challenging due to their universal dependence on *S*-adenosyl-L-methionine (SAM) as a methyl group donating substrate, which is transformed into *S*-adenosylhomocysteine (SAH) during this enzymatic reaction.(1, 2) SAH itself presents a pan methyltransferases inhibitor used as a starting point for several RNA MTase drug discovery campaigns.(3–5) A unique representative of this enzyme family is the RNA MTase DNA methyltransferase 2 (DNMT2) which is structurally related to other DNMTs, but this paralogous enzyme deviates in function and localization.(6–8) Instead of DNA, which is the main target of DNMT1 and DNMT3, DNMT2 introduces the m^5^C38 modification in several tRNAs.(6) Furthermore, DNMT2 is controversially discussed to be found in both nuclear compartments and the cytoplasm, hinting at its diverse roles and influence in protein biosynthesis.(9–11) Its distinctive catalytic MTase domain, featuring the unique cysteine-phenylalanine-threonine (CFT) motif, underscores its evolutionary and functional uniqueness within the DNMT family.(11) Physiologically, DNMT2 targets various tRNAs, including tRNA^Gly^, tRNA^Val^, and tRNA^Asp^, depending on the species investigated.(9, 12) tRNA methylation by DNMT2 plays a role in regulating RNA degradation by ribonucleases, thereby enhancing tRNA stability and overall protein expression.(10, 13) Additionally, tRNA m^5^C38 methylation improves codon recognition, particularly distinguishing between aspartate and glutamate, thus enhancing translational accuracy.(14) Given DNMT2’s role in regulating protein expression and translational accuracy, it is not surprising that DNMT2 misregulation has been implicated in cancer pathogenesis.(15) DNMT2 was found to be overexpressed in various cancer cells, such as cervical or bladder tissue, and leukemia, assuming DNMT2 plays a functional role in tumorigenesis.(15–17) Additionally, DNMT2 silencing was found to correlate with miRNA upregulation which is also related to proliferation and function as tumor suppressor.(18) Also, elevated m^5^C levels in small non-coding RNA (sncRNA), introduced by DNMT2, have been linked to metabolic disorders, where these influence the disorders’ inheritance induced by high-fat diets in mice.(19, 20) During oxidative stress, a DNMT2 up-regulation could be detected, leading to the assumption that DNMT2 also helps cells cope with exogenous stress factors, underscoring DNMT2’s multifaceted involvement in disease mechanisms.(21)

Despite its association with various diseases, DNMT2 remains a challenging target for drug development.(22) This is primarily due to the abundance of SAH and SAM and their ubiquitous binding proteins in cells with which inhibitors must compete.(23–25) Moreover, the fact that over 200 human MTases depend on SAM further complicates the discovery of selective inhibitors.(26) Efforts to develop inhibitors targeting DNMT2 have predominantly focused on SAH analogs.(1, 3, 4) While some inhibitors exhibit selectivity against the NSUN family which is also responsible for m^5^C modifications in tRNA, none have so far achieved selectivity towards DNMT3.(3) Current inhibitors also failed to inhibit DNMT2 within cellular assays, probably due to the low permeability of zwitterionic SAH derivates.(3) Notably, the most effective DNMT2 inhibitors to date possess enzymatic IC_50_ values around 2.5 µM.(4)

To address the development of cellularly potent DNMT2 inhibitors, in this study, we have applied the DNA-encoded library (DEL) screening technology on DNMT2 to identify novel modulating small molecule chemotypes. The goal was to develop the first selective chemical tools for DNMT2 investigation, modulation, and inhibition. DEL have gained widespread popularity in both industry and academia as a powerful tool for drug-like hit generation.(27, 28) Comprising unique molecular structures tagged with DNA barcodes, DEL enables efficient diverse compound library screening and groundbreaking identification of novel drug scaffolds.(29, 30) Recently, Wuxi AppTec enabled academic users to open access DEL, by providing a DELopen platform containing a library of 4.2 billion unique compounds.(31) Latest DELopen successes include the ligand discovery for diverse proteins such as thrombin, RNase L, E3 ligases, the transcription factor TEAD, and antibacterial drug targets, highlighting its broad utility across various target classes.(30, 32–36)

## MATERIAL AND METHODS

### DEL screening

DEL screening against human DNMT2 protein was performed with the fourth generation DELopen library (4.2 billion compounds, 27 sub-libraries) by WuXi AppTec. Screening was carried out according to the manufacturer’s protocol (available online: https://hits.wuxiapptec.com/assets/pdf/DELopen_Protocol.pdf). Briefly, the DEL panning was performed with recombinant human full-length DNMT2 protein immobilized via the His-tag on ThermoScientific HisPur™ Ni-NTA magnetic beads. The beads were first washed in a magnetic separation rack with washing buffer (50 mM HEPES, 150 mM NaCl, 5 mM MgCl_2_, 0.05% polysorbate-20, pH 7.5), and 250 μg beads were transferred for each selection condition to a separate tube. DNMT2 was diluted according to the manufacturer’s protocol in selection buffer (50 mM HEPES, 150 mM NaCl, 5 mM MgCl_2_, 0.05% Polysorbate-20, pH 7.5, 0.1 mg/mL sheared salmon sperm DNA, 1 mM TCEP, and 10 mM imidazole) and successful immobilization on the beads was confirmed by the SDS-PAGE protein capture assay described in manufacturer’s protocol. A differential panning procedure was employed with four different selection conditions (A–D): (**A**) Native DNMT2 without additives; (**B**) DNMT2 with SAH (100 µM); (**C**) DNMT2 with tRNA^Asp^ (10 µM); (**D**) No target control with no DNMT2 immobilized.

After three iterative selection rounds, the enriched DNA samples were transferred to WuXi AppTec for further processing including PCR amplification, quantitation by qPCR, and deep sequencing. All samples passed the DELopen quality control (SI Figure 1A, E) and were forwarded to Illumina NovaSeq deep sequencing. Subsequently, DEL hits selection (SI Figure 1C) was decided based on enrichment scores which were calculated by the DELopen unique algorithm described by Kuai et al.(37)

### Microscale thermophoresis (MST)

Screening for ligand binding and evaluation of dose-response affinity profiles was performed via microscale thermophoresis (MST) as described previously.(38) The measurements were conducted with a Nanotemper Technologies Monolith NT.115 device. All samples were measured in Monolith standard capillaries at 25 °C. The acquisition mode was set to “Nano-Blue” with an excitation power of 30% and a medium MST power, while the system was controlled using the software MO.Control (version 1.6.1) and for evaluation MO.Affinity Analysis (version 2.3) was used. The samples contained 2 µM DNMT2, 100 nM FTAD (5-FAM-triazolyl-adenosyl-Dab) probe, and 1.3% DMSO in MST buffer (50 mM HEPES, 150 mM NaCl, 1 mM dithiothreitol (DTT), 0.1% PEG-8000, 0.05% polysorbate-20, pH 7.4).(38) All compounds were employed at a final concentration of 100 µM for the initial ligand screening.

For *K_D_*-value determination of the DEL hit-derived fluorescent tracers **6**–**8**, the respective probe was used at a concentration of 100 nM (instead of FTAD) and the DNMT2 concentration was varied (50 µM–0 µM). All measurements were performed as triplicates. The *K_D_*-value was calculated and plotted in GraphPad Prism 8.0.1. MST experiments with fluorescently labeled tRNA (non-covalent labeling with SybrGold) were used to investigate if hit compounds **1**–**5** can dissociate the nucleic acid-protein complex formed by DNMT2 and were conducted as described previously.(39) To assess the DEL ligands’ effect on the RNA binding, the tRNA^Asp^ substrate (100 nM) was mixed with the DNMT2 enzyme (2 µM) supplemented with 1x SybrGold and 100 µM of the respective ligand.

### Isothermal titration calorimetry (ITC)

The DNMT2 protein was purified via size exclusion chromatography eluting in isothermal titration calorimetry (ITC) buffer (50 mM phosphate pH 8.0, 300 mM NaCl, 1 mM EDTA, 2 mM β-mercaptoethanol, 0.1% polysorbate-20) and directly used for measurements.(3) All compounds were dissolved in DMSO to a concentration of 5 mM and were then diluted in ITC buffer to a final concentration of 500 µM. DNMT2 was diluted to a final concentration of 50 µM (for compounds **5**, **14**, **47**, **11**, **16**), resp. 30 µM (for compounds **10**, **30**, **3**), resp. 1 mM (for compound **2**) with 10% DMSO to match the ligand stocks’ DMSO concentration. For the ITC measurements, a MicroCal PEAQ-ITC Automated workstation (Malvern Panalytical) with a 200 μL Hastelloy cell and an injection syringe volume of 40 μL was used. All experiments were performed in triplicates at 25 °C. For each measurement, 19 compound injections (2 μL each) with an injection speed of 0.5 μL/s were added to the reaction cell, containing DNMT2. The spacing time between sequential injections was 150 s (or 300 s for compounds **3**, **5**, **11**, **14**, **16**, **47**) and the stirring speed was set to 750 rpm with a reference power of 42 μW. As control experiments, ligands were titrated into ITC buffer. Data analysis and fitting were carried out using the MicroCal PEAQ-ITC Analysis Software 1.21 and plotted in GraphPad Prism 8.0.1.

Furthermore, displacement ITC experiments were performed as described for the direct titrations above with the following modifications: DNMT2 (50 µM) was preincubated with compound **10** (500 µM, 10% DMSO) or compound **3** (100 µM, 10% DMSO). Subsequently, SAH (500 µM, 10% DMSO) was titrated to the preincubated protein. During the same sample set, SAH (500 µM) was titrated directly into DNMT2 (50 µM) as a reference titration. Data analysis and displacement curve fitting were accomplished via the MicroCal PEAQ-ITC Analysis Software.

### Fluorescence polarization (FP)

Fluorescence polarization (FP) measurements with RNA MTases and small-molecule fluorescent tracers were conducted in black Greiner 96-well half-area plates using a Tecan Spark 10M plate reader as described previously.(40) The reader was equipped with polarization filters coupled to a monochromator setup (λ_ex_ = 480 nm, λ_em_ = 530 nm). For selectivity screening, reaction mixtures contained 2 µM recombinant enzyme (DNMT1, DNMT2, DNMT3A/3L, NSUN2, NSUN3, or NSUN6; preparation see below) and 10 nM DEL hit-derived probe **6** in MST buffer. For displacement experiments, 100 µM SAH or 10 µM tRNA^Asp^ were added from buffered stock solutions to the DNMT2-containing mixtures. All measurements were performed as triplicates. Polarization values (in mP) were calculated from polarization-specific parallel and orthogonal fluorescence intensities according to the Tecan in-built calculation routine. *K_D_*-values were calculated and plotted in GraphPad Prism 8.0.1.

### 3H-based methyltransferase enzyme activity assays

Enzymatic DNMT2 inhibition assays were carried out in 100 mM Tris-HCl, pH 8.0, 100 mM NH_4_OAc, 0.1 mM EDTA, 10 mM MgCl_2_, and 10 mM DTT as described previously.(3, 4) First, tRNA^Asp^ substrate was heated to 75 °C for 5 min and cooled down to room temperature (RT) before it was added to the reaction mixture to a final concentration of 5 μM. The reaction mixture contained 250 nM DNMT2 and the ligand compounds at variable concentrations (5% DMSO). Enzymatic reactions were started by the addition of 0.9 µM SAM as a mixture of ^3^H-SAM (Hartmann Analytics) resp. cold SAM (NEB) to a final activity of 0.025 μCi μL^−1^. Enzyme assays were carried out for 30 min at 37 °C. As a negative control served the reaction mixture without DNMT2 and as a positive control the reaction mixture without test compound. After 30 min, aliquots of 8 μL were taken from the reaction mixture and spotted on Whatman glass microfiber filters (GF/C, 25 mm). The RNA was precipitated on the filters with 5% ice-cold TCA for 15 min. The filters were washed twice with 5% TCA at RT for 20 min and 10 min and once in EtOH for 10 min. After drying, the filters were transferred into scintillation vials and 3 mL of PerkinElmer Gold MV liquid scintillation cocktail was added. Scintillation was measured with a scintillation counter (TriCarb Liquid Scintillation Analyzer 4810TR; measurement time of 1 min). For IC_50_-value determination, compounds were analyzed at a minimum of six different concentrations in triplicates. IC_50_-values were calculated evaluated and plotted in GraphPad Prism 8.0.1.

### Affinity selection-mass spectrometry (AS-MS)

Orthogonal *K_D_*-value determination for compound **3** was conducted by an affinity selection-mass spectrometry (AS-MS) assay according to Prudent et al.(41) and Simon et al.(42) Compound **3** was dissolved in DMSO (20 mM) and a dilution series in Tris buffer (25 mM Tris, 100 mM NaCl, pH 7.5) was prepared to give final ligand concentrations of 100 µM, 50 µM, 25 µM, 12.5 µM, 6.25 µM, 3.12 µM, 1.56 µM, and 0 µM. DNMT2 was diluted in Tris buffer (130 µM) and added to the dilution series to give a final concentration of 5 µM DNMT2. Zeba Spin Desalting columns 7K MWCO were used for the affinity-based separation. These columns were centrifuged at 2000 g for 1 min to remove the storage buffer. 20 µL of the prepared samples were loaded to the columns and the protein-ligand complex was eluted by centrifugation at 2000 g for 2 min. The centrifuged solutions were diluted with 10 µL ACN and analyzed by LC/MS with an HP Agilent 1100 Series HPLC system. The data evaluation and subsequent binding affinity determination were performed with MestReNova (version 14.0.1) and GraphPad Prism 8.0.1.

### Nano Differential Scanning Fluorimetry (nanoDSF)

Thermal shift assays were carried out in triplicates with the NanoTemper Prometheus NT.48 nanoDSF instrument using the manufacturer-designated capillaries.(43) Sample solutions contained 5 μM DNMT2 protein in buffer (25 mM Tris, 100 mM NaCl, pH 7.5) supplemented with variable concentrations of compound **2** and 10 vol% DMSO. In the capillaries, sample solutions were heated from 20 °C to 80 °C with a heating rate of 1.5 °C/min, and fluorescence emission was recorded at 330 nm and 350 nm, while excitation was carried out at 280 nm. The measured fluorescence ratio of the detected fluorescence at 330 nm and 350 nm was plotted as a function of temperature using GraphPad Prism 8.0.1. The melting temperature was calculated by the vertical slice method described by Bai et al.(44)

### In vitro permeability assay (PAMPA)

To determine the passive permeability of selected DNMT2 ligands, a parallel artificial membrane permeation assay (PAMPA) was performed as described previously.(45) The experiment was performed in duplicates with Sigma-Aldrich MAIPNTR10 donor plates with 5 µL of an artificial membrane (1% w/v L-α-phosphatidylcholine in *n*-dodecane) per well. After incubation for 7 h, the acceptor solutions were evaluated by LC/MS (HP Agilent 1100 Series). The peak areas in the UV chromatogram (λ = 210 nm) were used to calculate the areas under the curve (AUCs). The apparent permeability *P_app_* was calculated using the following equation with V_D_ and V_A_ as volumes of donor and acceptor solutions (0.15 cm^3^ and 0.4 cm^3^), AUC_acc_ and AUC_ref_ as the area of the analyt signal in the respective chromatogram of sample and reference solutions, A as the porosity-corrected filter area (0.2113 cm^2^) and t as the incubation time given in seconds: *P_app_*=−V_D_⋅V_A_⋅ln(1−AUC_acc_/AUC_ref_)/[(V_D_+V_A_)⋅A⋅t].

### Protein and tRNA production

DNMT2, NSUN2, NSUN6, and DNMT3A/3L were expressed from *E. coli* as described previously, while tRNA^Asp^ was synthesized by in vitro transcription as described previously.(3)

### DNMT2 cloning, expression, and purification for co-crystallization

For the co-crystallization of DNMT2Δ47 with compound **3**, the coding DNA sequence of human DNMT2 lacking residues 191–237 was amplified by PCR and cloned into a pETHis_1a vector.(46) For this purpose, overhangs for the restriction enzymes NcoI/BamHI were used. The plasmid was transformed in Rosetta II (DE3) *E. coli* (Merck-Novagen) and expressed in ZY autoinduction medium supplemented with trace elements, kanamycin, and chloramphenicol. The cells were grown to OD_600_ = 0.8 at 37 °C (220 rpm shaking) and further cultivated overnight at 20 °C. Harvesting was performed by centrifugation and washing with 1xPBS buffer (0.137 M NaCl, 2.7 mM KCl, 10 mM Na_2_HPO_4_, 1.8 mM KH_2_PO_4_). The resulting cell pellet was then resuspended in lysis buffer (20 mM HEPES, pH 7.5, 500 mM NaCl, 20 mM imidazole, 1 M urea, 4 mM β-mercaptoethanol), supplemented with a protease inhibitor mix (Roche) and 1:10000 benzonase. Lysis was performed with a microfluidizer (M1-10L, Microfluidics) followed by clearing the lysate applying centrifugation (40 minutes at 50,000 g) and filtering through a 0.45 μm membrane. The supernatant was loaded twice on 4 mL Ni-NTA Agarose (Qiagen) for Ni-IMAC using gravity flow columns (Biorad). The beads were washed with ten column volumes (10 CV) wash buffer (20 mM HEPES, pH 7.5, 150 mM NaCl, 4 mM β-mercaptoethanol), followed by 10 CV high salt buffer (20 mM HEPES, pH 7.5, 2 M NaCl, 4 mM β-mercaptoethanol) and again 10 CV wash buffer. Elution of the His-tagged DNMT2Δ47 was performed with 5 CV elution buffer (20 mM HEPES, pH 7.5, 150 mM NaCl, 300 mM imidazole, 4 mM β-mercaptoethanol). Protein concentration was measured with a NanoPhotometer NP80 (Implen) and TEV protease was added in a 1:75 ratio. Cleavage of the protein by TEV results in two additional residues, namely Gly-Ala, at the *N*-terminus. The DNMT2Δ47-protease mix was dialyzed overnight against wash buffer using a 6–8 kDa dialysis tube (Spectra/Por). The sample was applied to a reverse Ni-IMAC using 2 mL Ni-NTA Agarose which was washed with 5 CV wash buffer. The flow-through and wash fractions were pooled, concentrated with an Amicon Ultra concentrator (10 kDa cutoff, Merck KGaA) to 2 mL and loaded for further purification by SEC (size-exclusion chromatography) on a Superdex 75 16/600 gel filtration column (Cytiva) in wash buffer. The peak fractions were pooled, incubated with compound **3** in a 2-fold molar excess, and concentrated with an Amicon Ultra concentrator (10 kDa cutoff, Merck KGaA) to 10 mg/mL for subsequent crystallization.

### Analytical size-exclusion chromatography of DNMT2Δ47 with compound 3

To assess the oligomeric state of DNMT2Δ47 upon compound **3** binding, *in vitro* analytical SEC runs were performed on a Superdex 200 Increase 3.2/300 (Cytiva). For the first run, the apo-protein was injected at a concentration of 10 mg/mL, for the second run it was pre-incubated with a two-fold molar excess of compound **3**. The resulting chromatograms were analyzed by superposing their 280 nm signals using the Unicorn7 evaluation software (Cytiva).

### Crystallization, data collection, model building, and refinement

Crystals were grown at 291 K using sitting drop vapor diffusion. The crystallization reservoir was composed of 0.2 M MgCl_2_, 0.1 M Tris, pH 8.5, and 12% PEG-8000. Crystallization drops contained 300 nL reservoir solution and 300 nL concentrated protein. Needle-shaped crystals grew within three days. Crystals were frozen in liquid nitrogen using glycerol as the cryo-protectant. Data were collected at ESRF beamline ID30B at cryogenic conditions, integrated using XDS (47), and scaled using AIMLESS (48) as part of the CCP4 software package.(49) Phases were obtained by molecular replacement with PHASER (50) implemented in the Phenix package (51) using the structure of human DNMT2 ((52); PDB 1G55) as the search model. The resulting data set has space group P2_1_2_1_2_1_, a maximum resolution of 2.60 Å, and contains two DNMT2 molecules per asymmetric unit. Compound **3** was parametrized from SMILES description using phenix.elbow.(53) Iterative model building and refinement were performed with Coot (54) and phenix.refine.(55) C24, C79, and C287 side chains show additional electron density in both chains and therefore were built as *S*-hydroxycysteine residues (CSO). The quality of the resulting structural models was analyzed with MolProbity.(56) Structure figures were prepared with PyMol 2.5.7 (The PyMol Molecular Graphics System, Schrödinger LLC.). Hydrophobicity of the binding site was visualized by a molecular surface representation colored from high (white) to low (green) hydrophobicity according to the Eisenberg hydrophobicity scale.(57) The electrostatic potential was calculated and visualized using the PyMol APBS plugin.(58) Crystallographic data are summarized in SI Table 3. Coordinates and structure factors are deposited at the Protein Data Bank PDB under accession code 9HGM.

### SeeSAR analysis

BiosolveIT SeeSAR (v13.0.1) was used to visualize the estimated binding affinity contributions of the co-crystallized DNMT2 ligand. SeeSAR concedes a semi-quantitative calculation of the thermodynamics using the HYDE algorithm and was used to evaluate torsions, desolvation, and interaction contributions for each ligand’s heavy atom.(59) For this, the DNMT2Δ47-binding compound **3** complex (PDB: 9HGM) was imported to SeeSAR and evaluated using the “Analyzer” functionalities.

### Pull-Down DNMT2 interaction assay

For immobilization on streptavidin beads, DNMT2 was biotinylated using the EZ-Link™ Sulfo-NHS-LC-Biotinylation kit. A 20-fold excess of Sulfo-NHS-LC-Biotin (10 mM in water) was added to DNMT2 and incubated for 2 h on ice. To remove the excess biotin reagent, Zeba Spin desalting columns were equilibrated with 1× PBS buffer, and the reaction mixture was loaded and centrifuged for 2 min at 1000 g. The biotinylation level was estimated using a HABA assay included in the kit according to the manufacturer’s protocols. By this, 4.7 biotin molecules were found to be attached per DNMT2 molecule. To confirm that the biotinylation did not affect the binding competence of the SAH and drug binding pockets, MST measurements with FTAD as a fluorescent tracer were performed as described above.

For in vitro pull-down assays, biotinylated DNMT2 was immobilized on streptavidin-coated paramagnetic beads (GenScript Streptavidin MagBeads). For this, beads were washed with PBS buffer containing 0.05% triton X-100, and subsequently, DNMT2 (30 µg per 50 µL beads) was immobilized for 45 min at RT. As a negative control, beads were treated with a protein-free buffered solution. Beads were washed with PBS buffer, and for each sample, 5 µL beads were added to either 1.5 µg of recombinant NonO protein (Active Motif) or 3.0 µg of recombinant eIF4E protein (Cayman). Beads were incubated for 1 h at RT, and subsequently, the supernatant was removed. Afterward, 10 µL PBS-buffered elution solution (containing either: 100 µM sinefungin or 100 µM compound **16** or 10 µM tRNA) was added. The samples were incubated for 45 min at RT, and the supernatant was treated with 2 µL 5× Laemmli buffer. For analysis of the proteins still bound to the beads after elution treatment, 10 µL Laemmli buffer supplemented with 10 mM biotin was added to the residual beads. All samples were heated to 95 °C for 10 min and analyzed via SDS-PAGE. Fluorometric protein detection was conducted using ProteOrange staining (Lumiprobe) according to the manufacturer’s protocols. Protein bands were visualized using a GE Healthcare Typhoon Trio biomolecular scanner, with laser excitation at 488 nm and fluorescence detection at 580 nm.

### tRNA modification analysis by LC/MS

MOLM-13 cells were seeded at a density of 5×10^6^ cells in 15 mL RPMI-1640 medium (supplemented with 10% FBS) containing DNMT2 inhibitor **16** (5 µM and 50 µM), or DMSO as mock treatment. Cells were incubated for 72 h at 37 °C. Post-incubation, cells were harvested and washed with PBS. Cell pellets were lysed in 1.5 mL TRIzol, mixed thoroughly by pipetting, and incubated for 5 min at RT. Subsequently, 150 µL chloroform was added, vortexed, and incubated for 10 min at RT. Samples were centrifuged at 12,000 g for 15 min at 4 °C. The upper aqueous layer was transferred to new tubes and mixed with 750 µL isopropanol, vortexed, and incubated for 10 min at RT. Following another centrifugation at 12,000 g for 15 min at 4 °C, the supernatant was discarded. Pellets were washed with 750 µL of 75% ethanol, vortexed, and centrifuged again at 12,000 g for 15 min at 4 °C. After discarding the supernatant, the pellets were air-dried at 37 °C and dissolved in 50 µL nuclease-free water. Total RNA samples were further purified by New England Biolabs Monarch RNA Cleanup (500 µg) kits according to the manufacturer’s instructions (protocol for RNA size fractionation). Small RNA molecules (<100 nt) containing total tRNA were eluted in 50 µL nuclease-free water, and concentrations were determined using a Nanodrop spectrophotometer.

To enrich the DNMT2 tRNA substrates, a literature-known hybridization-based protocol for the purification of thermostable tRNA was adopted.(60) Therefore, three biotin-labeled anti-sense DNA oligonucleotides suitable for tRNA pull-down were designed and purchased from GenScript. Sequences complementary to tRNA^Asp-GUC^: 5’-GTGACAGGCGGGGATACTCACCACT-Biotin-3’, tRNA^Gly-GCC^: 5’-CGTGGCAGGCGAGAATTCTACCACT-Biotin-3’, tRNA^Val-AAC^: 5’-CGTGTTAGGCGAACGTGATAACCACT-Biotin-3’. Next, GenScript streptavidin MagBeads (200 µL) were washed thrice with 1 mL TBST. Biotin-labeled DNA-oligonucleotide solutions were prepared at 7.5 µM in 3×400 µL TBST. MagBeads were incubated with 400 µL of each Biotin-oligo for 10 min at RT, then washed with TBST. Unbound oligos were measured by Nanodrop to confirm loading, assuming a theoretical capacity of 3 nmol per 200 µL beads. MagBeads were pooled in 1200 µL TBST and distributed into 12 PCR tubes with 100 µL TBST each. Total tRNA samples (100 µL) were mixed with 100 µL hybridization buffer (20 mM Tris, pH 7.6, 1.8 M NaCl, 0.2 mM EDTA) and added to the prepared MagBeads. The mixture was heated RT to 75 °C over 40 min using a PCR cycler, then cooled to 4 °C. Following centrifugation, supernatants were collected and saved. The beads were washed repeatedly with TBST at 4 °C until the absorbance at 260 nm was less than 0.01. tRNA was eluted by heating the beads in 30 µL Tris buffer (10 mM Tris, pH 7.6) at 75 °C, with the supernatant collected after each elution step, repeated three times. tRNA concentration and purity were determined using a Nanodrop spectrophotometer and 10% denaturing PAGE, respectively.

100 ng of enriched tRNA per sample was enzymatically digested into nucleosides using a mixture of 0.6 U nuclease P1 from *Penicillium citrinum* (Sigma-Aldrich), 0.2 U snake venom phosphodiesterase from *Crotalus adamanteus* (Worthington), 0.2 U bovine intestine phosphatase (Sigma-Aldrich), 10 U benzonase (Sigma-Aldrich), along with 200 ng of Pentostatin (Sigma-Aldrich). This reaction was conducted for two hours at 37 °C in a buffer containing 5 mM Tris (pH 8) and 1 mM magnesium chloride.

Following digestion, the nucleosides were spiked with an internal standard of ^13^C stable isotope-labeled nucleosides derived from *Saccharomyces cerevisiae* and analyzed. 100 ng of the digested tRNA and 25 ng of the internal standard were analyzed using LC/MS (Agilent 1260 series coupled with an Agilent 6460 Triple Quadrupole mass spectrometer featuring an electrospray ion source). The mobile phase consisted of a 5 mM ammonium acetate buffer (pH 5.3) and LC/MS grade acetonitrile (VWR). The elution process began with 100% ammonium acetate buffer at a flow rate of 0.35 mL/min, transitioning to a linear gradient that reached 8% acetonitrile at 10 minutes and 40% at 20 minutes. The initial conditions were restored to 100% ammonium acetate buffer for an additional 10 minutes.

A Synergi Fusion column (4 µm particle size, 80 Å pore size, 250 × 2.0 mm; Phenomenex) was utilized for the separation. The UV signal at 254 nm was monitored using a multiple wavelength detector (MWD) to track the primary nucleosides. The ESI parameters were set as follows: gas temperature at 350 °C, gas flow at 8 L/min, nebulizer pressure at 50 psi, sheath gas temperature at 350 °C, sheath gas flow at 12 L/min, and capillary voltage at 3000 V. The mass spectrometer operated in positive ion mode, employing Agilent MassHunter software in dynamic MRM (multiple reaction monitoring) mode for analysis. Quantification was achieved through a combination of external and internal calibration methods, as outlined in Kellner et al.(61)

### Cell viability assays

Cell viability assays were conducted with MOLM-13 cells and HCT MTase mutant cell lines (WT, knock-down DNMT2, and knock-down NSUN2) to investigate the DNMT2 inhibiting compounds cellular effect. Cells were cultured according to standard protocols MOLM-13 (RPMI, 10% FBS, 1x Penicillin/Streptomycin, 37 °C, 5% CO_2_) and HCT (McCoys’s 5A Medium, 10% FBS, L-Glutamine, 1x Penicillin/Streptomycin, 37 °C, 5% CO_2_). Cell viability was assessed using Promega Cell Titer Glo assay kit (MOLM-13) or Lonza ViaLight™ Plus Cell Proliferation and Cytotoxicity BioAssay Kit (HCT cell lines). Briefly, 1000 cells were seeded in white Greiner half-area 96-well plates and incubated for 48 h with inhibitors/ligands from DMSO stocks (final: 0.1% DMSO) under standard cultivation conditions. Then, the cell titer reagent was added to each well according to the manufacturer’s protocols, and luminescence was measured using a Tecan Spark 10M plate reader (MOLM-13) or Molecular Devices FlexStation (HCT cell lines).

### Inhibitors and probes synthesis

Detailed chemical synthesis procedures can be found in the SI.

## RESULTS

### DELopen screening reveals novel DNMT2 ligand chemotypes

Recombinant human DNMT2 was screened against the fourth generation DELopen (WuXi AppTec) library (4.2 billion compounds, 27 sub-libraries) to identify new small molecule RNA methyltransferase ligands (Figure 1A). DEL pannings were carried out with His-tagged DNMT2 protein stably immobilized on nickel chelate (Tris-NTA) affinity resin (SI Figure 1B). Selection screens against Tris-NTA resin without protein loading served as a nonspecific binding control. Three successive panning rounds were conducted, while the second round was chosen for deep sequencing based on the total retained DNA conjugates (SI Figure 1A). Sequenced hits were decoded and filtered to remove commonly encountered false positives and nonspecific binders to the affinity resin.(62) After affinity enrichment, 1713 unique full-length molecules were identified across three positive selection conditions and their combinations: (**A**) native DNMT2 without additives (n = 28 compounds); (**B**) DNMT2 with SAH (competitor, 100 µM, n = 104 compounds); (**C**) DNMT2 with tRNA^Asp^ (competitor, 10 µM, n = 207 compounds); (**AB**) n = 159 compounds, (**AC**) n = 217 compounds, (**BC**) n = 251 compounds, (**ABC**) n = 747 compounds (Figure 1B).

**Figure 1:**
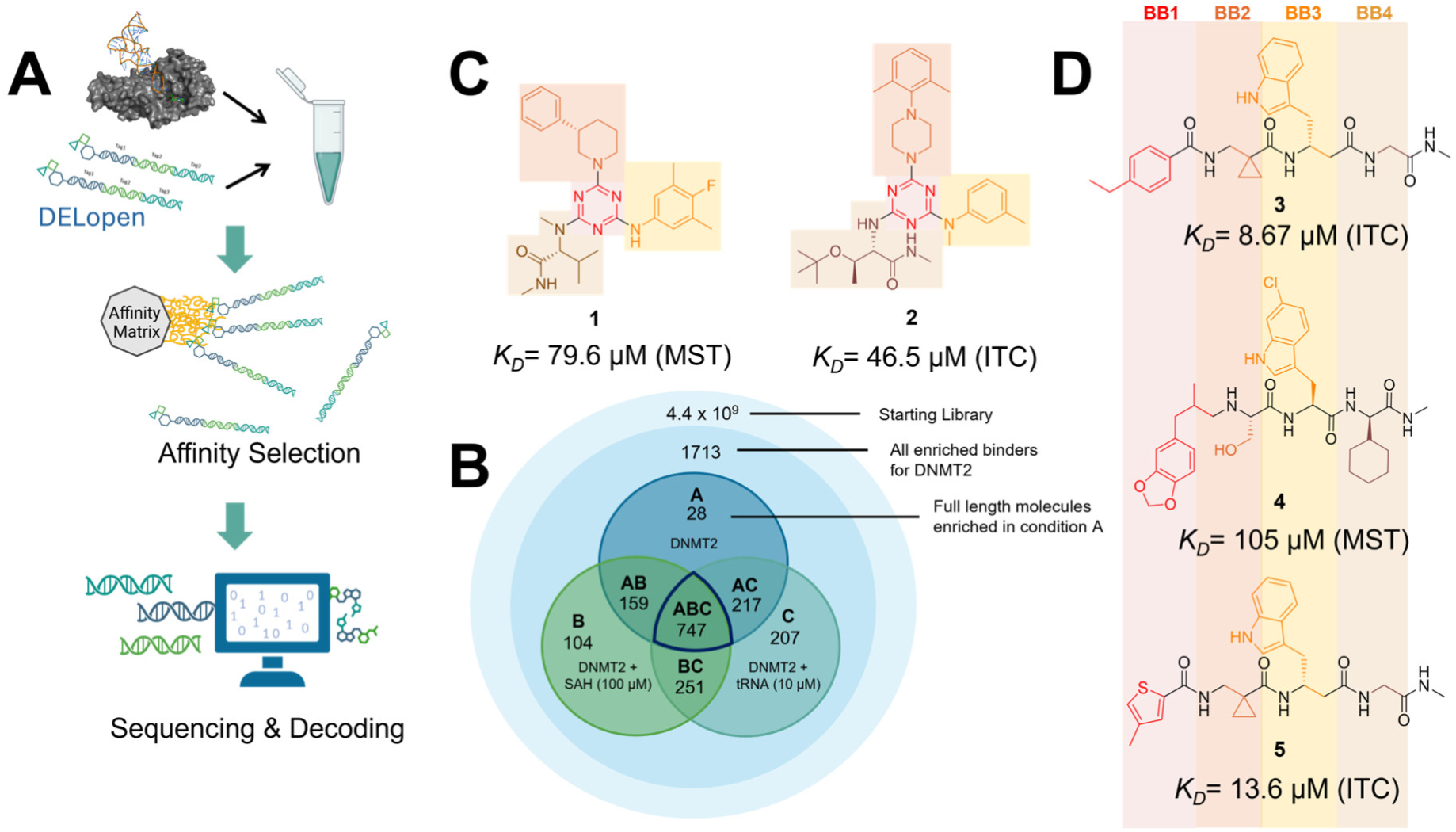
(**A**) DELopen screening workflow. DELopen libraries consist of 4.2B compounds encoded by individual DNA barcodes. During affinity selection, small molecule-DNA-conjugates are enriched, while non-binders are washed away. The pipeline’s last step is the sequencing and decoding of new ligand structures. (**B**) Conditions used during the DELopen screening: In condition A native DNMT2 without additives was used; condition B: DNMT2 and SAH (100 µM) were mixed to block the SAH binding site; condition C: DNMT2 was incubated with tRNA^Asp^ (10 µM). Most full-length molecules were enriched in condition ABC (unaffected by competitors), from which 5 selected compounds **1**–**5** were chosen for off-DNA synthesis and downstream characterization. (**C**) Enriched triazine-based compounds selected from ABC. (**D**) Peptidomimetic compounds enriched in ABC and chosen for further study. Building blocks BB1–BB4 are highlighted accordingly.

These post-DEL screening compounds had enrichment scores of at least 100 for DNMT2. The enrichment score was calculated by decoding the DNA tags from next-generation sequencing (NGS) results, considering the corrected copy number, library size, sequencing depth, and other normalization factors (see https://hits.wuxiapptec.com/delopen).(37) A molecule with an enrichment score of 100 is a hundred times more abundant in the sequencing data than the average molecule in the library. Interestingly, DEL sequencing analysis showed that most of the compounds enriched on DNMT2 were found within the condition **ABC**, which may reflect the inability of the low-affinity SAH competitor to completely occlude the active binding site during the panning. The five compounds with the highest confidence (enrichment scores > 200 000 and copy number > 10, SI Excel Table) and structural diversity (from three sub-libraries in total, see SI Figure 1F) were selected for further investigation and downstream validation studies (Figure 1C, D).

Across the 747 **ABC**-enriched and the five highest-scoring representative molecules (“hits”), recurring patterns were apparent in each of the four building block positions (denoted BB1, BB2, BB3, and BB4), indicating structurally related families: two triazine-based DEL hits (**1**–**2**) and three peptide-based DEL hits (**3**–**5**). Hit molecules **3**–**5** can be described as β-homo-tripeptides with non-natural side chain groups in building block positions BB2 resp. BB3 and incorporating an *N*-terminal substituent in BB1. The BB4 *C*-terminus (corresponding to the DNA tag attachment position) was synthetically amidated with methylamine. These hit compounds were resynthesized without the encoding DNA tag and were studied for DNMT2 active site binding using an MST displacement assay as previously described by Zimmermann et al.(38) For the thermophoretic detection, 5-FAM-triazolyl-adenosyl-diaminobutyric acid (FTAD) was used as a fluorescent probe, which can bind to the DNMT2 SAH binding pocket.(38) When FTAD is bound to DNMT2 versus its unbound state, there is a measurable difference in the F_norm_ value represented by a distinct positive MST shift, and thus, a ligand that can displace the FTAD-DNMT2 interaction can be detected by competitive MST screening assays (SI Figure 3).

In this regard, two of the five tested DEL hit compounds, **3** and **5**, induced a quantitative FTAD displacement in the MST assay (Figure 2A). In subsequent dose-response experiments, **3** and **5** showed *K_D_*-values of 48.2 µM and 241 µM, respectively (detailed information in Table 1). Interestingly, **3** and **5** are similar in structure, differing only by the first building block, 4-ethyl benzoic acid for **3** and 4-methylthiophene-2-carboxylic acid for **5** (Figure 1D), while compound **4** features a similar peptidomimetic structure but without β-homo amino acids. In contrast, **1** and **2** share a common triazine core structure, due to their origin from the same sub-library (Figure 1C). Notably, compounds **1** and **4** did not lead to FTAD displacement but rather induced a negative MST shift towards a higher FTAD-bound degree thermophoresis resp. strengthened interaction. We speculated about an allosteric binding mode of these compounds which agrees with the observation that these compounds were identified in the **ABC** panning cluster. Yet, only compound **2** did not lead to a significant thermophoresis behavior alteration even at millimolar concentrations, and hence, this compound was assayed by the alternative nanoDSF method, where **2** revealed an apparent *K_D_*-value of 2.98 µM due to thermal DNMT2 stabilization (Figure 2C). As an orthogonal biophysical binding experiment, we implemented isothermal titration calorimetry (ITC) ligand binding studies. In these experiments, **2**, **3**, and **5** showed significant binding enthalpy signals when titrated to DNMT2, with *K_D_*-values of 69.8, 8.67, and 13.6 µM, respectively (Figure 2B). Of note, **3** resulted in particularly long injection equilibration times in the minute magnitude, and thus, we speculated about a conformational change upon ligand binding which could be confirmed by subsequent crystallography as described below. Additionally, an orthogonal and label-free affinity selection-mass spectrometry (AS-MS) assay for lead compound **3** was conducted, revealing a *K_D_*-value of 8.40 µM, which is in agreement with the ITC and MST data.(41)

**Figure 2:**
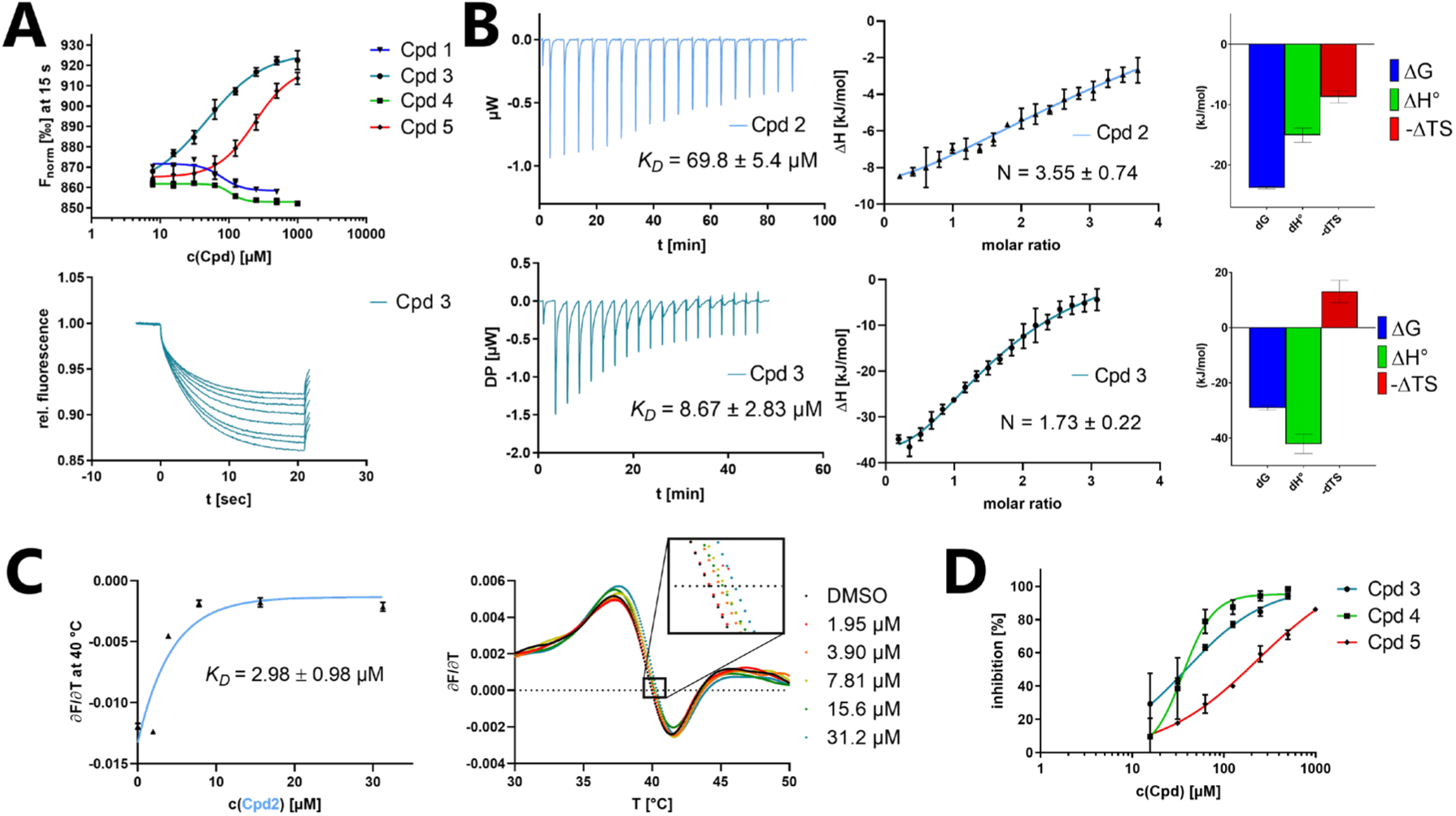
Selected biophysical and enzymatic characterizations of hit compounds **1**–**5** binding to DNMT2. (**A**) MST dose-response curves for compounds **1**, **3**, **4**, and **5** with detailed MST traces for compound **3**. (**B**) ITC data of **2** and **3** titrated to DNMT2. From left to right: Titration thermogram, stoichiometry plot, and signature plot. (**C**) nanoDSF data of **2** titrated to DNMT2. (**D**) DNMT2 enzyme inhibition results from enzymatic ^3^H-incorporation assay for **3**, **4**, and **5**. A comprehensive overview of the biophysical data for DNMT2 binding characterization can be found in the SI.

**Table 1:** Biophysical data of compounds **1**–**5** determined by various ligand binding and enzyme inhibition experiments. All results include the mean value and standard deviations from at least technical triplicate measurements. Raw data and analysis plots are depicted in Figure 2 and SI Figures 2–7.

| Cpd | IC<sub>50</sub> | K<sub>D</sub> |  |  |  |
| --- | --- | --- | --- | --- | --- |
|  | <sup>3</sup>H assay | MST | ITC | NanoDSF | AS-MS |
| **1** | n.i. | 79.6 ± 10.0 μM | n.b. | n.d. | n.d. |
| **2** | n.i. | n.b. | 69.8 ± 5.4 μM | 2.98 ± 0.98 μM | n.d. |
| **3** | 38.9 ± 6.9 μM | 48.2 ± 8.0 μM | 8.67 ± 2.83 μM | n.d. | 8.40 ± 2.20 μM |
| **4** | 44.4 ± 12.9 μM | 105 ± 9 μM | n.b. | n.d. | n.d. |
| **5** | 236 ± 63 μM | 241 ± 36 μM | 13.6 ± 1.6 μM | n.d. | n.d. |

n.i. = no inhibition / less than 50% inhibition at 500 μM // n.b. = no enthalpy signal observable (ITC) / no thermophoresis shift observable (MST) // n.d. = not determined

Besides biophysical characterization utilizing DNMT2 binding assays, all five DEL hits were evaluated regarding their inhibitory effects using an enzymatic ^3^H-incorporation assay, where the tritium-labeled methyl group incorporation from ^3^H-SAM to the DNMT2 substrate tRNA^Asp^ can be determined.(3, 4) In Figure 2D, it is shown that **3**, **4**, and **5** were able to inhibit DNMT2’s MTase activity with IC_50_-values of 38.9, 44.4, and 236 µM, respectively. Data and figures of additional ligand characterization experiments can be found in SI Figures 2–7.

Taken together with the data gleaned from the DEL sequencing results, the combined ligand characterization data suggest that the DEL hits identified during this screening campaign are not binding to the DNMT2’s active site nor tRNA pockets, because hit compounds were enriched in condition **ABC** and show only partly competitive behavior during ligand binding experiments. The experimental findings from MST, ITC, nanoDSF, and enzymatic ^3^H-incorporation assay suggest that triazine compounds **1** and **2** act as non-competitive DNMT2 binders without inhibition potency, whereas β-homo-tripeptides **3**–**5** function as DNMT2 enzyme inhibitors. The mode of action resp. their binding site remained elusive from the biophysical investigations but could be finally solved by DNMT2 crystallography (vide infra). We speculated that the minor discrepancies between the enrichment scores’ relative order and determined affinities may be due to the DNA tag sequence absence in the off-DNA synthesized hit compounds.

For further characterization of both triazine- and tripeptide-based DEL hits, fluorophore functionalized tracer compounds **6** and **7** were developed based on their parent compounds for target selectivity determination in fluorescence-based assays and as molecular probes for further investigations (Figure 3A).

**Figure 3:**
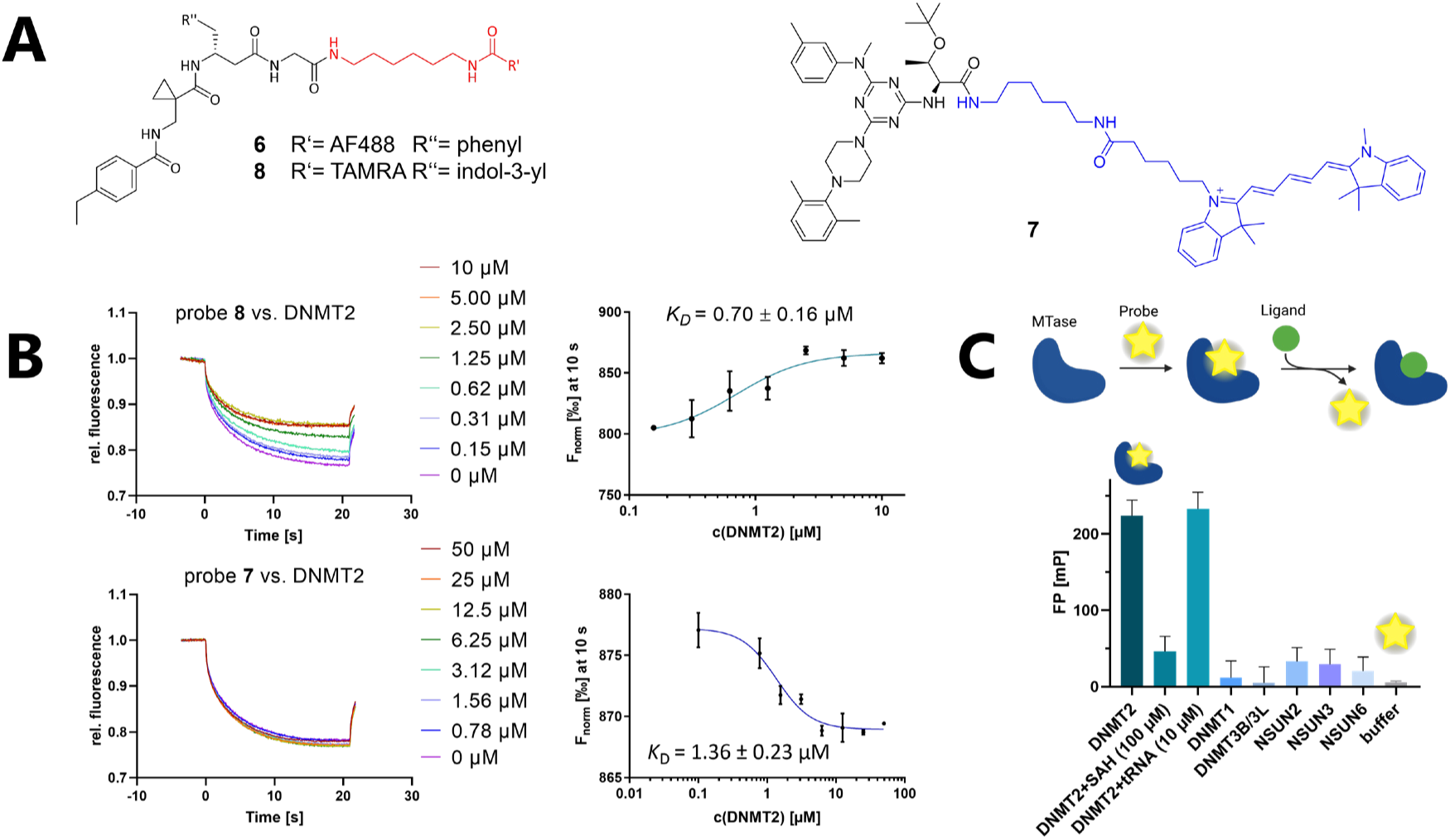
FP and MST experiments investigating DNMT2 and related MTases binding to fluorescence tracers/probes. (**A**) Chemical structures of synthesized probes. **6**: AF488-labeled β-homo-tripeptide. **7**: Cy5-labeled triazine probe. **8**: TAMRA-labeled β-homo-tripeptide. (**B**) Top: MST dose-response curves for probe **8** binding to DNMT2 and below analogous MST curves for probe **7**. (**C**) Fluorescence polarization assay for probe **6** binding to native DNMT2 (2 µM), DNMT2 (2 µM) preincubated with SAH (100 µM), DNMT2 (2 µM) with tRNA^Asp^ (10 µM), and various related MTase off-targets (all 2 µM).

Three fluorescent probes were designed during this study for both identified DEL chemotypes: AlexaFluor 488 (AF488) resp. tetramethylrhodamine (TAMRA) labeled β-homo-tripeptides (**6** and **8**), and a Cy5-labeled triazine-based probe (**7**). Apparent binding affinities of fluorescent probes were measured by MST experiments as described above with the modification that the respective probe was used instead of FTAD (Figure 3B and SI Figure 3E).(40) By this, probe binding affinities (**6**: 1.65 µM, **7**: 1.36 µM, and **8**: 0.70 µM) were determined which resemble the original DEL hit compounds’ **2** and **3** *K_D_*-values, suggesting that the chosen fluorophores do not significantly interfere with their binding behavior and the probes are suitable for fluorometric MTase investigations.

Next, peptidomimetic probe **6** was utilized to determine MTase binding selectivity towards related DNMT resp. NSUN proteins, and to investigate the competitivity of natural DNMT2 substrates (SAH and tRNA^Asp^) by fluorescence polarization (FP) displacement experiments (Figure 3C). Fluorometric assays using DNMT2 revealed that the compound **6**-DNMT2 complex yields high fluorescence polarization (220 mP) compared to enzyme-free buffer conditions (5 mP). By examining whether this effect can be reversed by the addition of natural ligands, we found that SAH (100 µM) is able to displace probe **6** while tRNA^Asp^ (10 µM) does not show a significant effect and high fluorescence polarization is retained during the tRNA substrate addition. These results were furthermore confirmed by an ITC displacement experiment, where **3** was able to displaceable SAH and vice versa (SI Figure 4F).(63) Analogous MST displacement experiments with in situ fluorescently labeled tRNA^Asp^ showed consistently that none of the hit compounds **1**–**5** is able to dissociate the nucleic acid-protein complex formed by DNMT2 and tRNA^Asp^ (*K_D_* = 210 nM, SI Figure 8A,B), and furthermore, no DEL compound (100 µM) had a significant influence on the thermophoresis of tRNA^Asp^ in protein-free solution (SI Figure 8C,D).(39)

Probe **6** also enabled the determination of ligand binding selectivity towards other m^5^C MTases using the FP assay setup. Since NSUN2, NSUN3, and NSUN6 introduce m^5^C in other tRNA and DNMT1 resp. DNMT3A/3L in DNA, these enzymes were selected for the ligand selectivity profile determination. By incubating probe **6** with various related m^5^C MTases (final concentration: 2 µM), it was observed, that this probe does not bind to related NSUN and DNMT families’ MTase representatives, highlighting its unique DNMT2 binding selectivity (Figure 3C).

### Crystallography unravels an allosteric binding pocket of hit compound 3

For our crystallographic studies, we used a construct of human DNMT2, dubbed DNMT2Δ47, in which residues 191 to 237 were deleted. A similar construct was already utilized for the X-ray structure of human DNMT2 with bound SAH ((52); PDB 1G55). We could obtain an X-ray structure of human DNMT2Δ47 with bound compound **3** at 2.6 Å resolution which shows an overall fold similar to DNMT2Δ47-SAH as indicated by an overall rmsd(C_α_) of 0.69 Å (SI Figure 17A). The asymmetric unit contains two very similar DNMT2Δ47-compound **3** complexes with an overall rmsd(C_α_) of 0.354 Å and largest differences in the loop region comprising residues 82 to 95 (Figure 4A). A large common surface area of 1685 Å^2^ is present between these two protein complexes. Interface and packing analysis with PISA (64) supports a stable dimer formed by the two complexes in the asymmetric unit. Therefore, we used analytical size exclusion chromatography to investigate the oligomeric state of DNMT2Δ47-compound **3** in solution. We could show that DNMT2Δ47 with bound compound **3** is indeed dimeric in solution at a protein concentration of 10 mg/mL (SI Figure 18), which was also used for crystallization. In contrast, DNMT2Δ47 without ligand is monomeric in solution under the same conditions.

**Figure 4:**
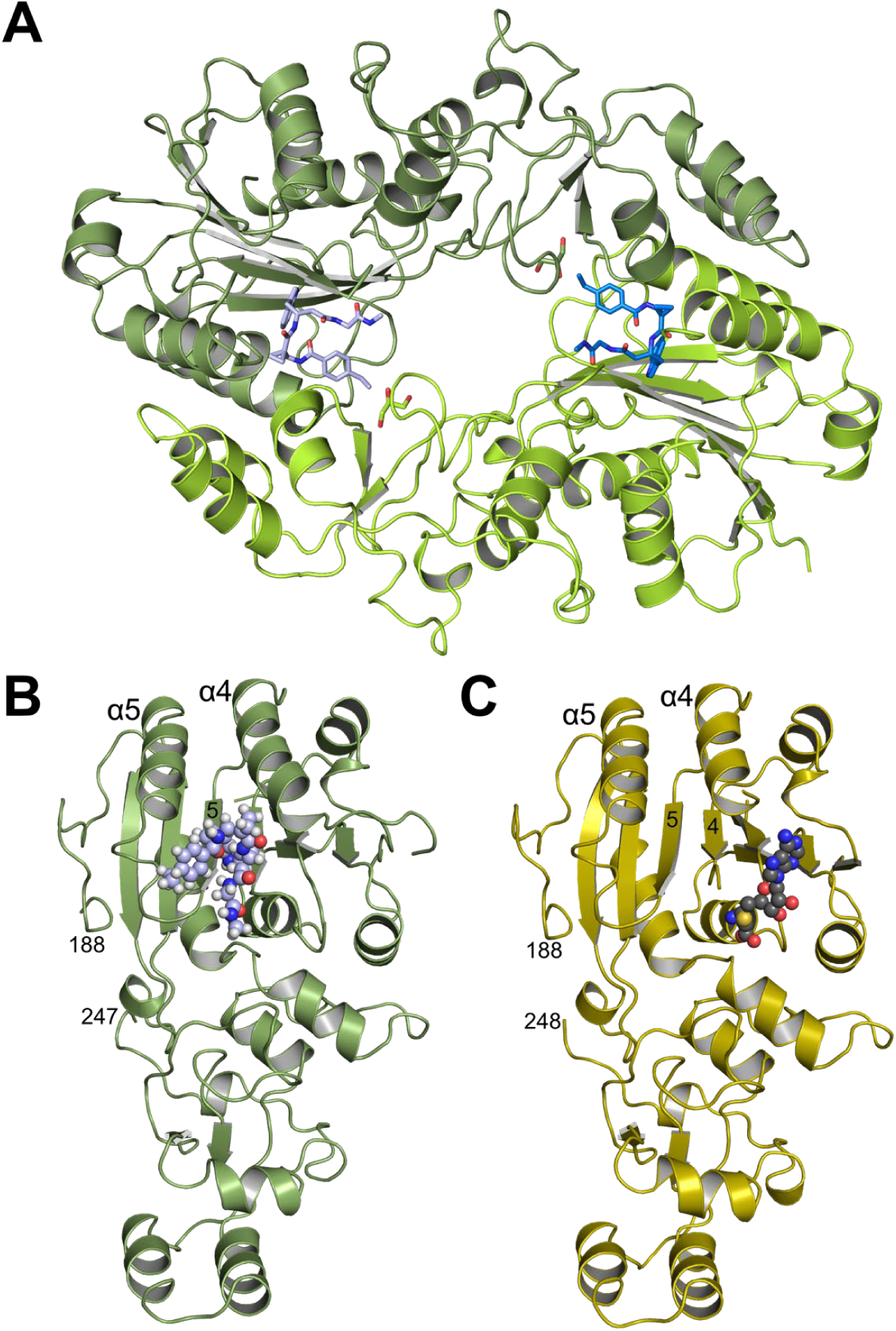
(**A**) X-ray structure of DNMT2Δ47 showing both molecules of the asymmetric unit. Bound compound **3** molecules are represented as sticks. Chain A is shown in green/light blue and chain B in light green/blue. (**B**) X-ray structure of DNMT2Δ47-compound **3** chain A depicted as cartoon (green) with compound **3** as spheres (carbon atoms in light blue). (**C**) The crystal structure of human DNMT2Δ47 (PDB 1G55) as cartoon (yellow) with bound SAH as spheres (carbon atoms in grey) is given for comparison.

Both complexes in the asymmetric unit show clear electron density for compound **3** (SI Figure 17C). Most interestingly, compound **3** is bound in a largely hydrophobic pocket adjacent to the active site (Figure 4B). For comparison, the structure with SAH bound in the active site is depicted in Figure 4C and superpositions of the two different complexes without and with ligands are given in supplemental SI Figures 17A and 17B.

Compound **3** is bound by residues mainly from the two regions β4-α4 and β5-α5, which is depicted in Figure 5A for chain A (interacting residues of chain B have been omitted for clarity). A comprehensive schematic ligand-protein interaction diagram was created with LigPlot+ (65) for both DNMT2Δ47-compound **3** complexes in the asymmetric unit (SI Figure 19). In the following, a detailed description of compound **3** binding to chain A and the differences between both complexes in the asymmetric unit are given. The indole ring of compound **3** is located in a hydrophobic cave formed by Y10, M75, backbone of S76, P77, F99, L118, F124, and L133. Additionally, the amine group of the indole ring forms a hydrogen bond with the backbone carbonyl of E119. The ethylbenzene moiety of compound **3** is bound between V121, F124, S127, T129 and L279_B_. The ethyl group is also in contact with a glycerol molecule (used for cryo-protection) bound in the dimer interface. The amide group, which connects the ethylbenzene and cyclopropane moieties, is in van der Waals contact with P78 and R95 and forms three hydrogen bonds, namely with the main chain carbonyl of L278_B_, with the side chain of N120 and an intramolecular H-bond to the amino group connecting the cyclopropane and indole moieties. The cyclopropane moiety is surrounded by the hydrophobic side chains of F99, L278_B_, and L279_B_. Finally, the two terminal amide groups of compound **3** form hydrogen bonds with the side chains of N120 and N375. Due to different side chain conformations in chain B, R95_B_ forms an additional hydrogen bond to this part of compound **3**_B_, and N281_A_ and E302_A_ are in non-polar contact with compound **3**_B_.

**Figure 5:**
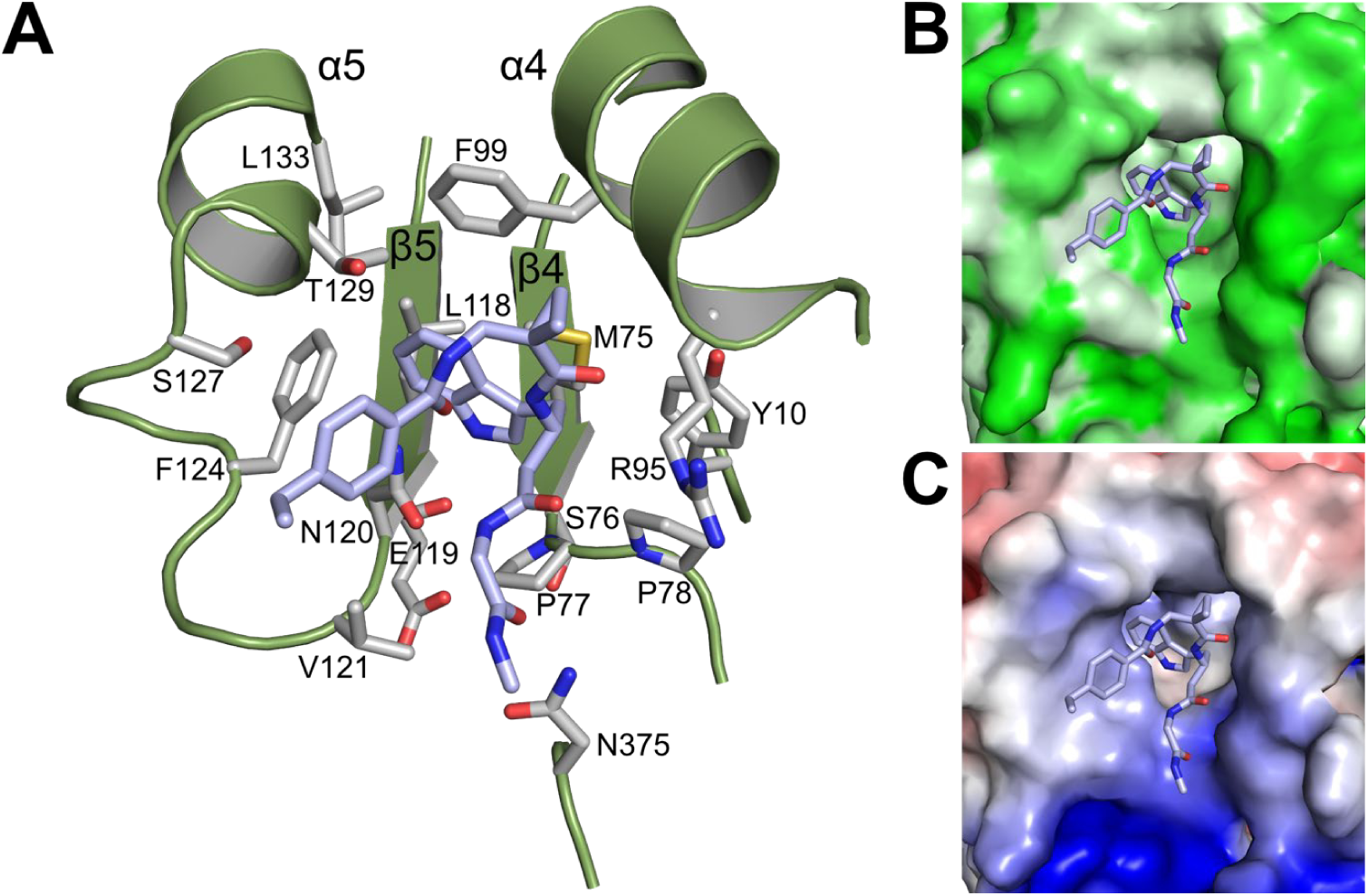
(**A**) A detailed view of compound **3** (carbon atoms in light blue) bound to DNMT2Δ47 chain A (green cartoon) with interacting protein side chains in light grey. Side chains of chain B are omitted for clarity. Molecular surface representation of the binding site with compound 3 colored by (**B**) hydrophobicity (white -> green = high -> low) and (**C**) electrostatic potential from -5 kT/e (red) to +5 kT/e (blue).

The molecular surface representations of the ligand binding site colored by hydrophobicity (Figure 5B) and electrostatic potential (Figure 5C) show a clear correlation with the properties of the individual moieties of the bound ligand, e.g., the binding of the ethylbenzene moiety to a hydrophobic patch and the interaction of the carbonyl groups with a highly positive patch.

In order to analyze the structural rearrangements induced by compound **3** upon binding to DNMT2Δ47, we created a local superposition with DNMT2Δ47-SAH onto the central β-sheet of the large domain, namely onto β3 to β7. As depicted in Figure 6, the largest local movement upon binding of compound **3** is observed in the loop following β4 (M75 to P78), where P77 and P78 are rotated and shifted towards the SAH binding site. Additionally, subsequent residues could be built in both complexes with compound **3** (A: C79-F82; B: C79-I87), whereas in DNMT2Δ47-SAH residues 79 to 96 are not resolved. Binding of compound **3** shifts residues into the SAH binding pocket. Moreover, the active site residue C79 is now resolved and moved into the SAH binding region. The superposition depicted in Figure 6 shows that binding of compound **3** narrows the active site which prohibits SAH binding due to steric hindrance by P77, P78, and C79. As a consequence, compound **3** acts as an allosteric inhibitor by reshaping the active site.

**Figure 6:**
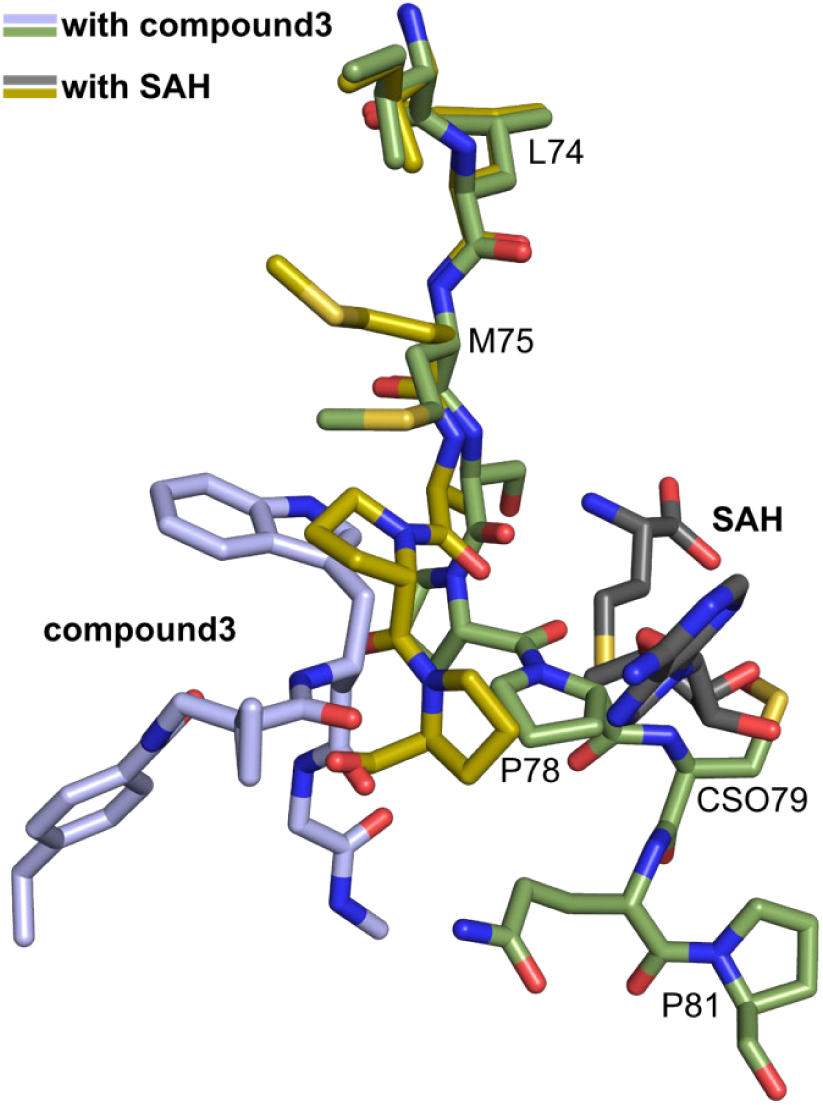
Local superposition onto the central β-sheet in the large domain of DNMT2Δ47-compound **3** (green/light blue) with DNMT2Δ47-SAH (yellow/grey) showing the end of β4 and following residues with respective ligands. Residues P77, P78, and C79 are shifted towards the SAH binding site upon compound **3** binding.

### Structure-activity study leads structural optimization of hit compound 3

As the next step, we aimed to increase the affinity of the DEL hits to generate a cellularly effective DNMT2 modulator. Compound **3** was chosen for structure-activity-relationship studies (SAR) due to its favorable binding characteristics observed in ITC, MST, and ^3^H-based enzyme assays. Its suitability for SAR investigation was enhanced by the feasibility of conducting solid-phase peptide synthesis (SPPS) to generate a diverse ligand analog set (syntheses procedures in the SI). A major advantage for compound **3** optimization is given by the elucidated crystal structure which allows rational considerations for improving ligand interactions in the new DNMT2 inhibitors design (Figure 5). First, a computational SeeSAR analysis of the ligand binding pose was performed which provided information on putative approaches for ligand optimization (Figure 7).(59)

**Figure 7:**
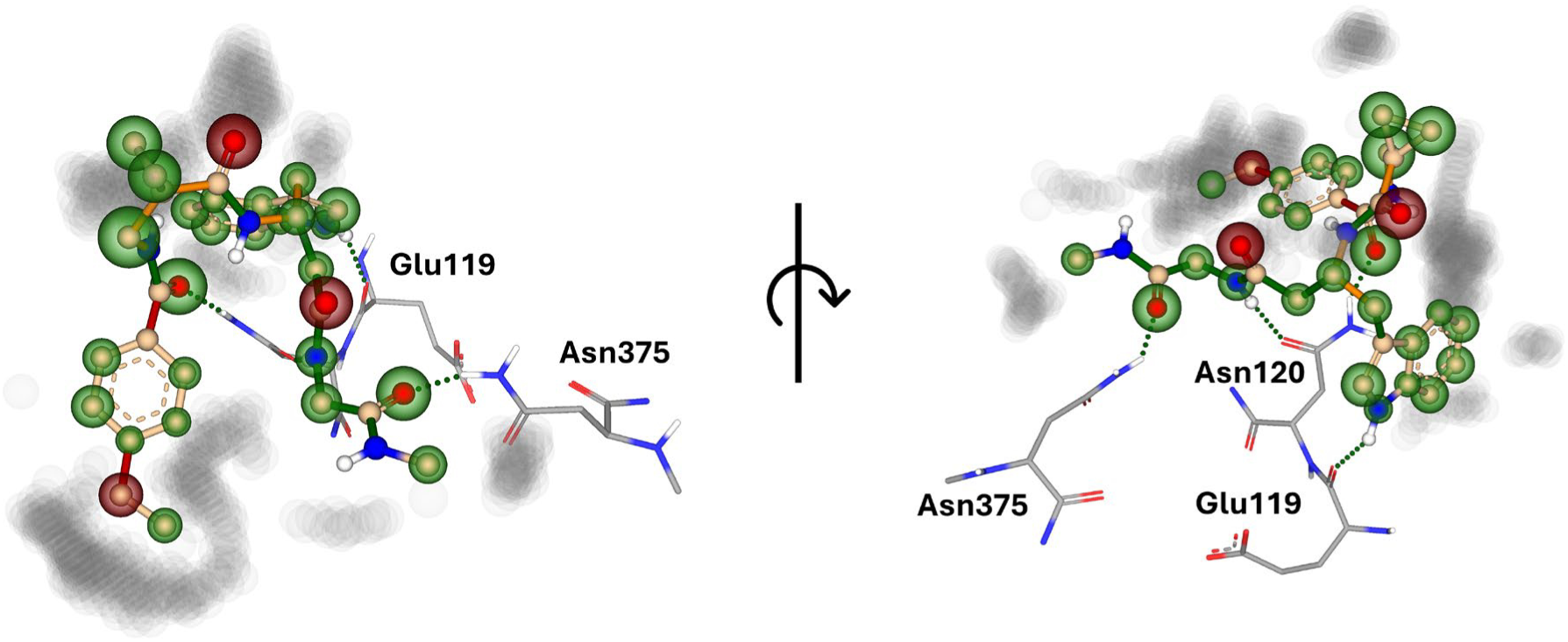
SeeSAR analysis by binding pose inspection of hit compound **3**. Hyde coloring enabled the identification of contributing resp. unfavorable atom placements (colored spheres; green: contributing, red: unfavored); the peptide backbone’s dihedral angles were evaluated for their torsional free energy (colored sticks; green: contributing, red: unfavored). Characteristic hydrogen bonds are formed with the protein residues Gln119, Asn120, and Asn375. Potentially druggable yet unoccupied space in the protein binding pocket is highlighted by grey shadings.

The following hypotheses on the ligand binding characteristics were derived from this computer-assisted analysis: Most atoms of ligand **3** are already optimally positioned for interaction with the DNMT2 protein (green spheres). A few atoms that are assumed to have an unfavorable effect on the DNMT2 interaction (red spheres of the BB2 and BB3 carbonyl oxygens) cannot be exchanged without the loss of the characteristic U-shaped ligand. In conclusion, we hypothesized that modification to the peptide backbone will lead to a significant loss of binding affinity. The peptide backbone’s dihedral angles are mostly in optimal shape for BB3 and BB4 but show minor potential for improvement in BB1 and BB2 which could favor a reduction of entropy penalty in the binding profile by rigidizing the ligand’s native binding conformation (see ITC results in Figure 2B). Interestingly, there are three regions in the ligand binding pose that allow ligand expansion (gray shading) and thus enable additional potentially beneficial interactions. These regions are localized 1.) between BB1 and the *C*-terminus which may allow ligand macrocyclization, 2.) in the homo-tryptophan residue 5-position, and 3.) near the BB2 cyclopropyl motif. These regions were to be investigated by substitutions and scaffold hopping strategies in the SAR study. Subsequently, a total of 40 derivatives derived from hit compound **3** were synthesized and screened for DNMT2 binding using the MST assay at a final ligand concentration of 100 µM. If significant binding to DNMT2 was measured (>25% FTAD displacement), the binding affinity was determined by following ITC experiments. For clarity, only a selection of the most potent DNMT2 ligands is shown in the main manuscript (Table 2). A comprehensive overview of compound and assay data can be found in SI Table 1.

**Table 2:**
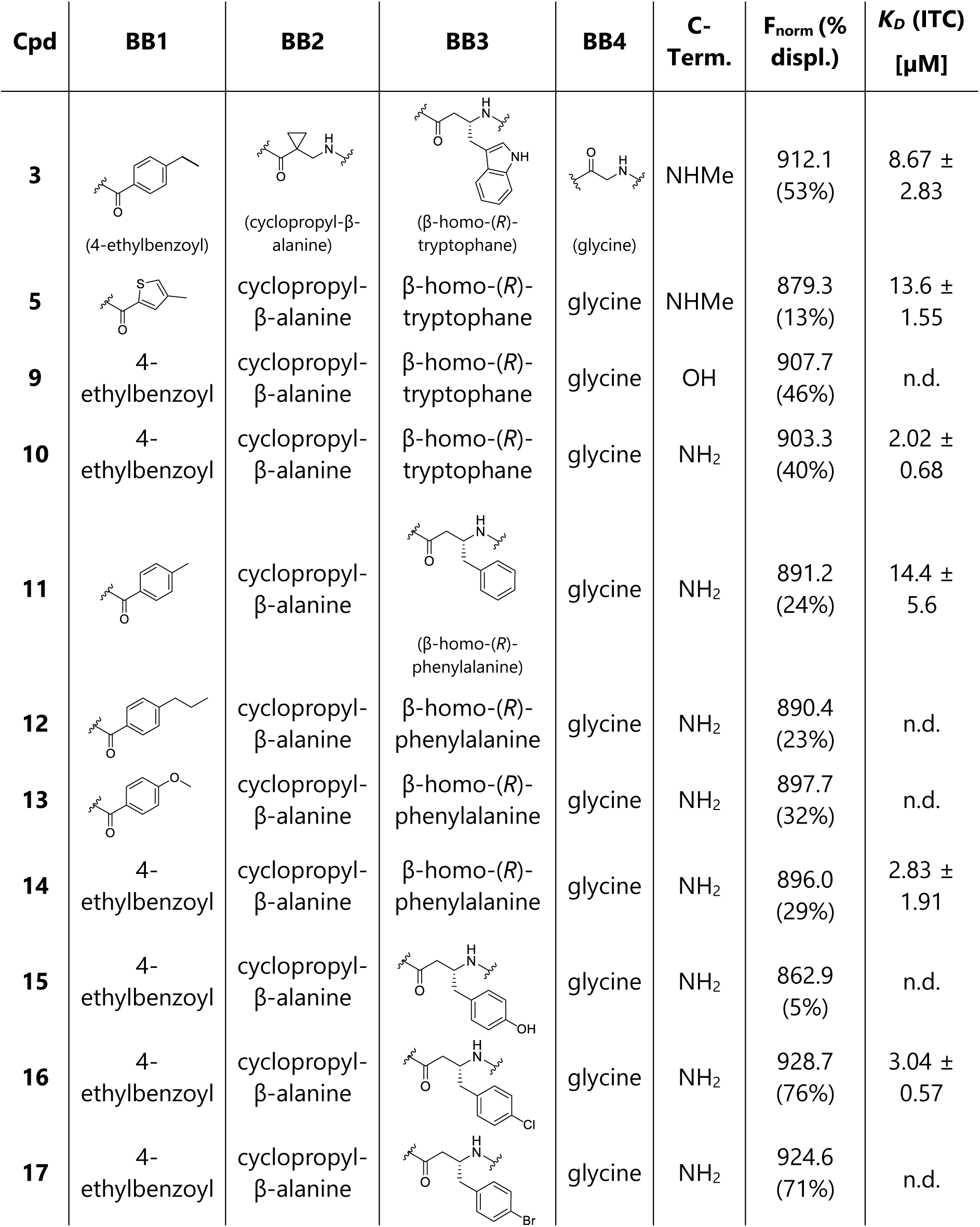
Overview of the eleven most potent β-homo-tripeptide derivatives as determined by MST experiments (F_norm_ and % FTAD displacement) and ITC assays. All results include the mean value and standard deviations from at least technical triplicate measurements. A table including all 40 synthesized analogs can be found in SI Table 1; n.d. = not determined.

During this SAR study all building block positions (BB1–BB4) were systematically varied: For the BB1 variation (4-ethylbenzoyl), it was observed that the alkyl chain at the phenyl ring’s *para* position was replaceable, still resulting in significant binding affinities. Notably, the presence of a (rigidized) alkyne group in this position resulted in the complete DNMT2 binding abolition. Furthermore, substitutions other than the *para* position, as well as, the aromatic moiety’s absence, failed to sustain ligand affinity. Next, we observed that any BB2 substitution, shortening, or decyclization (cyclopropyl-β-alanine) was also not tolerated, highlighting the critical necessity of this special β-amino acid. For the BB3 substitution (β-homo-tryptophan), all attempts to shorten the peptide backbone, altering the stereochemistry, and most tryptophan ring system replacements yielded no binding to DNMT2. However, replacement with the phenyl-containing analog (β-homo-(*R*)-phenylalanine, (**14**)) resulted in a significant binding (29% FTAD displacement). Subsequent synthesis of diverse *para*-substituted derivatives showed increased displacement values of up to 76% indicating improved binding affinity compared to parent compound **3**, which could be confirmed by subsequent dose-response MST and ITC experiments (*K_D_*(**16**) *=* 3.04±0.57 µM, Table 2, Figure 8). When substituting BB4 (glycine), extending the peptide backbone, or exchanging with either (*R*)- and (*S*)-alanine, BB4 derivates showed significantly lower DNMT2 affinities. Interestingly, amide group methylation between BB1 to BB4, aimed at potentially enhancing cellular permeability and stability, yielded no significant ligand affinity to DNMT2. As an exception, the peptide C-terminus methylation among other modifications (see also probes **6** and **8**) did not disturb the ligand affinity, and thus, represented a potential exit vector from the DNMT2 pocket suitable for ligand conjugation.

**Figure 8:**
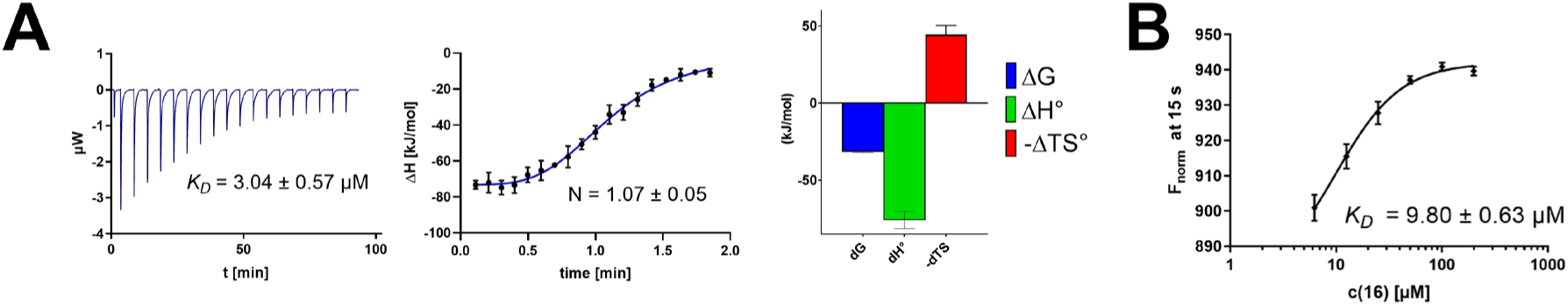
MST and ITC data of the SAR study’s most potent ligand. Compound **16** showed (**A**) a *K_D_* = 3.04±0.57 µM in ITC and (**B**) 9.80±0.63 µM in MST experiments.

In summary, our experimental SAR is a major confirmation of the SeeSAR hypotheses and agrees well with the crystal structure’s ligand pose. The BB3 replacement from β-homo-(*R*)-tryptophane to β-homo-(*R*)-(4-chloro)phenylalanine (**16**) led to an improvement in ligand affinity, while most other modifications resulted in similar or lower ligand affinities.

Since peptidomimetic drugs might be prone to cellular permeability issues, hit compounds **1**– **4** and the most potent ligands **10**, **14**, **16**, and **47** were tested for passive membrane transport using a parallel artificial membrane permeation assay (PAMPA). While compounds **1**–**4**, **10**, **14**, and **47** failed to pass through the artificial membrane, compound **16** showed moderate permeability (*P_app_* = 0.3 10^- 6^ cm/s) rendering this compound as the most promising lead structure of this SAR optimization. Detailed PAMPA results can be found in the supporting information (SI Table 2). Overall, compound **16** showed the best ligand binding properties, and it also demonstrated the ability for passive membrane transport in the PAMPA, making it the most promising candidate for cellular studies of DNMT2 inhibition.

### Cellular potency of allosteric inhibitors targeting DNMT2

Next, we aimed to investigate whether compound **16** can affect enzymatic DNMT2 activity in a cellular context and is thus able to reduce RNA modification contents in living systems. Previously, DNMT2^-/-^ mice were found to have reduced m^5^C levels in corresponding tRNAs (tRNA^Asp-GTC^, tRNA^Val-AAC^, and tRNA^Gly-GCC^), while only the double-knockout (DNMT2^-/-^ & NSUN2^-/-^) mutant resulted in a complete m^5^C elimination in tRNA substrates of DNMT2.(13) Further experiments utilizing siRNA-mediated DNMT2 knockdown and chemical inhibition with the pan-DNMT/NSUN inhibitor azacitidine in leukemia cells demonstrated a total C38 methylation loss.(66–68) Following these observations, we aimed to investigate whether **16** can reduce m^5^C modification levels in cells through isoacceptor-specific enrichment of DNMT2 tRNA substrates and modification analysis by means of an LC-MS/MS-based assay. In this regard, following previous studies investigating DNMT2’s role in leukemia cells,(67, 68) we treated MOLM-13 cells with **16** (5 and 50 µM) for 72 h, followed by the isolation of isoacceptor-specific tRNAs using biotinylated antisense DNA oligomers on magnetic beads. LC-MS/MS-based analysis of digested total tRNA isolates was used to determine relative changes in modification levels of 18 abundant tRNA modifications.(69)

As shown in Figure 9, levels of targeted tRNA modifications (m^5^C and m^5^Cm) were significantly reduced when MOLM-13 cells were treated with 50 µM of compound **16**. In this regard, the m^5^C reduction (∼25%) is in agreement with previous knock-down experiments highlighting that 75% of tRNA m^5^C are mediated by NSUNs.(13) Notably, treatment with 5 µM of compound **16** was found to be too low to stimulate any significant change in RNA modifications, probably due to its moderate cell permeability and proportionally lower cellular concentration. In addition, only minor other modifications changes were observed including mildly reduced pseudouridine and Um levels resp. slightly increased Am levels, highlighting our inhibitor chemotype’s DNMT2-selectivity. The origin of those off-target RNA modification alterations remains not entirely solved, but we speculate that RNA-modifying enzymes introducing those modifications are likely to be regulated by DNMT2-mediated m^5^C modification rendering them as hierarchical-type modifications.(70)

**Figure 9:**
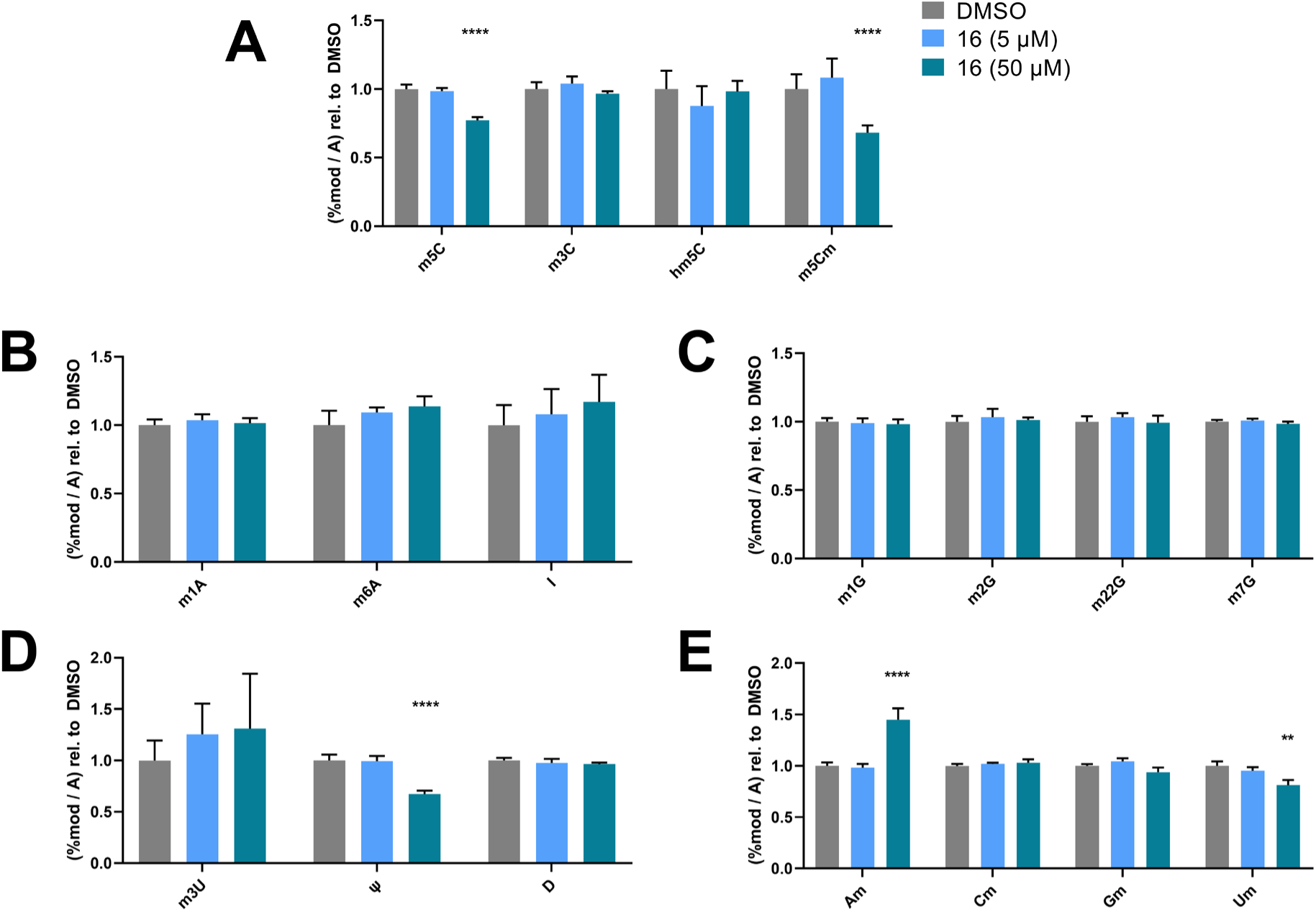
LC-MS/MS-based assay results for cellular tRNA modification changes analysis. MOLM-13 cells were treated for 72 h with **16** (5 and 50 µM); tRNA^Asp-GTC^, tRNA^Val-AAC^, and tRNA^Gly-GCC^ were enriched by an antisense pull-down strategy. (**A**) Changes in cytosine modifications relative to the DMSO control. (**B**) Changes in adenine modifications relative to the DMSO control. (**C**) Changes in guanine modifications relative to DMSO control. (**D**) Changes in uridine modifications relative to DMSO control. (**E**) Changes in 2’*O*-methylations relative to the DMSO control. * p<0.05, ** p<0.01, *** p<0.001, **** p<0.0001.

Since the genetic DNMT2 knockdown leads only to minor morphological changes in healthy and cancer cells, but the use of the FDA-approved pan-DNMT/NSUN inhibitor azacitidine is an effective cytostatic drug in leukemic cells, the question of whether DNMT2 is a cell viability-affecting drug target for cancer has been discussed for years.(71) We therefore investigated the effect of the DEL hit compounds **1**–**5** and **16** on the cell viability of MOLM-13 cells and compared it with the effect of azacitidine (SI Figure 9).(67, 68) Using an ATP-quantifying cell viability assay, after treatment with varying concentrations of test compounds for 48 h, triazines (**1** and **2**) exhibited cellular toxicity only at concentrations >100 µM, while the peptidomimetic hit compounds (**3**–**5** and **16**) showed only slight reduction in cell viability in MOLM-13 cells at high concentrations (CC_50_>50 µM). This suggests that the viability is not affected due to DNMT2 inhibition, since 50 µM of **16** leads to an efficient m^5^C level reduction in MOLM-13 cells. This is further supported by cell viability experiments in HCT mutant cells including the investigation of HCT DNMT2 knock-down and HCT NSUN2 knock-down cell lines highlighting low toxicity of all peptidomimetic compounds at 100 µM. From these cell-based investigations, we conclude that the cytotoxic effect of azacitidine (CC_50_<10 µM in all MOLM-13 and HCT cells) is not a cause of DNMT2 inhibition but mediated by DNMT1/3 and NSUNs inhibition. In summary, our novel DNMT2 inhibitor 16 is well capable of reducing tRNA m^5^C levels in a cellular context but at the same time not reducing the cell viability of MOLM-13 and HCT cell lines.

### Known DNMT2-interacting proteins do not bind to the allosteric pocket

The discovery of allosteric DNMT2 modulators mediating cellular m^5^C reduction implicates the question of whether this allosteric pocket may play a role in the cellular regulation of DNMT2 activity. The peptidomimetic structure of the identified hit compounds **3**–**5** might hint at a putative protein-based endogenous ligand, but so far, no allosteric DNMT2 interaction partners have been reported to regulate the enzymatic function. Yet, it was previously found by immunoprecipitation and peptide fingerprinting experiments that human DNMT2 shows conserved interaction with a handful of cellular proteins, which are involved in RNA processing and stress response, including eIF4E, splicing factors NonO resp. SfpQ, and proteins involved in RNA transport.(21) However, it is unknown in which conformation these proteins bind to DNMT2 and whether these can influence the DNMT2 activity.

To study a potential connection to our elucidated allosteric pocket, we probed the interaction between recombinant eIF4E resp. NonO proteins and DNMT2 by means of a pull-down competition assay. In this regard, we immobilized biotinylated DNMT2 on streptavidin-coated magnetic beads and incubated with recombinant eIF4E or NonO, which are able to bind to DNMT2. Subsequent treatment with the established DNMT2 ligands SFG (100 µM), compound **16** (100 µM), or tRNA^Asp^ (10 µM) was used to analyze if those ligands are able to dissociate the complex between eIF4E/NonO and DNMT2. SDS-PAGE analysis of the pulled-down fractions including the separated beads (solid) and supernatants (wash-out) revealed that NonO and eIF4E show efficient binding to DNMT2 on magnetic beads but do not bind to the unfunctionalized beads (Figure 10 and SI Figure 16), confirming the reports these proteins are in fact DNMT2 interactors. Yet, neither NonO nor eIF4E could be displaced from binding DNMT2 by competitive treatment with compound **16**, sinefungin (SFG), or tRNA^Asp^, indicating that both NonO and eIF4E do not bind to the allosteric binding pocket identified.

**Figure 10:**
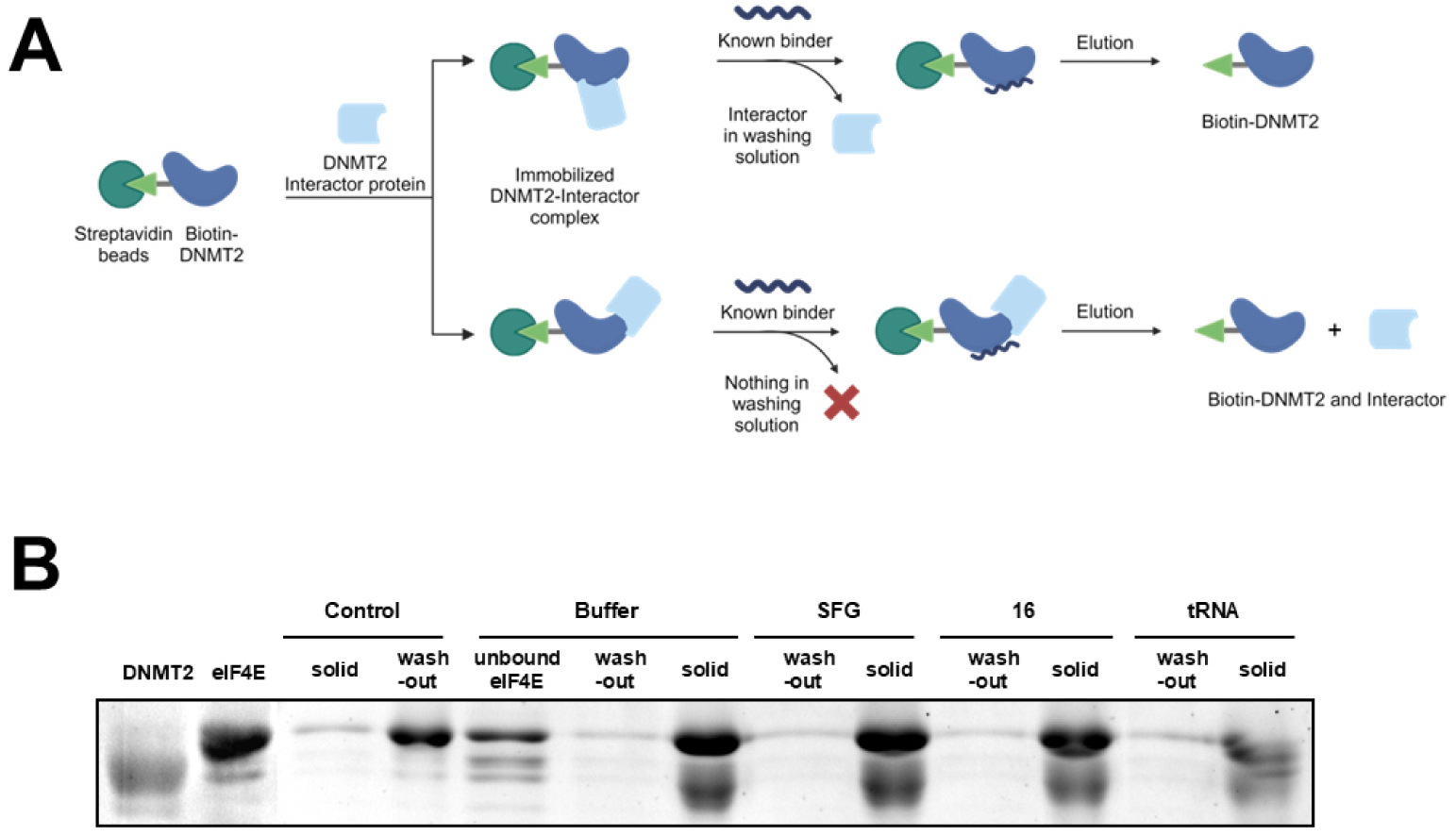
Known DNMT2-interacting proteins do not bind to the allosteric pocket. (**A**) Pull-down competition assay workflow. Biotinylated DNMT2 was immobilized on magnetic streptavidin beads and incubated with recombinant eIF4E protein. Subsequent treatment with the known ligands sinefungin (SFG), compound **16**, or tRNA^Asp^ was used to examine if such ligands can dissociate the DNMT2-eIF4E interaction. (**B**) Results of the SDS-PAGE-based analysis. Lane description from left to right: <u>DNMT2/eIF4E</u>: reference proteins for size comparison; <u>Control:</u> beads without immobilized DNMT2 served as the negative control, i.e. on these beads no binding of eIF4E was observed. <u>Treatment (buffer, SFG, **16**, tRNA):</u> The lane “unbound eIF4E” contains the supernatant after immobilization of eIF4E, which showed low protein levels, thus most eIF4E is effectively immobilized on DNMT2 beads. Lanes labeled with (wash-out) contain the supernatant of the respective treatment conditions (buffer, 100 µM SFG, 100 µM **16**, 10 µM tRNA^Asp^), while bands labeled with (solid) contain the solid fraction obtained by boiling the residual beads in Laemmli buffer.

We hypothesize, that both proteins might bind to a distinctive epitope outside of the MTase active site and reject the hypothesis that the newly identified allosteric pocket is an endogenous binding site at least for these known DNMT2 interacting proteins.

### Conclusion

In summary, our DEL screening successfully identified compounds that target a cryptic allosteric pocket of DNMT2, inducing a conformational change that blocks the SAH-binding active site and leads to subsequent DNMT2 inhibition. In-depth characterization of biomolecular interactions and the development of fluorescent DNMT2 probes revealed that this novel ligand chemotype showed strong selectivity for DNMT2 within the DNMT/NSUN family. DNMT2 ligand complex crystallography revealed the unique inhibitor binding mode by an active site loop reorganization. Synthetic structure-activity relationship studies were performed to improve the affinity of the allosteric ligand, which led to the discovery of a potent lead compound (*K_D_*∼3 µM) with improved passive cell permeability and the ability to inhibit DNMT2 in a cellular context as demonstrated by reduced tRNA m^5^C content. By this, we could also confirm that selective chemical DNMT2 inhibition is not affecting cell viability in different cancer cells, questioning its proposed role as a potential cancer drug target. Yet, the physiological role of the ligand-induced conformational change and potential allosteric interacting endogenous factors remain partly unelucidated and will need further experimentation (e.g., the question of whether this allosteric pocket may play a role in the cellular regulation of DNMT2 activity).

## DATA AVAILABILITY

Coordinates and structure factors of the DNMT2Δ47-compound 3 complex are deposited at the Protein Data Bank PDB under accession code 9HGM.

## SUPPLEMENTARY DATA

Supplementary Data are available online.

## AUTHOR CONTRIBUTION

A.F.F. performed DEL panning, analysis of hit clustering data, hit selection, microscale thermophoresis, ^3^H-assays, FP assays, synthesis of compounds, SeeSAR analysis, and writing of the manuscript; M.S. performed cloning, expression, purification, and characterization of DNMT2Δ47, protein crystallization, data collection, model building, and writing of the manuscript; A.C.W. performed ITC and nanoDSF measurements; V.K. performed synthesis, MST measurements, AS-MS assay, and pull-down assays; J.K. performed protein crystallization, data collection, model building, and writing of the manuscript; Z.N. performed protein expression and tRNA preparation; R.A.Z. performed protein expression and developed the idea for DNMT2Δ47 mutant; L.G. performed cell viability assays and tRNA LC/MS sample preparation; C.Z. performed PAMPA; M.J. performed cell viability assays; K.F. concept of study, supervision, and writing of the manuscript; M.H. concept of study, supervision, and writing of the manuscript; I.S. concept of study, supervision, and writing of the manuscript; F.B. concept of study, supervision, MOLM-13 cultivation, and writing of the manuscript. All authors commented on the manuscript.

## ACKNOWLEDGEMENTS

We thank Francesca Tuorto (Heidelberg) for providing NSUN2 knock-down cell lines, Ganna Podoprygorina (Mainz) for support during LC/MS measurements, WuXi AppTec for support during sequencing and data analysis of the DEL screenings, Hanna Bodenstaff (Mainz) for support during the synthesis of the inhibitors, Karin Pauly (Mainz) for support during cell viability assays, and Michael Klein (Mainz) for support during the preparation of Biorender illustrations. We acknowledge access to ESRF Grenoble beamline ID30B and thank the ESRF staff for their support. We thank Claudia Siegmann from the CCTP/BZH crystallization platform for excellent technical support. F.B. acknowledges the support from TRR319 (Project-ID 439669440) TP A01. I.S., M.S., and J.K. acknowledge the service SDS@hd supported by the Ministry of Science, Research and Arts Baden-Württemberg and the DFG through grants INST 35/1314-1 FUGG, INST 35/1503-1 FUGG.

## FUNDING

Financial support by the Deutsche Forschungsgemeinschaft (DFG, German Research Foundation) via TRR319 (Project-ID 439669440) B03 to I.S. and TP C01 to M.H. is gratefully acknowledged.

## CONFLICT OF INTEREST

Mark Helm is a consultant for Moderna Inc.

