## Supplementary material for "DNA-encoded Library Screening Uncovers Potent DNMT2 Inhibitors Targeting a Cryptic Allosteric Binding Site": SI

**ABSTRACT**

The human RNA methyltransferase DNMT2 is thought to be involved in various pathophysiological processes, yet, a major challenge in drug targeting DNMT2 is given by the fact that current SAH-derived inhibitors have poor target selectivity and limited cellular permeability. In this study, we have performed a DNA-encoded library (DEL) screening on DNMT2 yielding five non-SAH-like hit structures, three of which feature a peptidomimetic scaffold. All DEL hits could be validated by orthogonal biophysical and biochemical assays for DNMT2 binding. At the same time, the lead structure did not interact with related methyltransferases from the DNMT and NSUN families highlighting an unmatched DNMT2-targeting selectivity profile. Subsequent crystallographic studies revealed the unique ligand binding mode including an active site loop rearrangement and the formation of a cryptic allosteric binding pocket able to modulate the enzymatic activity by non-covalent DNMT2 dimerization. Based on the crystallographic results, we performed a structure-activity relationship study around the inhibitor lead structure resulting in an optimized DNMT2 inhibitor ($K_D$=3.04 µM), which was able to reduce $m^5C$ levels in MOLM-13 tRNA.

#### Supplementary data of DNA encoded library screening

DEL screenings were performed as described in the method section. SI Figure 1 shows additional information about quality controls performed during the execution, enrichment scores, and the libraries featuring hit compounds **1–5**.

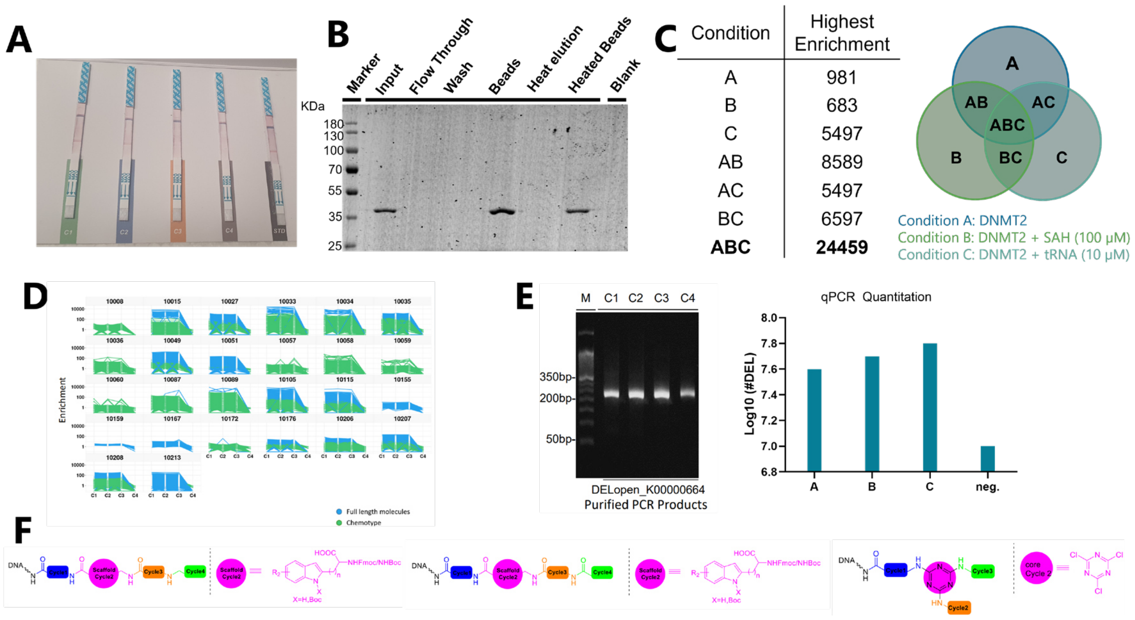

**Figure 1:** Supplementary data for the DNA encoded library screening on DNMT2. (**A**) Quality control test as provided by Wuxi AppTec. The four tested conditions mentioned in the manuscript are labeled here as C1–C4 (in the manuscript A–D). STD represents the internal standard according to the WuXi protocol (https://hits.wuxiapptec.com/assets/pdf/DELopen_Protocol.pdf). (**B**) DNMT2 capture assay was performed according to the protocol instructions to verify that DNMT2 can be stably immobilized on magnetic beads. The labeling is according to the original protocol. (**C**) Total count of highest enrichment scores for different panning conditions. The conditions are explained in the Venn diagram. (**D**) Overall performance and enrichment scores of the different libraries after the removal of NTC (no target control) binders. Most hits were enriched across C1–C3 equally, meaning neither SAH nor tRNA led to DEL hit displacement. (**E**) Quality control (gel electrophoresis and qPCR) was performed by Wuxi AppTec for all selection conditions after the second panning round. (**F**) Structures of the libraries featuring hit compounds chosen for off-DNA synthesis and further investigations.

#### Summary of all synthesized compound 3 analogs

**Table 1** Overview of all synthesized compounds during the SAR study around compound **3** and their characterization by MST ($F_{norm}$) and ITC ($K_D$). All results include the mean value and standard deviations from at least technical triplicate measurements. Raw data and analysis plots are depicted in Figure 2 and SI Figures 3–7.

| Cpd | BB1 | BB2 | BB3 | BB4 | C-Term. | $F_{norm}$ (% displ.) | $K_D$ (ITC) [µM] |
|---|---|---|---|---|---|---|---|
| **3** | 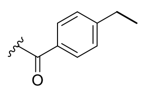 (4-ethylbenzoyl) | 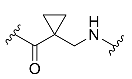 (cyclopropyl-β-alanine) | 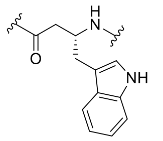 (β-homo-(*R*)-tryptophane) | 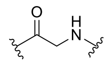 (glycine) | NH-Me | 912.1 (53%) | 8.67 ± 2.83 |
| **5** | 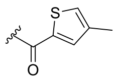 | cyclopropyl-β-alanine | β-homo-(*R*)-tryptophane | glycine | NH-Me | 879.3 (13%) | 13.6 ± 1.6 |
| **9** | 4-ethylbenzoyl | cyclopropyl-β-alanine | β-homo-(*R*)-tryptophane | glycine | OH | 907.7 (46%) | n.d. |
| **10** | 4-ethylbenzoyl | cyclopropyl-β-alanine | β-homo-(*R*)-tryptophane | glycine | $NH_2$ | 903.3 (40%) | 2.02 ± 0.68 |
| **11** | 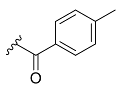 | cyclopropyl-β-alanine | β-homo-(*R*)-phenylalanine | glycine | $NH_2$ | 891.2 (24%) | 14.4 ± 5.6 |
| **12** | 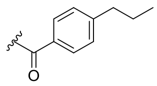 | cyclopropyl-β-alanine | β-homo-(*R*)-phenylalanine | glycine | $NH_2$ | 890.4 (23%) | n.d. |
| **13** | 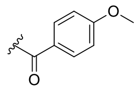 | cyclopropyl-β-alanine | β-homo-(*R*)-phenylalanine | glycine | $NH_2$ | 897.7 (32%) | n.d. |

| | | | | | | | |
|---|---|---|---|---|---|---|---|
| **14** | 4-ethylbenzoyl | cyclopropyl-β-alanine | β-homo-(*R*)-phenylalanine | glycine | $NH_2$ | 896.0 (29%) | 2.83 ± 1.91 |
| **15** | 4-ethylbenzoyl | cyclopropyl-β-alanine | 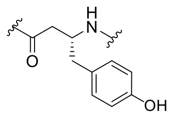 | glycine | $NH_2$ | 862.9 (5%) | n.d. |
| **16** | 4-ethylbenzoyl | cyclopropyl-β-alanine | 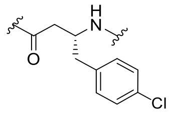 | glycine | $NH_2$ | 928.7 (76%) | 3.04 ± 0.57 |
| **17** | 4-ethylbenzoyl | cyclopropyl-β-alanine | 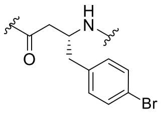 | glycine | $NH_2$ | 924.6 (71%) | n.d. |
| **18** | 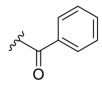 | cyclopropyl-β-alanine | 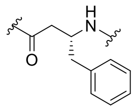 (β-homo-(*R*)-phenylalanine) | glycine | $NH_2$ | 868.2 (7%) | n.d. |
| **19** | 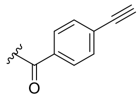 | cyclopropyl-β-alanine | β-homo-(*R*)-phenylalanine | glycine | $NH_2$ | 871.4 (8%) | n.d. |
| **20** | 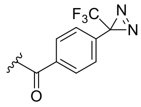 | cyclopropyl-β-alanine | β-homo-(*R*)-phenylalanine | glycine | $NH_2$ | 876.5 (11%) | n.d. |
| **21** | 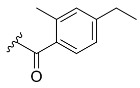 | cyclopropyl-β-alanine | β-homo-(*R*)-phenylalanine | glycine | $NH_2$ | 862.4 (5%) | n.d. |
| **22** | 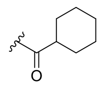 | cyclopropyl-β-alanine | β-homo-(*R*)-phenylalanine | glycine | $NH_2$ | 864.4 (5%) | n.d. |

| | | | | | | | |
|---|---|---|---|---|---|---|---|
| **23** | 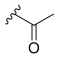 | cyclopropyl-β-alanine | β-homo-(*R*)-phenylalanine | glycine | $NH_2$ | 864.4 (5%) | n.d. |
| **24** | 4-ethylbenzoyl | 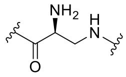 | β-homo-(*R*)-tryptophane | glycine | $NH_2$ | 860.8 (4%) | n.d. |
| **25** | 4-ethylbenzoyl | 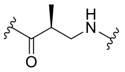 | β-homo-(*R*)-tryptophane | glycine | $NH_2$ | 855.5 (3%) | n.d. |
| **26** | 4-ethylbenzoyl | 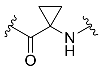 | β-homo-(*R*)-tryptophane | glycine | $NH_2$ | 861.9 (5%) | n.d. |
| **27** | 4-ethylbenzoyl | 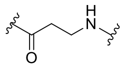 | β-homo-(*R*)-tryptophane | glycine | $NH_2$ | 863.8 (5%) | n.d. |
| **28** | 4-ethylbenzoyl | 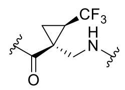 | β-homo-(*R*)-phenylalanine | glycine | $NH_2$ | 871.9 (8%) | n.d. |
| **29** | 4-ethylbenzoyl | 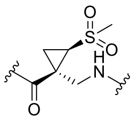 | β-homo-(*R*)-phenylalanine | glycine | $NH_2$ | 872.3 (9%) | n.d. |
| **30** | 4-ethylbenzoyl | cyclopropyl-β-alanine | 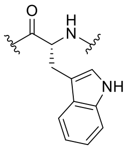 | glycine | $NH_2$ | 861.5 (5%) | n.b. |
| **31** | 4-ethylbenzoyl | cyclopropyl-β-alanine | 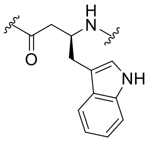 | glycine | $NH_2$ | 857.0 (4%) | n.d. |
| **32** | 4-ethylbenzoyl | cyclopropyl-β-alanine | 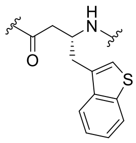 | glycine | $NH_2$ | 856.7 (3%) | n.d. |

| | | | | | | | |
|---|---|---|---|---|---|---|---|
| **33** | 4-ethylbenzoyl | cyclopropyl-β-alanine | 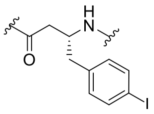 | glycine | $NH_2$ | 885.6 (18%) | n.d. |
| **34** | 4-ethylbenzoyl | cyclopropyl-β-alanine | 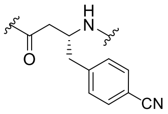 | glycine | $NH_2$ | 879.4 (13%) | n.d. |
| **35** | 4-ethylbenzoyl | cyclopropyl-β-alanine | 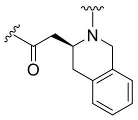 | glycine | $NH_2$ | 861.5 (5%) | n.d. |
| **36** | 4-ethylbenzoyl | cyclopropyl-β-alanine | 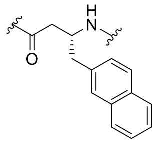 | glycine | $NH_2$ | 861.7 (5%) | n.d. |
| **37** | 4-ethylbenzoyl | cyclopropyl-β-alanine | 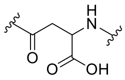 | glycine | $NH_2$ | 858.6 (4%) | n.d. |
| **38** | 4-ethylbenzoyl | cyclopropyl-β-alanine | 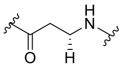 | glycine | $NH_2$ | 859.4 (4%) | n.d. |
| **39** | 4-ethylbenzoyl | cyclopropyl-β-alanine | β-homo-(*R*)-phenylalanine | 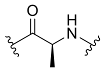 | $NH_2$ | 871.4 (8%) | n.d. |
| **40** | 4-ethylbenzoyl | cyclopropyl-β-alanine | β-homo-(*R*)-phenylalanine | 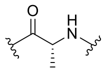 | $NH_2$ | 870.1 (8%) | n.d. |
| **41** | 4-ethylbenzoyl | cyclopropyl-β-alanine | β-homo-(*R*)-phenylalanine | 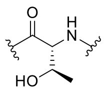 | $NH_2$ | 865.3 (6%) | n.d. |
| **42** | 4-ethylbenzoyl | cyclopropyl-β-alanine | β-homo-(*R*)-phenylalanine | 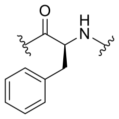 | $NH_2$ | 858.8 (4%) | n.d. |
| **43** | 4-ethylbenzoyl | cyclopropyl-β-alanine | β-homo-(*R*)-phenylalanine | 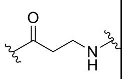 | $NH_2$ | 874.7 (10%) | n.d. |

| | | | | | | | |
|---|---|---|---|---|---|---|---|
| **44** | 4-ethylbenzoyl | 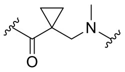 | β-homo-(*R*)-phenylalanine | glycine | $NH_2$ | 869.7 (7%) | n.d. |
| **45** | 4-ethylbenzoyl | cyclopropyl-β-alanine | 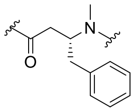 | glycine | $NH_2$ | 862.6 (5%) | n.d. |
| **46** | 4-ethylbenzoyl | cyclopropyl-β-alanine | β-homo-(*R*)-phenylalanine | 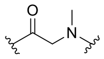 | $NH_2$ | 868.4 (7%) | n.d. |
| **47** | 4-ethylbenzoyl | cyclopropyl-β-alanine | β-homo-(*R*)-phenylalanine | glycine | $NMe_2$ | 901.0 (36%) | 10.9 ± 7.3 |
| **48** | 4-ethylbenzoyl | cyclopropyl-β-alanine | β-homo-(*R*)-phenylalanine | glycine | NH-PEG4 | 894.1 (27%) | n.d. |

#### General procedure for solid phase synthesis (SPPS)

Rink amide AM resin (loading capacity: 0.6 – 0.8 mmol/g) or Wang Gly resin (loading capacity: 0.135 mmol/g) was swelled twice for 5 min in DMF. Fmoc-deprotection was performed by treating with 40% piperidine in DMF (2x 10 min shaking). The resin was rinsed with DCM (3x) and DMF (3x). In the next step, the corresponding building block was coupled to the resin by dissolving the amino acid (3.0 eq or 1.2 eq.), TBTU (3.0 eq or 1.2 eq.), and NMM (9.0 eq.) in DMF and shaking for 1.5 hours (or 16 hours for 1.2 eq). Subsequently, the resin was rinsed with DCM (3x) and DMF (3x). To couple the next amino acids, the procedure of deprotection and coupling was repeated. When the last amino acid was coupled, the resin was rinsed with DMF (3x) and DCM (3x). To cleave the peptide from the resin, the resin was treated with a mixture of TFA:TIPS:DCM (50:10:40) for 1.5 h. The resin was washed with DCM (3x) and treated again with TFA:TIPS:DCM (50:10:40) for 16 h. All cleavage and wash solutions combined were evaporated under reduced pressure. The crude product was purified by reversed-phase flash column chromatography (Biotage Isolera, ACN/$H_2O$ + TFA), and the product containing fractions were lyophilized.

#### Characterization of compounds

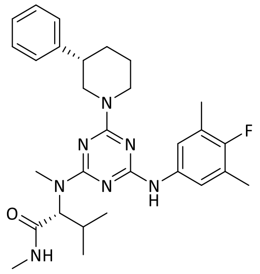

**1**

Compound **1**: $^{1}$H-NMR (400 MHz, DMSO-*d*$_6$): δ [ppm] = 8.94 (s, 1H), 7.74 – 7.16 (m, 6H), 4.76 (s, 2H), 3.32 (d, *J* = 2.8 Hz, 2H), 3.05 – 2.83 (m, 3H), 2.58 (s, 13H), 2.34 – 1.90 (m, 6H), 1.79 (d, *J* = 13.0 Hz, 1H), 0.93 (d, *J* = 4.3 Hz, 2H), 0.79 (d, *J* = 6.6 Hz, 2H).

$^{13}$C-NMR (101 MHz, DMSO): δ [ppm] = 170.7, 170.1, 165.9, 165.8, 143.9, 143.9, 136.0, 128.4, 127.0, 126.4, 123.1, 119.8, 62.6, 49.7, 43.4, 42.2, 31.8, 26.4, 25.7, 25.5, 25.1, 20.0, 19.1, 18.8, 14.5.

LC/MS: m/z calculated for $C_{29}H_{38}FN_7O$ $[M+H]^+$: 520.3, found: 520.3.

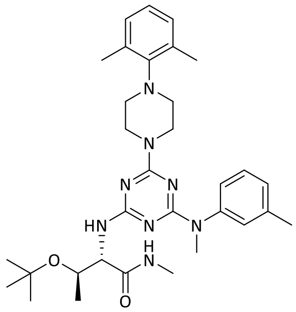

**2**

Compound **2**: $^{1}$H-NMR (400 MHz, DMSO-$d_6$): δ [ppm] = 7.77 – 7.68 (m, 1H), 7.27 – 7.07 (m, 3H), 6.95 (h, *J* = 7.7, 7.1 Hz, 4H), 4.02 (t, *J* = 5.3 Hz, 1H), 3.78 – 3.68 (m, 3H), 3.38 (s, 3H), 3.32 (s, 3H), 2.97 (d, *J* = 6.0 Hz, 4H), 2.57 (d, *J* = 4.4 Hz, 3H), 2.32 – 2.23 (m, 9H), 1.06 (dd, *J* = 15.2, 8.3 Hz, 12H).

$^{13}$C-NMR (101 MHz, DMSO): δ [ppm] = 171.0, 165.6, 165.2, 164.4, 148.0, 144.7, 144.5, 137.6, 137.3, 136.1, 128.9, 128.1, 127.9, 127.0, 126.7, 125.7, 125.4, 124.9, 123.7, 123.0, 73.3, 67.2, 59.2, 49.3, 44.1, 37.3, 28.0, 25.7, 21.0, 20.7, 20.5, 19.4.

LC/MS: m/z calculated for $C_{32}H_{46}N_8O_2$ $[M+H]^+$: 575.3, found: 575.2.

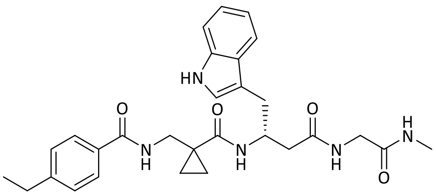

**3**

Compound **3**: $^{1}$H-NMR (600 MHz, DMSO-$d_6$): δ [ppm] = 10.80 – 10.76 (m, 1H), 8.58 (t, *J* = 6.1 Hz, 1H), 8.13 (t, *J* = 5.9 Hz, 1H), 8.05 (d, *J* = 7.9 Hz, 1H), 7.76 (d, *J* = 8.2 Hz, 2H), 7.66 (d, *J* = 4.6 Hz, 1H), 7.59 (d, *J* = 7.9 Hz, 1H), 7.29 (t, *J* = 8.2 Hz, 3H), 7.09 (d, *J* = 2.2 Hz, 1H), 7.07 – 7.01 (m, 1H), 6.98 – 6.92 (m, 1H), 4.38 – 4.31 (m, 1H), 3.60 (ddd, *J* = 25.4, 15.6, 6.2 Hz, 2H), 3.48 (dd, *J* = 16.5, 5.7 Hz, 1H), 3.41 (dd, *J* = 14.8, 6.2 Hz, 1H), 2.83 (d, *J* = 6.5 Hz, 2H), 2.64 (q, *J* = 7.6 Hz, 2H), 2.54 (d, *J* = 4.6 Hz, 3H), 2.40 – 2.30 (m, 2H), 1.18 (t, *J* = 7.6 Hz, 3H), 0.91 (t, *J* = 3.6 Hz, 2H), 0.79 (dd, *J* = 9.4, 3.6 Hz, 2H).

$^{13}$C-NMR (151 MHz, DMSO): δ [ppm] = 171.6, 170.8, 169.3, 167.2, 147.5, 136.1, 131.4, 127.7, 127.6, 127.5, 123.3, 120.9, 118.6, 118.2, 111.3, 111.2, 47.8, 42.2, 42.1, 29.9, 28.1, 25.5, 25.3, 15.4, 13.2, 12.9.

LC/MS: m/z calculated for $C_{29}H_{35}N_5O_4$ $[M+H]^+$: 518.3, found: 518.2.

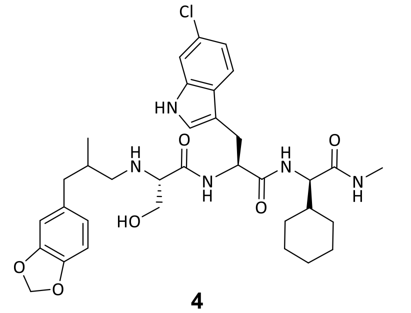

**4**

Compound **4**: $^{1}$H-NMR (400 MHz, DMSO-$d_6$): δ [ppm] = 10.97 (s, 1H), 8.03 (d, *J* = 8.2 Hz, 1H), 7.90 (d, *J* = 10.1 Hz, 2H), 7.51 (d, *J* = 8.5 Hz, 1H), 7.31 (d, *J* = 6.4 Hz, 1H), 7.09 (s, 1H), 6.92 (d, *J* = 8.5 Hz, 1H), 6.76 (d, *J* = 7.8 Hz, 1H), 6.66 (s, 1H), 6.52 (d, *J* = 7.9 Hz, 1H), 5.94 (s, 2H), 4.92 (s, 1H), 4.62 (q, *J* = 6.9 Hz, 1H), 4.08 (t, *J* = 8.2 Hz, 1H), 3.47 (d, *J* = 11.1 Hz, 1H), 3.03 (ddt, *J* = 37.3, 22.9, 5.8 Hz, 3H), 2.62 – 2.54 (m, 4H), 2.37 – 1.94 (m, 4H), 1.58 (dt, *J* = 18.1, 11.0 Hz, 6H), 1.45 (d, *J* = 12.6 Hz, 1H), 1.21 – 0.77 (m, 6H), 0.63 (d, *J* = 6.4 Hz, 3H).

$^{13}$C-NMR (101 MHz, DMSO): δ [ppm] = 171.8, 171.0, 170.5, 147.0, 145.1, 136.3, 134.7, 126.4, 125.5, 124.9, 121.7, 120.0, 118.4, 110.7, 109.9, 109.2, 107.8, 100.5, 64.8, 62.1, 57.1, 53.9, 52.6, 35.1, 29.1, 28.3, 27.5, 25.8, 25.5, 25.4, 17.3.

LC/MS: m/z calculated for $C_{34}H_{44}ClN_5O_6$ $[M+H]^+$: 654.3, found: 654.3.

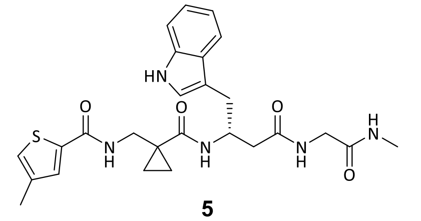

**5**

Compound **5**: $^{1}$H-NMR (400 MHz, DMSO-$d_6$): δ [ppm] = 10.76 (s, 1H), 8.49 (t, *J* = 7.0 Hz, 1H), 8.10 (d, *J* = 6.6 Hz, 1H), 7.86 – 7.80 (m, 1H), 7.60 (q, *J* = 8.9, 7.7 Hz, 3H), 7.35 (s, 1H), 7.30 (d, *J* = 8.2 Hz, 1H), 7.09 (s, 1H), 7.07 – 7.00 (m, 1H), 6.99 – 6.92 (m, 1H), 4.41 – 4.30 (m, 1H), 2.82 (t, *J* = 4.7 Hz, 2H), 2.54 (d, *J* = 4.3 Hz, 4H), 2.34 (d, *J* = 7.0 Hz, 2H), 2.21 (d, *J* = 3.2 Hz, 3H), 1.24 (s, 1H), 1.03 – 0.85 (m, 3H), 0.76 (s, 2H).

$^{13}$C-NMR (101 MHz, DMSO): δ [ppm] = 172.0, 171.3, 169.7, 162.4, 139.2, 138.3, 136.6, 130.9, 128.0, 126.9, 123.8, 121.3, 119.0, 118.6, 111.7, 111.6, 48.3, 42.6, 30.4, 25.9, 25.6, 23.8, 15.9, 13.4, 13.1.

LC/MS: m/z calculated for $C_{26}H_{31}N_5O_4S$ $[M+H]^+$: 510.2, found: 510.1.

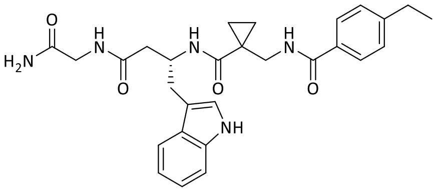

**10**

Compound **10**: Yield: 2.9 mg (5.80 µmol ≙ 13% of th.), colorless solid.

$^{1}$H-NMR (600 MHz, DMSO-$d_6$): δ [ppm] = 10.77 (s, 1H), 8.59 (t, *J* = 6.2 Hz, 1H), 8.07 (t, *J* = 5.9 Hz, 1H), 8.04 (d, *J* = 7.9 Hz, 1H), 7.81 – 7.71 (m, 2H), 7.59 (d, *J* = 7.9 Hz, 1H), 7.33 – 7.26 (m, 3H), 7.24 (s, 1H), 7.09 (d, *J* = 2.2 Hz, 1H), 7.04 (t, *J* = 7.3 Hz, 2H), 6.95 (t, *J* = 7.4 Hz, 1H), 4.34 (h, *J* = 6.9 Hz, 1H), 3.61 (dd, *J* = 16.8, 6.0 Hz, 1H), 3.56 (dd, *J* = 14.8, 6.1 Hz, 1H), 3.48 (dd, *J* = 16.8, 5.5 Hz, 1H), 3.41 (dd, *J* = 14.8, 6.1 Hz, 1H), 2.82 (d, *J* = 6.7 Hz, 2H), 2.64 (q, *J* = 7.6 Hz, 2H), 2.33 (d, *J* = 6.8 Hz, 1H), 1.18 (t, *J* = 7.6 Hz, 3H), 0.98 – 0.72 (m, 4H).

$^{13}$C-NMR (151 MHz, DMSO): δ [ppm] = 171.6, 171.1, 170.6, 167.2, 147.5, 136.1, 131.4, 127.7, 127.6, 127.5, 123.3, 120.8, 118.6, 118.2, 111.3, 111.2, 47.7, 42.3, 41.9, 40.1, 29.8, 28.1, 25.3, 15.4, 13.3, 12.9.

LC/MS: m/z calculated for $C_{28}H_{33}N_5O_4$ $[M+H]^+$: 504.3, found: 504.1.

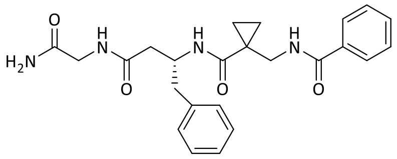

**18**

Compound **18**: Yield: 15.0 mg (0.034 mmol ≙ 75% of th.), colorless solid.

$^{1}$H-NMR (300 MHz, DMSO-$d_6$): δ [ppm] = 8.62 (t, *J* = 6.1 Hz, 1H), 8.10 (t, *J* = 5.9 Hz, 1H), 7.96 (d, *J* = 8.1 Hz, 1H), 7.91 – 7.75 (m, 2H), 7.67 – 7.36 (m, 3H), 7.24 (s, 1H), 7.20 – 7.03 (m, 5H), 7.03 (s, 1H), 4.30 (q, *J* = 7.5 Hz, 1H), 3.74 – 3.39 (m, 4H), 2.93 – 2.59 (m, 2H), 2.32 (d, *J* = 6.8 Hz, 2H), 1.04 – 0.56 (m, 4H).

$^{13}$C-NMR (75 MHz, DMSO): δ [ppm] = 171.5, 171.1, 170.4, 167.2, 138.8, 134.0, 131.5, 129.2, 128.3, 128.0, 127.4, 126.0, 48.4, 42.3, 41.9, 40.3, 25.2, 13.0.

LC/MS: m/z calculated for $C_{24}H_{28}N_4O_4$ $[M+H]^+$: 437.2, found: 437.2.

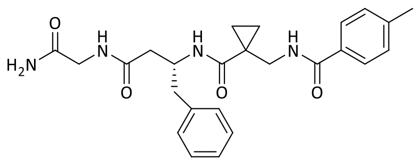

**11**

Compound **11**: Yield: 15.0 mg (0.033 mmol ≙ 75% of th.), colorless solid.

$^{1}$H-NMR (300 MHz, DMSO-$d_6$): δ [ppm] = 8.55 (t, $J$ = 6.1 Hz, 1H), 8.09 (t, $J$ = 5.9 Hz, 1H), 8.00 (d, $J$ = 8.1 Hz, 1H), 7.82 – 7.69 (m, 2H), 7.32 – 7.26 (m, 2H), 7.24 (s, 1H), 7.17 – 7.07 (m, 5H), 7.03 (s, 1H), 4.30 (q, $J$ = 7.0 Hz, 1H), 3.72 – 3.38 (m, 4H), 2.90 – 2.59 (m, 2H), 2.36 (s, 3H), 2.31 (d, $J$ = 6.9 Hz, 2H), 0.96 – 0.68 (m, 4H).

$^{13}$C-NMR (75 MHz, DMSO): δ [ppm] = 171.5, 171.1, 170.3, 167.1, 141.4, 138.8, 131.1, 129.2, 128.9, 128.0, 127.4, 126.0, 48.4, 42.3, 41.9, 25.3, 21.0, 13.0.

LC/MS: m/z calculated for $C_{25}H_{30}N_4O_4$ $[M+H]^+$: 451.2, found: 451.1.

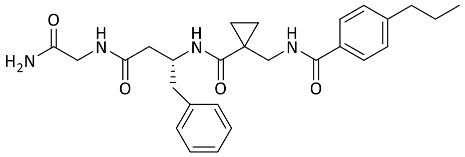

**12**

Compound **12**: Yield: 5.0 mg (0.010 mmol ≙ 23% of th.), colorless solid.

$^{1}$H-NMR (400 MHz, DMSO-$d_6$): δ [ppm] = 8.54 (t, $J$ = 6.3 Hz, 1H), 8.08 (t, $J$ = 6.0 Hz, 1H), 7.98 (d, $J$ = 8.1 Hz, 1H), 7.81 – 7.73 (m, 2H), 7.33 – 7.26 (m, 2H), 7.23 (s, 1H), 7.17 – 7.05 (m, 5H), 7.01 (s, 1H), 4.29 (p, $J$ = 7.2 Hz, 1H), 3.70 – 3.43 (m, 4H), 2.87 – 2.64 (m, 2H), 2.61 (t, $J$ = 7.7 Hz, 2H), 2.31 (d, $J$ = 6.9 Hz, 2H), 1.61 (h, $J$ = 7.1 Hz, 2H), 0.93 – 0.72 (m, 7H).

$^{13}$C-NMR (101 MHz, DMSO): δ [ppm] = 171.5, 171.1, 170.3, 167.2, 145.9, 138.8, 131.4, 129.1, 128.3, 127.9, 127.4, 125.9, 48.4, 42.3, 41.9, 40.3, 37.0, 25.2, 23.9, 13.6, 13.0, 13.0.

LC/MS: m/z calculated for $C_{27}H_{34}N_4O_4$ $[M+H]^+$: 479.3, found: 479.2.

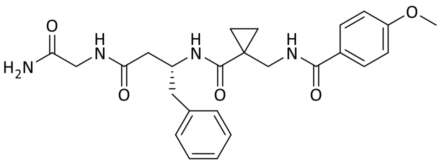

**13**

Compound **13**: Yield: 18.0 mg (0.040 mmol ≙ 86% of th.), colorless solid.

[1]H-NMR (300 MHz, DMSO-$d_6$): δ [ppm] = 8.50 (t, *J* = 6.1 Hz, 1H), 8.16 – 7.98 (m, 2H), 7.91 – 7.76 (m, 2H), 7.24 (s, 1H), 7.20 – 7.07 (m, 4H), 7.06 – 6.91 (m, 3H), 4.29 (q, *J* = 8.5 Hz, 1H), 3.82 (s, 3H), 3.72 – 3.40 (m, 4H), 2.90 – 2.58 (m, 2H), 2.31 (d, *J* = 6.8 Hz, 2H), 0.98 – 0.61 (m, 4H).

[13]C-NMR (75 MHz, DMSO): δ [ppm] = 171.5, 171.1, 170.4, 166.8, 161.8, 138.8, 129.3, 129.2, 128.0, 126.1, 126.0, 113.5, 55.4, 48.4, 42.3, 41.9, 25.3, 13.1.

LC/MS: m/z calculated for $C_{25}H_{30}N_4O_5$ $[M+H]^+$: 467.2, found: 467.1.

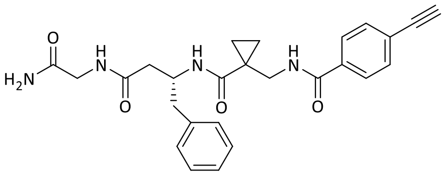

**19**

Compound **19**: Yield: 15.0 mg (0.033 mmol ≙ 71% of th.), pale brown solid.

[1]H-NMR (300 MHz, DMSO-$d_6$): δ [ppm] = 8.69 (t, *J* = 6.1 Hz, 1H), 8.10 (t, *J* = 5.8 Hz, 1H), 7.96 – 7.76 (m, 3H), 7.59 (d, *J* = 8.2 Hz, 2H), 7.24 (s, 1H), 7.20 – 7.06 (m, 5H), 7.03 (s, 1H), 4.39 (s, 1H), 4.30 (q, *J* = 8.2 Hz, 1H), 3.70 – 3.38 (m, 4H), 2.86 – 2.57 (m, 2H), 2.32 (d, *J* = 6.8 Hz, 2H), 0.96 – 0.61 (m, 4H).

[13]C-NMR (75 MHz, DMSO): δ [ppm] = 171.4, 171.1, 170.4, 166.4, 138.8, 134.0, 131.7, 129.1, 128.0, 127.7, 126.0, 124.7, 83.0, 82.9, 48.4, 42.5, 41.9, 40.2, 25.1, 12.9.

LC/MS: m/z calculated for $C_{26}H_{28}N_4O_4$ $[M+H]^+$: 461.2, found: 461.1.

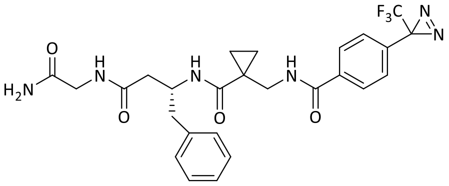

**20**

Compound **20**: Yield: 17.0 mg (0.031 mmol ≙ 69% of th.), colorless solid.

[1]H-NMR (300 MHz, DMSO-$d_6$): δ [ppm] = 8.74 (t, *J* = 6.0 Hz, 1H), 8.09 (t, *J* = 5.9 Hz, 1H), 8.01 – 7.89 (m, 2H), 7.82 (d, *J* = 8.2 Hz, 1H), 7.39 (d, *J* = 8.1 Hz, 2H), 7.23 (s, 1H), 7.18 – 7.03 (m, 4H), 7.03 (s, 1H), 4.29 (p, *J* = 7.1 Hz, 1H), 3.80 – 3.39 (m, 4H), 2.94 – 2.57 (m, 2H), 2.31 (d, *J* = 6.8 Hz, 2H), 0.99 – 0.69 (m, 4H).

[13]C-NMR (75 MHz, DMSO): δ [ppm] = 171.4, 171.1, 170.4, 166.1, 138.8, 135.6, 130.4, 129.1, 128.3, 127.9, 126.4, 125.9, 123.1, 120.4, 48.3, 42.5, 41.9, 40.2, 39.4, 28.3, 27.9, 25.1, 12.8.

LC/MS: m/z calculated for $C_{26}H_{27}F_3N_6O_4$ $[M+H]^+$: 545.2, found: 545.1.

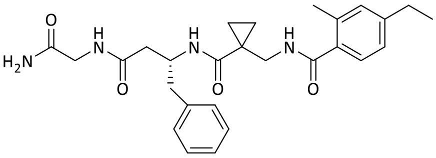

**21**

Compound **21**: Yield: 10.1 mg (0.021 mmol ≙ 46% of th.), colorless solid.

$^1$H-NMR (300 MHz, DMSO-$d_6$): δ [ppm] = 8.36 (t, *J* = 6.2 Hz, 1H), 8.09 (t, *J* = 5.8 Hz, 1H), 7.93 (d, *J* = 8.1 Hz, 1H), 7.30 – 7.11 (m, 7H), 7.11 – 6.92 (m, 3H), 4.40 – 4.20 (m, 1H), 3.69 – 3.45 (m, 3H), 3.43 – 3.25 (m, 1H), 2.74 (qd, *J* = 13.5, 6.8 Hz, 2H), 2.58 (q, *J* = 7.6 Hz, 2H), 2.39 – 2.16 (m, 5H), 1.17 (t, *J* = 7.6 Hz, 3H), 0.99 – 0.62 (m, 4H).

$^{13}$C-NMR (75 MHz, DMSO): δ [ppm] = 171.6, 171.1, 170.3, 170.0, 145.4, 138.9, 135.3, 134.0, 130.0, 129.2, 128.1, 127.4, 126.0, 124.9, 48.6, 41.9, 40.4, 28.0, 25.2, 19.6, 15.6, 12.9, 12.8.

LC/MS: m/z calculated for $C_{27}H_{34}N_4O_4$ [M+H]$^+$: 479.3, found: 479.2.

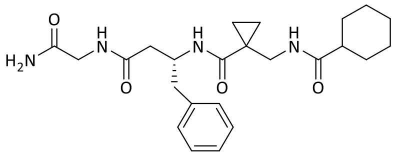

**22**

Compound **22**: Yield: 16.0 mg (0.036 mmol ≙ 79% of th.), colorless solid.

$^1$H-NMR (300 MHz, DMSO-$d_6$): δ [ppm] = 8.09 (t, *J* = 5.8 Hz, 1H), 7.96 (t, *J* = 6.2 Hz, 1H), 7.86 (d, *J* = 8.1 Hz, 1H), 7.34 – 7.09 (m, 6H), 7.02 (s, 1H), 4.38 – 4.10 (m, 1H), 3.60 (qd, *J* = 16.8, 5.8 Hz, 2H), 3.45 – 3.21 (m, 1H), 3.12 (dd, *J* = 14.7, 6.1 Hz, 1H), 2.89 – 2.56 (m, 2H), 2.27 (d, *J* = 6.9 Hz, 2H), 2.19 – 1.98 (m, 1H), 1.80 – 1.47 (m, 5H), 1.47 – 1.02 (m, 5H), 0.91 – 0.70 (m, 2H), 0.69 – 0.55 (m, 2H).

$^{13}$C-NMR (75 MHz, DMSO): δ [ppm] = 176.3, 171.5, 171.1, 170.3, 138.9, 129.2, 128.1, 126.0, 48.4, 43.9, 41.9, 41.3, 40.3, 29.2, 29.1, 25.4, 25.3, 25.1, 12.9, 12.8.

LC/MS: m/z calculated for $C_{24}H_{34}N_4O_4$ [M+H]$^+$: 443.3, found: 443.2.

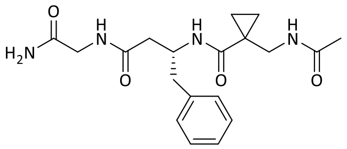

**23**

Compound **23**: Yield: 7.2 mg (0.018 mmol ≙ 41% of th.), colorless solid.

$^{1}$H-NMR (300 MHz, DMSO-$d_6$): δ [ppm] = 8.08 (q, $J$ = 5.9 Hz, 2H), 7.78 (d, $J$ = 8.0 Hz, 1H), 7.42 – 7.10 (m, 6H), 7.03 (s, 1H), 4.25 (p, $J$ = 6.9 Hz, 1H), 3.80 – 3.46 (m, 2H), 3.22 (qd, $J$ = 14.8, 6.2 Hz, 2H), 2.93 – 2.56 (m, 2H), 2.29 (d, $J$ = 6.9 Hz, 2H), 1.80 (s, 3H), 1.01 – 0.72 (m, 2H), 0.71 – 0.56 (m, 2H).

$^{13}$C-NMR (75 MHz, DMSO): δ [ppm] = 171.4, 171.1, 170.3, 170.3, 138.8, 129.2, 128.1, 126.0, 48.3, 41.9, 41.7, 40.0, 39.4, 25.1, 22.5, 12.9.

LC/MS: m/z calculated for $C_{19}H_{26}N_4O_4$ $[M+H]^+$: 375.2, found: 375.2.

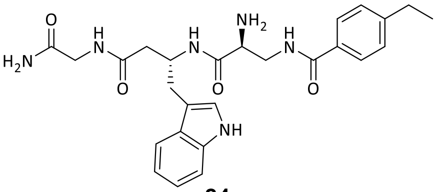

**24**

Compound **24**: Yield: 5.0 mg (0.010 mmol ≙ 23% of th.), colorless solid.

$^{1}$H-NMR (400 MHz, DMSO-$d_6$): δ [ppm] = 10.82 (s, 1H), 8.49 (d, $J$ = 8.0 Hz, 1H), 8.41 (t, $J$ = 5.7 Hz, 1H), 8.21 (s, 3H), 8.02 (t, $J$ = 5.8 Hz, 1H), 7.76 (d, $J$ = 7.9 Hz, 2H), 7.59 (d, $J$ = 7.9 Hz, 1H), 7.43 – 7.21 (m, 4H), 7.13 (s, 1H), 7.10 – 7.00 (m, 2H), 6.95 (t, $J$ = 7.4 Hz, 1H), 4.35 (q, $J$ = 7.0 Hz, 1H), 3.94 (t, $J$ = 6.1 Hz, 1H), 3.63 (d, $J$ = 5.7 Hz, 2H), 3.59 (t, $J$ = 5.9 Hz, 2H), 2.83 (m, 2H), 2.65 (q, J = 7.5 Hz, 2H), 2.44 – 2.29 (m, 2H), 1.18 (t, $J$ = 7.6 Hz, 3H).

$^{13}$C-NMR (101 MHz, DMSO): δ [ppm] = 171.1, 170.3, 167.1, 166.0, 147.6, 136.2, 131.3, 127.6, 127.5, 127.4, 123.5, 120.9, 118.5, 118.2, 111.3, 110.7, 52.3, 47.8, 41.7, 40.2, 39.0, 29.9, 28.0, 15.4.

LC/MS: m/z calculated for $C_{26}H_{32}N_6O_4$ $[M+H]^+$: 493.3, found: 493.2.

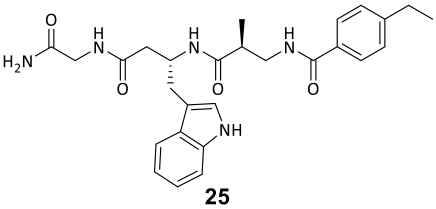

**25**

Compound **25**: Yield: 1.2 mg (2.520 µmol ≙ 1% of th.), colorless solid.

$^{1}$H-NMR (400 MHz, DMSO-$d_6$): δ [ppm] = 10.75 (s, 1H), 8.33 (t, $J$ = 5.8 Hz, 1H), 7.96 (t, $J$ = 5.8 Hz, 1H), 7.78 (d, $J$ = 8.2 Hz, 1H), 7.73 (d, $J$ = 7.9 Hz, 2H), 7.58 (d, $J$ = 7.9 Hz, 1H), 7.31 (d, $J$ = 8.1 Hz, 1H), 7.25 (d, $J$ = 7.5 Hz, 3H), 7.09 (s, 1H), 7.07 – 6.99 (m, 2H), 6.94 (t, $J$ = 7.4 Hz, 1H),

4.32 (q, *J* = 7.0 Hz, 1H), 3.69 – 3.48 (m, 2H), 3.28 – 3.12 (m, 2H), 2.81 (m, 2H), 2.69 – 2.53 (m, 3H), 2.38 – 2.16 (m, 2H), 1.21 – 1.09 (m, 3H), 0.98 (d, *J* = 6.8 Hz, 3H).

$^{13}$C-NMR (101 MHz, DMSO): δ [ppm] = 173.7, 171.1, 170.6, 166.3, 147.0, 136.1, 132.1, 127.6, 127.6, 127.2, 123.3, 120.8, 118.6, 118.2, 111.3, 111.1, 47.1, 42.5, 41.9, 30.2, 28.0, 15.6, 15.4.

LC/MS: m/z calculated for $C_{27}H_{33}N_5O_4$ $[M+H]^+$: 492.3, found: 492.1.

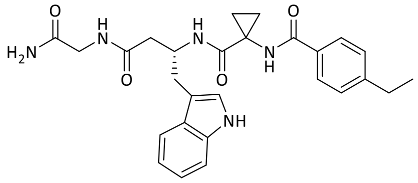

**26**

Compound **26**: Yield: 5.0 mg (0.010 mmol ≙ 23% of th.), colorless solid.

$^{1}$H-NMR (400 MHz, DMSO-*d*$_6$): δ [ppm] = 10.81 (s, 1H), 8.85 (s, 1H), 7.94 (t, *J* = 5.9 Hz, 1H), 7.87 – 7.73 (m, 3H), 7.60 (d, *J* = 7.9 Hz, 1H), 7.42 – 7.27 (m, 3H), 7.25 (s, 1H), 7.16 (s, 1H), 7.09 – 6.97 (m, 2H), 6.94 (t, *J* = 7.4 Hz, 1H), 4.30 (h, *J* = 6.6 Hz, 1H), 3.61 (t, *J* = 5.2 Hz, 2H), 2.99 – 2.71 (m, 2H), 2.66 (q, *J* = 7.6 Hz, 2H), 2.29 (ddd, *J* = 48.3, 14.5, 6.2 Hz, 2H), 1.44 – 1.28 (m, 2H), 1.26 – 1.10 (m, 3H), 1.06 – 0.86 (m, 2H).

$^{13}$C-NMR (101 MHz, DMSO): δ [ppm] = 171.1, 170.8, 170.8, 167.4, 147.4, 136.1, 131.8, 127.8, 127.5, 127.5, 123.6, 120.8, 118.5, 118.2, 111.2, 110.9, 41.8, 38.8, 34.7, 29.7, 28.1, 16.2, 15.5.

LC/MS: m/z calculated for $C_{27}H_{31}N_5O_4$ $[M+H]^+$: 490.2, found: 490.1.

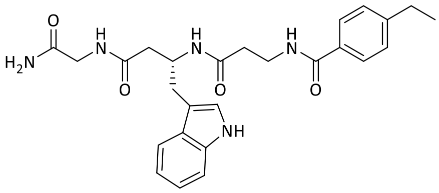

**27**

Compound **27**: Yield: 11.0 mg (0.023 mmol ≙ 50% of th.), colorless solid.

$^{1}$H-NMR (300 MHz, DMSO-*d*$_6$): δ [ppm] = 10.79 (d, *J* = 2.4 Hz, 1H), 8.38 (t, *J* = 5.6 Hz, 1H), 8.03 (t, *J* = 5.8 Hz, 1H), 7.87 (d, *J* = 8.2 Hz, 1H), 7.79 – 7.66 (m, 2H), 7.58 (d, *J* = 7.8 Hz, 1H), 7.37 – 7.19 (m, 4H), 7.11 (d, *J* = 2.2 Hz, 1H), 7.09 – 7.00 (m, 2H), 7.00 – 6.90 (m, 1H), 4.33 (q, *J* = 7.2 Hz, 1H), 3.69 – 3.50 (m, 3H), 3.49 – 3.29 (m, 2H), 2.83 (d, *J* = 6.5 Hz, 2H), 2.64 (q, *J* = 7.6 Hz, 2H), 2.42 – 2.18 (m, 3H), 1.18 (t, *J* = 7.6 Hz, 3H).

$^{13}$C-NMR (75 MHz, DMSO): δ [ppm] = 171.1, 170.6, 169.9, 166.1, 147.1, 136.1, 132.0, 127.6, 127.2, 123.4, 120.8, 118.6, 118.2, 111.3, 111.1, 47.2, 41.9, 40.0, 36.1, 35.7, 29.8, 28.0, 15.4.

LC/MS: m/z calculated for $C_{26}H_{31}N_5O_4$ $[M+H]^+$: 478.2, found: 478.1.

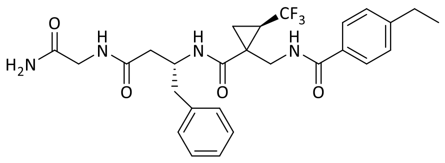

**28**

Compound **28**: Yield: 23.2 mg (0.040 mmol ≙ 87% of th.), colorless solid.

$^1$H-NMR (300 MHz, DMSO-$d_6$): δ [ppm] = 8.51 (s, 1H), 8.15 (d, *J* = 7.5 Hz, 1H), 7.81 – 7.74 (m, 2H), 7.37 – 7.13 (m, 6H), 7.07 (s, 4H), 3.70 (s, 4H), 3.63 – 3.56 (m, 2H), 2.72 – 2.58 (m, 3H), 2.41 – 2.29 (m, 2H), 2.05 – 1.83 (m, 1H), 1.48 – 1.28 (m, 2H), 1.19 (ddd, *J* = 7.5, 5.1, 2.7 Hz, 3H).

$^{13}$C-NMR (75 MHz, DMSO): δ [ppm] = 244.5, 237.0, 214.3, 204.3, 166.2, 148.9, 147.6, 138.2, 131.3, 129.3, 129.1, 128.0, 127.9, 127.7, 127.5, 127.5, 40.3, 40.0, 39.1, 38.8, 38.5, 28.1, 15.4.

LC/MS: m/z calculated for $C_{27}H_{31}F_3N_4O_4$ $[M+H]^+$: 533.2, found: 533.2.

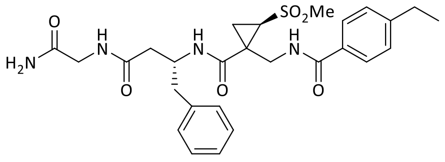

**29**

Compound **29**: Yield: 19.5 mg (0.036 mmol ≙ 72% of th.), colorless solid.

$^1$H-NMR (300 MHz, DMSO-$d_6$): δ [ppm] = 8.58 – 8.41 (m, 1H), 8.26 (t, *J* = 8.8 Hz, 1H), 8.12 (q, *J* = 5.8 Hz, 1H), 7.79 (d, *J* = 7.3 Hz, 2H), 7.36 – 7.09 (m, 6H), 7.04 (s, 3H), 4.33 – 4.16 (m, 5H), 3.83 (ddd, *J* = 30.8, 14.8, 5.2 Hz, 1H), 3.64 – 3.48 (m, 2H), 3.14 (s, 1H), 3.00 – 2.80 (m, 1H), 2.66 (qd, *J* = 7.4, 3.3 Hz, 3H), 2.41 – 2.26 (m, 2H), 1.73 – 1.62 (m, 2H), 1.19 (td, *J* = 7.6, 2.7 Hz, 3H).

$^{13}$C-NMR (75 MHz, DMSO): δ [ppm] = 171.1, 171.1, 170.3, 170.2, 167.8, 167.6, 167.3, 167.2, 147.8, 147.7, 138.6, 138.5, 131.2, 131.2, 129.3, 129.1, 128.1, 127.9, 127.7, 127.6, 127.5, 42.8, 42.8, 32.9, 32.8, 28.1, 15.4.

LC/MS: m/z calculated for $C_{27}H_{34}N_4O_6S$ $[M+H]^+$: 543.2, found: 543.2.

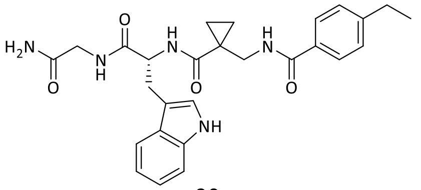

**30**

Compound **30**: Yield: 24.0 mg (0.049 mmol ≙ quant.), colorless solid.

$^{1}$H-NMR (300 MHz, DMSO-$d_6$): δ [ppm] = 10.77 (d, *J* = 2.4 Hz, 1H), 8.63 (t, *J* = 6.1 Hz, 1H), 8.44 (d, *J* = 6.7 Hz, 1H), 8.32 (t, *J* = 5.9 Hz, 1H), 7.80 – 7.67 (m, 2H), 7.60 (d, *J* = 7.7 Hz, 1H), 7.35 – 7.23 (m, 3H), 7.20 (d, *J* = 2.3 Hz, 1H), 7.14 – 7.01 (m, 3H), 6.97 (ddd, *J* = 8.0, 6.9, 1.1 Hz, 1H), 4.41 (dt, *J* = 9.4, 6.3 Hz, 1H), 3.74 – 3.56 (m, 2H), 3.56 – 3.44 (m, 2H), 3.17 (dd, *J* = 14.7, 4.8 Hz, 1H), 3.01 (dd, *J* = 14.7, 9.3 Hz, 1H), 2.65 (q, *J* = 7.6 Hz, 2H), 1.19 (t, *J* = 7.6 Hz, 3H), 0.98 – 0.71 (m, 4H).

$^{13}$C-NMR (75 MHz, DMSO): δ [ppm] = 172.9, 172.1, 171.1, 167.4, 147.5, 136.1, 131.4, 127.7, 127.5, 127.3, 123.6, 120.9, 118.4, 118.3, 111.4, 110.3, 54.8, 42.3, 42.2, 28.1, 27.1, 25.2, 15.4, 13.6.

LC/MS: m/z calculated for $C_{27}H_{31}N_5O_4$ $[M+H]^+$: 490.2, found: 490.1.

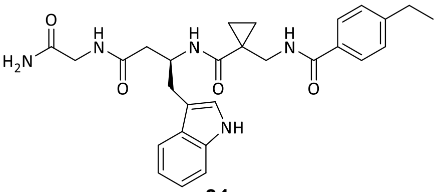

**31**

Compound **31**: Yield: 8.0 mg (0.016 mmol ≙ 35% of th.), colorless solid.

$^{1}$H-NMR (300 MHz, DMSO-$d_6$): δ [ppm] = 10.76 (d, *J* = 2.4 Hz, 1H), 8.57 (t, *J* = 6.1 Hz, 1H), 8.16 – 7.90 (m, 2H), 7.84 – 7.68 (m, 2H), 7.60 (d, *J* = 7.8 Hz, 1H), 7.35 – 7.25 (m, 3H), 7.23 (s, 1H), 7.09 (d, *J* = 2.2 Hz, 1H), 7.07 – 6.98 (m, 2H), 6.98 – 6.88 (m, 1H), 4.34 (q, *J* = 6.9 Hz, 1H), 3.72 – 3.36 (m, 4H), 2.83 (d, *J* = 6.5 Hz, 2H), 2.65 (q, *J* = 7.6 Hz, 2H), 2.34 (d, *J* = 6.8 Hz, 2H), 1.19 (t, *J* = 7.6 Hz, 3H), 1.06 – 0.59 (m, 4H).

$^{13}$C-NMR (75 MHz, DMSO): δ [ppm] = 171.5, 171.1, 170.6, 167.2, 147.5, 136.1, 131.5, 127.7, 127.6, 127.4, 123.3, 120.8, 118.6, 118.2, 111.3, 111.2, 47.7, 42.3, 41.9, 39.9, 29.8, 28.1, 25.3, 15.4, 13.2, 12.9.

LC/MS: m/z calculated for $C_{28}H_{33}N_5O_4$ $[M+H]^+$: 504.3, found: 504.1.

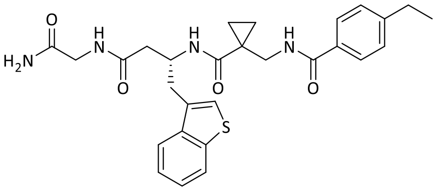

**32**

Compound **32**: Yield: 10.0 mg (0.019 mmol ≙ 43% of th.), colorless solid.

$^{1}$H-NMR (300 MHz, DMSO-$d_6$): δ [ppm] = 8.58 (t, *J* = 6.1 Hz, 1H), 8.22 – 8.05 (m, 2H), 8.02 – 7.93 (m, 1H), 7.93 – 7.85 (m, 1H), 7.81 – 7.68 (m, 2H), 7.47 – 7.18 (m, 6H), 7.03 (s, 1H), 4.42 (q, *J* = 7.0 Hz, 1H), 3.75 – 3.43 (m, 4H), 3.00 (qd, *J* = 14.2, 6.6 Hz, 2H), 2.65 (q, *J* = 7.5 Hz, 2H), 2.41 (d, *J* = 6.9 Hz, 2H), 1.19 (t, *J* = 7.6 Hz, 3H), 0.98 – 0.68 (m, 4H).

$^{13}$C-NMR (75 MHz, DMSO): δ [ppm] = 171.7, 171.0, 170.4, 167.2, 147.5, 139.6, 139.0, 133.2, 131.4, 127.7, 127.4, 124.2, 124.0, 123.3, 122.8, 122.1, 46.7, 42.3, 41.9, 40.2, 32.8, 28.1, 25.3, 15.4, 13.1.

LC/MS: m/z calculated for $C_{28}H_{32}N_4O_4S$ $[M+H]^+$: 521.2, found: 521.1.

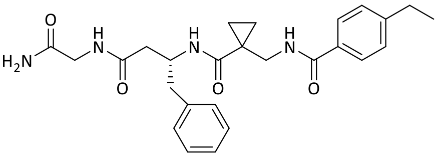

**14**

Compound **14**: Yield: 16.0 mg (0.034 mmol ≙ 77% of th.), colorless solid.

$^{1}$H-NMR (600 MHz, DMSO-$d_6$): δ [ppm] = 8.58 (t, *J* = 6.2 Hz, 1H), 8.11 (t, *J* = 5.9 Hz, 1H), 8.01 (d, *J* = 8.2 Hz, 1H), 7.82 – 7.74 (m, 2H), 7.32 (d, *J* = 8.2 Hz, 2H), 7.25 (s, 1H), 7.15 – 7.12 (m, 4H), 7.12 – 7.07 (m, 1H), 7.04 (s, 1H), 4.33 – 4.22 (m, 1H), 3.61 (dd, *J* = 16.8, 6.0 Hz, 1H), 3.54 – 3.45 (m, 3H), 2.78 (dd, *J* = 13.6, 5.1 Hz, 1H), 2.68 – 2.63 (m, 3H), 2.30 (d, *J* = 6.9 Hz, 2H), 1.19 (t, *J* = 7.6 Hz, 3H), 0.92 – 0.80 (m, 2H), 0.80 – 0.71 (m, 2H).

$^{13}$C-NMR (151 MHz, DMSO): δ [ppm] = 171.5, 171.1, 170.4, 167.2, 147.6, 138.9, 131.4, 129.2, 128.0, 127.7, 127.5, 126.0, 48.5, 42.3, 41.9, 40.3, 40.1, 28.1, 25.3, 15.5, 13.1.

LC/MS: m/z calculated for $C_{26}H_{32}N_4O_4$ $[M+H]^+$: 465.3, found: 465.1.

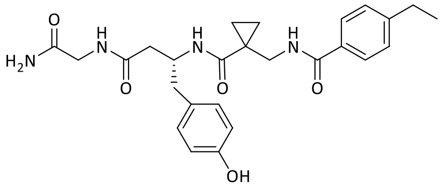

**15**

Compound **15**: Yield: 15.0 mg (0.031 mmol ≙ 69% of th.), colorless solid.

$^{1}$H-NMR (600 MHz, DMSO-$d_6$): δ [ppm] = 9.11 (s, 1H), 8.57 (t, *J* = 6.1 Hz, 1H), 8.05 (t, *J* = 5.8 Hz, 1H), 7.95 (d, *J* = 8.1 Hz, 1H), 7.85 – 7.71 (m, 2H), 7.41 – 7.27 (m, 2H), 7.22 (s, 1H), 7.02 (s, 1H), 6.97 – 6.86 (m, 2H), 6.64 – 6.49 (m, 2H), 4.21 (h, *J* = 7.3 Hz, 1H), 3.69 – 3.44 (m, 4H), 2.74 – 2.53 (m, 4H), 2.27 (d, *J* = 6.8 Hz, 2H), 1.19 (t, *J* = 7.6 Hz, 3H), 0.96 – 0.60 (m, 4H).

$^{13}$C-NMR (151 MHz, DMSO): δ [ppm] = 171.5, 171.1, 170.4, 167.2, 155.5, 147.6, 131.4, 130.0, 128.9, 127.7, 127.5, 114.8, 48.7, 42.3, 41.9, 40.2, 38.8, 28.1, 25.2, 15.4, 13.0.

LC/MS: m/z calculated for $C_{26}H_{32}N_4O_5$ $[M+H]^+$: 481.2, found: 481.1.

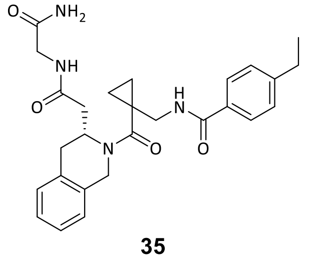

**35**

Compound **35**: Yield: 11.0 mg (0.023 mmol ≙ 50% of th.), colorless solid.

$^{1}$H-NMR (400 MHz, DMSO-$d_6$): δ [ppm] = 8.42 (t, *J* = 6.1 Hz, 1H), 8.08 (t, *J* = 6.0 Hz, 1H), 7.70 (d, *J* = 7.8 Hz, 2H), 7.26 (d, *J* = 7.9 Hz, 2H), 7.24 – 7.18 (m, 1H), 7.18 – 7.08 (m, 4H), 7.00 (s, 1H), 5.39 – 4.83 (m, 2H), 4.58 (s, 1H), 3.78 – 3.50 (m, 3H), 2.82 – 2.67 (m, 1H), 2.63 (q, *J* = 7.6 Hz, 3H), 2.45 – 2.23 (m, 2H), 1.18 (t, *J* = 7.6 Hz, 3H), 0.93 – 0.49 (m, 4H).

$^{13}$C-NMR (101 MHz, DMSO): δ [ppm] = 170.9, 170.2, 170.1, 166.5, 147.2, 131.8, 127.6, 127.3, 126.6, 126.0, 41.9, 37.0, 32.0, 28.0, 26.0, 15.4, 9.8, 9.5.

LC/MS: m/z calculated for $C_{27}H_{32}N_4O_4$ $[M+H]^+$: 477.3, found: 477.1.

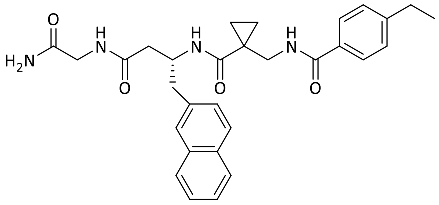

**36**

Compound **36**: Yield: 19.0 mg (0.037 mmol ≙ 80% of th.), colorless solid.

$^{1}$H-NMR (300 MHz, DMSO-$d_6$): δ [ppm] = 8.59 (t, *J* = 6.1 Hz, 1H), 8.23 – 8.03 (m, 2H), 7.88 – 7.72 (m, 3H), 7.69 (d, *J* = 8.4 Hz, 1H), 7.64 – 7.49 (m, 2H), 7.50 – 7.36 (m, 2H), 7.36 – 7.27 (m, 3H), 7.25 (s, 1H), 7.04 (s, 1H), 4.41 (p, *J* = 6.9 Hz, 1H), 3.77 – 3.46 (m, 3H), 3.46 – 3.26 (m, 1H), 2.98 (dd, *J* = 13.6, 4.9 Hz, 1H), 2.91 – 2.75 (m, 1H), 2.67 (q, *J* = 7.6 Hz, 2H), 2.43 – 2.28 (m, 2H), 1.21 (t, *J* = 7.6 Hz, 3H), 0.98 – 0.66 (m, 4H).

$^{13}$C-NMR (75 MHz, DMSO): δ [ppm] = 171.5, 171.1, 170.3, 167.2, 147.6, 136.6, 132.9, 131.6, 131.3, 127.9, 127.7, 127.5, 127.4, 127.4, 127.3, 127.2, 125.8, 125.2, 48.4, 42.3, 41.9, 40.5, 39.7, 28.1, 25.3, 15.4, 13.3, 13.0.

LC/MS: m/z calculated for $C_{30}H_{34}N_4O_4$ $[M+H]^+$: 515.3, found: 515.2.

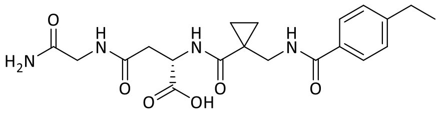

**37**

Compound **37**: Yield: 5.0 mg (0.012 mmol ≙ 26% of th.), colorless solid.

$^{1}$H-NMR (600 MHz, DMSO-$d_6$): δ [ppm] = 8.60 (t, *J* = 6.1 Hz, 1H), 8.26 – 8.20 (m, 2H), 7.81 – 7.74 (m, 2H), 7.35 (s, 1H), 7.30 (d, *J* = 8.1 Hz, 2H), 7.08 (s, 1H), 4.49 (q, *J* = 7.0 Hz, 1H), 3.62 (dd, *J* = 16.9, 6.1 Hz, 1H), 3.58 – 3.50 (m, 2H), 3.47 (dd, *J* = 16.9, 5.6 Hz, 1H), 2.71 – 2.60 (m, 3H), 1.19 (t, *J* = 7.6 Hz, 4H), 0.99 – 0.91 (m, 2H), 0.87 – 0.78 (m, 2H).

$^{13}$C-NMR (151 MHz, DMSO): δ [ppm] = 173.1, 172.2, 172.1, 171.2, 169.6, 167.3, 147.5, 131.5, 127.7, 127.6, 49.8, 42.1, 42.1, 40.1, 37.1, 28.1, 24.9, 15.5, 13.3, 13.0.

LC/MS: m/z calculated for $C_{20}H_{26}N_4O_6$ $[M+H]^+$: 419.2, found: 419.1.

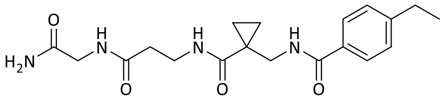

**38**

Compound **38**: Yield: 5.0 mg (0.013 mmol ≙ 29% of th.), colorless solid.

$^{1}$H-NMR (600 MHz, DMSO-$d_6$): δ [ppm] = 8.60 (t, *J* = 6.1 Hz, 1H), 8.10 (dt, *J* = 15.2, 5.6 Hz, 2H), 7.82 – 7.75 (m, 2H), 7.36 – 7.26 (m, 3H), 7.05 (s, 1H), 3.58 (d, *J* = 5.8 Hz, 2H), 3.51 (d, *J* = 6.0 Hz, 2H), 3.28 (td, *J* = 7.2, 5.3 Hz, 2H), 2.65 (q, *J* = 7.6 Hz, 2H), 2.30 (t, *J* = 7.3 Hz, 2H), 1.18 (t, *J* = 7.6 Hz, 3H), 0.95 (q, *J* = 3.7 Hz, 2H), 0.81 (q, *J* = 3.7 Hz, 2H).

$^{13}$C-NMR (151 MHz, DMSO): δ [ppm] = 172.2, 171.2, 170.8, 167.2, 147.6, 131.5, 127.8, 127.5, 42.2, 41.9, 40.1, 36.0, 35.2, 28.1, 25.1, 15.5, 13.1.

LC/MS: m/z calculated for $C_{19}H_{26}N_4O_4$ $[M+H]^+$: 375.2, found: 375.1.

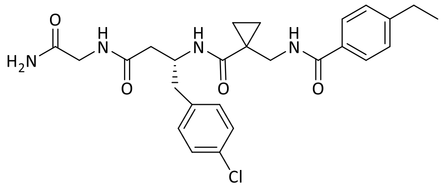

**16**

Compound **16**: Yield: 21.0 mg (0.042 mmol ≙ 26% of th.), colorless solid.

$^{1}$H-NMR (300 MHz, DMSO-$d_6$): δ [ppm] = 8.58 (t, *J* = 6.1 Hz, 1H), 8.11 (t, *J* = 5.8 Hz, 1H), 8.01 (d, *J* = 8.1 Hz, 1H), 7.77 (d, *J* = 8.2 Hz, 2H), 7.31 (d, *J* = 8.2 Hz, 2H), 7.25 (s, 1H), 7.13 (s, 4H), 7.03 (s, 1H), 4.28 (q, *J* = 7.8 Hz, 1H), 3.68 – 3.40 (m, 4H), 2.82 (dd, *J* = 13.6, 4.8 Hz, 1H), 2.74 – 2.60 (m, 3H), 2.32 (d, *J* = 6.9 Hz, 2H), 1.20 (t, *J* = 7.6 Hz, 3H), 0.93 – 0.72 (m, 4H).

$^{13}$C-NMR (75 MHz, DMSO): δ [ppm] = 171.6, 171.1, 170.3, 167.2, 147.6, 137.8, 131.4, 131.0, 130.6, 127.8, 127.7, 127.5, 54.9, 48.2, 42.3, 41.9, 40.2, 38.6, 28.1, 25.3, 15.4, 13.2, 13.1.

LC/MS: m/z calculated for $C_{26}H_{31}ClN_4O_4$ $[M+H]^+$: 499.2, found: 498.9.

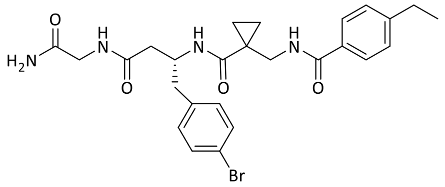

**17**

Compound **17**: Yield: 29.1 mg (0.053 mmol ≙ 34% of th.), colorless solid.

$^1$H-NMR (300 MHz, DMSO-$d_6$): δ [ppm] = 8.59 (t, $J$ = 6.1 Hz, 1H), 8.12 (t, $J$ = 5.9 Hz, 1H), 8.01 (d, $J$ = 8.1 Hz, 1H), 7.77 (d, $J$ = 8.3 Hz, 2H), 7.35 – 7.29 (m, 2H), 7.28 – 7.20 (m, 3H), 7.11 – 6.99 (m, 3H), 4.28 (td, $J$ = 7.8, 4.8 Hz, 1H), 3.69 – 3.47 (m, 3H), 3.42 (d, $J$ = 5.9 Hz, 1H), 2.80 (dd, $J$ = 13.6, 4.8 Hz, 1H), 2.65 (p, $J$ = 7.6 Hz, 3H), 2.32 (d, $J$ = 6.9 Hz, 2H), 1.20 (t, $J$ = 7.6 Hz, 3H), 0.95 – 0.71 (m, 4H).

$^{13}$C-NMR (75 MHz, DMSO): δ [ppm] = 172.0, 171.5, 170.7, 167.6, 148.0, 138.7, 131.8, 131.2, 128.2, 127.9, 119.6, 48.6, 42.8, 42.4, 28.5, 25.7, 15.9, 13.7, 13.5.

LC/MS: m/z calculated for $C_{26}H_{31}BrN_4O_4$ $[M+H]^+$: 543.1, found: 543.1.

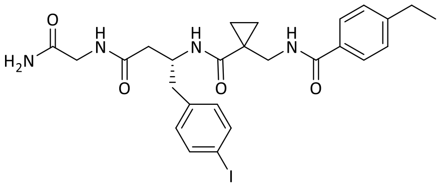

**33**

Compound **33**: Yield: 32.1 mg (0.054 mmol ≙ 34% of th.), colorless solid.

$^1$H-NMR (300 MHz, DMSO-$d_6$): δ [ppm] = 8.59 (t, $J$ = 6.1 Hz, 1H), 8.11 (t, $J$ = 5.9 Hz, 1H), 8.00 (d, $J$ = 8.1 Hz, 1H), 7.77 (d, $J$ = 8.3 Hz, 2H), 7.46 – 7.37 (m, 2H), 7.36 – 7.28 (m, 2H), 7.25 (s, 1H), 7.03 (s, 1H), 6.93 (d, $J$ = 8.3 Hz, 2H), 4.27 (q, $J$ = 7.2 Hz, 1H), 3.74 – 3.47 (m, 4H), 2.78 (dd, $J$ = 13.7, 4.8 Hz, 1H), 2.72 – 2.56 (m, 3H), 2.31 (d, $J$ = 6.9 Hz, 2H), 1.21 (t, $J$ = 7.6 Hz, 3H), 0.95 – 0.71 (m, 4H).

$^{13}$C-NMR (75 MHz, DMSO): δ [ppm] = 172.0, 171.5, 170.7, 167.6, 148.0, 139.0, 137.1, 132.0, 131.8, 128.2, 127.9, 92.2, 48.6, 42.8, 42.4, 40.7, 28.6, 25.7, 15.9, 13.7, 13.5.

LC/MS: m/z calculated for $C_{26}H_{31}ClN_4O_4$ $[M+H]^+$: 591.1, found: 591.1.

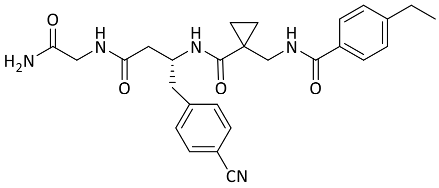

**34**

Compound **34**: Yield: 26.0 mg (0.044 mmol ≙ 28% of th.), colorless solid.

$^1$H-NMR (300 MHz, DMSO-$d_6$): δ [ppm] = 8.59 (t, $J$ = 6.1 Hz, 1H), 8.15 (t, $J$ = 5.9 Hz, 1H), 8.06 (d, $J$ = 8.1 Hz, 1H), 7.74 (d, $J$ = 8.3 Hz, 2H), 7.50 (d, $J$ = 8.3 Hz, 2H), 7.30 (t, $J$ = 7.9 Hz, 5H), 7.03 (s, 1H), 4.33 (dq, $J$ = 11.7, 6.7 Hz, 1H), 3.68 – 3.49 (m, 3H), 3.33 (d, $J$ = 5.9 Hz, 1H), 2.95 (dd,

$J$ = 13.6, 4.6 Hz, 1H), 2.81 – 2.60 (m, 3H), 2.35 (d, $J$ = 7.0 Hz, 2H), 1.21 (t, $J$ = 7.6 Hz, 3H), 0.94 – 0.70 (m, 4H).

$^{13}$C-NMR (75 MHz, DMSO): δ [ppm] = 172.1, 171.5, 170.6, 167.6, 148.1, 145.3, 132.2, 131.7, 130.6, 128.1, 127.9, 119.4, 109.2, 48.3, 42.8, 42.3, 40.7, 28.5, 25.7, 15.8, 13.7, 13.5.

LC/MS: m/z calculated for $C_{27}H_{31}N_5O_4$ $[M+H]^+$: 490.2, found: 490.2.

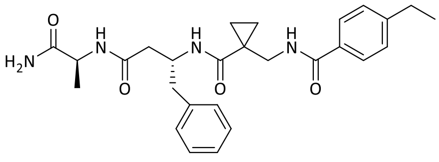

**39**

Compound **39**: Yield: 14 mg (0.029 mmol ≙ 64% of th.), colorless solid.

$^{1}$H-NMR (300 MHz, DMSO-$d_6$): δ [ppm] = 8.53 (t, $J$ = 6.1 Hz, 1H), 8.03 – 7.90 (m, 2H), 7.85 – 7.69 (m, 2H), 7.39 – 7.28 (m, 2H), 7.26 (s, 1H), 7.20 – 7.01 (m, 5H), 6.96 (s, 1H), 4.29 (h, $J$ = 7.4 Hz, 1H), 4.16 (p, $J$ = 7.1 Hz, 1H), 3.48 (p, $J$ = 8.5 Hz, 2H), 2.78 (dd, $J$ = 13.5, 5.2 Hz, 1H), 2.72 – 2.58 (m, 3H), 2.28 (d, $J$ = 6.9 Hz, 2H), 1.24 – 1.11 (m, 6H), 0.92 – 0.71 (m, 4H).

$^{13}$C-NMR (75 MHz, DMSO): δ [ppm] = 174.4, 171.5, 169.7, 167.2, 147.6, 138.8, 131.5, 129.1, 128.0, 127.7, 127.5, 126.0, 48.5, 48.0, 42.2, 28.1, 25.2, 18.2, 15.4, 13.0, 12.9.

LC/MS: m/z calculated for $C_{27}H_{34}N_4O_4$ $[M+H]^+$: 479.3, found: 479.2.

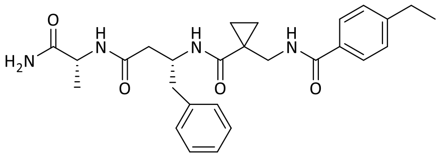

**40**

Compound **40**: Yield: 3.0 mg (6.263 µmol ≙ 14% of th.), colorless solid.

$^{1}$H-NMR (400 MHz, DMSO-$d_6$): δ [ppm] = 8.54 (t, $J$ = 6.0 Hz, 1H), 8.01 – 7.90 (m, 2H), 7.77 (d, $J$ = 7.8 Hz, 2H), 7.31 (d, $J$ = 7.8 Hz, 2H), 7.25 (s, 1H), 7.18 – 7.02 (m, 5H), 6.95 (s, 1H), 4.28 (q, $J$ = 7.0 Hz, 1H), 4.16 (p, $J$ = 7.2 Hz, 1H), 3.62 – 3.51 (m, 1H), 3.44 – 3.34 (m, 1H), 2.83 – 2.60 (m, 4H), 2.30 (d, $J$ = 6.9 Hz, 2H), 1.28 – 1.09 (m, 6H), 0.95 – 0.67 (m, 4H).

$^{13}$C-NMR (101 MHz, DMSO): δ [ppm] = 174.3, 171.4, 169.7, 167.2, 147.6, 138.8, 131.4, 129.1, 127.9, 127.7, 127.5, 125.9, 48.4, 48.0, 42.3, 40.2, 28.1, 25.2, 18.2, 15.4, 13.0, 12.9.

LC/MS: m/z calculated for $C_{27}H_{34}N_4O_4$ $[M+H]^+$: 479.3, found: 479.1.

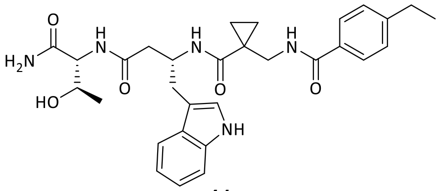

**41**

Compound **41**: Yield: 7.0 mg (0.013 mmol ≙ 28% of th.), colorless solid.

$^{1}$H-NMR (300 MHz, DMSO-$d_6$): δ [ppm] = 10.75 (d, *J* = 2.4 Hz, 1H), 8.56 (t, *J* = 6.1 Hz, 1H), 8.00 (d, *J* = 7.9 Hz, 1H), 7.86 (d, *J* = 8.6 Hz, 1H), 7.82 – 7.68 (m, 3H), 7.58 (d, *J* = 7.8 Hz, 1H), 7.37 – 7.19 (m, 5H), 7.12 – 6.98 (m, 3H), 6.98 – 6.88 (m, 1H), 4.35 (q, *J* = 6.9 Hz, 1H), 4.21 (dd, *J* = 8.6, 6.1 Hz, 1H), 3.84 (p, *J* = 6.3 Hz, 1H), 2.82 (d, *J* = 6.5 Hz, 2H), 2.65 (q, *J* = 7.6 Hz, 2H), 2.38 (d, *J* = 6.9 Hz, 2H), 1.19 (t, *J* = 7.6 Hz, 4H), 1.03 (d, *J* = 6.3 Hz, 3H), 0.93 – 0.86 (m, 2H), 0.81 – 0.73 (m, 2H).

$^{13}$C-NMR (75 MHz, DMSO): δ [ppm] = 172.3, 171.6, 170.5, 167.2, 147.5, 136.1, 131.5, 127.7, 127.7, 127.6, 127.4, 127.4, 123.2, 120.8, 118.6, 118.2, 111.3, 111.2, 66.8, 58.1, 47.9, 42.2, 40.2, 29.7, 28.1, 25.3, 19.5, 15.4, 13.3, 13.0.

LC/MS: m/z calculated for $C_{30}H_{37}N_5O_5$ $[M+H]^+$: 548.3, found: 547.8.

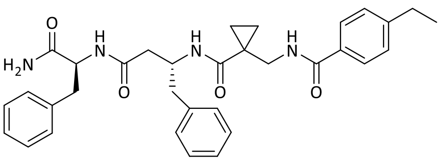

**42**

Compound **42**: Yield: 15.0 mg (0.027 mmol ≙ 59% of th.), colorless solid.

$^{1}$H-NMR (300 MHz, DMSO-$d_6$): δ [ppm] = 8.54 (t, *J* = 6.1 Hz, 1H), 8.14 (d, *J* = 8.4 Hz, 1H), 7.98 (d, *J* = 8.2 Hz, 1H), 7.87 – 7.71 (m, 2H), 7.39 (s, 1H), 7.31 (d, *J* = 8.2 Hz, 2H), 7.27 – 7.22 (m, 3H), 7.22 – 6.95 (m, 8H), 4.55 – 4.35 (m, 1H), 4.24 (q, *J* = 7.3 Hz, 1H), 3.59 – 3.42 (m, 2H), 3.02 (dd, *J* = 13.8, 4.7 Hz, 1H), 2.84 – 2.42 (m, 5H), 2.23 (qd, *J* = 14.2, 6.9 Hz, 2H), 1.19 (t, *J* = 7.6 Hz, 3H), 0.97 – 0.51 (m, 4H).

$^{13}$C-NMR (75 MHz, DMSO): δ [ppm] = 173.3, 171.5, 170.0, 167.2, 147.6, 138.8, 138.3, 131.5, 129.1, 128.1, 127.9, 127.7, 127.5, 126.2, 125.9, 53.9, 48.4, 42.3, 37.5, 28.1, 25.2, 15.4, 13.0, 13.0.

LC/MS: m/z calculated for $C_{33}H_{38}N_4O_4$ $[M+H]^+$: 555.3, found: 555.2.

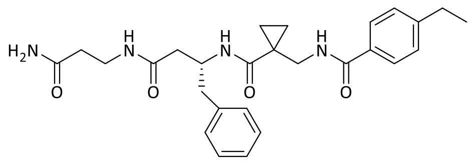

**43**

Compound **43**: Yield: 18.0 mg (0.038 mmol ≙ 82% of th.), colorless solid.

$^1$H-NMR (300 MHz, DMSO-$d_6$): δ [ppm] = 8.57 (t, *J* = 6.1 Hz, 1H), 8.05 – 7.84 (m, 2H), 7.85 – 7.70 (m, 2H), 7.40 – 7.24 (m, 3H), 7.22 – 7.01 (m, 5H), 6.82 (s, 1H), 4.26 (h, *J* = 7.1 Hz, 1H), 3.61 – 3.39 (m, 2H), 3.29 – 3.06 (m, 2H), 2.81 – 2.57 (m, 4H), 2.32 – 2.04 (m, 4H), 1.19 (t, *J* = 7.6 Hz, 3H), 0.98 – 0.69 (m, 4H).

$^{13}$C-NMR (75 MHz, DMSO): δ [ppm] = 172.6, 171.4, 169.9, 167.2, 147.6, 138.8, 131.4, 129.2, 128.0, 127.7, 127.5, 125.9, 48.4, 42.2, 40.2, 35.2, 35.0, 28.1, 25.2, 15.4, 13.0.

LC/MS: m/z calculated for $C_{27}H_{34}N_4O_4$ $[M+H]^+$: 479.3, found: 479.2.

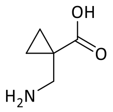

**49**

Compound **49**: 1-Cyanocyclopropane-1-carboxylic acid (500 mg, 4.500 mmol, 1.0 eq.) was dissolved in methanol (10 mL), ammonia in methanol (7 M, 1.0 mL), and Raney-nickel (50 mg, 5 w%) were added. The mixture was stirred under a hydrogen atmosphere (4 bar) for 16 h at RT. The reaction mixture was adjusted to alkaline pH (pH = 8) by adding ammonia in methanol (7 M, 0.5 mL), filtered over Celite S, and the solvent was removed under reduced pressure.

Yield: 574.0 mg (4.986 mmol ≙ quant.), colorless solid.

$^1$H-NMR (300 MHz, MeO*D*): δ [ppm] = 3.06 (s, 2H), 1.36 – 1.29 (m, 2H), 1.02 – 0.94 (m, 2H).

$^{13}$C-NMR (75 MHz, MeOD): δ [ppm] = 177.8, 101.2, 45.2, 15.0.

LC/MS: m/z calculated for $C_5H_9NO_2$ $[M+H]^+$: 116.1, found: 116.0.

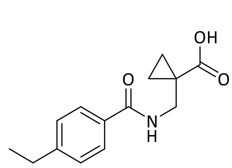

**50**

Compound **50**: The crude product **49** (518 mg, 4.500 mmol, 1.0 eq.) was suspended in DCM (10 mL), DMF (0.5 mL), and TEA (620 µL, 4.473 mmol, 1.0 eq.). The mixture was cooled to 0 °C,

and a solution of acid 4-ethylbenzoyl chloride (662 µL, 4.500 mmol, 1.0 eq.) in DCM (0.3 mL) was added. The mixture was warmed to RT and stirred for 16 h. The reaction mixture was washed with 1 M hydrochloric acid thrice (5 mL each). The organic phase was separated, dried over sodium sulfate, and the solvents were removed under reduced pressure. The crude product was purified by column chromatography (mobile phase: CH:EtOAc 2:1 with 0.1% TFA)
Yield: 413.0 mg (1.670 mmol ≙ 37% of th.), colorless solid.

$^{1}$H-NMR (300 MHz, DMSO-$d_6$): δ [ppm] = 9.99 (s, 1H), 7.80 – 7.62 (m, 2H), 7.29 – 7.20 (m, 2H), 7.00 (t, *J* = 6.2 Hz, 1H), 3.62 (d, *J* = 6.1 Hz, 2H), 2.68 (q, *J* = 7.6 Hz, 2H), 1.35 (q, *J* = 4.2 Hz, 2H), 1.24 (t, *J* = 7.6 Hz, 3H), 1.15 (q, *J* = 4.3 Hz, 2H).

$^{13}$C-NMR (75 MHz, DMSO): δ [ppm] = 181.1, 167.9, 148.5, 131.6, 128.2, 127.3, 42.9, 28.9, 24.8, 16.0, 15.5.

LC/MS: m/z calculated for $C_{14}H_{17}NO_3$ $[M+H]^+$: 248.1, found: 248.0.

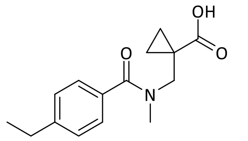

**51**

Compound **51**: Compound **50** (50 mg, 0.202 mmol, 1.0 eq.) was dissolved in anhydrous THF (5 mL) and cooled to 0 °C. Sodium hydride (60% suspension, 21 mg, 0.525 mmol, 2.6 eq.) was added and the mixture was stirred for 5 min. Then, methyl iodide (38 µL, 0.610 mmol, 3.0 eq.) was added dropwise. The mixture was warmed to RT and stirred for 16 h. A solution of sodium hydroxide (18 mg, 0.450 mmol, 2.2 eq.) in water (5 mL) was added and the mixture was stirred for 1 h. The mixture was adjusted to acidic pH (pH = 1) with 1 M hydrochloric acid and the solvents were removed under reduced pressure. The crude product was dissolved in sodium hydroxide solution (pH = 12) and was three times extracted with DCM (3 mL each). Then, the aqueous phase was brought to an acidic pH (pH = 1) with 1 M hydrochloric acid and extracted thrice with DCM (3 mL each). The organic phases from the acidic extraction were combined, and dried over sodium sulfate, and the solvent was removed under reduced pressure.
Yield: 44.0 mg (0.168 mmol ≙ 83% of th.), colorless solid.

$^{1}$H-NMR (300 MHz, *CD*$Cl_3$, 60 °C): δ [ppm] = 9.87 (s, 1H), 7.30 (d, *J* = 8.2 Hz, 2H), 7.19 (d, *J* = 7.9 Hz, 2H), 3.82 (s, 2H), 3.05 (s, 3H), 2.66 (q, *J* = 7.6 Hz, 2H), 1.39 (q, *J* = 4.1 Hz, 2H), 1.24 (t, *J* = 7.6 Hz, 3H), 1.15 – 0.94 (m, 2H).

$^{13}$C-NMR (75 MHz, CDCl$_3$, 60 °C): δ [ppm] = 179.1, 173.0, 146.2, 133.7, 128.1, 127.9, 127.4, 127.3, 28.8, 23.2, 16.2, 15.3.

LC/MS: m/z calculated for $C_{15}H_{19}NO_3$[M+H]$^+$: 262.2, found: 262.0.

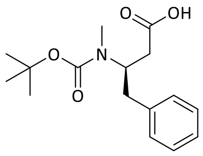

**52**

Compound **52**: (*R*)-3-*tert*-butoxycarbonylamino-4-phenylbutyric acid (100 mg, 0.358 mmol, 1.0 eq.) was dissolved in anhydrous THF (7 mL) and cooled to 0 °C. Sodium hydride (60% suspension, 39 mg, 0.975 mmol, 2.7 eq.) was added and the mixture was stirred for 5 min. Then, methyl iodide (67 µL, 1.074 mmol, 3.0 eq.) was added dropwise. The mixture was warmed to RT and stirred for 16 h. Because the reaction was not complete, methyl iodide (46 µL, 0.739 mmol, 2.1 eq.) and sodium hydride (34 mg, 0.850 mmol, 2.4 eq.) were added and the mixture was stirred for an additional 16 h. Water (2 mL) was added, and the pH was confirmed to be >12. The mixture was stirred for 1 h and all solvents were removed under reduced pressure and the crude product was purified by reversed-phase flash column chromatography (Biotage Isolera, ACN/$H_2O$ + TFA).

Yield: 39.0 mg (0.133 mmol ≙ 37% of th.), colorless solid.

$^1$H-NMR (300 MHz, CDCl$_3$, 60 °C): δ [ppm] = 8.92 (s, 1H), 7.28 – 6.94 (m, 5H), 4.45 (tt, *J* = 8.9, 6.1 Hz, 1H), 2.93 – 2.52 (m, 5H), 2.45 (dd, *J* = 15.5 Hz, 5.8 Hz, 1H), 1.27 (s, 9H).

$^{13}$C-NMR (75 MHz, CDCl$_3$, 60 °C): δ [ppm] = 176.1, 155.9, 138.2, 129.2, 128.7, 126.8, 80.3, 55.7, 38.9, 37.6, 30.8, 28.5.

LC/MS: m/z calculated for $C_{16}H_{23}NO_4$ [M-Boc+H]$^+$: 194.1, found: 194.0.

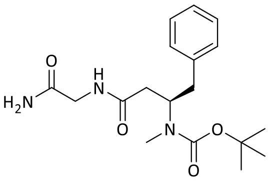

**53**

Compound **53**: Compound **52** (35 mg, 0.119 mmol, 1.0 eq.), glycine amide hydrochloride (14 mg, 0.127 mmol, 1.1 eq.), and TBTU (38 mg, 0.119 mmol, 1.0 eq.) were dissolved in anhydrous DMF (1.5 mL), and DIPEA (63 µL, 0.357 mmol, 3.0 eq.) was added. The mixture was stirred for 16 h at RT. All solvents were removed under reduced pressure, and the crude product was purified by reversed-phase flash column chromatography (Biotage Isolera, ACN/$H_2O$ + TFA).

Yield: 33.0 mg (0.094 mmol ≙ 79% of th.), colorless solid.

$^1$H-NMR (300 MHz, C*D*$Cl_3$): δ [ppm] = 7.37 – 6.99 (m, 6H), 6.85 (s, 1H), 5.92 (s, 1H), 4.62 (s, 1H), 4.17 – 3.65 (m, 2H), 3.11 (s, 1H), 2.97 – 2.54 (m, 5H), 2.55 – 2.34 (m, 1H), 1.33 (s, 9H).

$^{13}$C-NMR (75 MHz, C*D*$Cl_3$): δ [ppm] = 138.0, 129.1, 128.6, 126.7, 80.2, 55.6, 43.0, 38.9, 30.6, 28.4.

LC/MS: m/z calculated for $C_{18}H_{27}N_3O_4$ $[M\text{-}Boc+H]^+$: 250.2, found: 250.1.

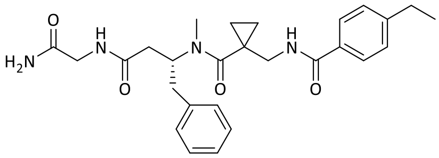

**45**

Compound **45**: Compound **53** (26 mg, 0.074 mmol, 1.0 eq.) was dissolved in DCM (1 mL) and TFA (1 mL) was added. The mixture was stirred at RT for 1.5 h. The solvents were removed under reduced pressure. The deprotected crude product (14 mg, 0.127 mmol, 1.1 eq.), **51** (18 mg, 0.074 mmol, 1.0 eq.), and TBTU (25 mg, 0.078 mmol, 1.1 eq.) were dissolved in anhydrous DMF (1.5 mL), and DIPEA (39 µL, 0.224 mmol, 3.0 eq.) was added. The mixture was stirred for 16 h at RT. The solvents were removed under reduced pressure and the crude product was purified by reversed-phase flash column chromatography (Biotage Isolera, ACN/$H_2O$ + TFA).

Yield: 32.0 mg (0.669 mmol ≙ 90% of th.), colorless solid.

$^1$H-NMR (300 MHz, DMSO-$d_6$): δ [ppm] = 8.34 – 8.08 (m, 2H), 7.71 (d, *J* = 7.9 Hz, 2H), 7.31 – 7.08 (m, 8H), 7.03 (s, 1H), 4.82 (s, 1H), 3.60 (s, 2H), 3.35 (s, 2H), 3.08 – 2.70 (m, 5H), 2.63 (q, *J* = 7.6 Hz, 2H), 2.57 – 2.33 (m, 2H), 1.18 (t, *J* = 7.6 Hz, 3H), 0.70 – 0.28 (m, 4H).

$^{13}$C-NMR (75 MHz, DMSO): δ [ppm] = 171.2, 171.0, 170.5, 166.4, 147.3, 138.6, 131.8, 128.9, 128.1, 127.6, 127.3, 126.1, 41.9, 41.5, 37.4, 37.1, 30.7, 28.0, 26.0, 15.4, 9.5.

LC/MS: m/z calculated for $C_{27}H_{34}N_4O_4$ $[M+H]^+$: 479.3, found: 479.1.

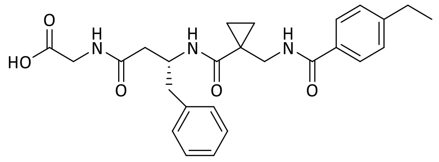

**54**

Compound **54**: Yield: 48.0 mg (0.103 mmol ≙ 76% of th.): colorless solid.

$^{1}$H-NMR (300 MHz, DMSO-$d_6$): δ [ppm] = 12.51(s, 1H), 8.57 (t, *J* = 6.0 Hz, 1H), 8.24 (t, *J* = 5.9 Hz, 1H), 7.99 (d, *J* = 8.0 Hz, 1H), 7.78 (d, *J* = 7.9 Hz, 2H), 7.32 (d, *J* = 7.9 Hz, 2H), 7.23 – 6.96 (m, 5H), 4.41 – 4.10 (m, 1H), 3.88 – 3.56 (m, 2H), 3.58 – 3.16 (m, 2H), 2.82 (dd, *J* = 13.5, 5.0 Hz, 1H), 2.66 (q, *J* = 7.6 Hz, 3H), 2.40 – 2.14 (m, 2H), 1.20 (t, *J* = 7.5 Hz, 3H), 0.97 – 0.59 (m, 4H).

$^{13}$C-NMR (75 MHz, DMSO): δ [ppm] = 171.3, 170.4, 167.2, 147.6, 131.4, 138.9, 129.1, 128.0, 125.9, 127.7, 127.5, 48.5, 42.3, 39.8, 39.1, 28.1, 25.2, 15.4, 13.1.

LC/MS: m/z calculated for $C_{26}H_{31}N_3O_5$ $[M+H]^+$: 466.2, found: 466.1.

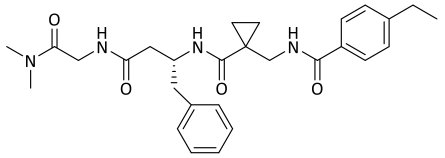

**47**

Compound **47**: Compound **54** (10 mg, 0.022 mmol, 1.0 eq.), dimethylamine hydrochloride (2.9 mg, 0.036 mmol, 1.7 eq.), DCC (5.6 mg, 0.027 mmol, 1.3 eq.) and HOBt (3.6 mg, 0.027 mmol, 1.3 eq.) were dissolved in DCM (1 mL). DIPEA (16 µL, 0.092 µmol, 4.3 eq.) was added dropwise and the mixture was stirred for 16 h at RT. More solvent (DCM, 1 mL), dimethylamine hydrochloride (2.6 mg, 0.032 mmol, 1.5 eq.), DCC (5 mg, 0.024 mmol, 1.1 eq.), and DIPEA (11.7 µL, 0.067 mmol, 2.5 eq.) were added and the mixture was stirred for additional 16 h at RT because the reaction was not completed. The solvents were removed under reduced pressure and the crude product was purified by reversed-phase column chromatography (Biotage Isolera, ACN/$H_2O$ + TFA).

Yield: 8.0 mg (0.016 mmol ≙ 76% of th.), colorless solid.

$^{1}$H-NMR (300 MHz, DMSO-$d_6$): δ [ppm] = 8.54 (t, *J* = 6.0 Hz, 1H), 8.00 – 7.89 (m, 2H), 7.81 – 7.73 (m, 2H), 7.35 – 7.25 (m, 2H), 7.17 – 7.06 (m, 5H), 4.25 (p, *J* = 7.1 Hz, 1H), 3.96 – 3.66 (m, 2H), 3.47 (qd, *J* = 14.8, 6.1 Hz, 2H), 2.89 (s, 3H), 2.80 (s, 3H), 2.74 – 2.59 (m, 2H), 2.41 – 2.22 (m, 2H), 1.19 (t, *J* = 7.6 Hz, 3H), 0.95 – 0.67 (m, 4H).

$^{13}$C-NMR (75 MHz, DMSO): δ [ppm] = 171.4, 170.1, 168.1, 167.1, 147.5, 138.9, 131.4, 129.2, 128.0, 127.6, 127.5, 125.9, 48.5, 42.3, 40.5, 39.1, 35.6, 35.1, 28.1, 25.2, 15.4, 13.1, 13.0.

LC/MS: m/z calculated for $C_{28}H_{36}N_4O_4$ $[M+H]^+$: 493.3, found: 493.2.

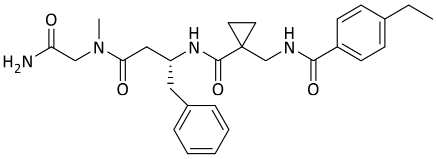

**46**

Compound **46**: Yield: 16.0 mg (0.033 mmol ≙ 74% of th.), colorless solid.

$^{1}$H-NMR (300 MHz, DMSO-$d_6$, 60 °C): δ [ppm] = 8.41 – 8.23 (m, 1H), 7.93 – 7.79 (m, 1H), 7.79 – 7.68 (m, 2H), 7.36 – 7.25 (m, 2H), 7.24 – 7.00 (m, 6H), 4.33 (q, $J$ = 7.3 Hz, 1H), 4.04 – 3.69 (m, 2H), 3.62 – 3.34 (m, 2H), 2.99 – 2.72 (m, 4H), 2.72 – 2.60 (m, 2H), 2.60 – 2.27 (m, 2H), 1.21 (t, $J$ = 7.6 Hz, 3H), 1.02 – 0.63 (m, 4H).

$^{13}$C-NMR (75 MHz, DMSO, 60 °C): δ [ppm] = 171.3, 170.3, 170.0, 166.9, 147.2, 138.8, 131.4, 128.8, 127.7, 127.3, 127.1, 125.6, 50.0, 48.2, 48.0, 42.2, 37.2, 35.9, 27.7, 25.0, 14.9, 12.6.

LC/MS: m/z calculated for $C_{27}H_{34}N_4O_4$ $[M+H]^+$: 479.3, found: 479.2.

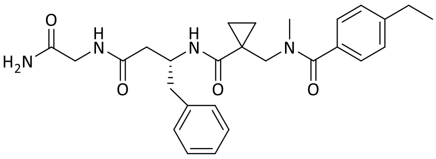

**44**

Compound **44**: Yield: 17.0 mg (0.036 mmol ≙ 79% of th.), colorless solid.

$^{1}$H-NMR (300 MHz, DMSO-$d_6$, 60 °C): δ [ppm] = 7.85 (t, $J$ = 5.7 Hz, 1H), 7.53 (s, 1H), 7.28 – 7.10 (m, 10H), 4.34 (h, $J$ = 7.1 Hz, 1H), 3.80 – 3.45 (m, 4H), 2.88 – 2.70 (m, 5H), 2.65 (q, $J$ = 7.6 Hz, 2H), 2.38 – 2.28 (m, 2H), 1.21 (t, $J$ = 7.6 Hz, 3H), 1.03 – 0 .81 (m, 2H), 0.80 – 0.58 (m, 2H).

$^{13}$C-NMR (75 MHz, DMSO, 60 °C): δ [ppm] = 170.9, 170.9, 170.7, 170.0, 144.9, 138.5, 133.5, 128.8, 127.7, 127.2, 126.7, 125.7, 47.9, 41.8, 40.0, 39.4, 27.7, 23.7, 14.8, 12.0.

LC/MS: m/z calculated for $C_{27}H_{34}N_4O_4$ $[M+H]^+$: 479.3, found: 479.2.

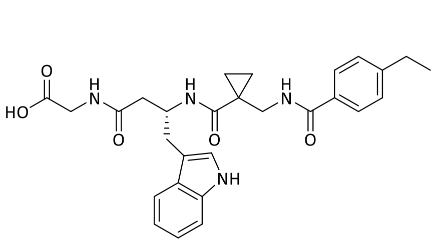

**9**

Compound **9**: Yield: 1.4 mg (0.028 mmol ≙ 43% of th.), colorless solid.

$^{1}$H-NMR (400 MHz, MeO*D*): δ [ppm] = 8.24 (d, *J* = 7.6 Hz, 1H), 7.66 (d, *J* = 7.8 Hz, 2H), 7.60 (d, *J* = 7.7 Hz, 1H), 7.25 (d, *J* = 6.5 Hz, 3H), 7.09 – 6.94 (m, 4H), 6.65 (d, *J* = 7.7 Hz, 1H), 4.56 (d, *J* = 6.4 Hz, 1H), 3.74 (q, *J* = 17.8 Hz, 2H), 3.55 (s, 2H), 3.01 (p, *J* = 7.9 Hz, 2H), 2.69 (q, *J* = 7.5 Hz, 2H), 2.51 (d, *J* = 6.5 Hz, 2H), 2.22 (s, 1H), 1.45 (s, 1H), 1.25 (t, *J* = 7.5 Hz, 3H), 1.10 (q, *J* = 9.8 Hz, 2H), 0.86 (s, 2H).

$^{13}$C-NMR (101 MHz, MeOD): δ [ppm] = 174.9, 173.8, 170.8, 156.0, 149.9, 138.0, 132.5, 130.8, 129.6, 129.2, 129.0, 128.6, 124.3, 122.2, 119.7, 119.6, 116.0, 112.2, 44.6, 40.9, 30.8, 29.7, 28.0, 26.8, 20.5, 15.8, 14.6, 14.4.

LC/MS: m/z calculated for $C_{28}H_{32}N_4O_5$ $[M+H]^+$: 505.2, found: 504.7.

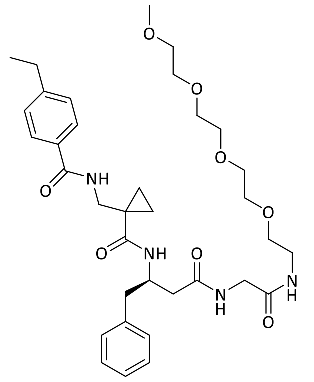

**48**

Compound **48**: Yield: 5.0 mg (7.641 µmol ≙ 44% of th.), colorless solid.

$^{1}$H-NMR (400 MHz, DMSO-*d*$_6$): δ [ppm] = 8.55 (t, *J* = 6.1 Hz, 1H), 8.13 (t, *J* = 5.9 Hz, 1H), 7.99 (d, *J* = 8.1 Hz, 1H), 7.83 (t, *J* = 5.7 Hz, 1H), 7.80 – 7.73 (m, 2H), 7.31 (d, *J* = 8.1 Hz, 2H), 7.21 – 7.02 (m, 5H), 4.29 (q, *J* = 7.1 Hz, 1H), 3.79 – 3.44 (m, 14H), 3.44 – 3.27 (m, 4H), 3.26 – 3.12 (m, 5H), 2.79 (dd, *J* = 13.5, 5.2 Hz, 1H), 2.72 – 2.58 (m, 3H), 2.31 (d, *J* = 6.8 Hz, 2H), 1.20 (t, *J* = 7.6 Hz, 3H), 0.96 – 0.66 (m, 4H).

$^{13}$C-NMR (101 MHz, DMSO): δ [ppm] = 171.5, 170.4, 169.0, 167.2, 147.6, 138.8, 131.4, 129.1, 128.0, 127.7, 127.5, 126.0, 71.3, 69.8, 69.7, 69.6, 69.0, 58.1, 48.4, 42.3, 42.0, 38.6, 30.7, 28.1, 25.2, 15.4, 13.0.

LC/MS: m/z calculated for $C_{35}H_{50}N_4O_8$ $[M+H]^+$: 655.4, found: 655.3.

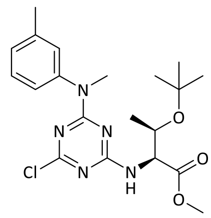

**55**

Compound **55**: Cyanuric chloride (195 mg, 1.057 mmol, 1.0 eq.) and methyl *O*-(*tert*-butyl)-L-threoninate (200 mg, 1.057 mmol, 1.0 eq.) were suspended in 2 mL of DCM. The solution was cooled to 0 °C and DIPEA (370 µL, 2.114 mmol, 2.0 eq.) was added. The reaction was allowed to stir for 10 minutes at 0 °C. Subsequently, *N*,3-dimethylaniline (128 mg, 1.057 mmol, 1.0 eq.), and DIPEA (370 µL, 2.114 mmol, 2.0 eq.) were added at 0 °C and the mixture was stirred for 12 hours at 40 °C in a sealed tube. The crude product was purified by column chromatography (mobile phase: CH:EtOAc 20:1–10:1).

Yield: 331.0 mg (0.786 mmol ≙ 74% of th.), colorless oil.

$^{1}$H-NMR (300 MHz, C*D*Cl$_3$): δ [ppm] = 7.26 (s, 1H), 7.14 – 6.91 (m, 3H), 6.20 – 5.73 (m, 1H), 4.41 – 3.97 (m, 1H), 3.68 (d, *J* = 13.7 Hz, 3H), 3.44 (d, *J* = 7.4 Hz, 3H), 2.36 (d, *J* = 8.2 Hz, 3H), 1.38 – 0.97 (m, 12H).

$^{13}$C-NMR (75 MHz, *C*DCl$_3$): δ [ppm] = 171.6, 165.3, 143.5, 138.8, 129.0, 127.4, 127.0, 124.0, 123.6, 77.4, 74.2, 67.3, 59.6, 52.2, 38.8, 28.4, 21.5.

LC/MS: m/z calculated for $C_{20}H_{28}ClN_5O_3$ [M+H]$^+$: 422.2, found: 422.1.

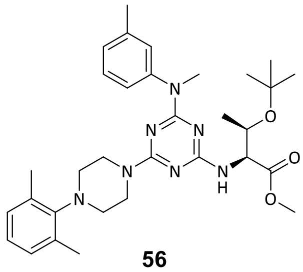

**56**

Compound **56**: Compound **55** (80 mg, 0.190 mmol, 1.0 eq.) was dissolved in DMF, 1-(2,6-dimethylphenyl)piperazine (43 mg, 0.228 mmol, 1.2 eq.) and potassium carbonate (65 mg, 0.475 mmol, 2.5 eq.) were added, and the mixture was allowed to stir for 45 minutes at 100 °C. The solvent was removed in vacuo and the residue was dissolved in DCM, which was washed three times with water. The organic phase was filtered over silica and the product was obtained as an orange oil.

Yield: 103.0 mg (0.179 mmol ≙ 94% of th.), orange oil.

$^1$H-NMR (300 MHz, *C*DCl$_3$): δ [ppm] = 7.28 – 7.09 (m, 3H), 6.99 (h, *J* = 4.6, 4.1 Hz, 4H), 4.27 – 4.11 (m, 1H), 3.81 (d, *J* = 22.4 Hz, 3H), 3.69 (s, 3H), 3.47 (d, *J* = 11.3 Hz, 3H), 3.15 – 3.00 (m, 4H), 2.95 (s, 1H), 2.88 (s, 1H), 2.34 (dd, *J* = 8.9, 2.4 Hz, 9H), 1.24 (dd, *J* = 9.4, 6.3 Hz, 3H), 1.13 (d, *J* = 4.8 Hz, 9H).

$^{13}$C-NMR (75 MHz, *C*DCl$_3$): δ [ppm] = 172.8, 162.6, 161.1, 148.3, 137.9, 136.9, 129.0, 128.0, 127.1, 125.7, 125.2, 123.5, 77.4, 73.9, 67.6, 59.6, 59.4, 51.9, 50.7, 49.8, 47.3, 44.6, 41.6, 36.5, 31.5, 28.4, 21.2, 19.8.

LC/MS: m/z calculated for $C_{32}H_{45}N_7O_3$ [M+H]$^+$: 576.4, found: 576.3.

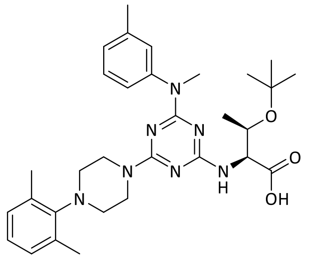

**57**

Compound **57**: Compound **56** (88 mg, 0.152 mmol, 1.0 eq.) was dissolved in 1 mL water and 1 mL THF, and sodium hydroxide (30 mg, 0.750 mmol, 5.0 eq) was added. The mixture was stirred at RT for 48 hours. The solvents were removed under reduced pressure and the crude product was purified by reversed-phase column chromatography (Biotage Isolera, ACN/$H_2O$ + TFA).

Yield: 15.0 mg (0.027 mmol ≙ 18% of th.), colorless solid.

$^1$H-NMR (300 MHz, *C*DCl3): δ [ppm] = 7.39 (t, *J* = 7.5 Hz, 1H), 7.06 – 6.95 (m, 7H), 4.55 (d, *J* = 7.9 Hz, 1H), 4.21 (dd, *J* = 6.4, 2.3 Hz, 1H), 3.92 (s, 3H), 3.47 (s, 3H), 3.12 (t, *J* = 5.4 Hz, 4H), 2.37 (d, *J* = 8.3 Hz, 3H), 2.33 (s, 7H), 1.16 (d, *J* = 20.0 Hz, 12H).

$^{13}$C-NMR (75 MHz, *C*DCl$_3$): δ [ppm] = 184.5, 161.2, 147.6, 137.0, 136.8, 129.3, 129.3, 127.5, 125.9, 123.9, 77.4, 76.9, 75.3, 68.8, 67.0, 49.6, 46.0, 45.9, 39.4, 28.1, 21.4, 20.4, 19.8.

LC/MS: m/z calculated for $C_{31}H_{43}N_7O_3$ [M+H]$^+$: 562.3, found: 562.3.

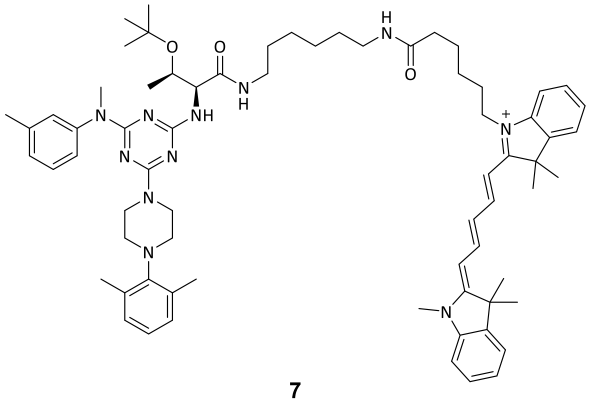

**7**

Compound **7**: Compound **57** (1.0 mg, 1.700 µmol, 1.0 eq.), Cy5 amine (1.0 mg, 1.700 µmol, 1.0 eq.), HATU (1 mg, 2.000 µmol, 1.2 eq.), and DIPEA (3 µL, 16.000 µmol, 9.5 eq.) were dissolved in 20 µL anhydrous DMF. The reaction was allowed to incubate for 16 hours. The crude product was purified by reversed-phase column chromatography (Biotage Isolera, ACN/$H_2O$ + TFA).

Yield: 1.0 mg (0.817 µmol ≙ 48% of th.), blue solid.

LC/MS: m/z calculated for $C_{69}H_{94}N_{11}O_3^+$ $[M+H]^{2+}$: 562.8, found: 562.9.

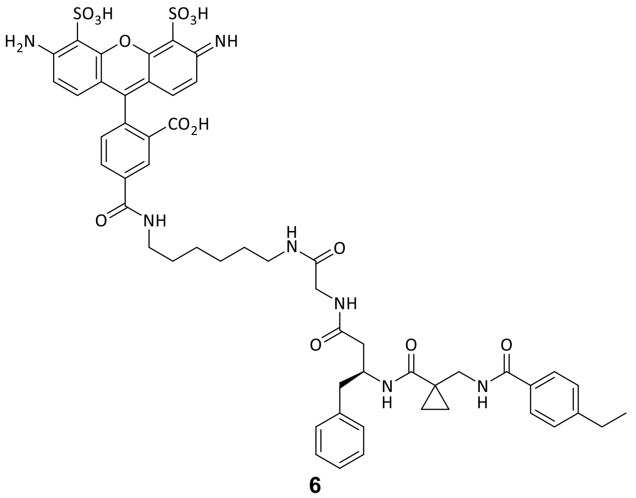

**6**

Compound **6**: Compound **54** (0.8 mg, 1.640 µmol, 1.2 eq.), AF 488 amine (1.0 mg, 1.390 µmol, 1.0 eq.), TBTU (0.5 mg, 1.680 µmol, 1.2 eq.), and DIPEA (2.3 µL, 13.200 µmol, 9.5 eq.) were dissolved in anhydrous DMF (0.6 mL) and stirred at RT for 48 hours. The crude product was purified by reversed-phase column chromatography (Biotage Isolera, ACN/$H_2O$ + TFA).

Yield: 0.4 mg (0.37 mmol ≙ 27% of th.), orange solid.

$^1$H-NMR (600 MHz, DMSO-$d_6$): δ [ppm] = 9.00 (s, 2H), 8.86 (s, 1H), 8.69 (s, 1H), 8.58 (t, *J* = 6.2 Hz, 1H), 8.50 (s, 2H), 8.26 (dd, *J* = 8.0, 1.8 Hz, 1H), 8.16 (t, *J* = 6.0 Hz, 1H), 8.03 (d,

*J* = 8.1 Hz, 1H), 7.81 – 7.73 (m, 2H), 7.50 (d, *J* = 7.8 Hz, 1H), 7.31 (d, *J* = 8.0 Hz, 1H), 7.15 – 7.11 (m, 2H), 7.11 – 7.06 (m, 1H), 7.02 – 6.88 (m, 3H), 4.28 (dt, *J* = 14.2, 6.9 Hz, 1H), 3.69 – 3.60 (m, 1H), 3.58 – 3.50 (m, 1H), 3.50 – 3.45 (m, 2H), 3.33 – 3.22 (m, 1H), 3.07 – 2.99 (m, 2H), 2.78 (dd, *J* = 13.6, 5.2 Hz, 1H), 2.70 – 2.59 (m, 3H), 2.38 (p, *J* = 1.8 Hz, 1H), 2.31 (d, *J* = 6.9 Hz, 2H), 2.11 (s, 1H), 2.08 (s, 1H), 2.04 – 1.91 (m, 2H), 1.58 – 1.48 (m, 2H), 1.38 (q, *J* = 9.4 Hz, 2H), 1.35 – 1.20 (m, 5H), 1.19 (t, *J* = 7.6 Hz, 2H), 1.13 (s, 1H), 0.91 – 0.69 (m, 4H).

LC/MS: m/z calculated for $C_{53}H_{57}N_7O_{14}S_2$ $[M+2H]^{2+}$: 540.7 found: 540.6.

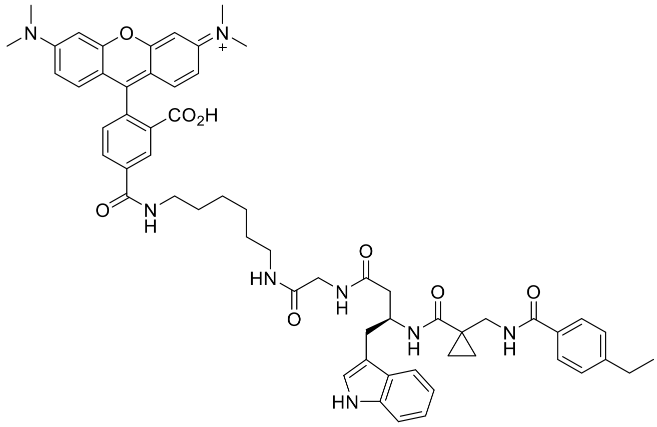

**8**

Compound **8**: Compound **9** (1.1 mg, 2.112 µmol, 1.2 eq.), TBTU (0.7 mg, 2.112 µmol, 1.2 eq.), and TAMRA-amine (1.0 mg, 1.760 µmol, 1.0 eq.) were dissolved in DMF (3 mL), and DIPEA (0.4 µL, 2.112 µmol, 1.2 eq.) was added. The reaction mixture was shaken for 16 hours at RT. Afterward, the solvent was removed under reduced pressure and the crude product was purified by reversed-phase flash column chromatography (Biotage Isolera, ACN/$H_2O$ + TFA).

Yield: 1.9 mg (1.871 µmol ≙ 89% of th.), red solid.

$^1$H-NMR (600 MHz, MeO*D*): δ [ppm] = 8.51 (s, 1H), 8.02 (d, *J* = 7.8 Hz, 1H), 7.88 (d, *J* = 8.2 Hz, 1H), 7.68 (dd, *J* = 8.2, 5.1 Hz, 1H), 7.61 (dd, *J* = 15.0, 8.3 Hz, 2H), 7.50 (d, *J* = 7.9 Hz, 1H), 7.33 – 7.23 (m, 3H), 7.20 – 7.12 (m, 3H), 6.98 – 6.85 (m, 5H), 4.60 (s, 2H), 4.52 (dd, *J* = 14.4, 5.9 Hz, 1H), 3.73 (d, *J* = 16.7 Hz, 1H), 3.66 (s, 1H), 3.63 – 3.52 (m, 3H), 3.45 – 3.35 (m, 3H), 3.32 (t, 1H), 3.21 (d, *J* = 2.8 Hz, 6H), 3.17 – 3.06 (m, 2H), 2.98 – 2.88 (m, 2H), 2.70 – 2.60 (m, 4H), 2.52 (dd, *J* = 14.3, 4.9 Hz, 1H), 2.43 (dd, *J* = 14.4, 8.7 Hz, 1H), 1.62 – 1.54 (m, 2H), 1.49 – 1.41 (m, 2H), 1.39 – 1.28 (m, 4H), 1.27 – 1.16 (m, 6H), 1.06 – 0.97 (m, 2H), 0.93 (t, *J* = 7.4 Hz, 1H), 0.89 – 0.76 (m, 3H).

$^{13}$C-NMR (151 MHz, MeOD): δ [ppm] = 175.0, 174.1, 171.6, 170.7, 169.0, 162.1, 159.0, 158.7, 149.8, 137.9, 132.5, 132.3, 131.0, 129.8, 129.1, 129.0, 128.9, 128.6, 128.5, 128.4, 124.2, 122.2, 119.6, 119.6, 115.1, 114.8, 112.2, 112.1, 97.3, 57.6, 57.5, 57.3, 52.6, 49.9, 49.6, 44.6, 43.7, 41.7,

40.9, 40.7, 40.2, 32.7, 31.1, 30.3, 29.9, 29.8, 27.5, 27.4, 26.9, 21.2, 17.4, 17.3, 17.2, 15.9, 15.9, 14.9, 14.6, 14.6, 14.2.

LC/MS: m/z calculated for $C_{59}H_{67}N_8O_8^+$ $[M+H]^{2+}$: 508.3, found: 508.4.

#### MST supplementary data

MST was employed as the primary screening method for all DEL compounds, measuring the degree of FTAD displacement from DNMT2. Compounds were measured at a final concentration of 100 µM, with sinefungin (SFG) and DMSO included as positive and negative controls. DMSO showed no probe displacement, whereas SFG exhibited complete displacement (SI Figure 2). Compounds demonstrating the most effective FTAD displacement were further investigated via dose-response experiments to determine $K_D$-values. The determination of the additional $K_D$-values for the hit compounds and probe **6** is shown in SI Figure 3.

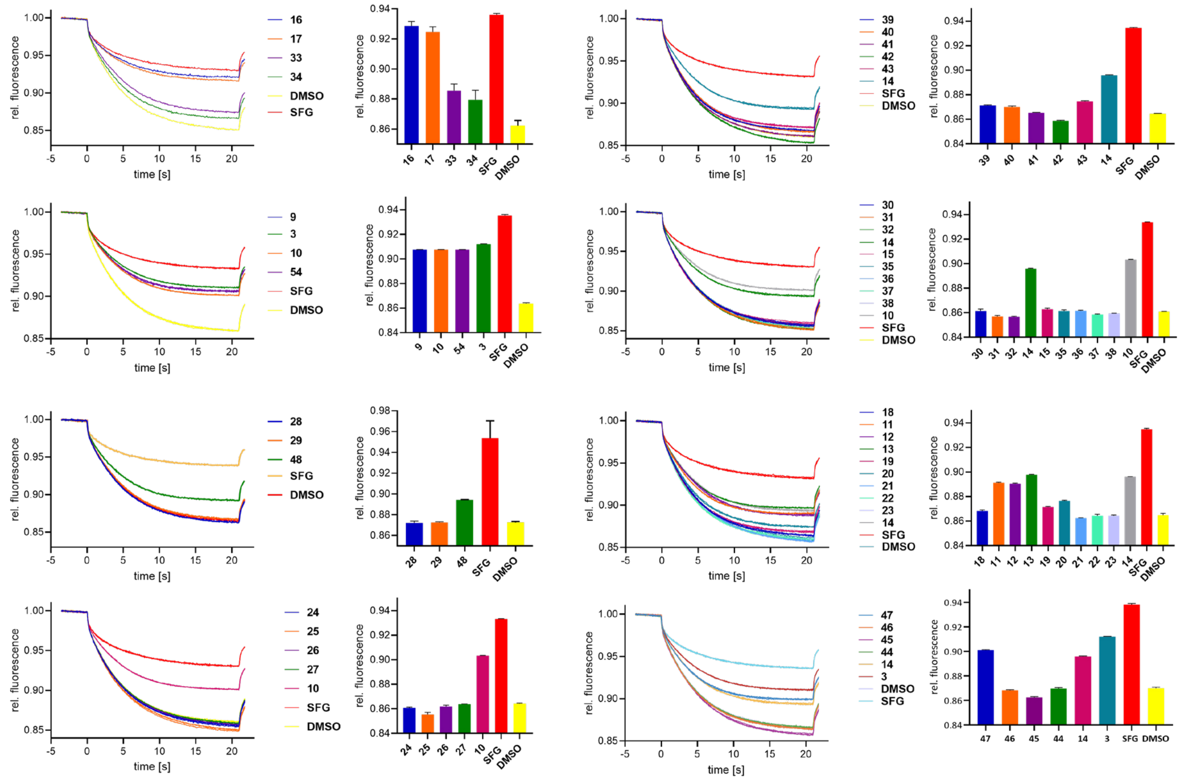

**Figure 2:** MST screening results of all compounds synthesized during the SAR study. Compounds were measured at a concentration of 100 µM. MST experiments were performed as described in the methods section using the fluorescent FTAD probe.

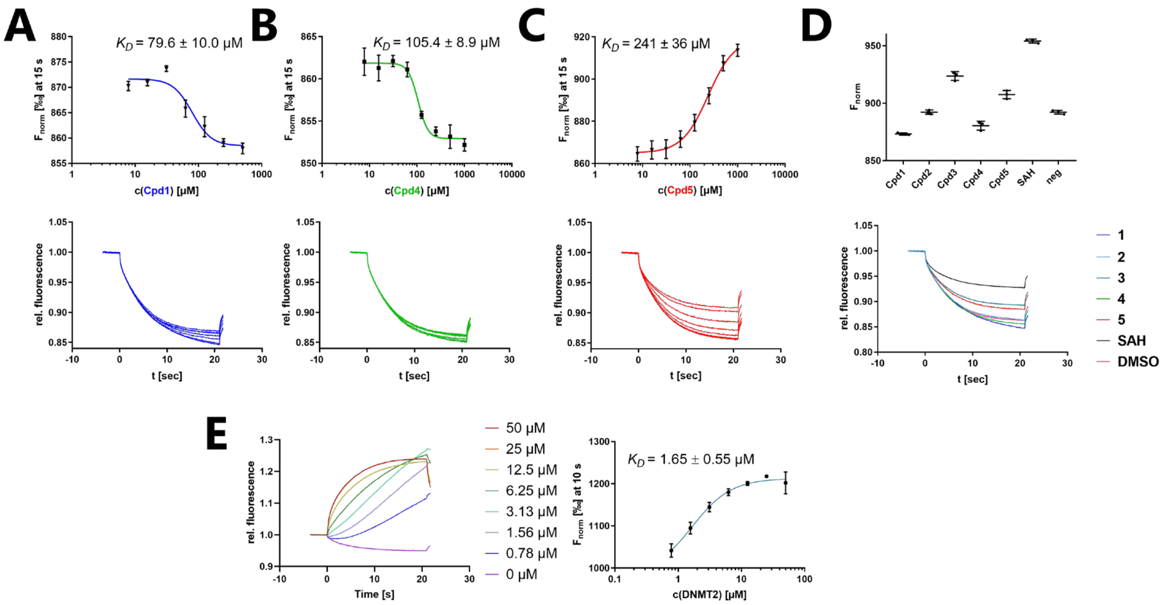

**Figure 3:** Supplementary MST results for the hit compounds selected from the DEL screening. MST experiments were performed as described in the methods section using the fluorescent FTAD probe. (**A**) $K_D$-value determination of compound **1**. (**B**) $K_D$-value determination of compound **4**. (**C**) $K_D$-value determination of compound **5**. (**D**) Screening of all hit compounds **1–5** from the DEL Screening at 100 µM. (**E**) $K_D$-value determination of probe **6** by dose-response experiments using variable concentrations of DNMT2.

#### ITC supplementary data

As an orthogonal method, isothermal titration calorimetry (ITC) was employed to investigate compounds identified as DEL hits and their analogs from the SAR study. Additionally, compound **3** was investigated by ITC displacement experiments using SAH as the competitive titrant. Thermograms are complemented by a signature blot illustrating changes in enthalpy (ΔH), free energy (ΔG), and entropy (-TΔS) together with its stoichiometry plot. Figures can be found in SI Figures 4 and 5. Experiments were conducted as described in the method section.

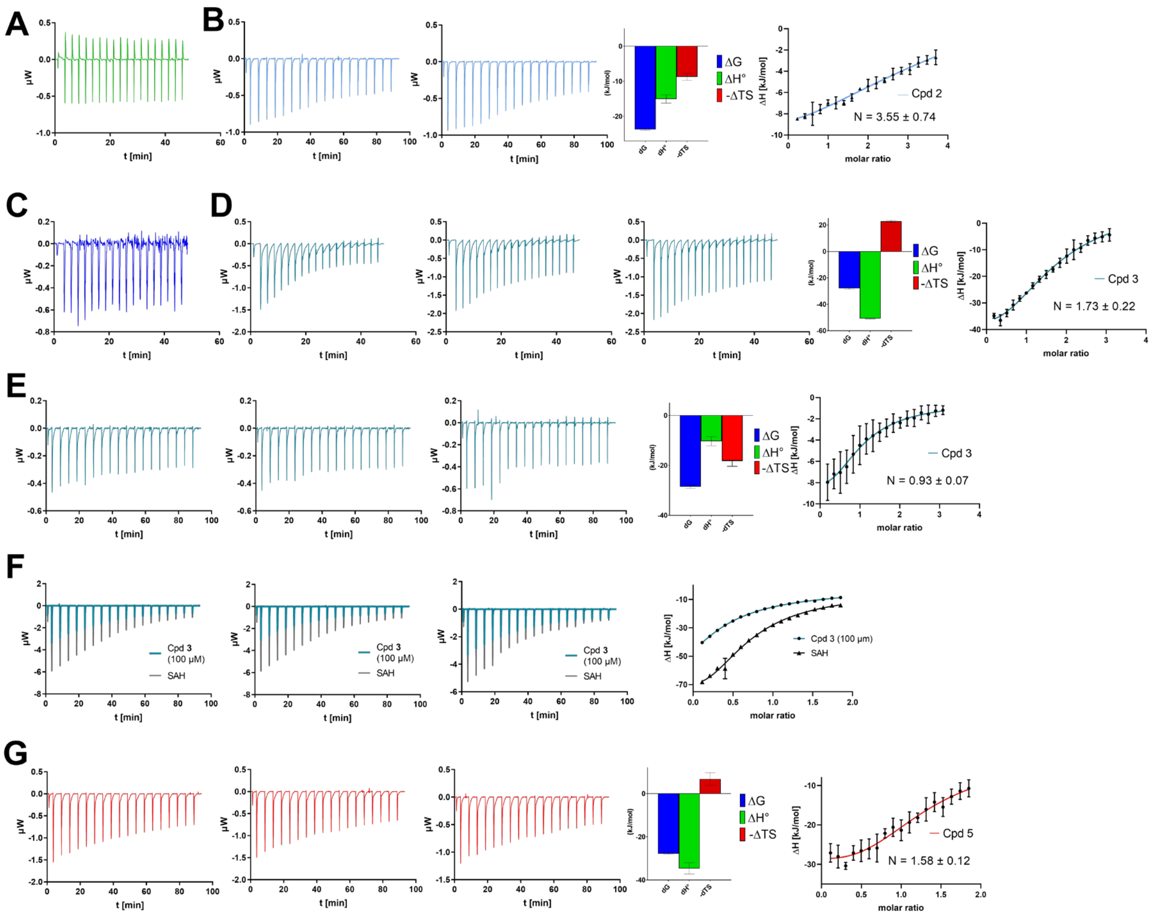

**Figure 4:** Supplementary ITC data for compounds **1–5.** (**A**) Compound **4** was titrated into DNMT2. No enthalpy signal change was observed. (**B**) Compound **2** was titrated into DNMT2. (**C**) Compound **1** was titrated into DNMT2. No enthalpy signal change was observed. (**D**) Compound **3** was titrated into DNMT2. (**E**) Compound **3** titrated into DNMT2 with a spacing time of 300 s. (**F**) DNMT2-SAH displacement experiment for compound **3**. (**G**) Compound **5** was titrated into DNMT2.

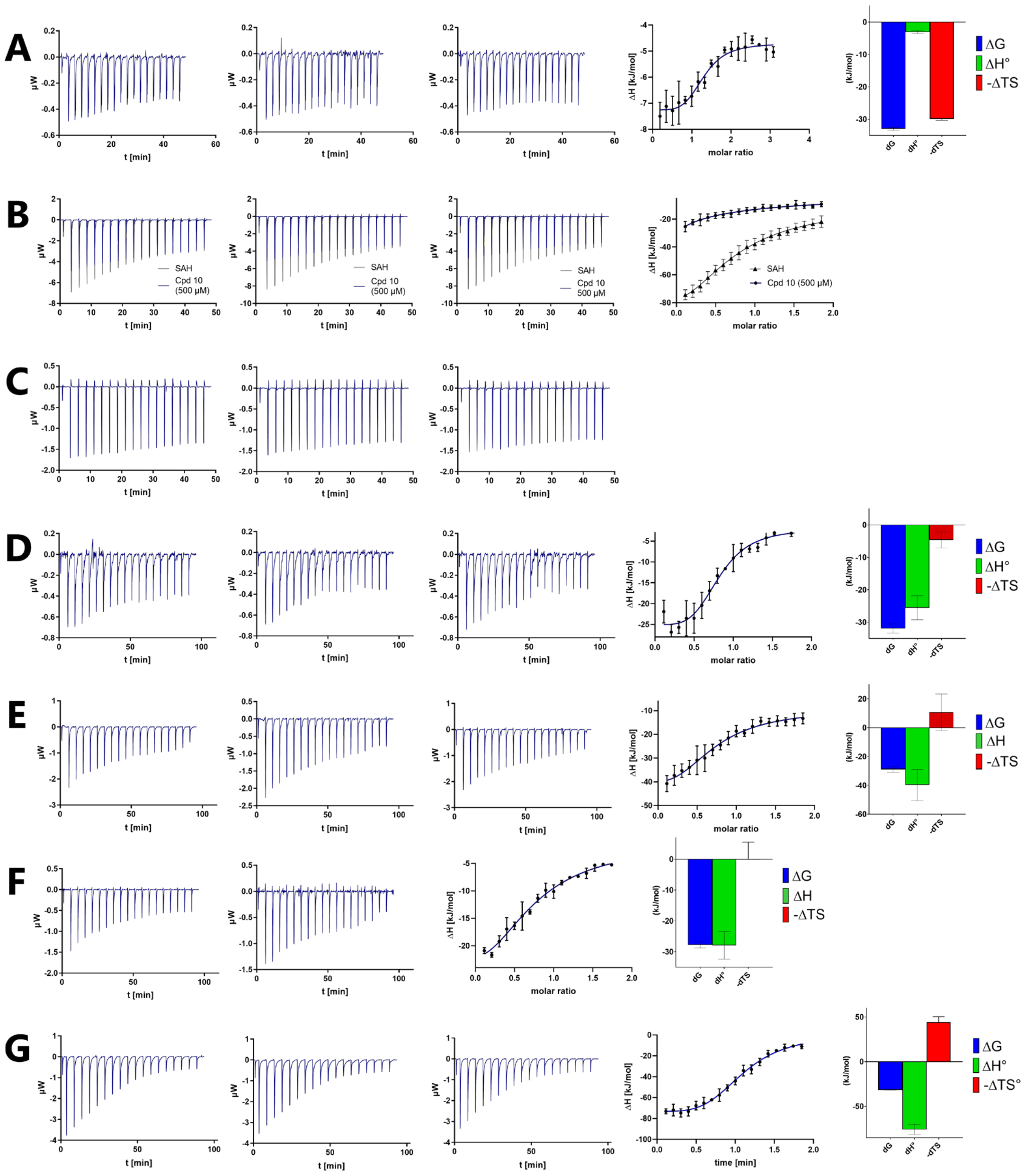

**Figure 5:** Supplementary ITC data for synthesized compound **3** analogs. For compound **30** no binding was observed, so no stoichiometry plot and signature plot was created. (**A**) Compound **10** (**B**) Compound **10** measured by SAH displacement (**C**) **30** (**D**) **14** (**E**) **47** (**F**) **11** (**G**) **16**.

#### Affinity selection-mass spectrometry (AS-MS)

Affinity selection-mass spectrometry assays were utilized to provide an orthogonal characterization of compound **3** binding to DNMT2. The experimental procedure was conducted as described in the method section.

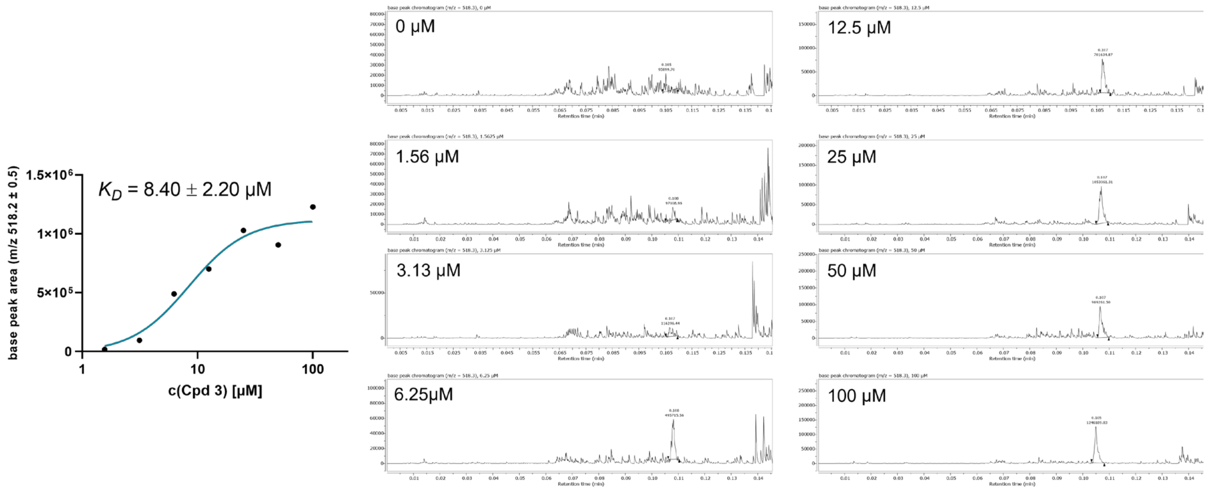

**Figure 6:** AS-MS-based determination of compound **3** binding affinity to DNMT2. (**left**) Dose-response curve for $K_D$ determination. (**right**) Base peak chromatograms for compound **3** content analysis in the analyzed fractions.

#### Supplementary data of $^3$H-based methyltransferase DNMT2 activity assays

To determine the DNMT2 inhibitory properties of the DEL hit compounds a $^3$H-based incorporation assay screening was performed as described in the method section. Since compounds **3**–**5** showed significant inhibition, $IC_{50}$-values were determined (see Figure 2 in main manuscript)

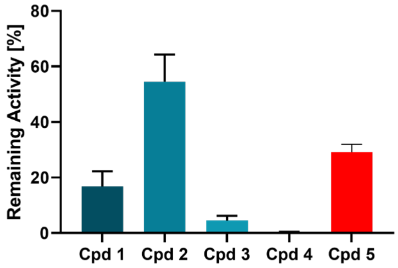

**Figure 7:** Screening data from $^3$H-based methyltransferase enzyme activity assay. For compounds **1**–**5**, the remaining DNMT2 activity was determined at 500 µM of the respective compound.

##### DNMT2-tRNA MST displacement experiments

Here, we used a recently described MST protocol for the investigation of hit compounds on the influence of DNMT2-tRNA complex formation. MST displacement experiments with in situ fluorescently labelled (1x SybrGold) $tRNA^{Asp}$ showed consistently that none of the hit compounds **1**–**5** was able to dissociate the tRNA-protein complex formed by DNMT2 and $tRNA^{Asp}$ ($K_D$=210 nM, SI Figure 8 A, B) and no hit compound (100 μM) had a significant influence on the thermophoresis of $tRNA^{Asp}$ in protein-free solution (SI Figure 8 C, D).

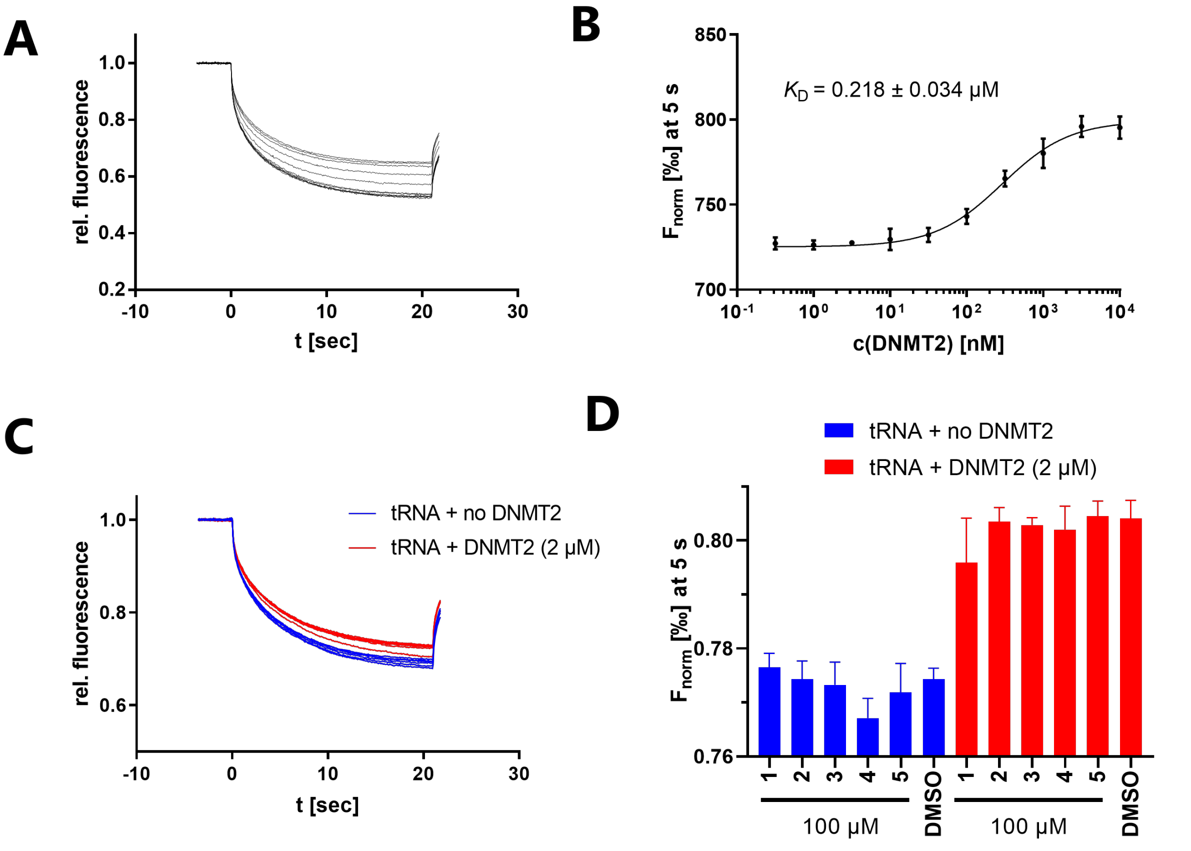

**Figure 8:** Analysis of DNMT2-RNA interaction by in situ labeling with SybrGold (1×). (**A**, **B**) Dissociation constant ($K_D$) of $tRNA^{Asp}$ (100 nM, labeled with SybrGold) towards DNMT2 (varying concentrations) was determined by dose-response experiments. (**C**, **D**). Neither the native $tRNA^{Asp}$ samples (blue) nor the DNMT2-tRNA complex (red) showed significant thermophoresis alterations when treated with the hit compounds **1**–**5**.

#### Cell viability

Figure 9 presents cell viability data for treatment with compounds **1**–**5** and **16** across various cell lines, including wild-type HCT cells, HCT cells with NSUN2 knock-down, HCT cells with DNMT2 knock-down, and wild-type MOLM-13 cells. Notably, peptidomimetic compounds **3**–**5** and **16** exhibit low toxicity at all concentrations, while compounds **1** and **2** demonstrate significant toxicity only at higher concentrations (100 μM).

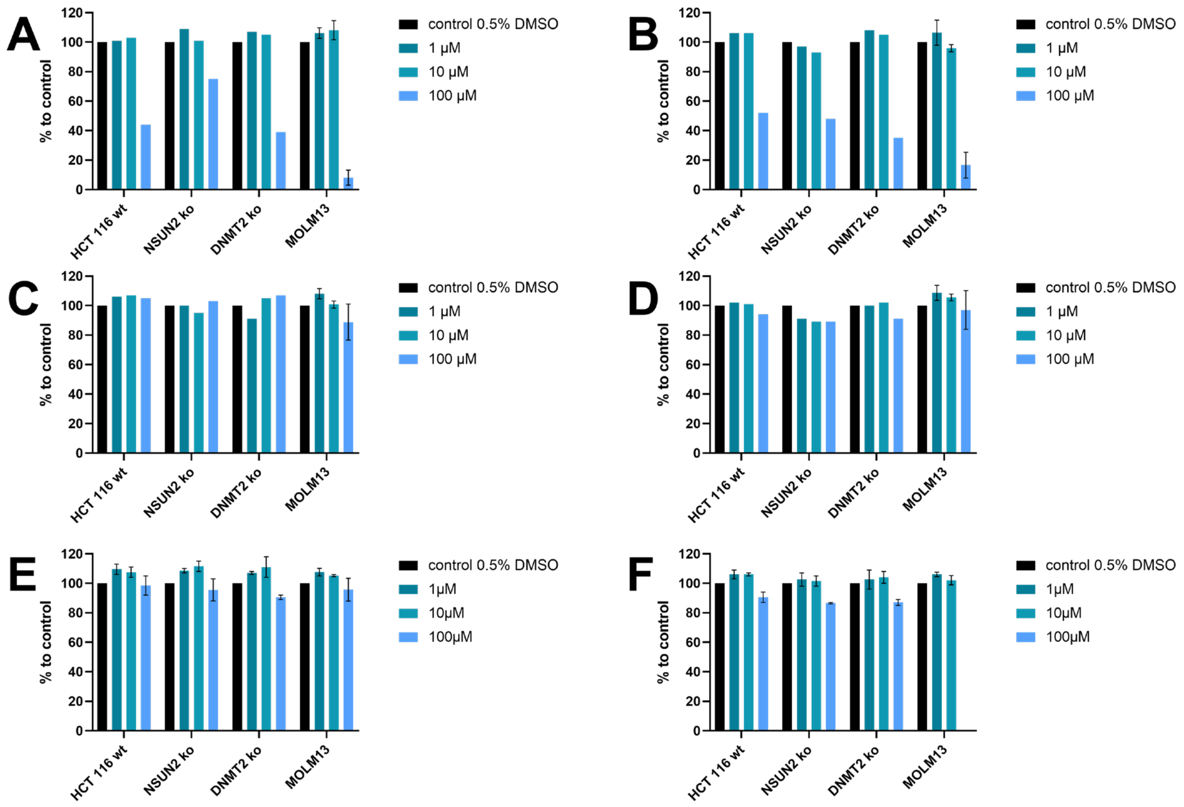

**Figure 9:** Cell viability assay data for compounds **1–5** and **16** used for treatment of HCT wild-type cells, HCT NSUN2 knock-down cells, HCT DNMT2 knock-down cells, and MOLM13 wild-type cells. (**A**) Data for compound **1**. (**B**) Data for compound **2**. (**C**) Data for compound **3**. (**D**) Data for compound **4**. (**E**) Data for compound **5**. (**F**) Data for compound **16**.

#### Parallel Artificial Membrane Permeation Assay (PAMPA)

Table 2 presents the PAMPA assay results, conducted as described in the method section, revealing that only **16** could diffuse across the artificial membrane. SI Figures 10–15 highlight the corresponding chromatograms. Compounds **1** and **2** were not sufficiently soluble for evaluation by PAMPA.

**Table 2:** PAMPA results of the investigated compounds determined in duplicates.

| Compound | Compound structure | $P_{app}$ [*$10^{-6}$ cm/s] |
|---|---|---|
| **3** | 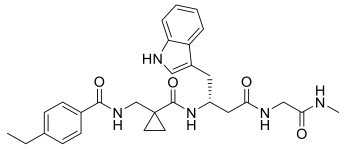 | below detection limit |
| **10** | 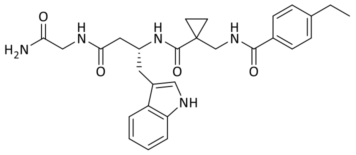 | below detection limit |
| **14** | 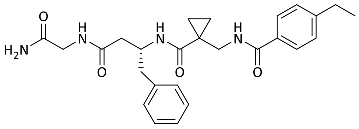 | below detection limit |
| **16** | 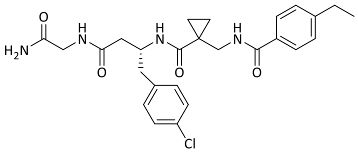 | 0.30 |
| **47** | 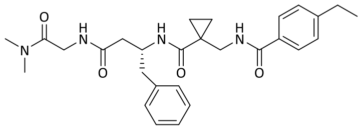 | below detection limit |
| **54** | 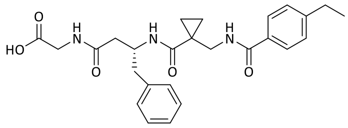 | below detection limit |

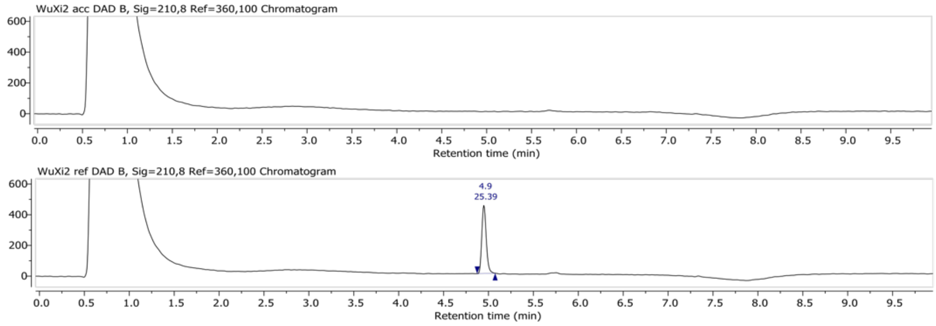

**Figure 10:** Base peak chromatogram at m/z = 518.3 ± 0.5 of acceptor and reference solution after incubating for compound **3**. The peak of compound **3** is marked and the area under the peak is stated above the peak.

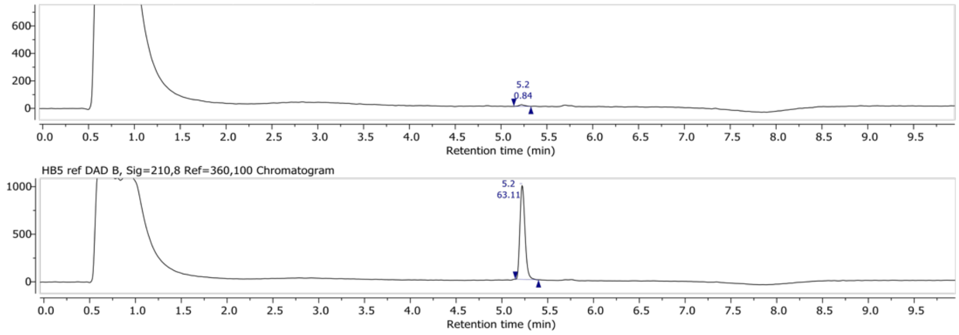

**Figure 11:** Base peak chromatogram at m/z = 499.2 ± 0.5 of acceptor and reference solution after incubating for compound **16**. The peak of compound **16** is marked and the area under the peak is stated above the peak.

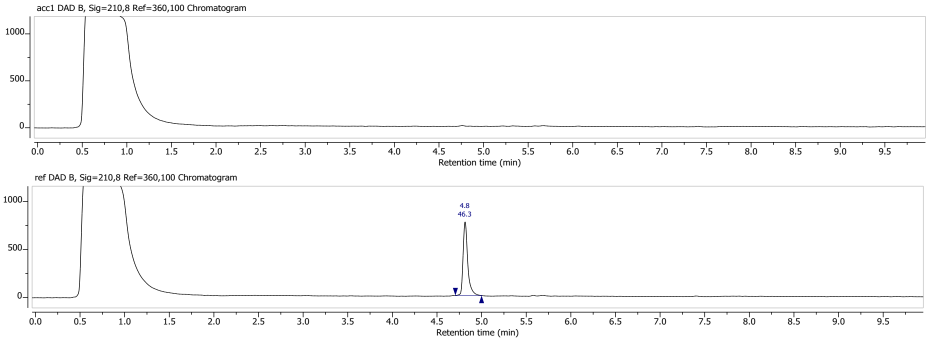

**Figure 12:** Base peak chromatogram at m/z = 504.3 ± 0.5 of acceptor and reference solution after incubating for compound **10**. The peak of compound **10** is marked and the area under the peak is stated above the peak.

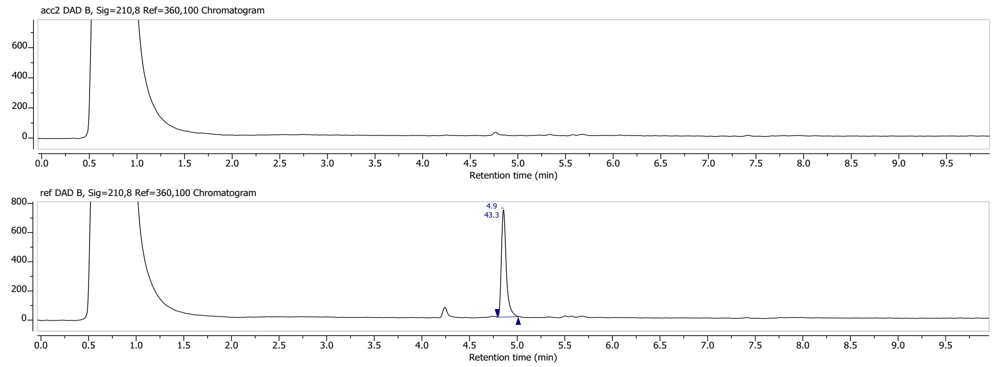

**Figure 13:** Base peak chromatogram at m/z = 465.3 ± 0.5 of acceptor and reference solution after incubating for compound **14**. The peak of compound **14** is marked and the area under the peak is stated above the peak.

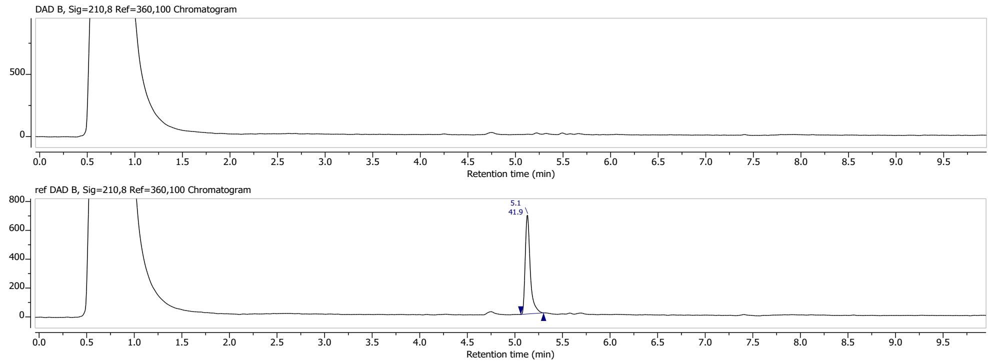

**Figure 14:** Base peak chromatogram at m/z = 493.3 ± 0.5 of acceptor and reference solution after incubating for compound **47**. The peak of compound **47** is marked and the area under the peak is stated above the peak.

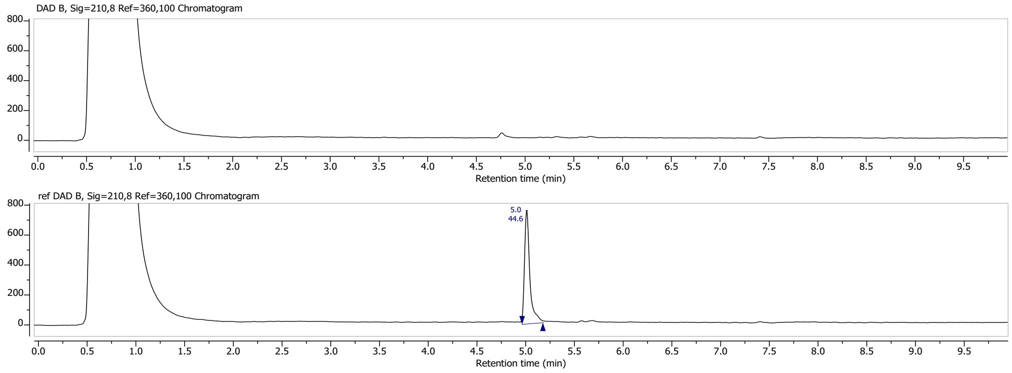

**Figure 15:** Base peak chromatogram at m/z = 466.2 ± 0.5 of acceptor and reference solution after incubating for compound **54**. The peak of compound **54** is marked and the area under the peak is stated above the peak.

#### Pull-down interaction assay

In SI Figure 16, supplementary data for the pull-down interaction assay between DNMT2 and NonO is presented. The methodology employed for these assays is detailed in the method section and Figure 10, where the results are also discussed.

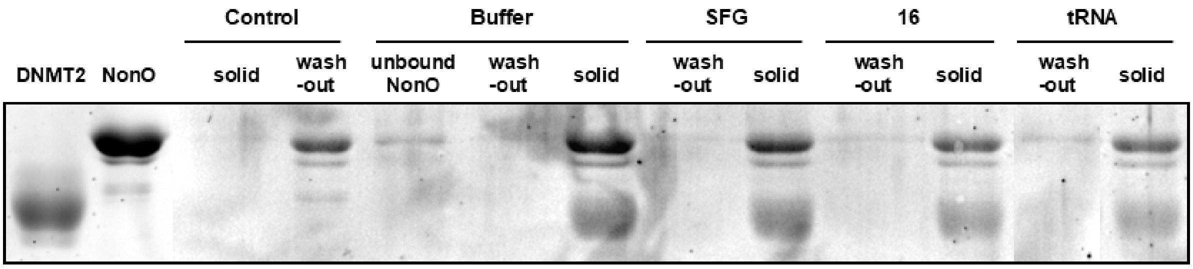

**Figure 16:** Known DNMT2-interacting proteins do not bind to the allosteric pocket. Results of the SDS-PAGE-based analysis. Lane description from left to right: DNMT2/NonO: reference proteins for size comparison; Control: beads without immobilized DNMT2 served as the negative control, i.e. on these beads no binding of NonO was observed. Treatments (buffer, SFG, **16**, tRNA): The lane "unbound NonO" contains the supernatant after immobilization of NonO, which showed low protein levels, thus most NonO is effectively immobilized on DNMT2 beads. Lanes labeled with (wash-out) contain the supernatant of the respective treatment conditions (buffer, 100 µM SFG, 100 µM **16**, 10 µM $tRNA^{Asp}$), while bands labeled with (solid) contain the solid fraction obtained by boiling the residual beads in Laemmli buffer. NonO could not be displaced from binding DNMT2 by competitive treatment with compound **16**, SFG, nor $tRNA^{Asp}$, indicating that NonO does not bind to the allosteric binding pocket identified.

**Supplementary data for crystallographic investigations**

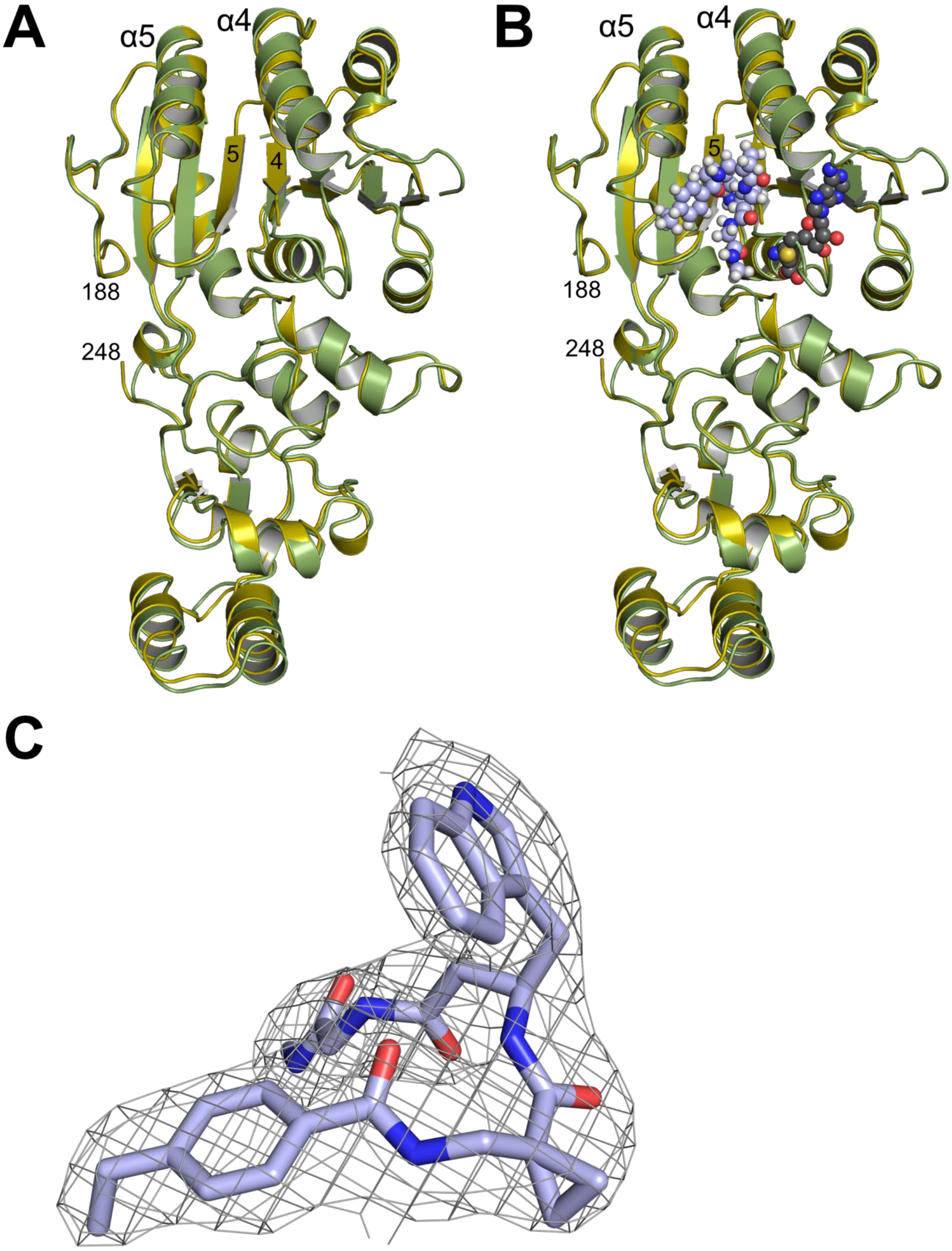

**Figure 17:** Global superposition of the X-ray structures of human DNMT2Δ47-compound **3** (green / light blue) and human DNMT2Δ47-SAH (yellow / grey) without (**A**) and with (**B**) ligands. (**C**) Compound **3** bound to DNMT2Δ47 (chain A) with 2m$F_{obs}$-D$F_{calc}$ electron density map at a contour level of +1σ.

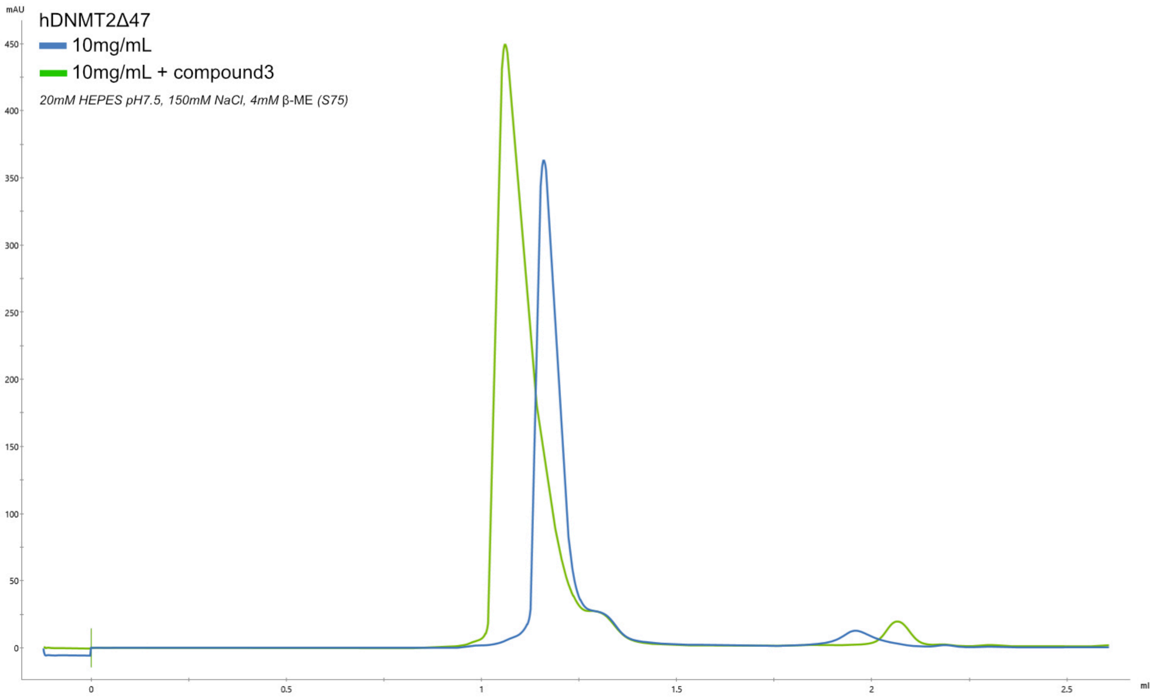

**Figure 18:** Analytical SEC elution profile of DNMT2Δ47 with and without compound **3**. Superposition of two chromatograms showing the 280 nm signal of analytical SEC runs for hDNMT2Δ47 without ligand (blue) and pre-incubated with a 2-fold molar excess of compound **3** (green). For both runs the protein concentration was at 10 mg/mL as used for crystallization. A Superdex S75 3.2/300 column was used with a buffer consisting of 20 mM HEPES pH 7.5, 150 mM NaCl and 4 mM β-ME. The peak shift indicates the dimerization of DNMT2Δ47 upon compound **3** binding.

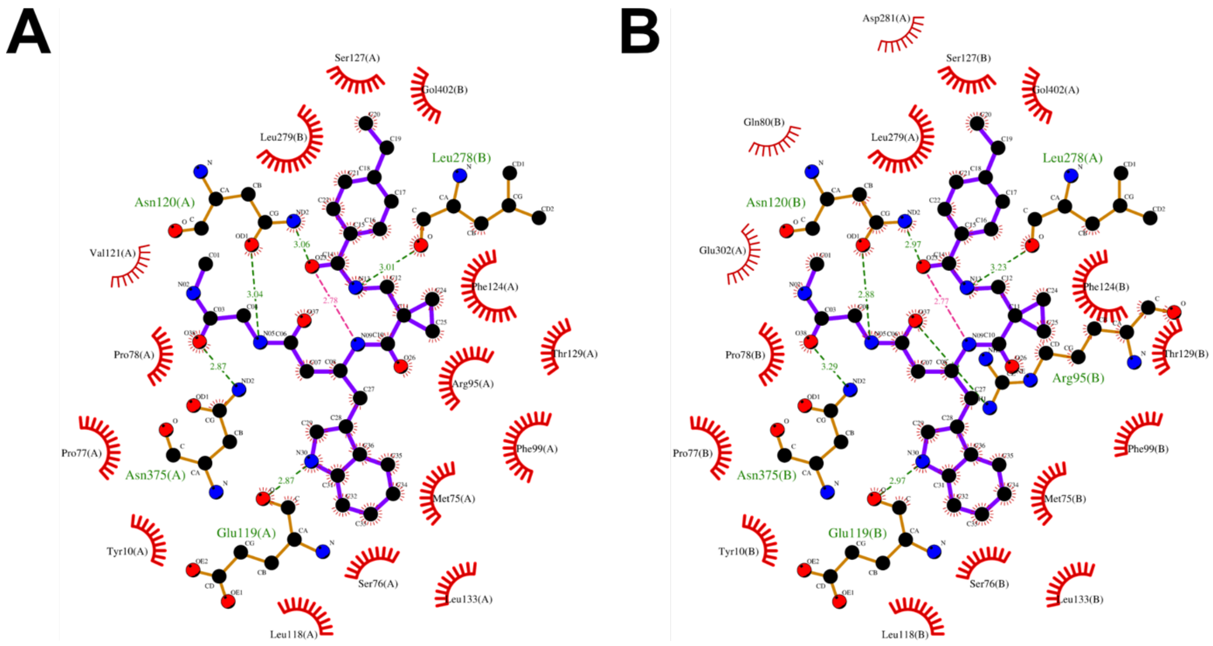

**Figure 19:** A schematic ligand-protein interaction diagram created with LigPlot+ for compound **3** bound to chain A (**A**) and chain B (**B**). Intermolecular hydrogen bonds are given in green, and the intramolecular hydrogen bond in compound **3** was added to the diagram in magenta.

**Table 3:** X-ray data collection refinement statistics.

| **PDB entry 9HGM** | |
|---|---|
| **Data Collection** | |
| Beamline | ESRF Grenoble ID30B |
| Wavelength (Å) | 0.8731 |
| Resolution range (Å) | 48.67-2.60 (2.72-2.60) |
| Space group | $P2_12_12_1$ |
| a,b,c (Å) | 71.20, 97.35, 125.02 |
| α,β,ɣ (°) | 90, 90, 90 |
| Total reflections | 263886 (33598) |
| Unique reflections | 27445 (3300) |

| | |
|---|---|
| Multiplicity | 9.6 (10.2) |
| Completeness (%) | 100 (100) |
| Mean I/sigma(I) | 10.6 (1.2) |
| R-merge | 0.117 (1.966) |
| R-pim | 0.042 (0.688) |
| CC1/2 | 0.999 (0.626) |
| **Refinement** | |
| Resolution range (Å) | 48.67-2.60 (2.69-2.60) |
| Reflections used in refinement | 27366 (2667) |
| Reflections used for R-free | 1396 (132) |
| R-work | 0.1995 (0.3377) |
| R-free | 0.2454 (0.3798) |
| Number of non-hydrogen atoms | 5410 |
| Macromolecules | 5276 |
| Ligands | 88 |
| Solvent | 46 |
| Protein residues | 656 |
| RMS (bonds) (Å) | 0.005 |
| RMS (angles) (°) | 0.580 |
| Ramachandran favored (%) | 97.4 |
| Ramachandran allowed (%) | 2.6 |
| Ramachandran outliers (%) | 0 |
| Rotamer outliers (%) | 0.84 |
| Clashscore | 2.02 |
| Average B-factor (Å$^2$) | 83.3 |
| Macromolecules (Å$^2$) | 83.6 |
| Ligands (Å$^2$) | 68.1 |
| Solvent (Å$^2$) | 74.1 |

Statistics for the highest-resolution shell are shown in parentheses.

### Spectral Appendix

#### Compound 1:

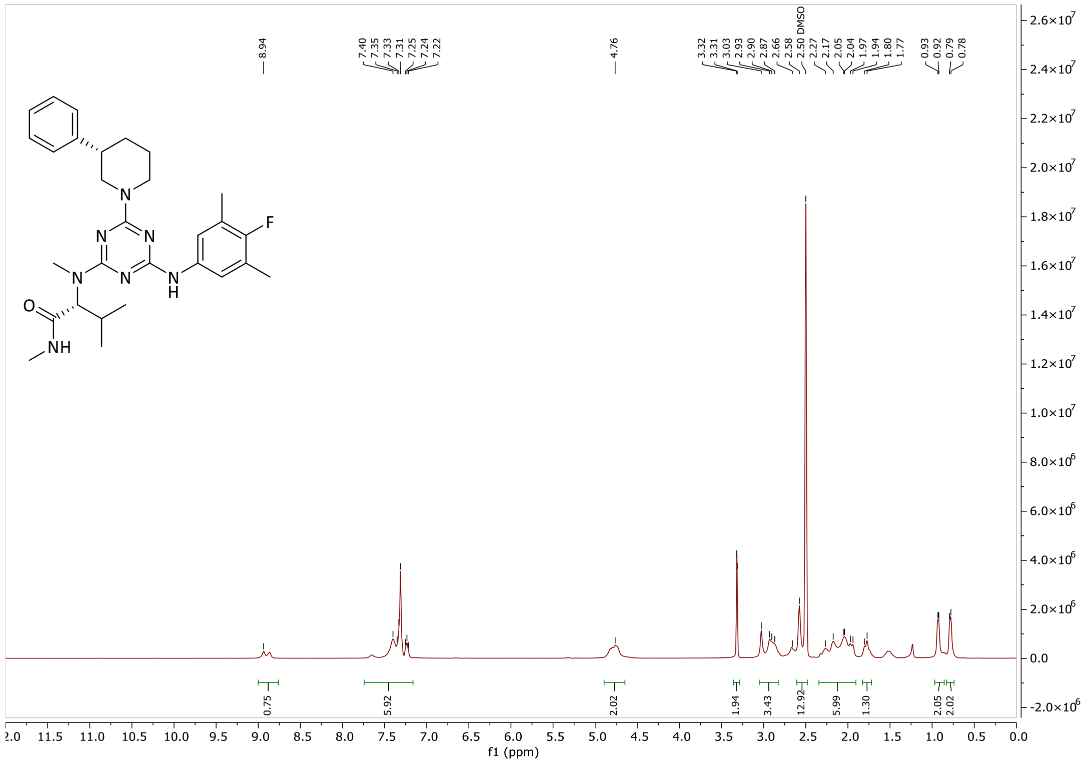

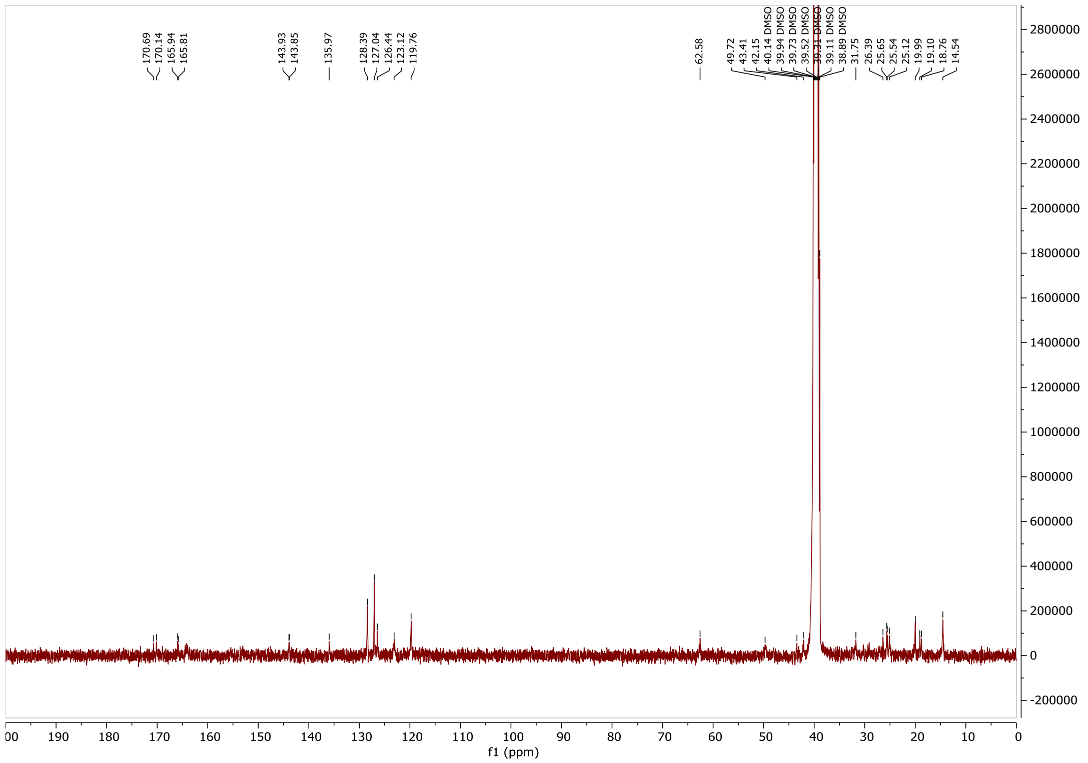

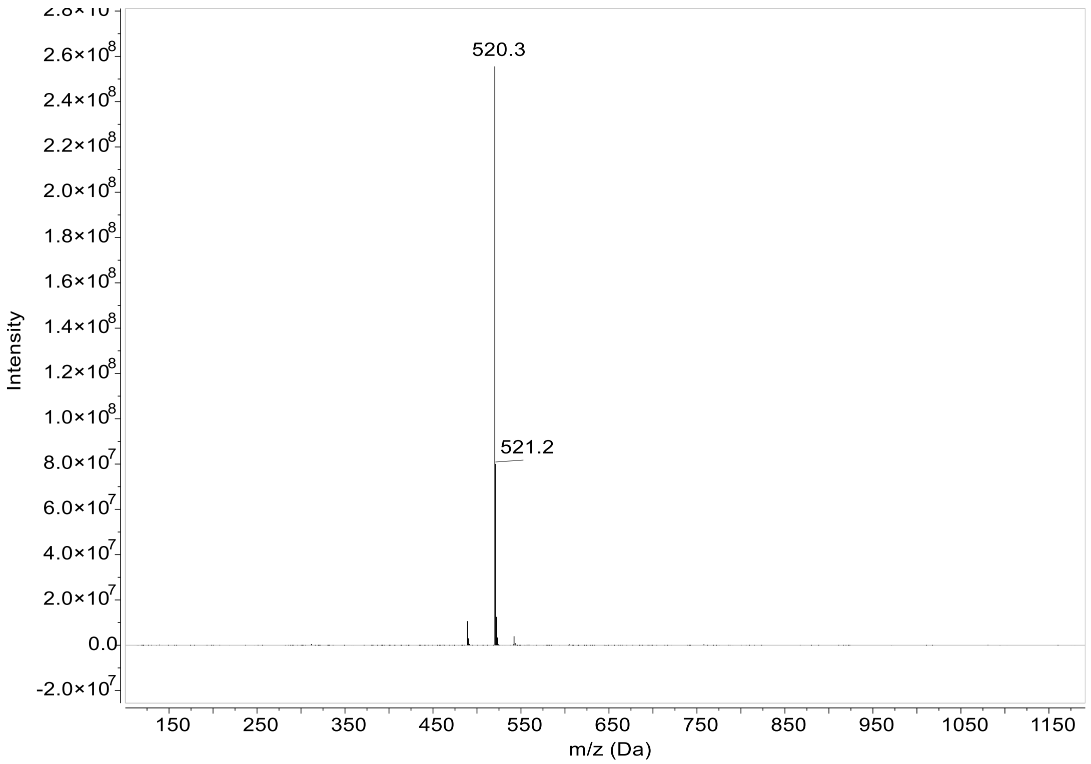
520.3
521.2
Intensity
m/z (Da)

#### Compound 2:

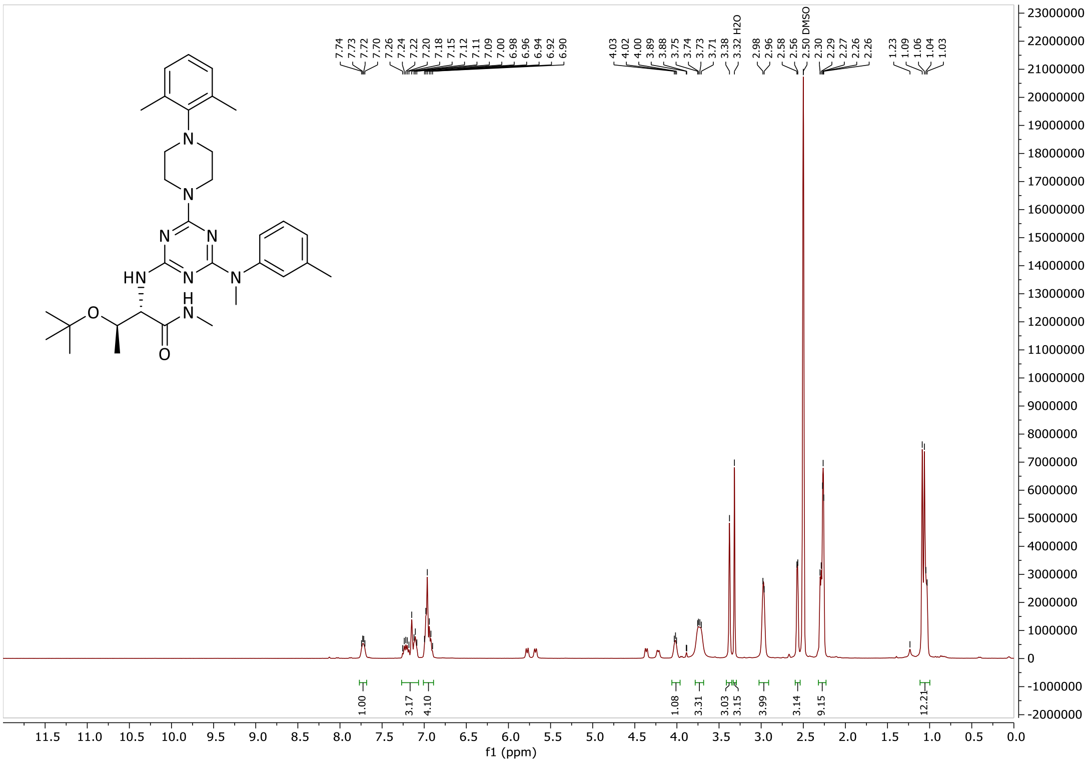
7.74
7.73
7.72
7.70
7.26
7.24
7.22
7.20
7.18
7.15
7.12
7.11
7.09
7.00
6.98
6.96
6.94
6.92
6.90
4.03
4.02
4.00
3.89
3.88
3.75
3.74
3.73
3.71
3.38
3.32 H2O
2.98
2.96
2.58
2.56
2.50 DMSO
2.30
2.29
2.27
2.26
2.26
1.23
1.09
1.06
1.04
1.03
1.00
3.17
4.10
1.08
3.31
3.03
3.15
3.99
3.14
9.15
12.21
f1 (ppm)

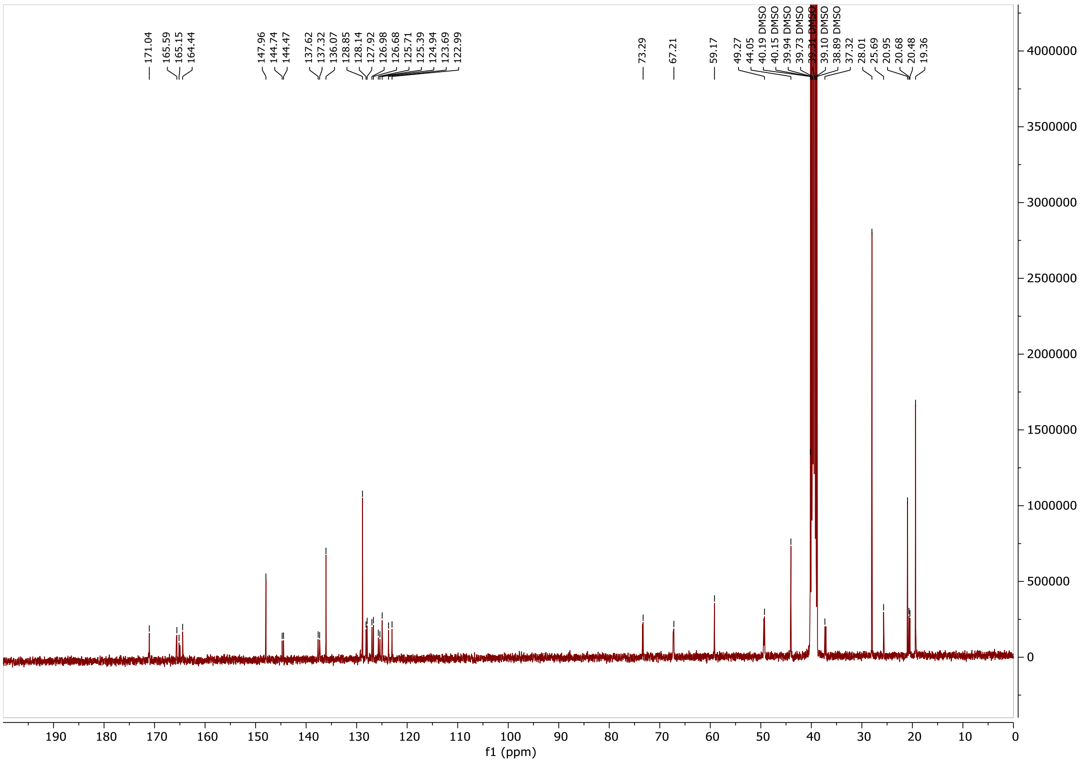
171.04
165.59
165.15
164.44
147.96
144.74
144.47
137.62
137.32
136.07
128.85
128.14
127.92
126.98
126.68
125.71
125.39
124.94
123.69
122.99
73.29
67.21
59.17
49.27
44.05
40.19 DMSO
40.15 DMSO
39.94 DMSO
39.73 DMSO
39.10 DMSO
38.89 DMSO
37.32
28.01
25.69
20.95
20.68
20.48
19.36
f1 (ppm)

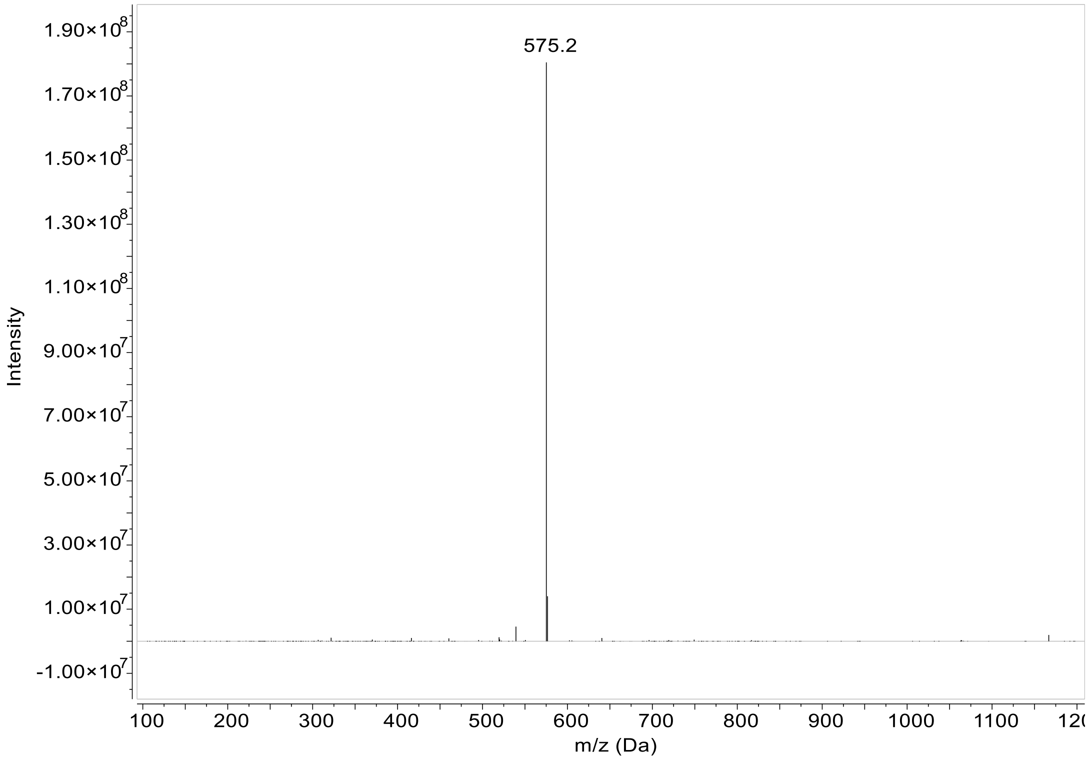
575.2
Intensity
m/z (Da)
1.90×10^8
1.70×10^8
1.50×10^8
1.30×10^8
1.10×10^8
9.00×10^7
7.00×10^7
5.00×10^7
3.00×10^7
1.00×10^7
-1.00×10^7
100
200
300
400
500
600
700
800
900
1000
1100

### Compound 3:

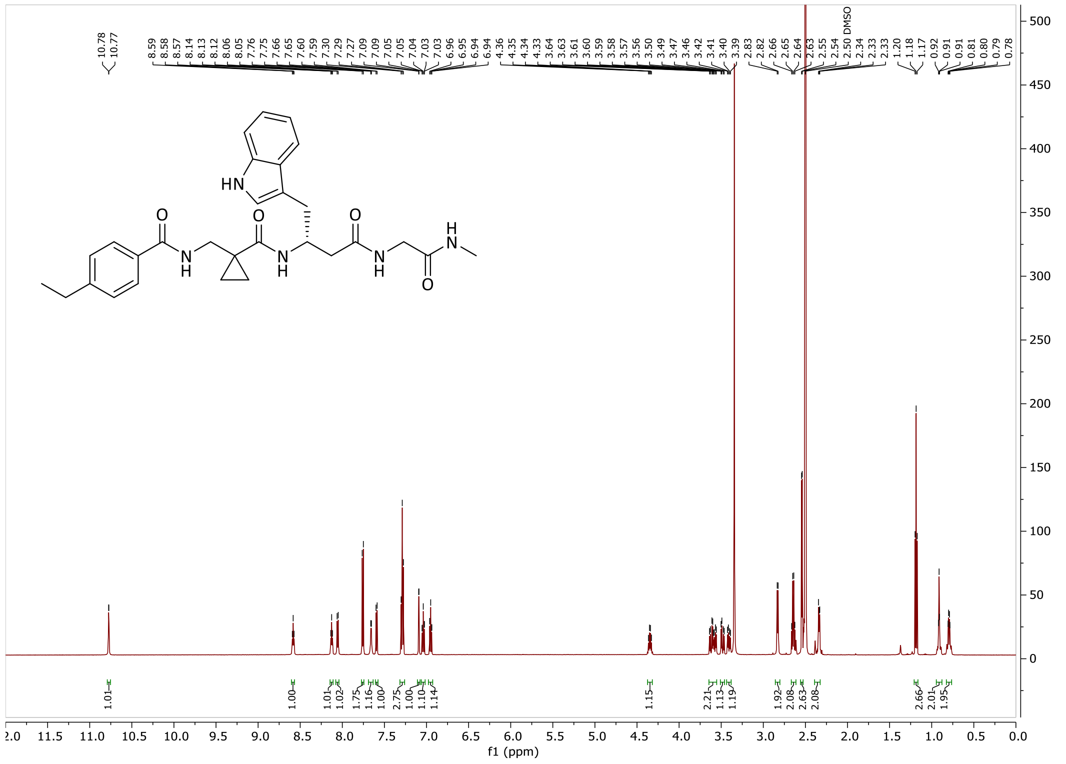

10.78
10.77
8.59
8.58
8.57
8.14
8.13
8.12
8.06
8.05
7.76
7.75
7.66
7.65
7.60
7.59
7.30
7.29
7.27
7.09
7.09
7.05
7.05
7.04
7.03
7.03
6.96
6.95
6.94
6.94
4.36
4.35
4.34
4.33
3.64
3.63
3.61
3.60
3.59
3.58
3.57
3.56
3.50
3.49
3.47
3.46
3.42
3.41
3.40
3.39
2.83
2.82
2.66
2.65
2.64
2.63
2.55
2.54
2.50 DMSO
2.34
2.33
2.33
1.20
1.18
1.17
0.92
0.91
0.91
0.81
0.80
0.79
0.78
HN
O
O
O
N
H
N
H
N
H
H
N
O
1.01
1.00
1.01
1.02
1.75
1.16
1.00
2.75
1.00
1.10
1.14
1.15
2.21
1.13
1.19
1.92
2.08
2.63
2.08
2.66
2.01
1.95
f1 (ppm)

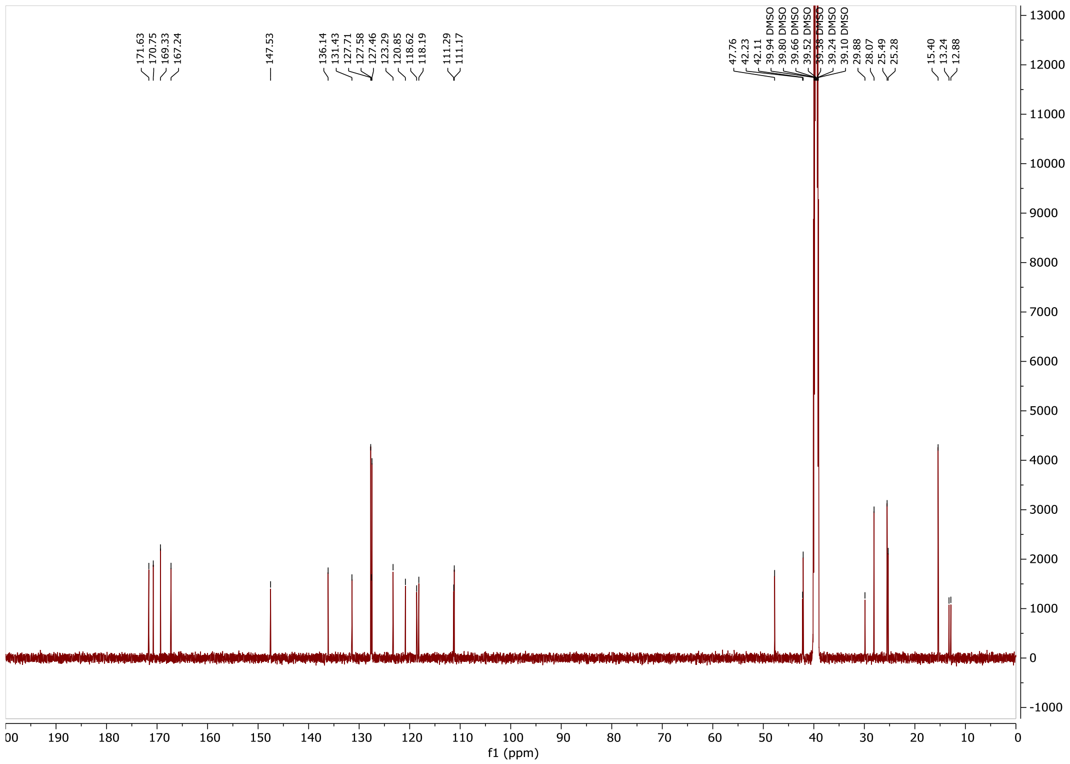

171.63
170.75
169.33
167.24
147.53
136.14
131.43
127.71
127.58
127.46
123.29
120.85
118.62
118.19
111.29
111.17
47.76
42.23
42.11
39.94 DMSO
39.80 DMSO
39.66 DMSO
39.52 DMSO
39.38 DMSO
39.24 DMSO
39.10 DMSO
29.88
28.07
25.49
25.28
15.40
13.24
12.88
f1 (ppm)

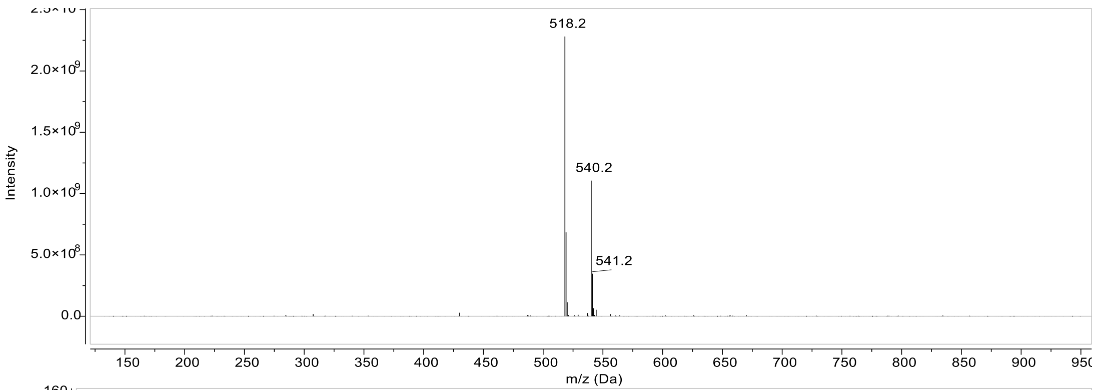

518.2
540.2
541.2
Intensity
2.0×10^9
1.5×10^9
1.0×10^9
5.0×10^8
0.0
m/z (Da)

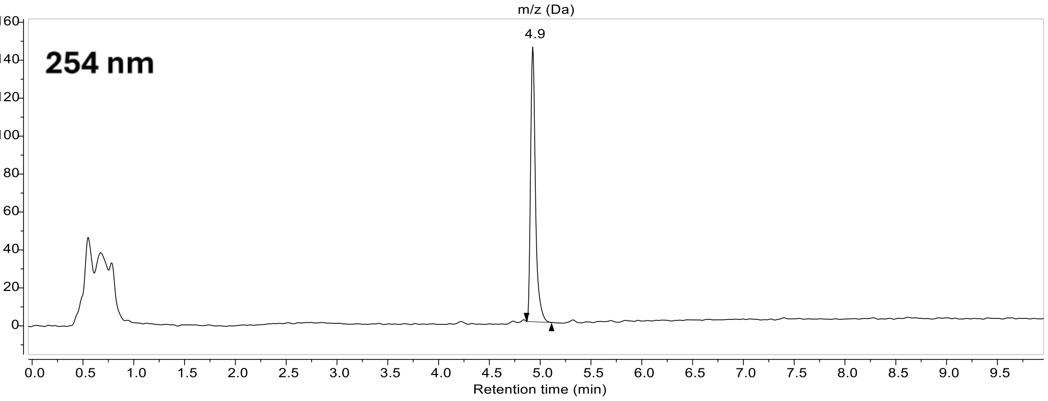

254 nm
4.9
Retention time (min)

### Compound 4:

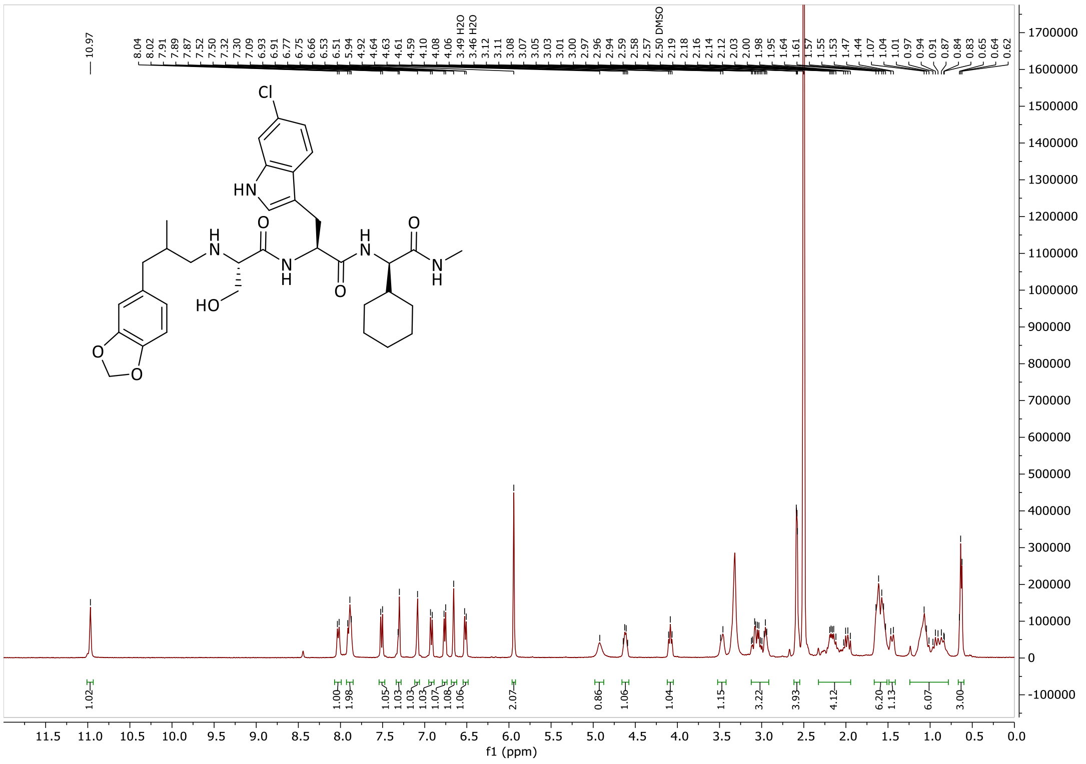

10.97
8.04 8.02 7.91 7.89 7.87 7.52 7.50 7.32 7.30 7.09 6.93 6.91 6.77 6.75 6.66 6.53 6.51 5.94 4.92 4.64 4.63 4.61 4.59 4.10 4.08 4.06 3.49 H2O 3.46 H2O 3.12 3.11 3.08 3.07 3.05 3.03 3.01 3.00 2.97 2.96 2.94 2.59 2.58 2.57 2.50 DMSO 2.19 2.18 2.16 2.14 2.12 2.03 2.00 1.98 1.95 1.64 1.61 1.57 1.55 1.53 1.47 1.44 1.07 1.04 1.01 0.97 0.94 0.91 0.87 0.84 0.83 0.65 0.64 0.62
1.02 1.00 1.98 1.05 1.03 1.03 1.03 1.07 1.08 1.06 2.07 0.86 1.06 1.04 1.15 3.22 3.93 4.12 6.20 1.13 6.07 3.00
Cl
HN
H
N
O
N
H
O
H
N
O
N
H
HO
O
O
f1 (ppm)

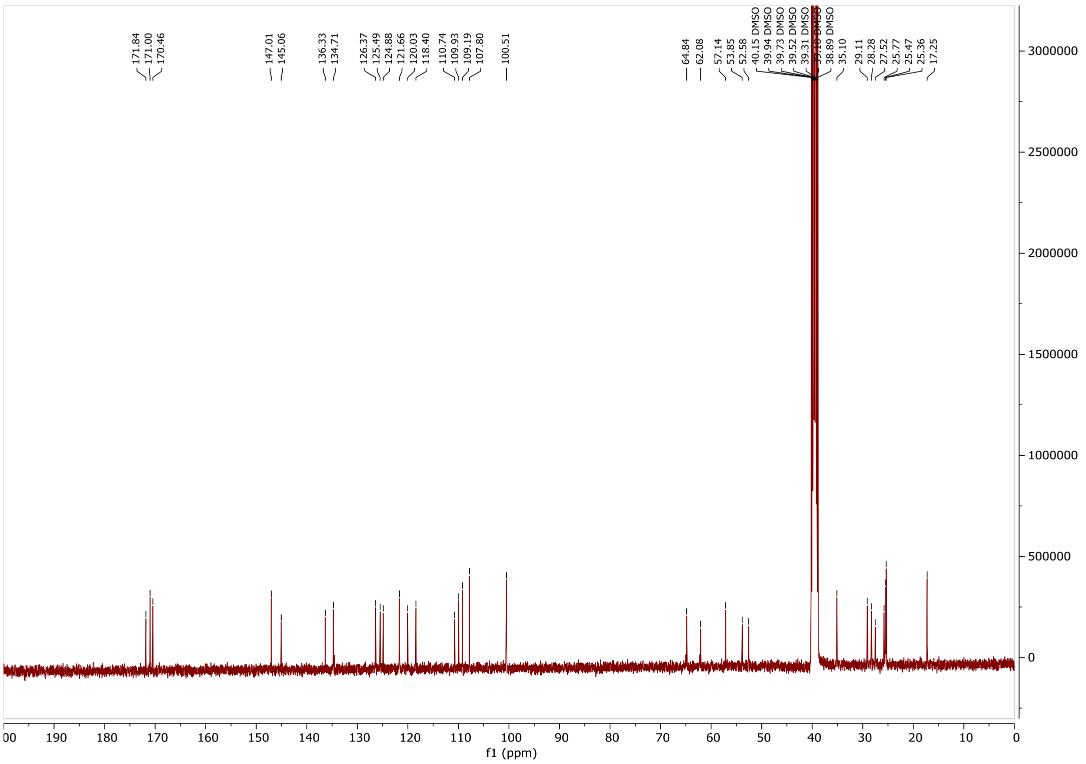

171.84 171.00 170.46 147.01 145.06 136.33 134.71 126.37 125.49 124.88 121.66 120.03 118.40 110.74 109.93 109.19 107.80 100.51 64.84 62.08 57.14 53.85 52.58 40.15 DMSO 39.94 DMSO 39.73 DMSO 39.52 DMSO 39.31 DMSO 39.10 DMSO 38.89 DMSO 35.10 29.11 28.28 27.52 25.77 25.47 25.36 17.25
f1 (ppm)

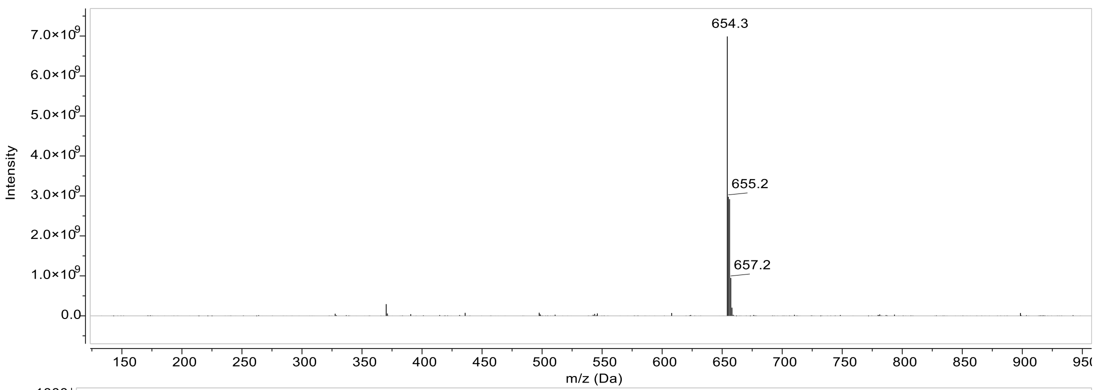

654.3
655.2
657.2
Intensity
m/z (Da)

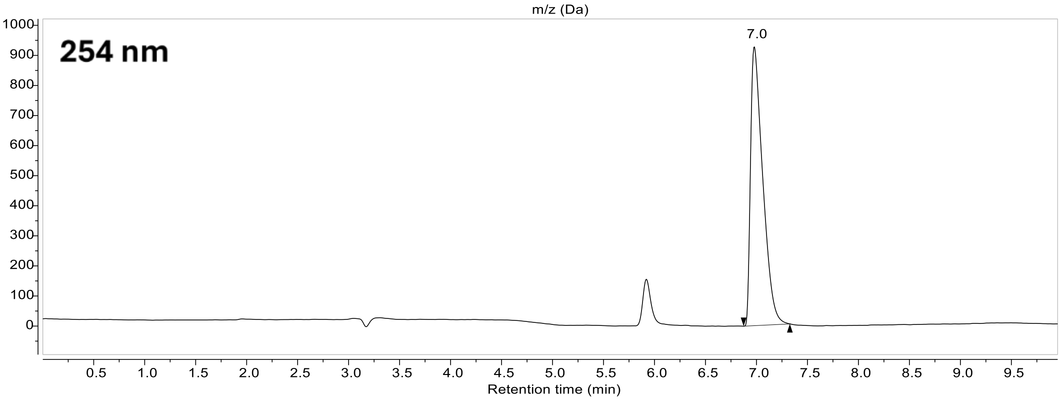

254 nm
7.0
Retention time (min)

#### Compound 5:

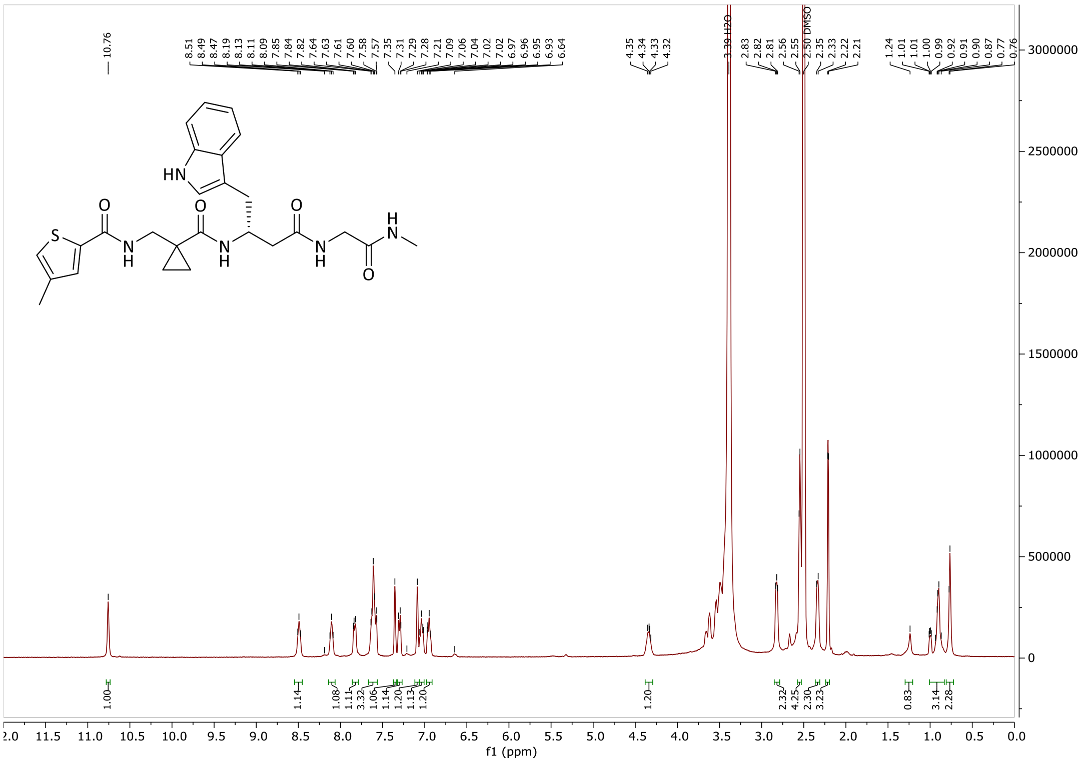

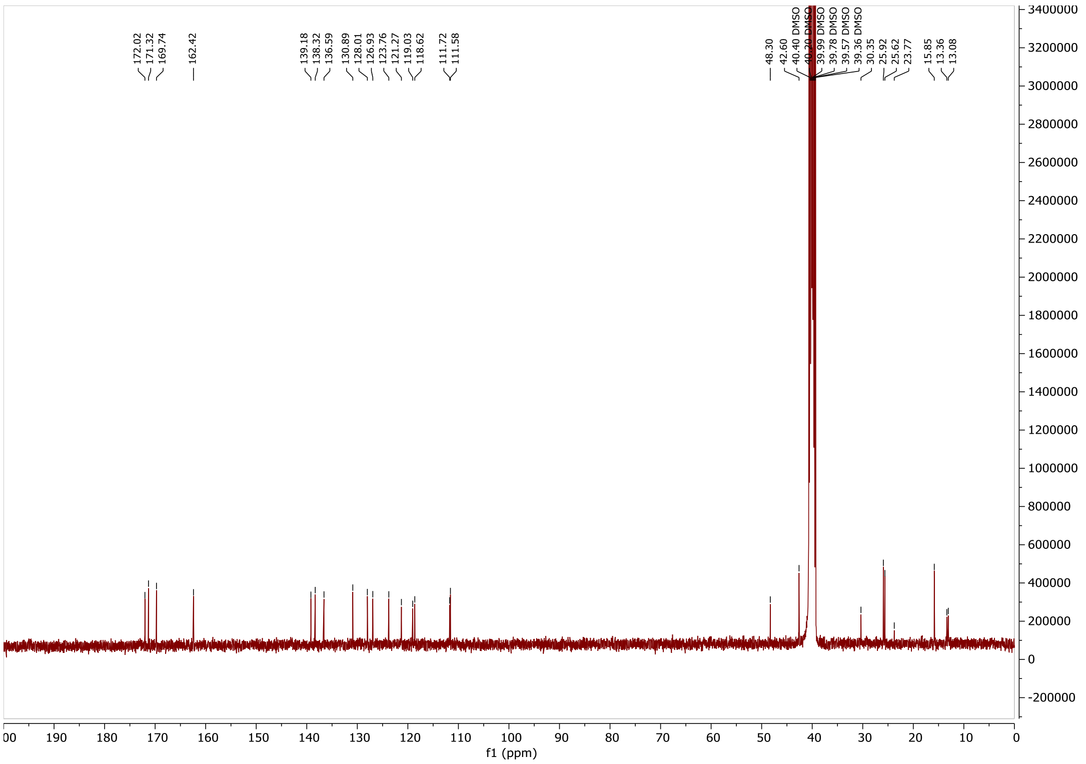

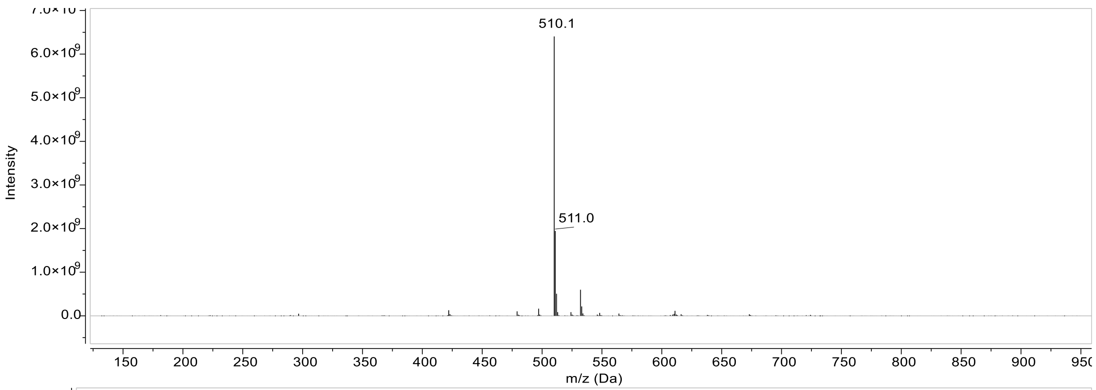

510.1
511.0
Intensity
6.0×10^9
5.0×10^9
4.0×10^9
3.0×10^9
2.0×10^9
1.0×10^9
0.0
150 200 250 300 350 400 450 500 550 600 650 700 750 800 850 900 950
m/z (Da)

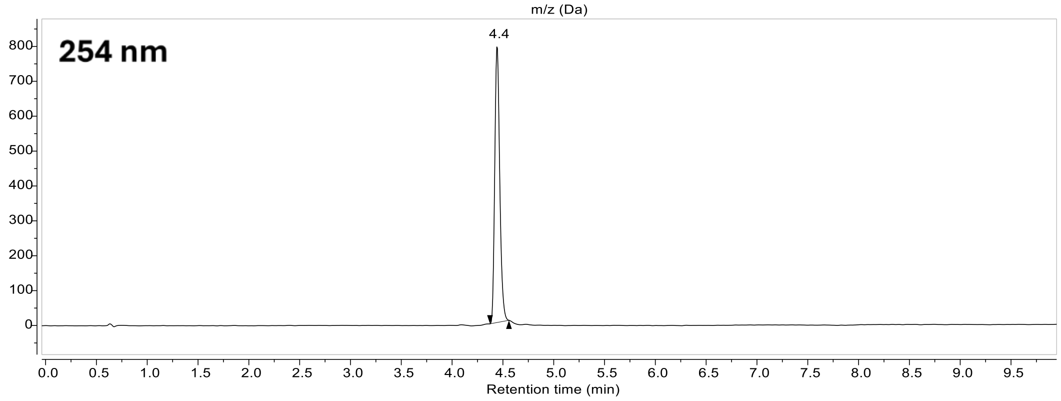

254 nm
4.4
800 700 600 500 400 300 200 100 0
0.0 0.5 1.0 1.5 2.0 2.5 3.0 3.5 4.0 4.5 5.0 5.5 6.0 6.5 7.0 7.5 8.0 8.5 9.0 9.5
Retention time (min)

#### Compound 10:

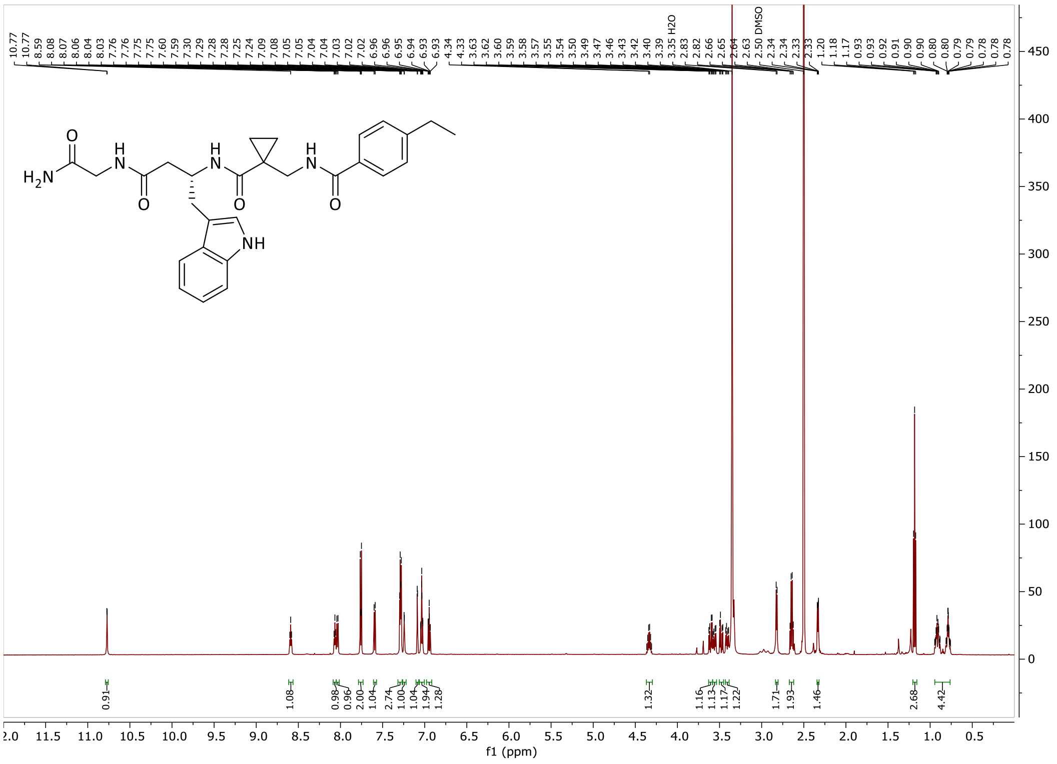

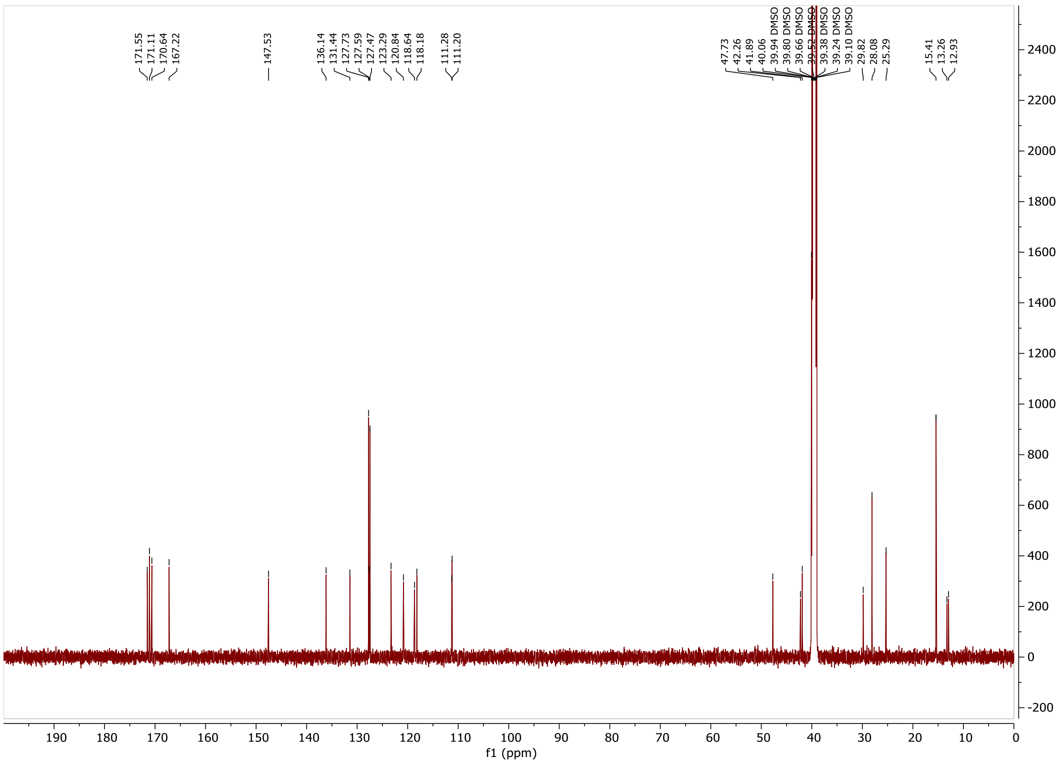

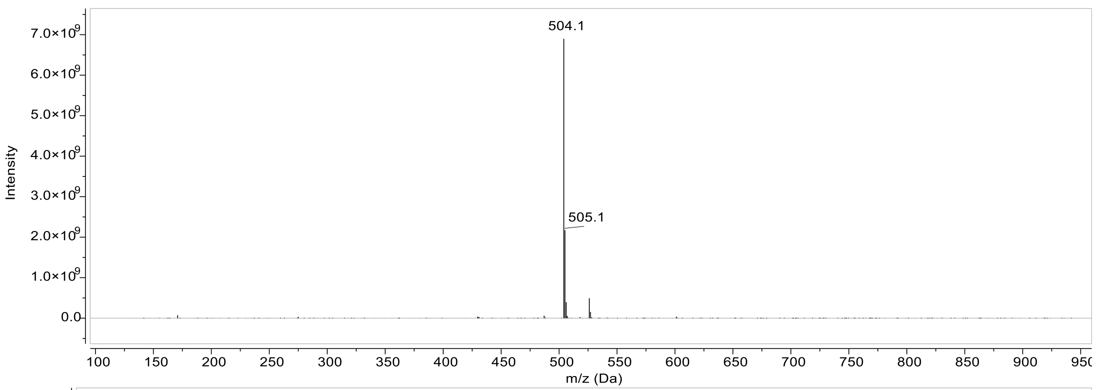

504.1
505.1
Intensity
m/z (Da)

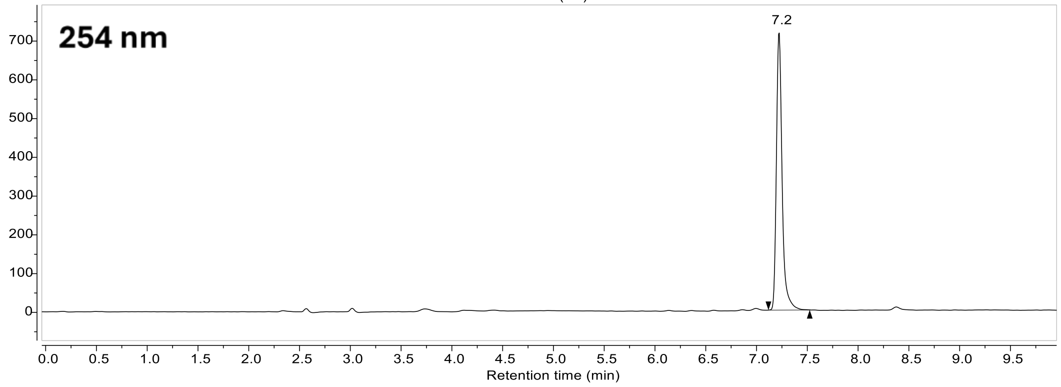

254 nm
7.2
Retention time (min)

#### Compound 18:

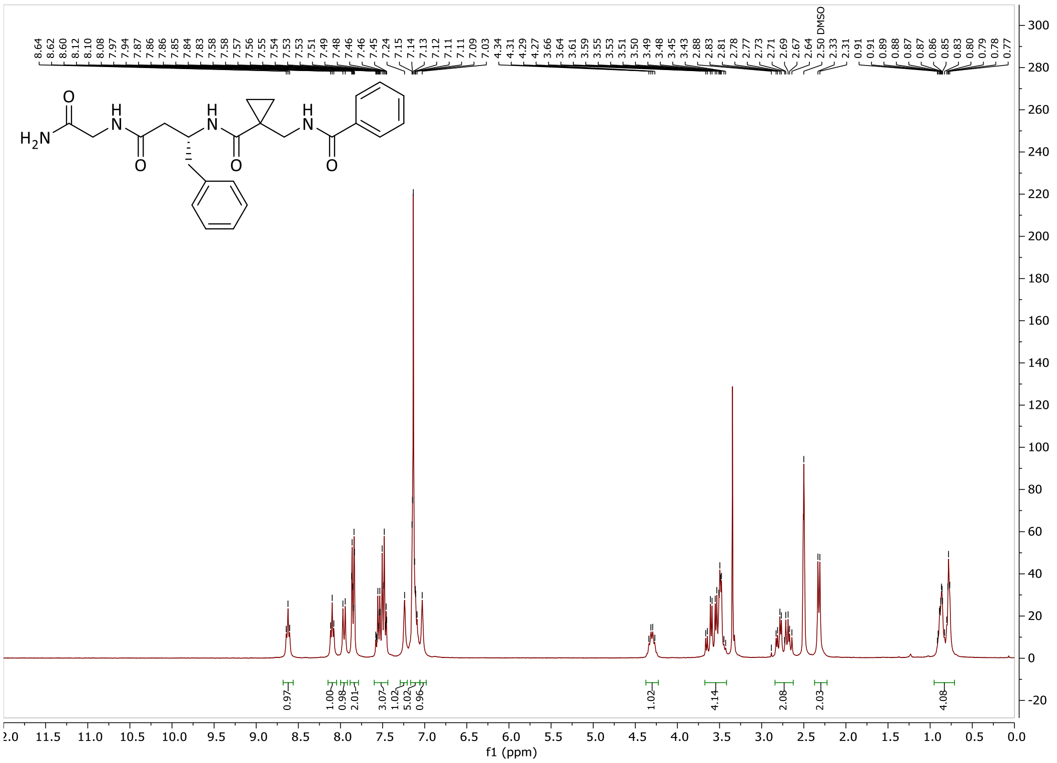

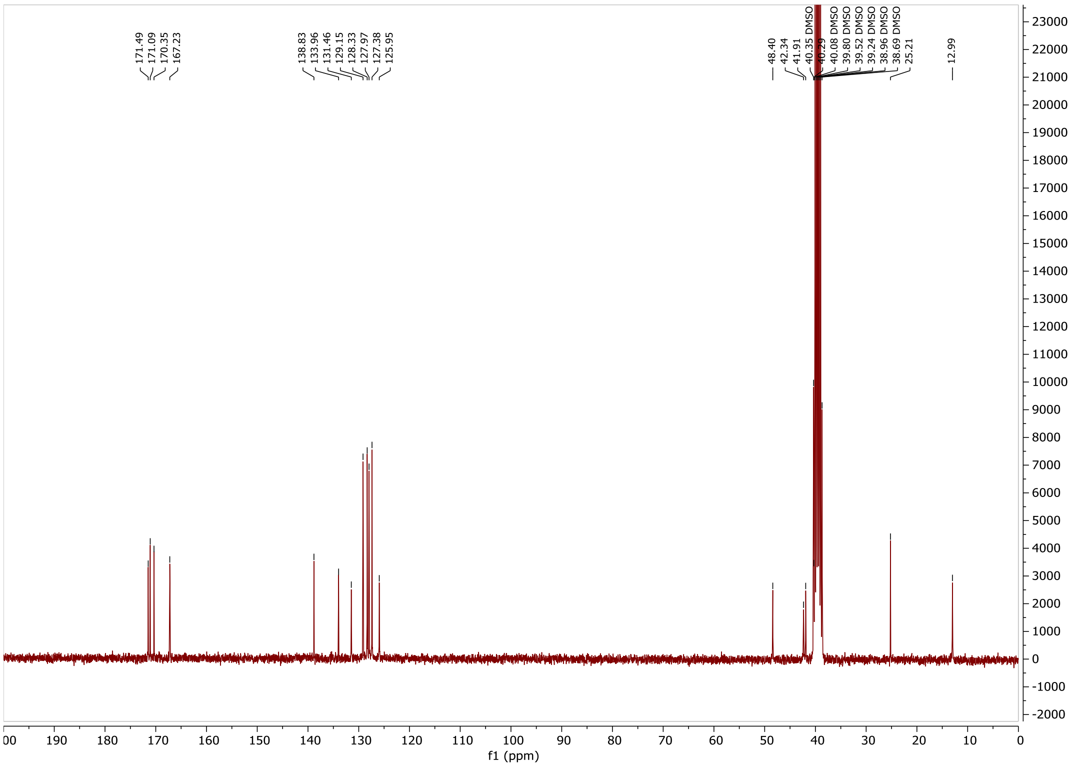

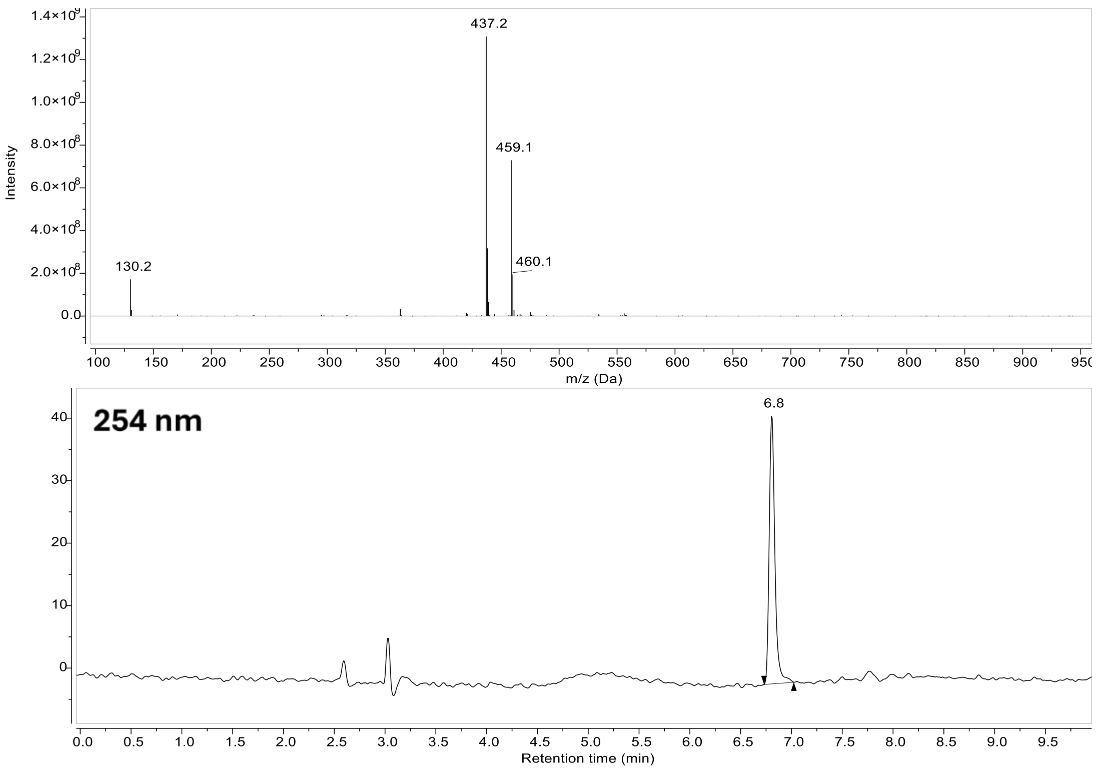

437.2
459.1
460.1
130.2
Intensity
m/z (Da)
254 nm
6.8
Retention time (min)

#### Compound 11:

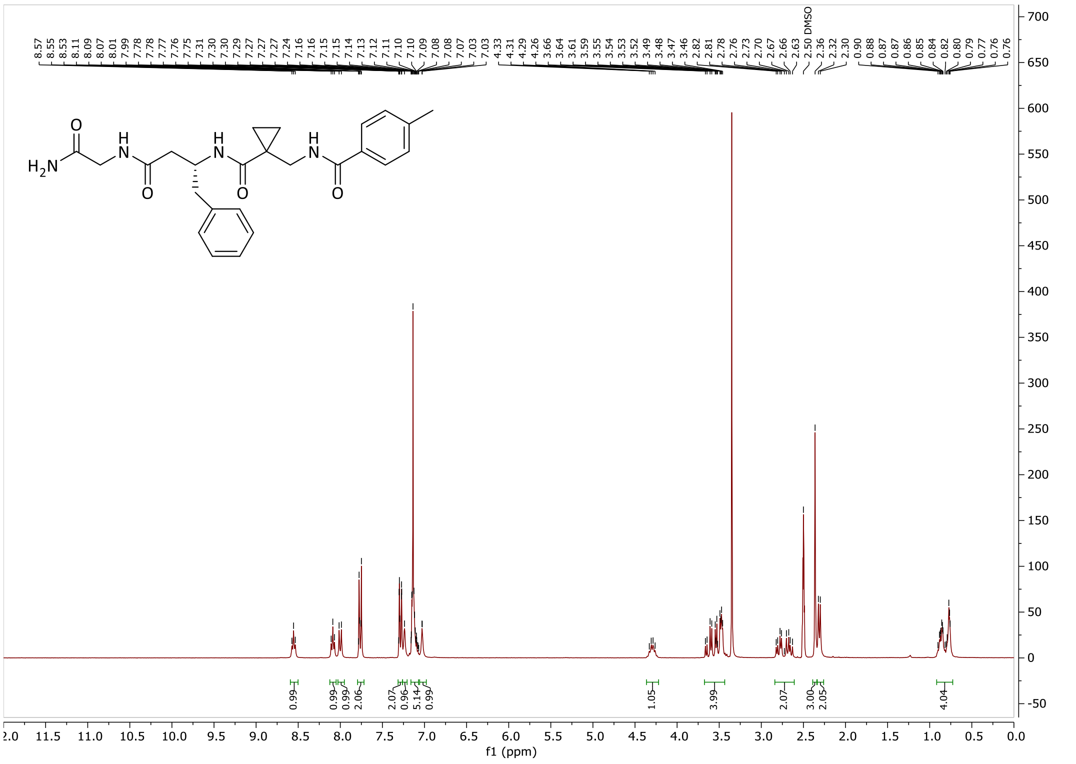

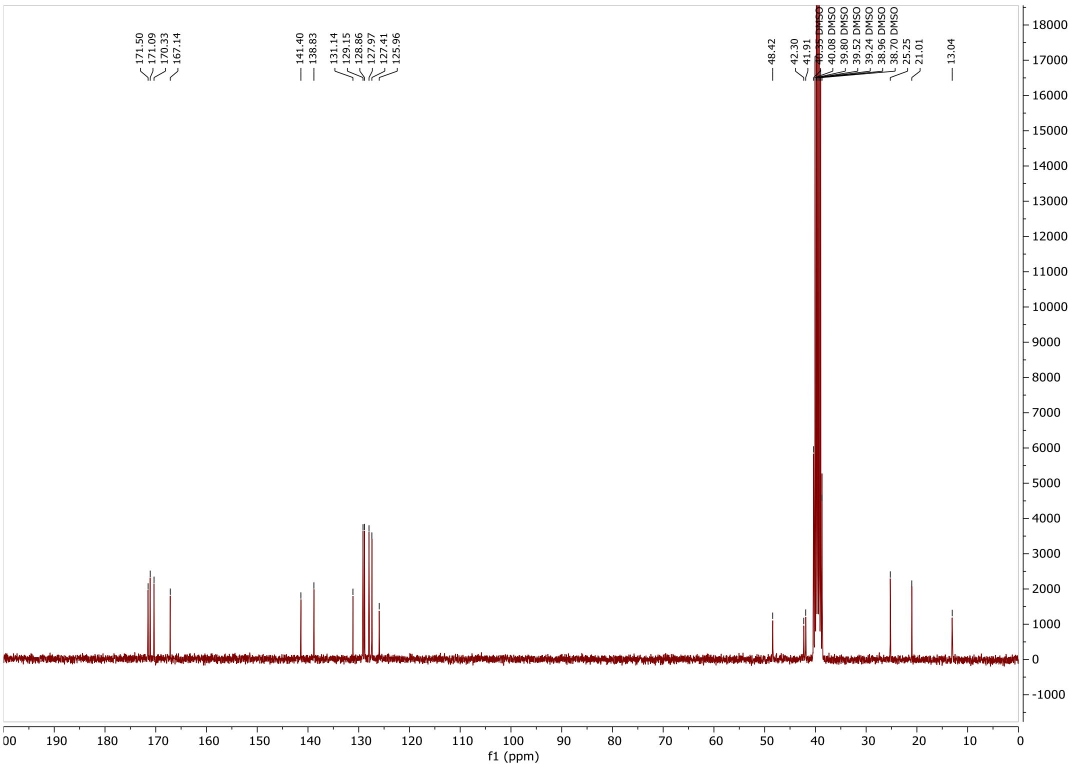

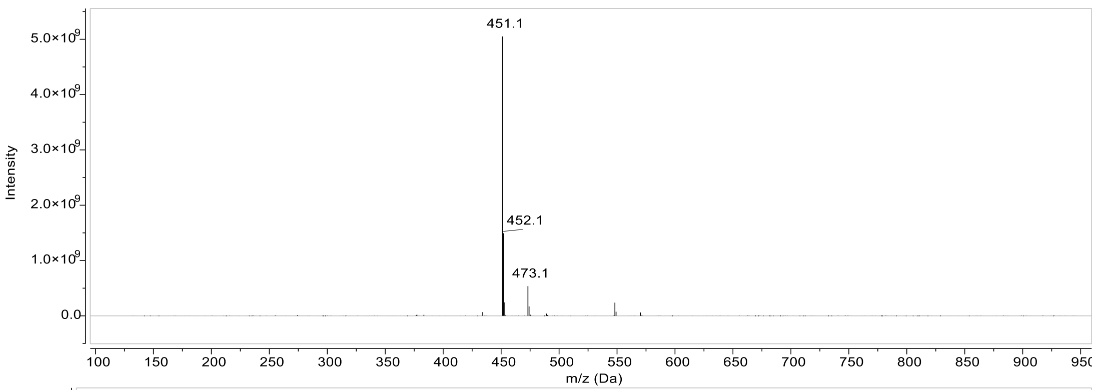
451.1
452.1
473.1
5.0×10^9
4.0×10^9
3.0×10^9
2.0×10^9
1.0×10^9
0.0
Intensity
100 150 200 250 300 350 400 450 500 550 600 650 700 750 800 850 900 950
m/z (Da)

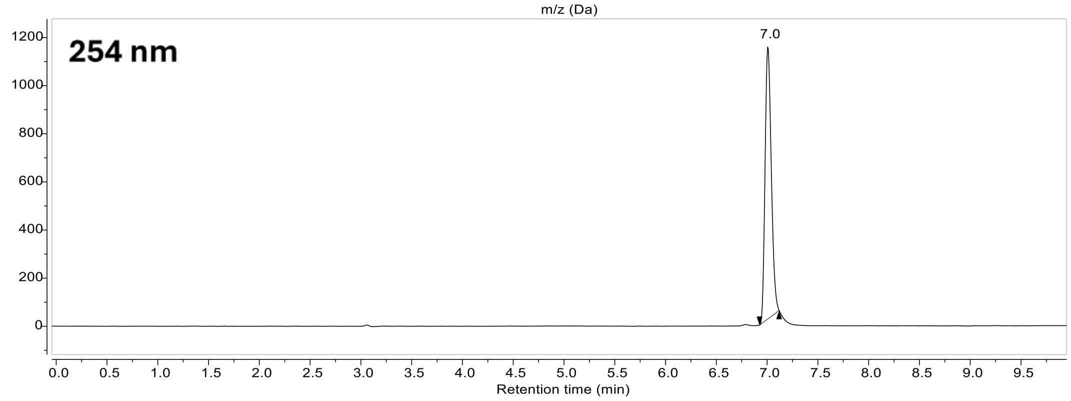
254 nm
7.0
1200
1000
800
600
400
200
0
0.0 0.5 1.0 1.5 2.0 2.5 3.0 3.5 4.0 4.5 5.0 5.5 6.0 6.5 7.0 7.5 8.0 8.5 9.0 9.5
Retention time (min)

#### Compound 12:

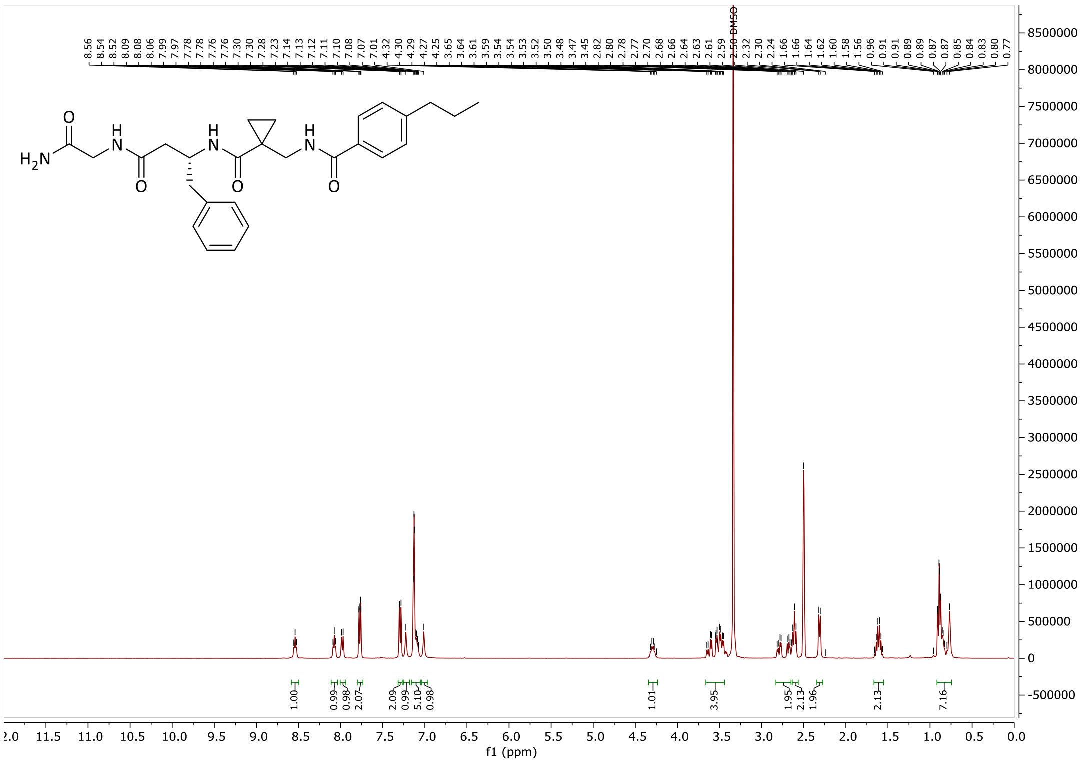

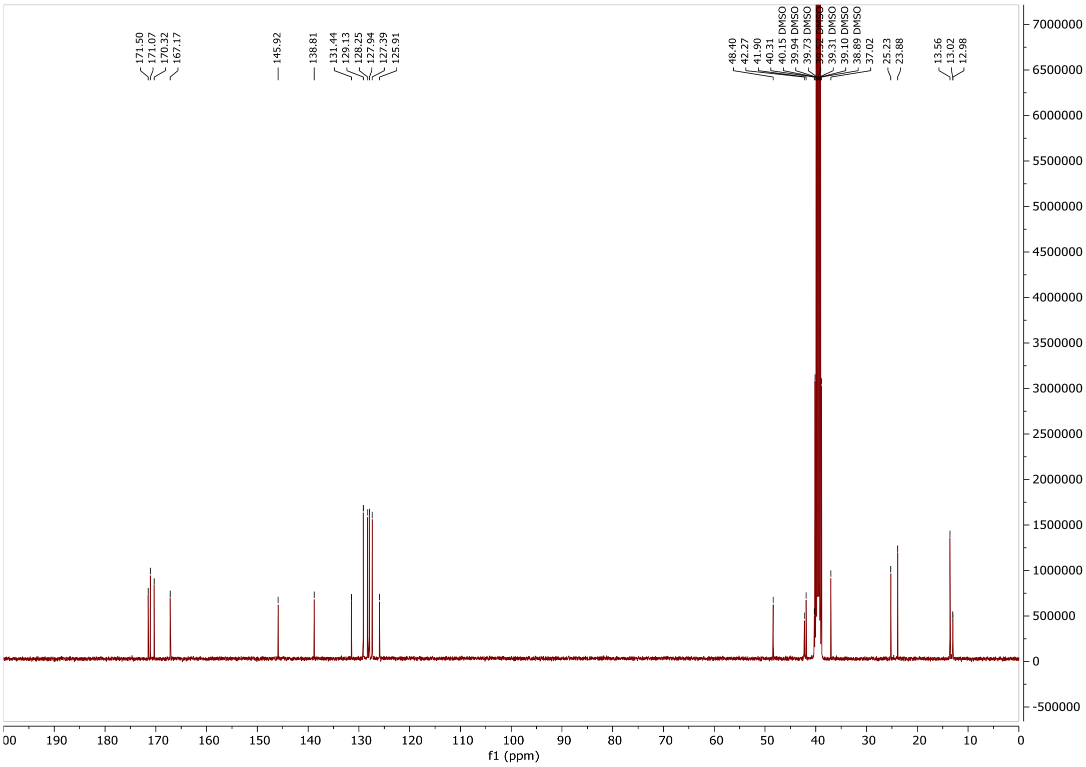

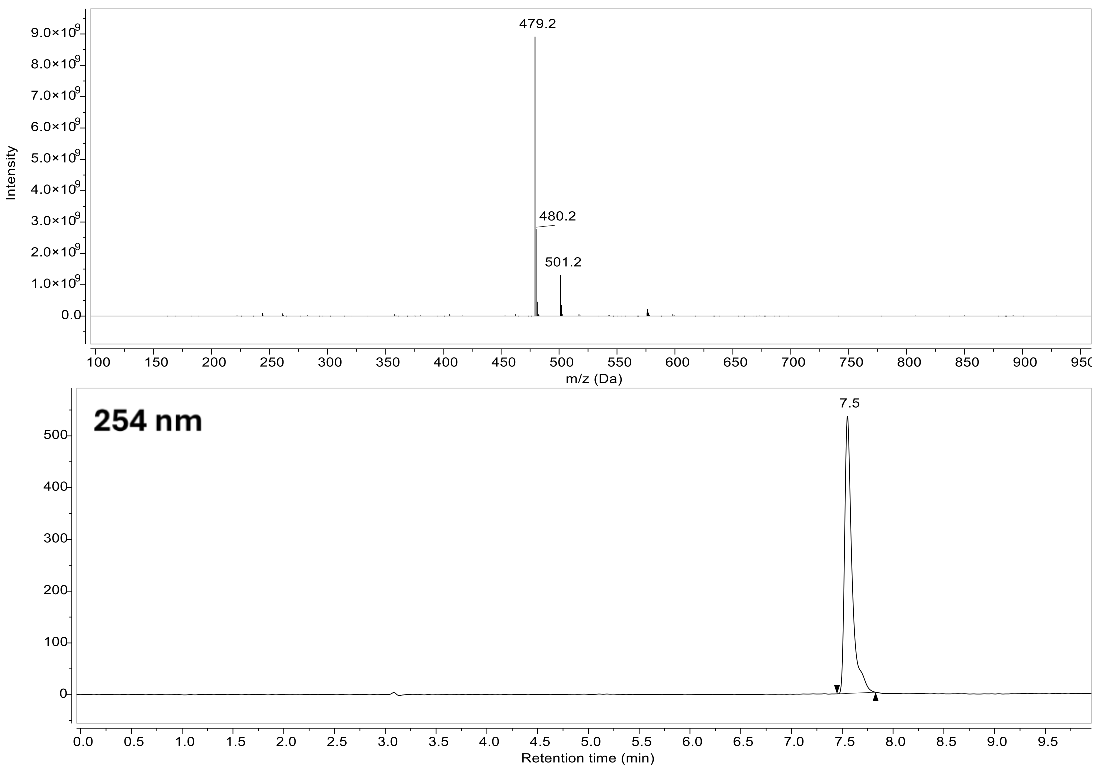

479.2
480.2
501.2
Intensity
m/z (Da)
254 nm
7.5
Retention time (min)

**Compound 13:**

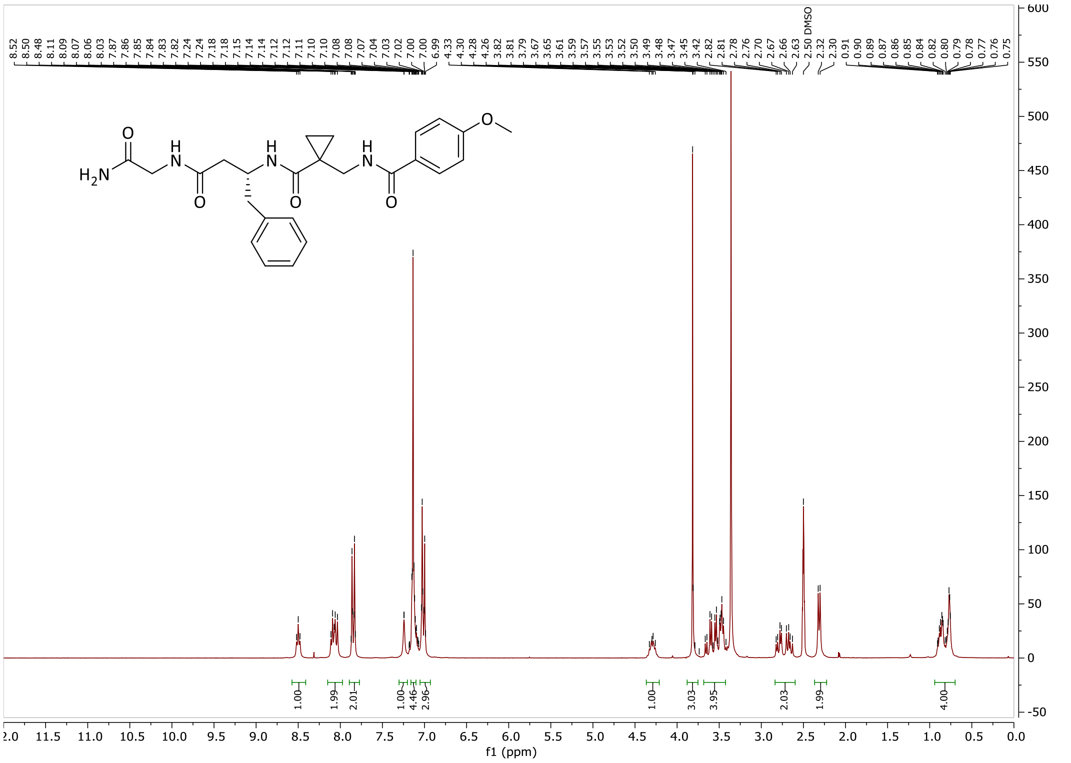

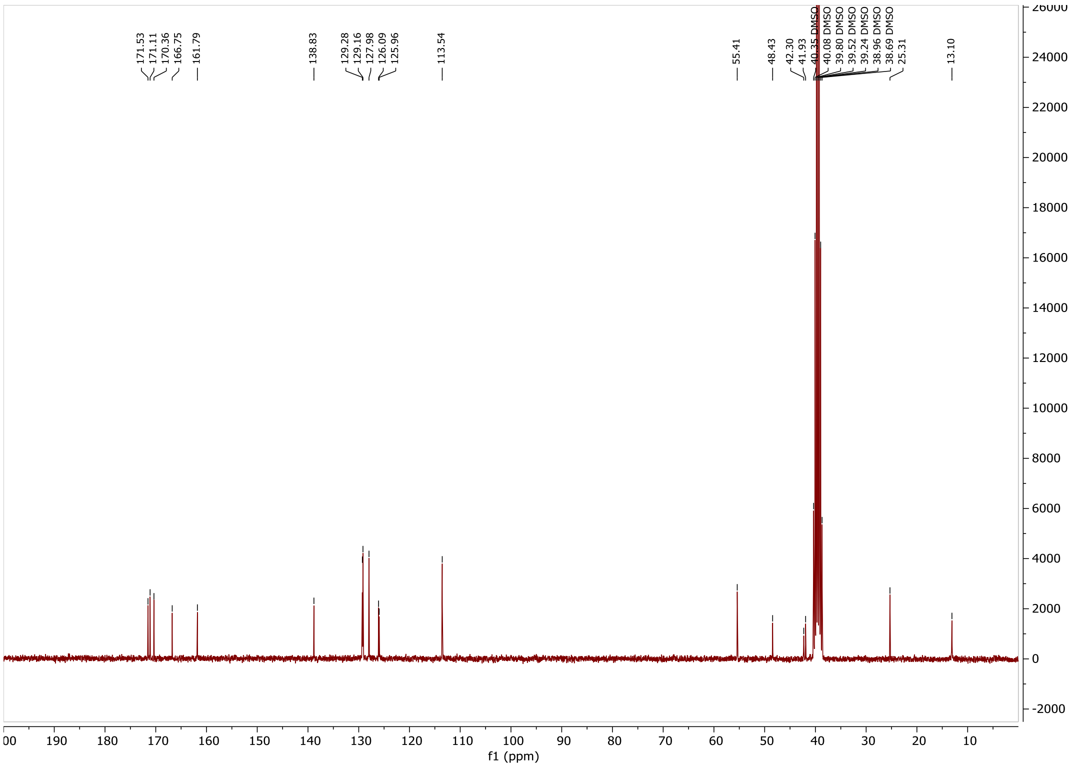

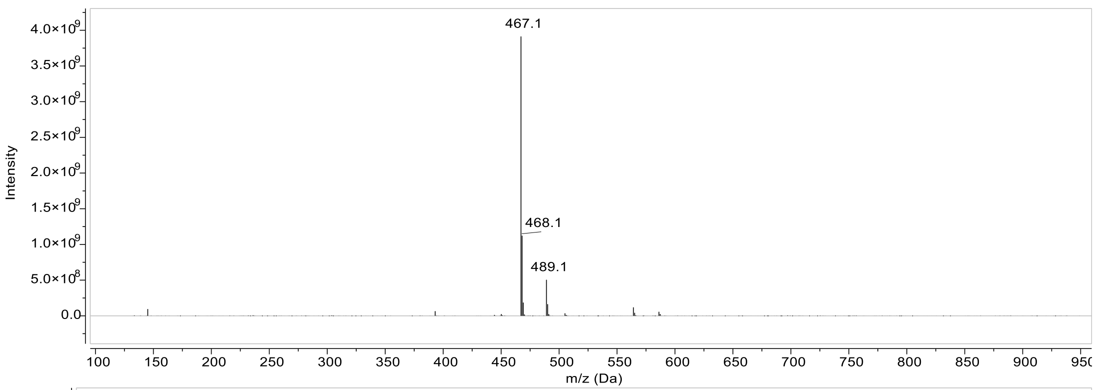

467.1
468.1
489.1
Intensity
m/z (Da)

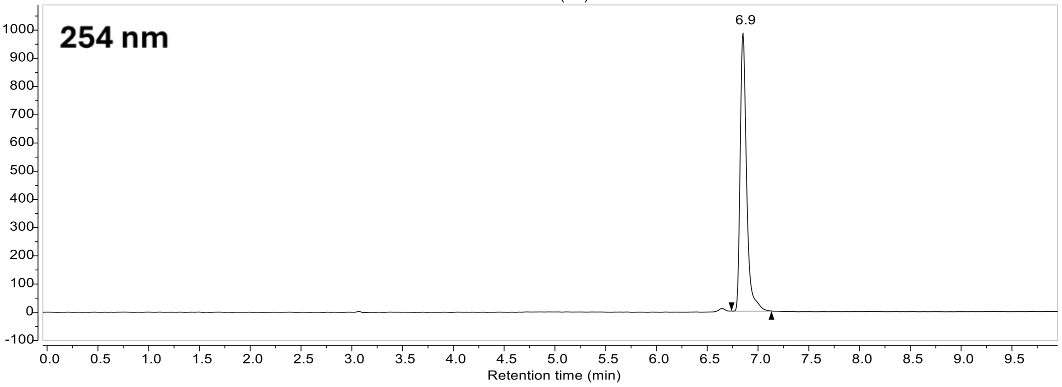

254 nm
6.9
Retention time (min)

#### Compound 19:

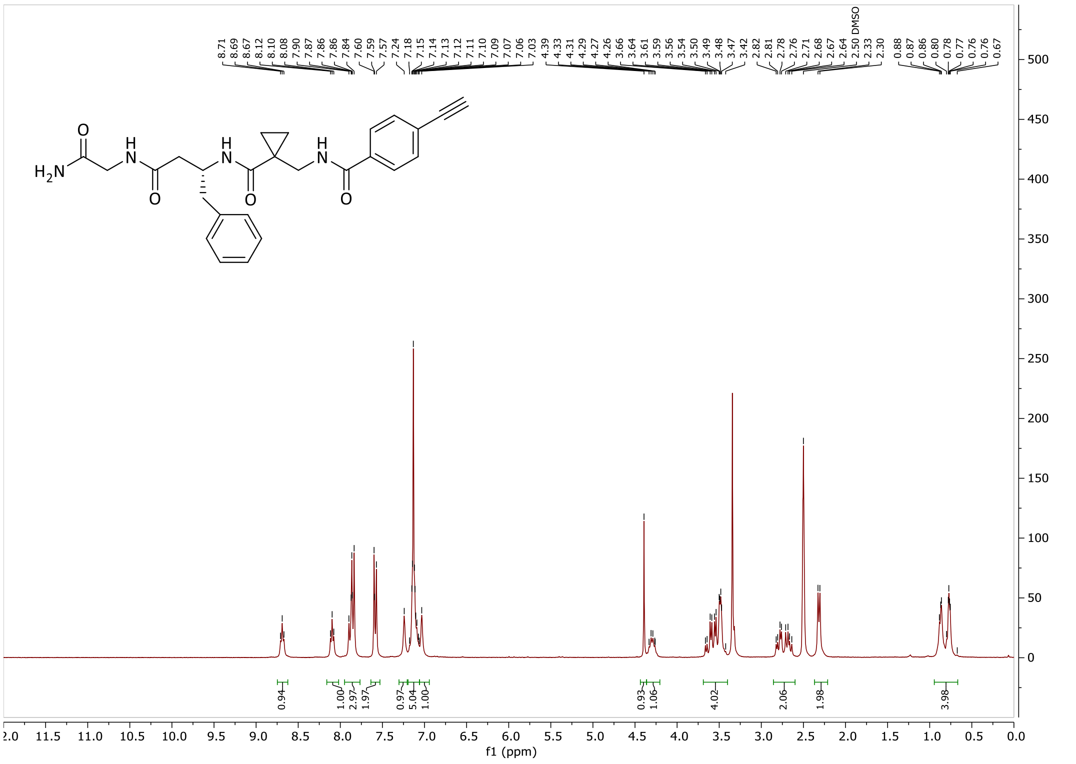

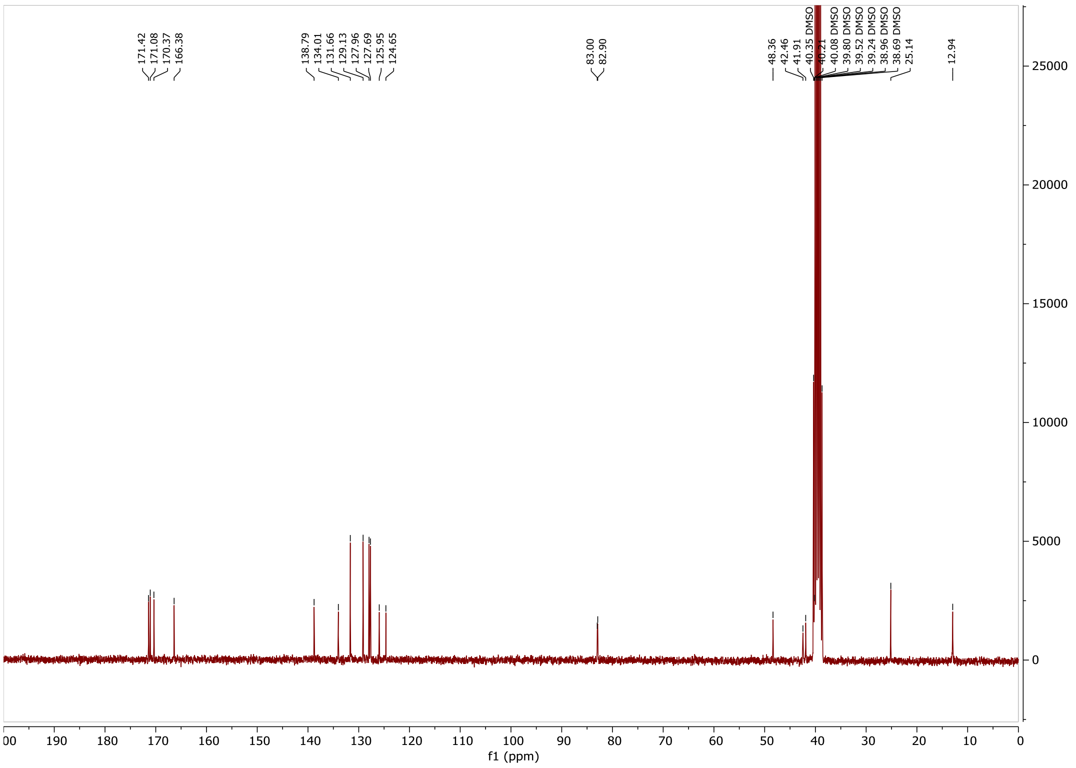

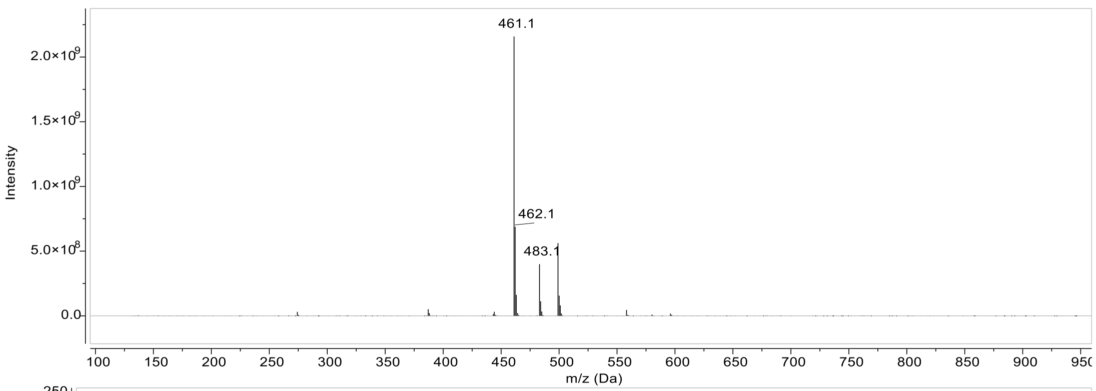
461.1
462.1
483.1
Intensity
m/z (Da)

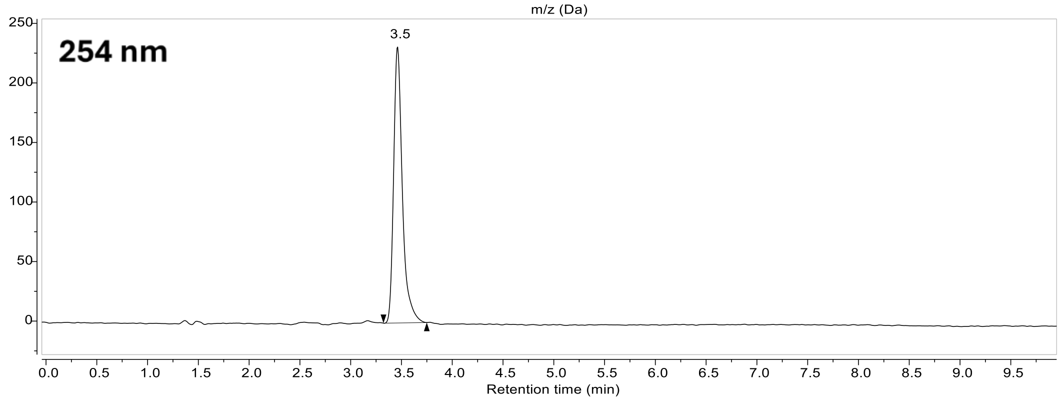
254 nm
3.5
Retention time (min)

#### Compound 20:

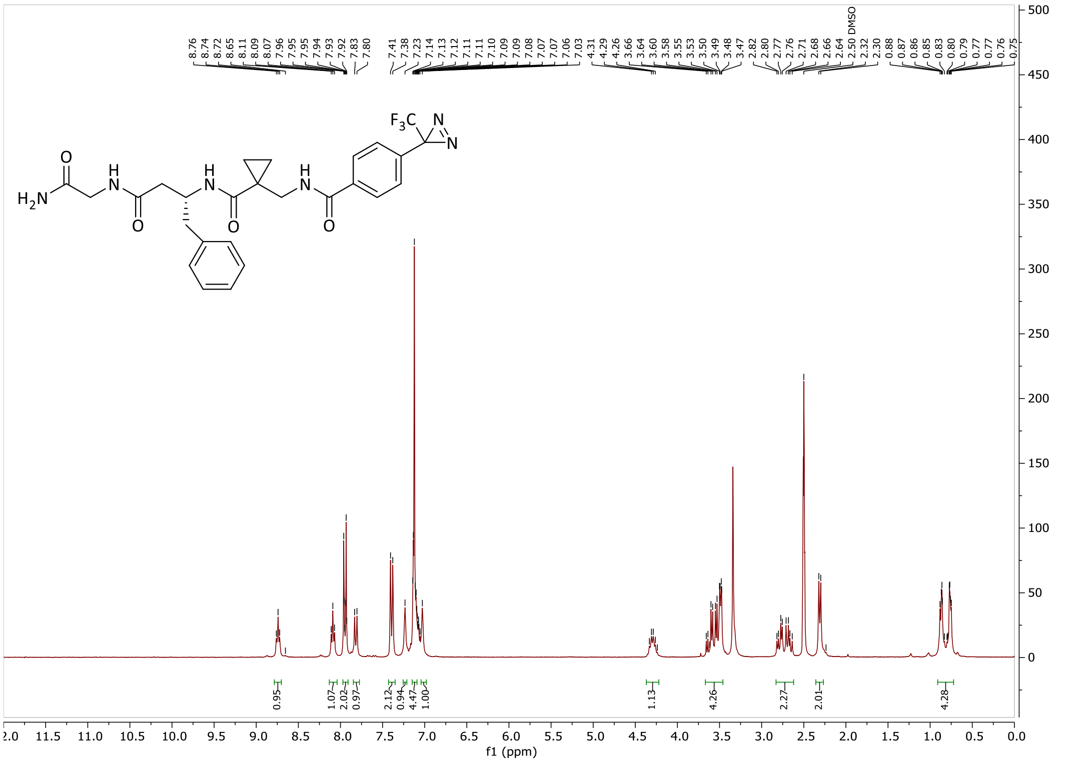

8.76
8.74
8.72
8.65
8.11
8.09
8.07
7.96
7.95
7.95
7.94
7.93
7.92
7.83
7.80
7.41
7.38
7.23
7.14
7.13
7.12
7.11
7.11
7.10
7.09
7.09
7.08
7.07
7.07
7.06
7.03
4.31
4.29
4.26
3.66
3.60
3.58
3.55
3.53
3.50
3.49
3.48
3.47
2.82
2.80
2.77
2.76
2.71
2.68
2.66
2.64
2.50 DMSO
2.32
2.30
0.88
0.87
0.86
0.85
0.83
0.80
0.79
0.77
0.77
0.76
0.75
$F_3C$
N
N
O
$H_2N$
H
N
O
H
N
O
H
N
O
0.95
1.07
2.02
0.97
2.12
0.94
4.47
1.00
1.13
4.26
2.27
2.01
4.28
f1 (ppm)

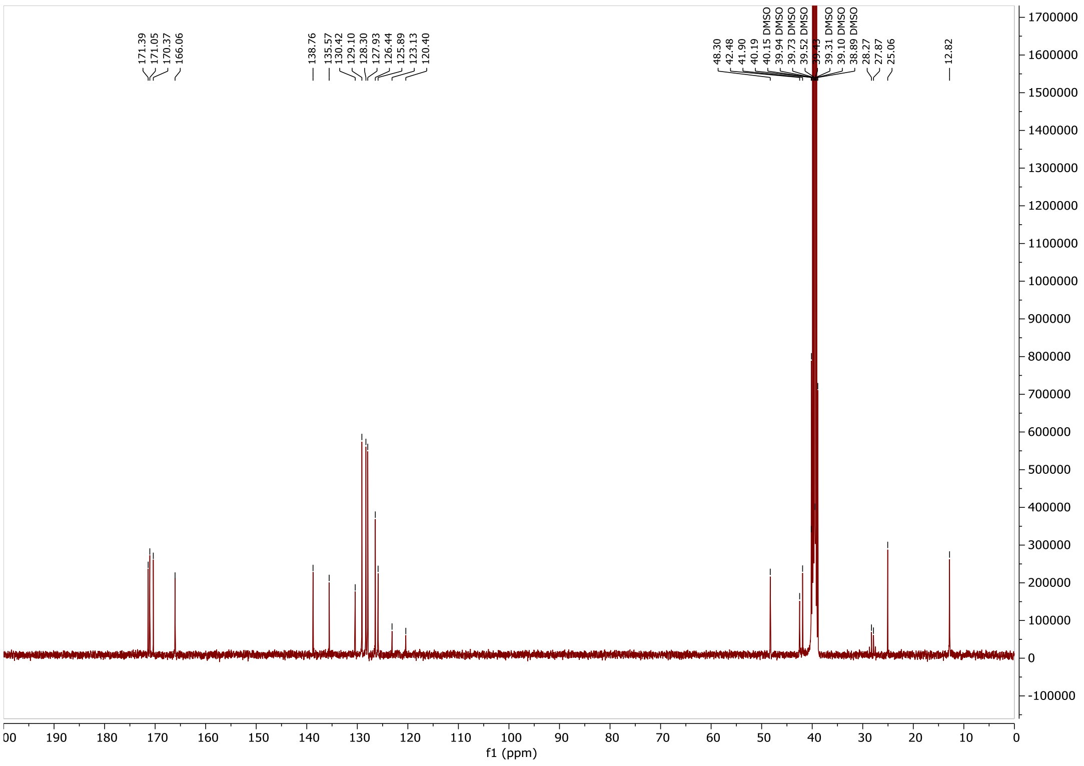

171.39
171.05
170.37
166.06
138.76
135.57
130.42
129.10
128.30
127.93
126.44
125.89
123.13
120.40
48.30
42.48
41.90
40.19
40.15 DMSO
39.94 DMSO
39.73 DMSO
39.52 DMSO
39.31 DMSO
39.10 DMSO
38.89 DMSO
28.27
27.87
25.06
12.82
f1 (ppm)

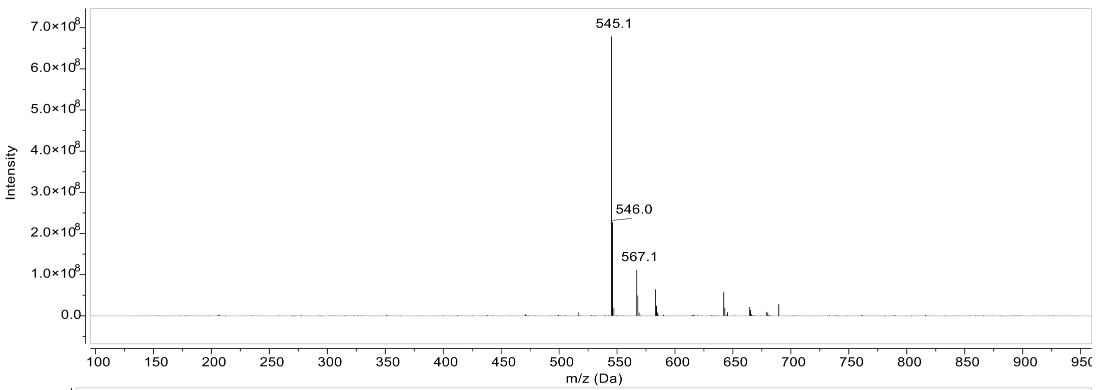
545.1
546.0
567.1
Intensity
m/z (Da)

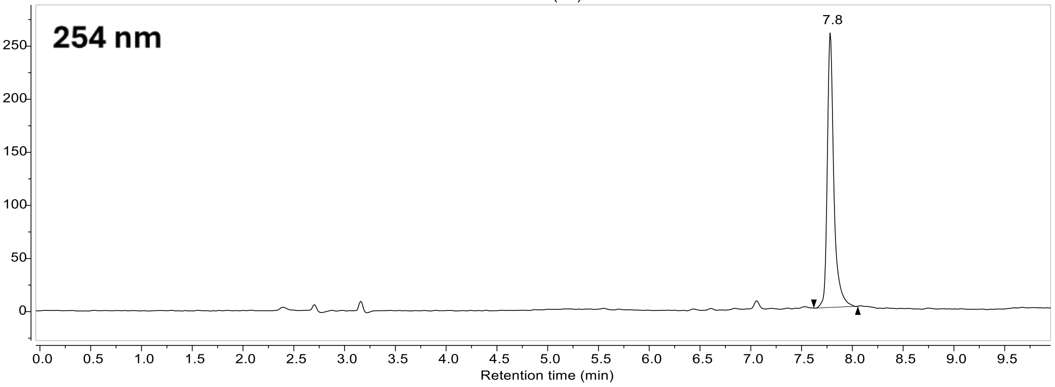
254 nm
7.8
Retention time (min)

#### Compound 21:

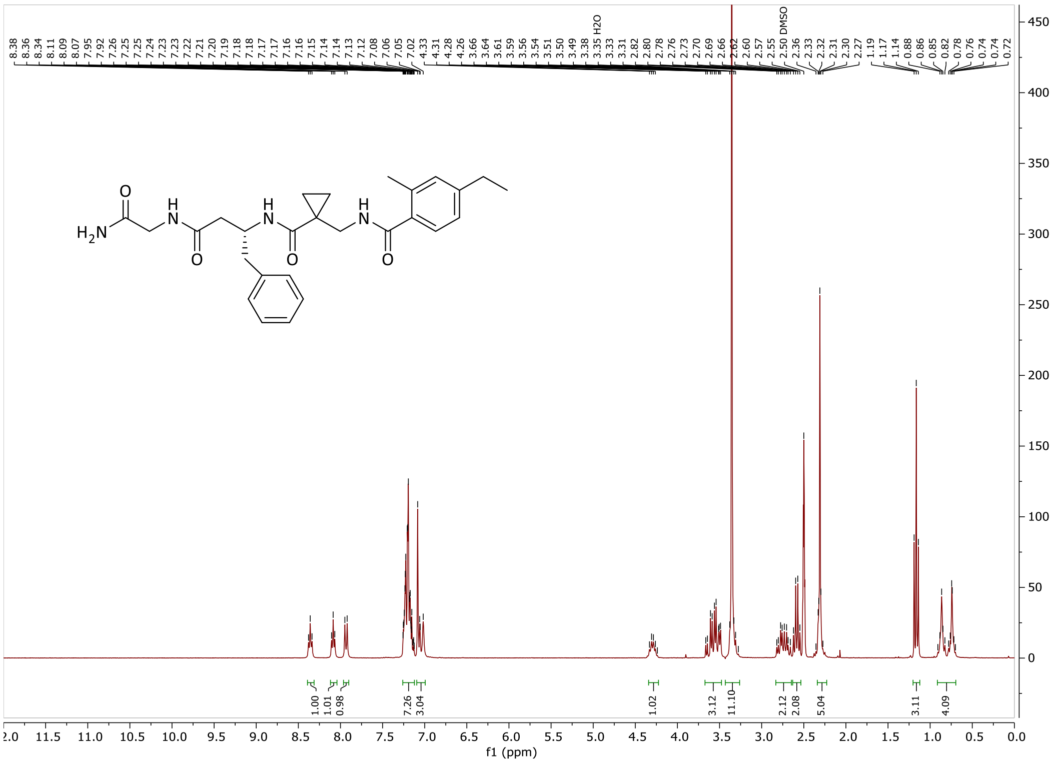

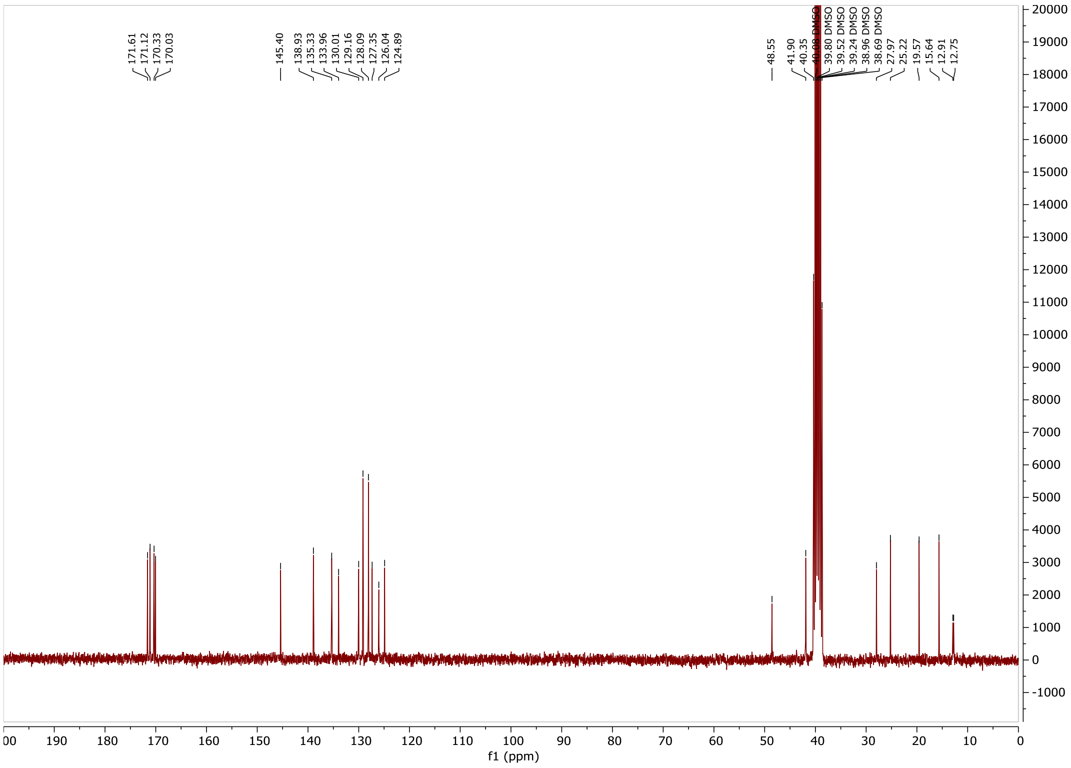

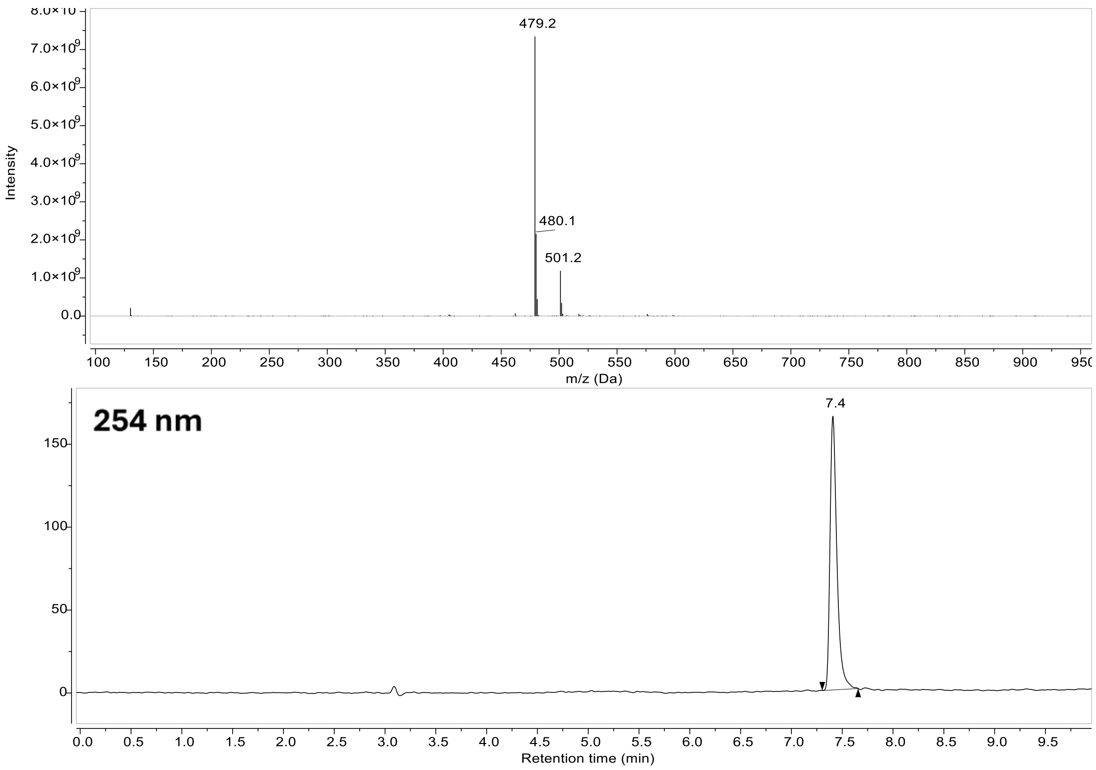

479.2
480.1
501.2
Intensity
m/z (Da)
254 nm
7.4
Retention time (min)

**Compound 22:**

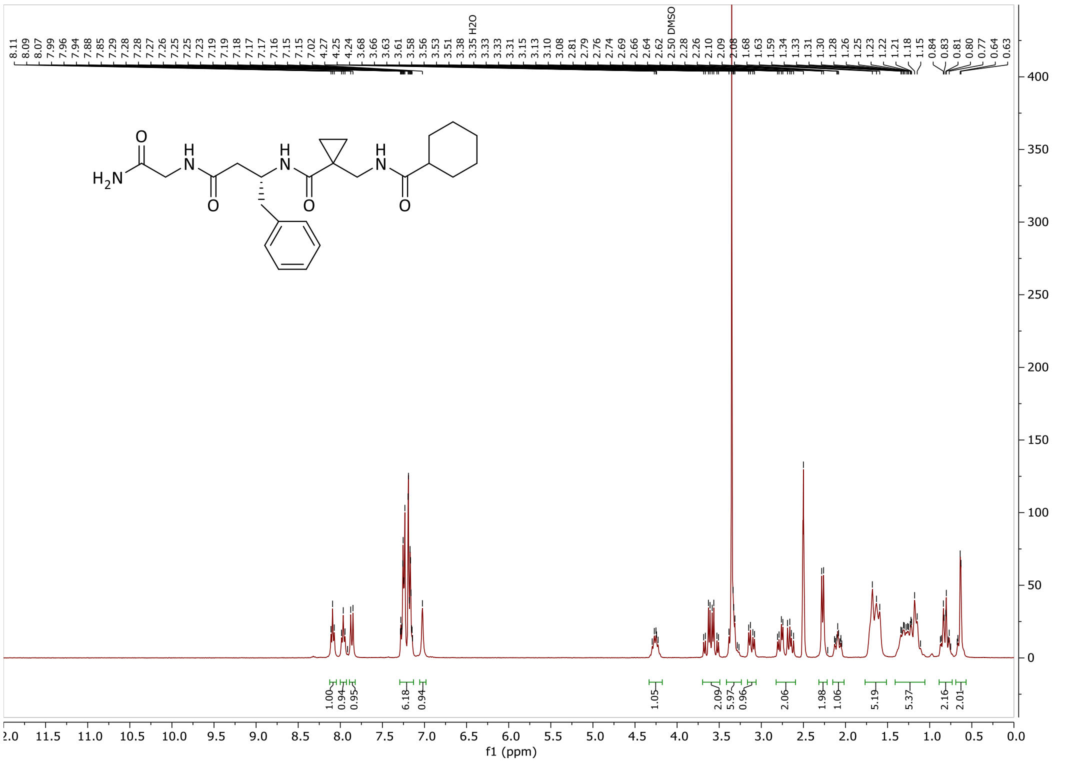

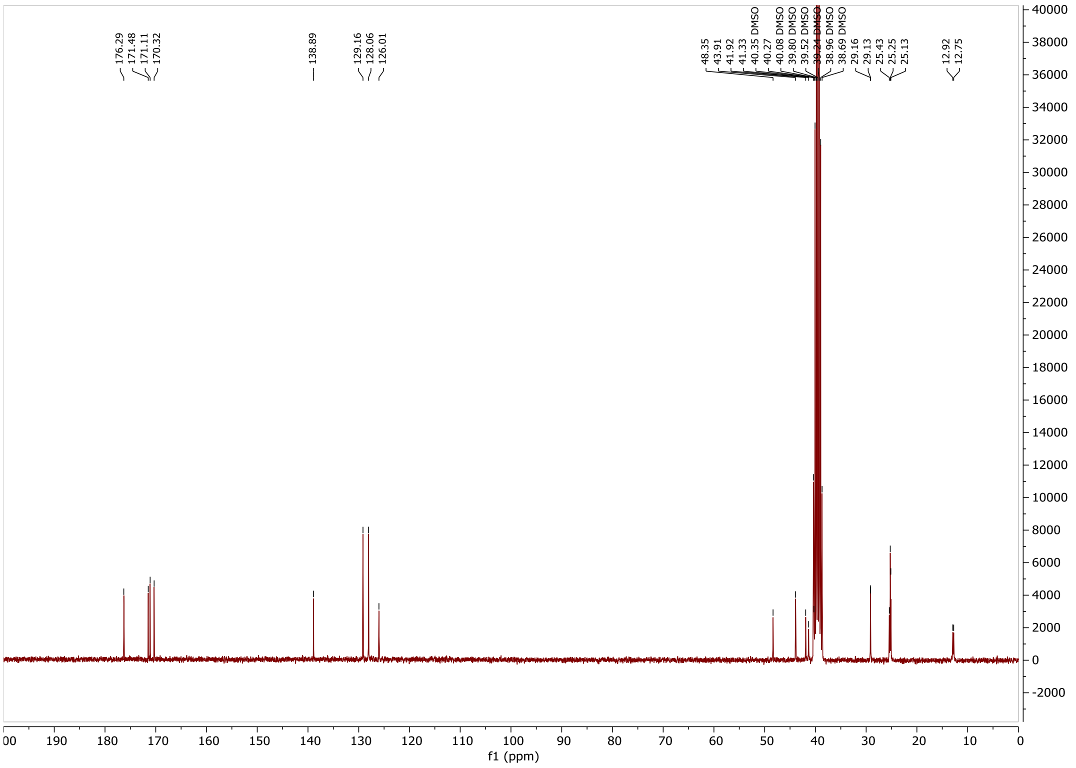

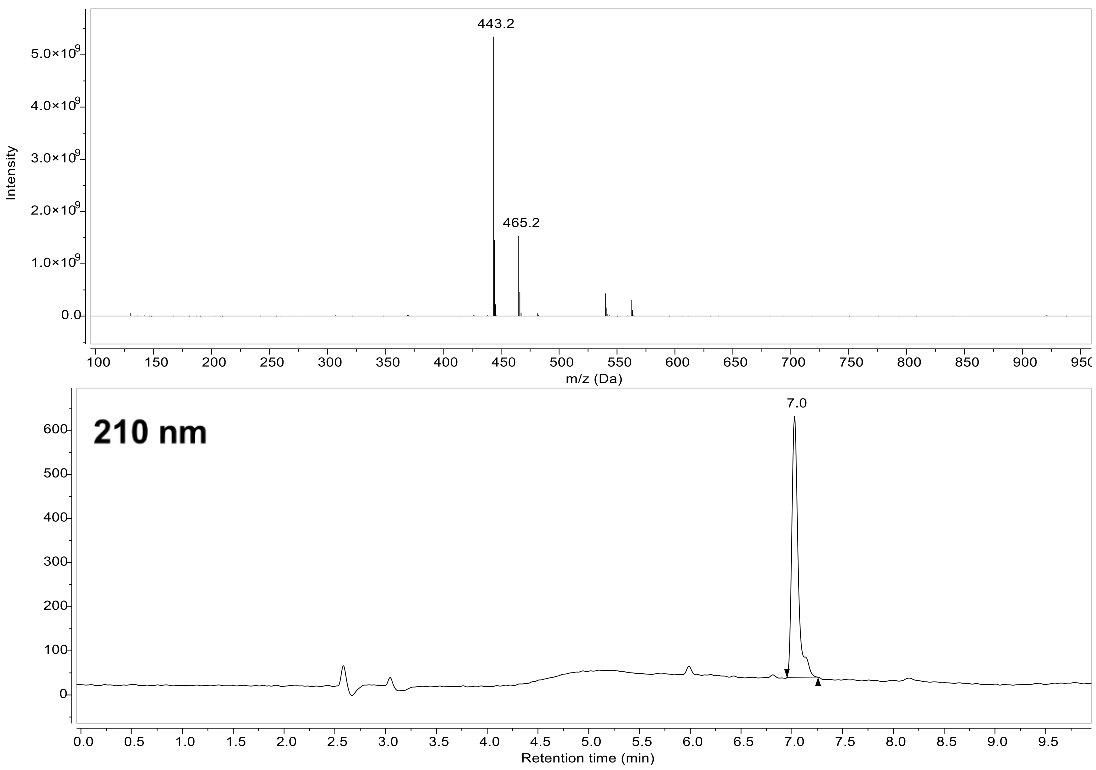
443.2
465.2
Intensity
m/z (Da)
210 nm
7.0
Retention time (min)

### Compound 23:

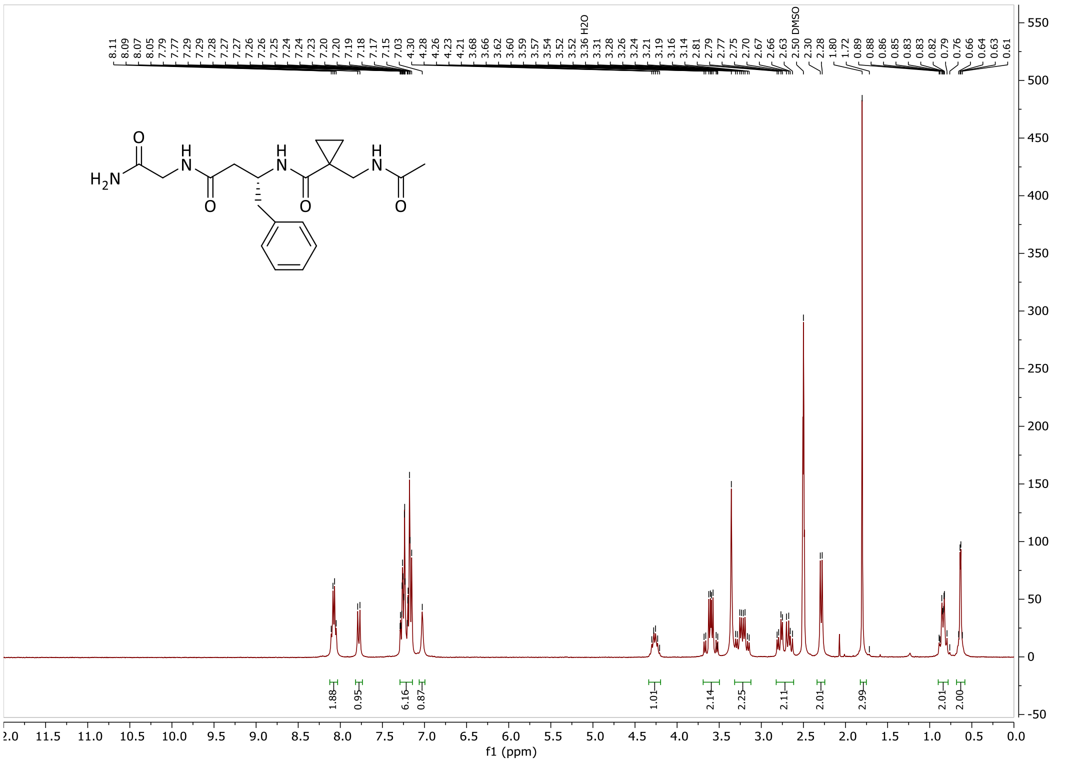
8.11 8.09 8.07 8.05 7.79 7.77 7.29 7.29 7.28 7.27 7.27 7.26 7.26 7.25 7.24 7.24 7.23 7.20 7.20 7.19 7.18 7.17 7.15 7.03 4.30 4.28 4.26 4.23 4.21 3.68 3.66 3.62 3.60 3.59 3.57 3.54 3.52 3.52 3.36 H2O 3.31 3.28 3.26 3.24 3.21 3.19 3.16 3.14 2.81 2.79 2.77 2.75 2.70 2.67 2.66 2.63 2.50 DMSO 2.30 2.28 1.80 1.72 0.89 0.88 0.86 0.85 0.83 0.83 0.82 0.79 0.76 0.66 0.64 0.63 0.61
1.88 0.95 6.16 0.87 1.01 2.14 2.25 2.11 2.01 2.99 2.01 2.00
f1 (ppm)

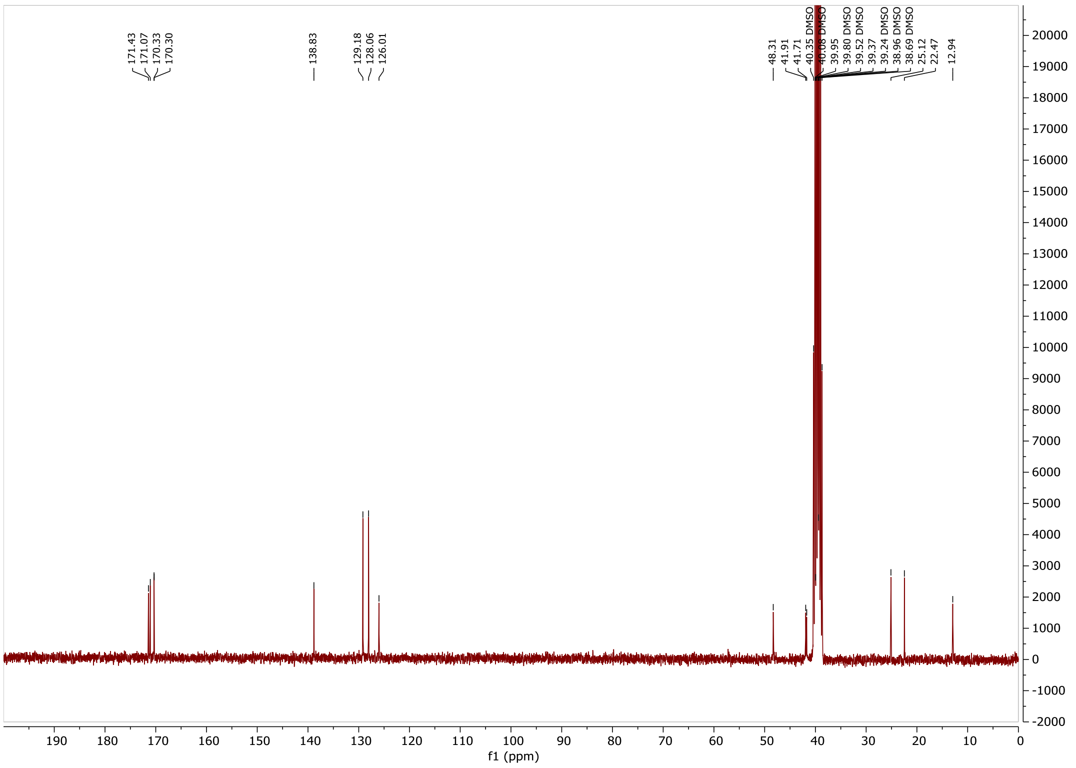
171.43 171.07 170.33 170.30 138.83 129.18 128.06 126.01 48.31 41.91 41.71 40.35 DMSO 40.08 DMSO 39.95 39.80 DMSO 39.52 DMSO 39.37 39.24 DMSO 38.96 DMSO 38.69 DMSO 25.12 22.47 12.94
f1 (ppm)

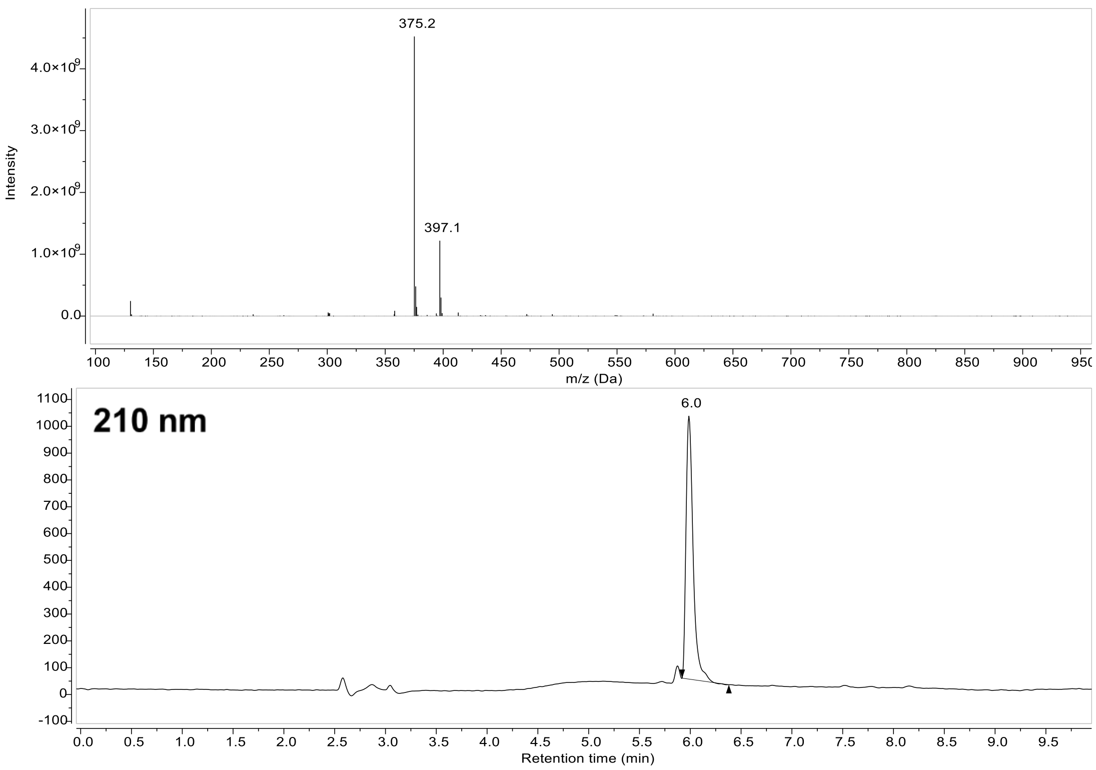

375.2
397.1
Intensity
4.0×10⁹
3.0×10⁹
2.0×10⁹
1.0×10⁹
0.0
m/z (Da)
210 nm
6.0
Retention time (min)

**Compound 24**:

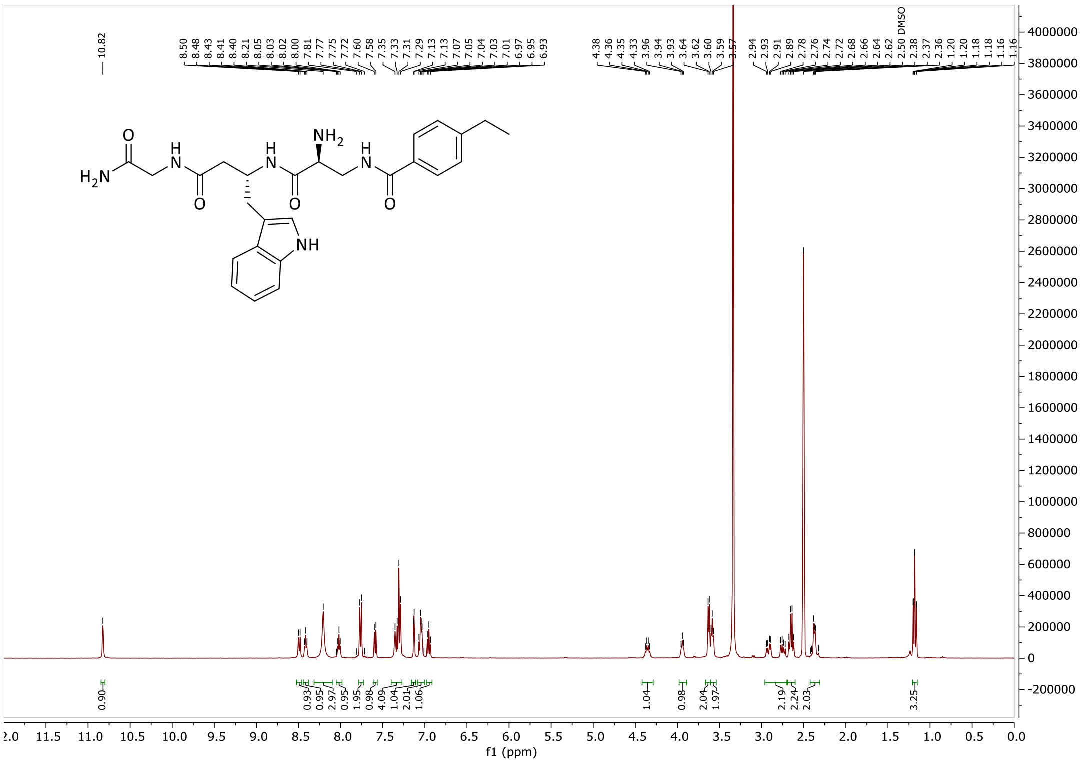

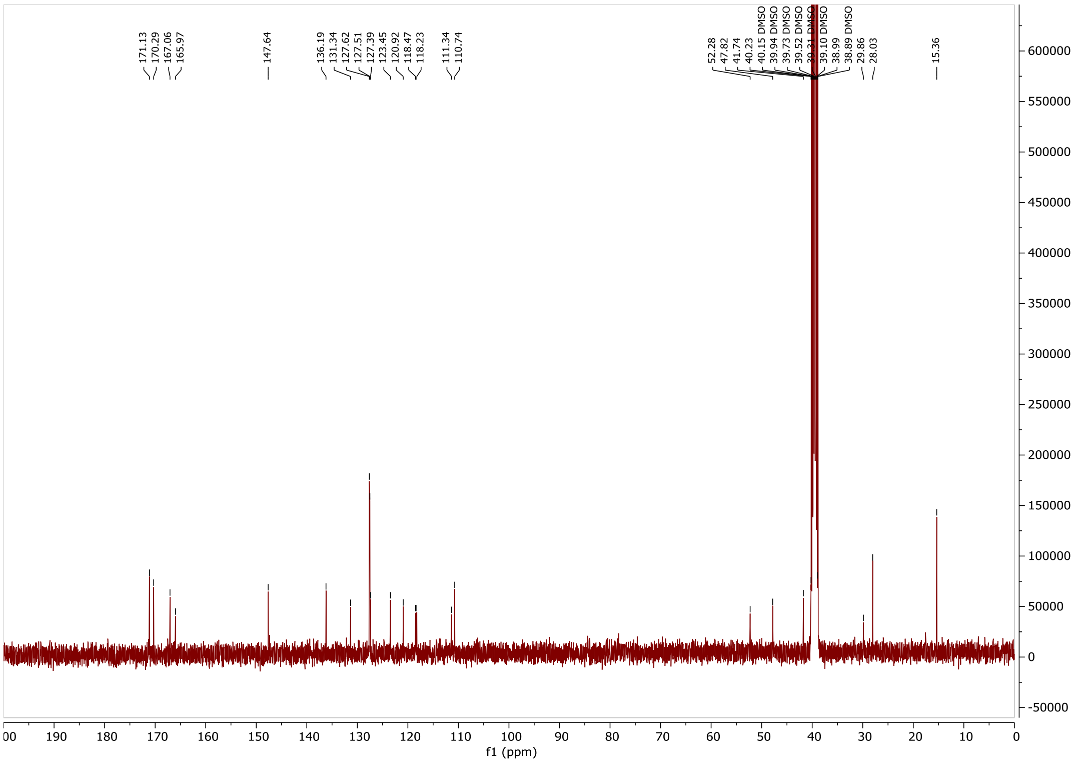

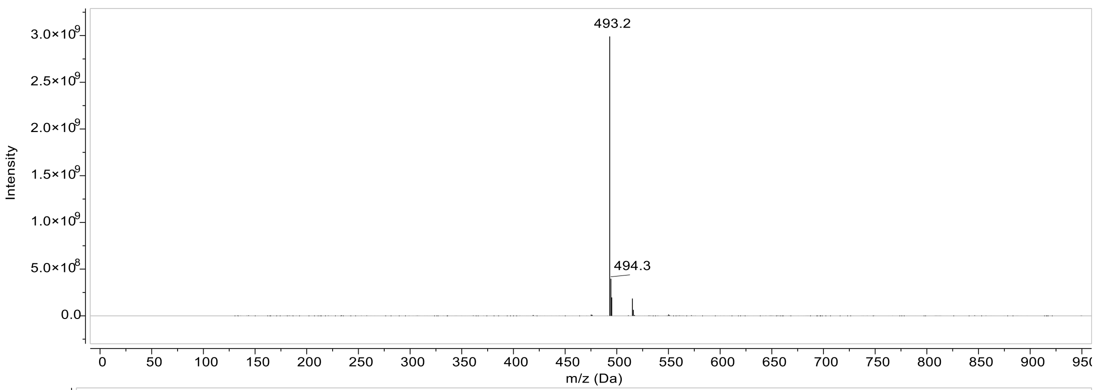

493.2
494.3
Intensity
3.0×10^9
2.5×10^9
2.0×10^9
1.5×10^9
1.0×10^9
5.0×10^8
0.0
0 50 100 150 200 250 300 350 400 450 500 550 600 650 700 750 800 850 900 950
m/z (Da)

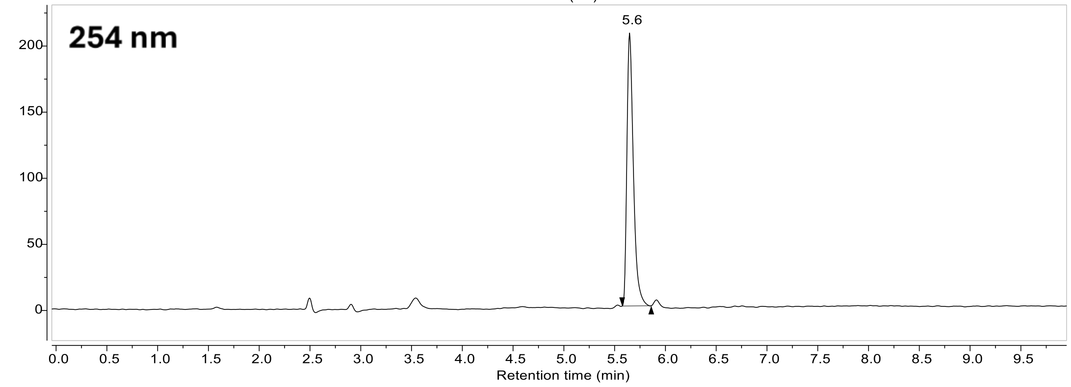

254 nm
5.6
200
150
100
50
0
0.0 0.5 1.0 1.5 2.0 2.5 3.0 3.5 4.0 4.5 5.0 5.5 6.0 6.5 7.0 7.5 8.0 8.5 9.0 9.5
Retention time (min)

#### Compound 25:

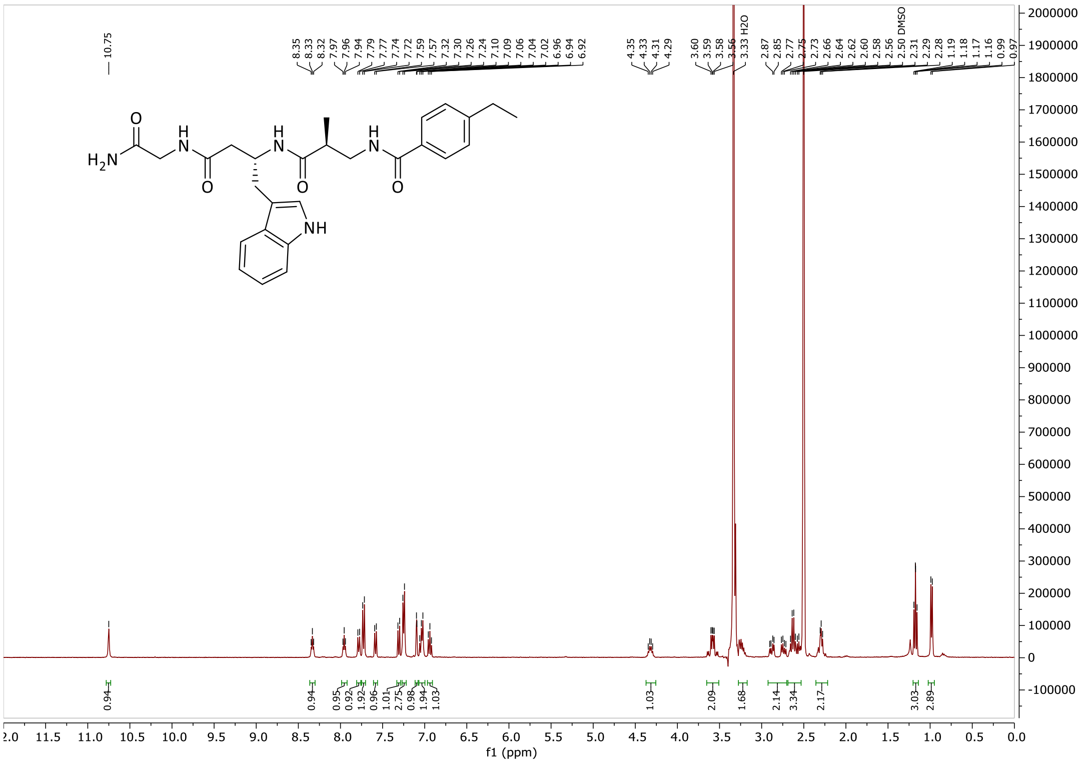

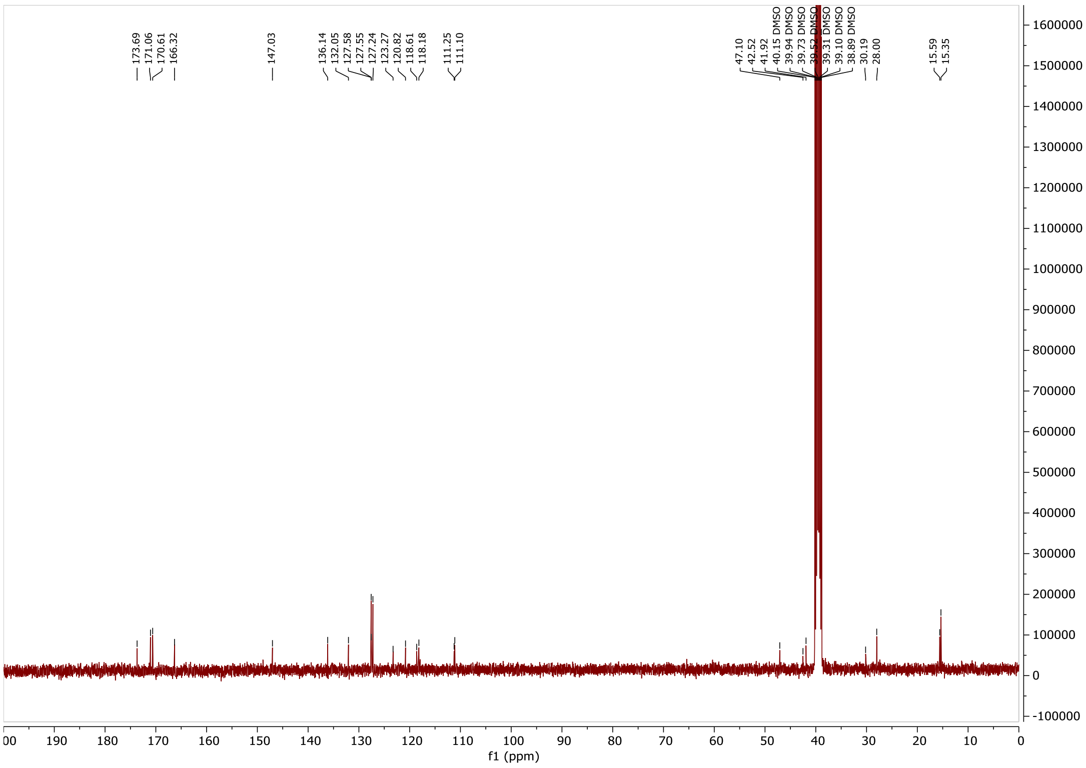

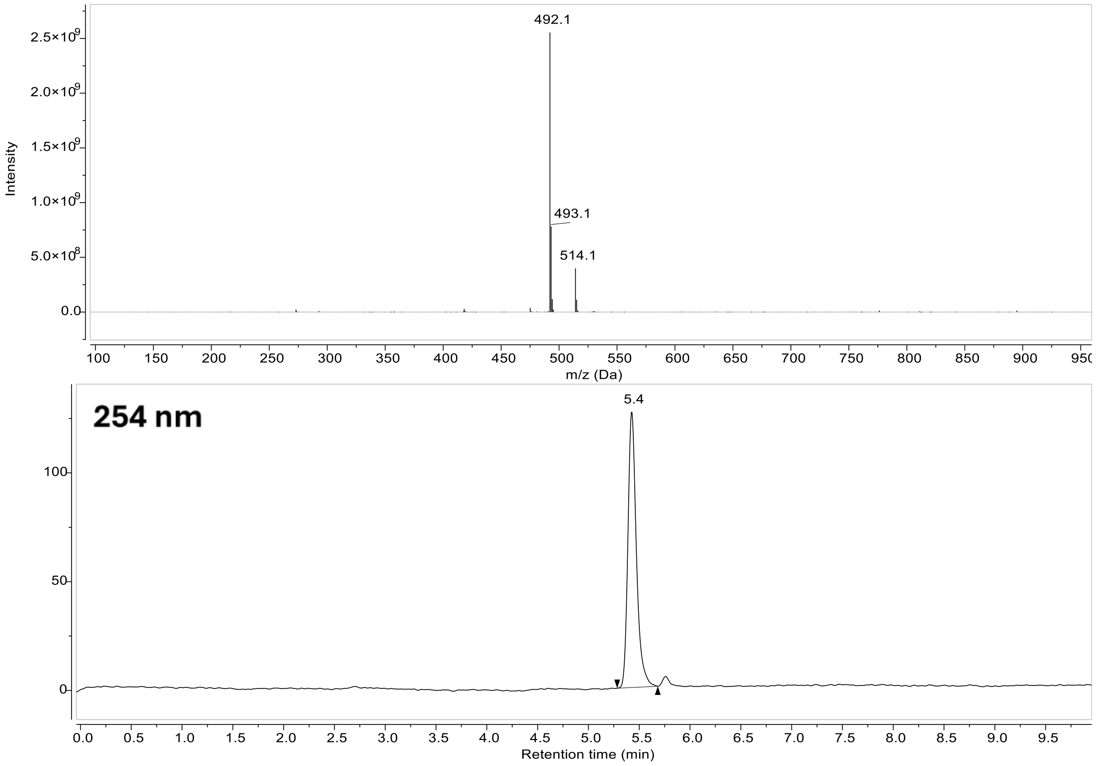
492.1
493.1
514.1
Intensity
m/z (Da)
254 nm
5.4
Retention time (min)

### Compound 26:

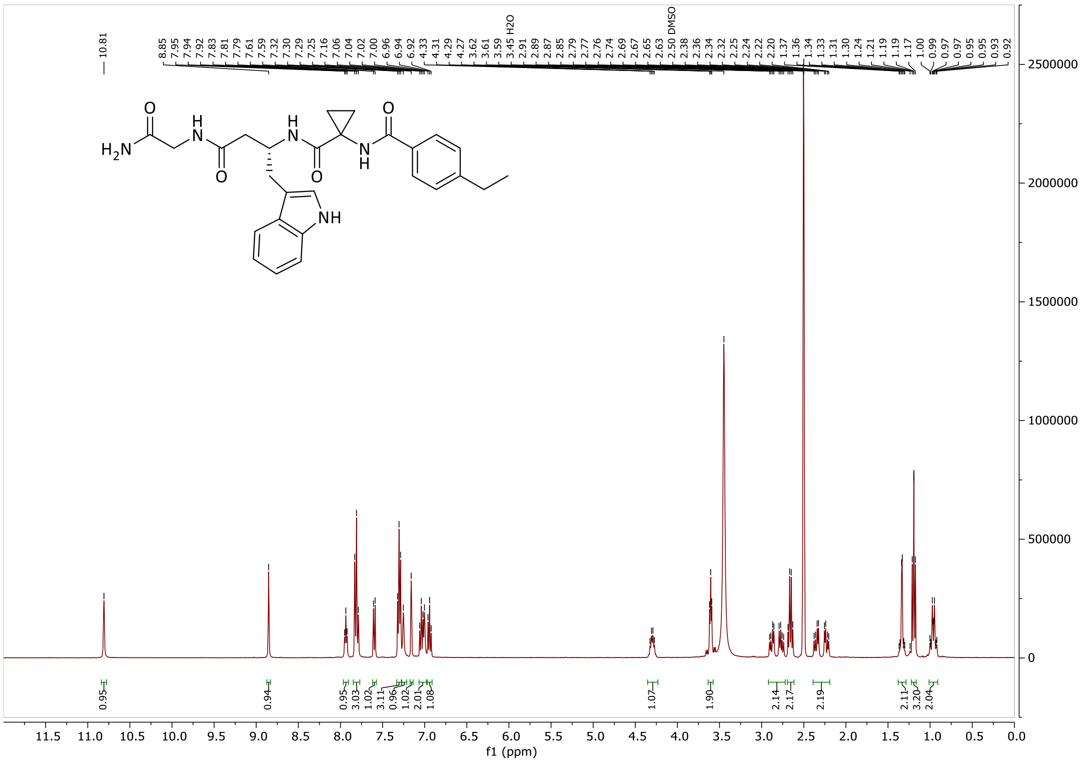

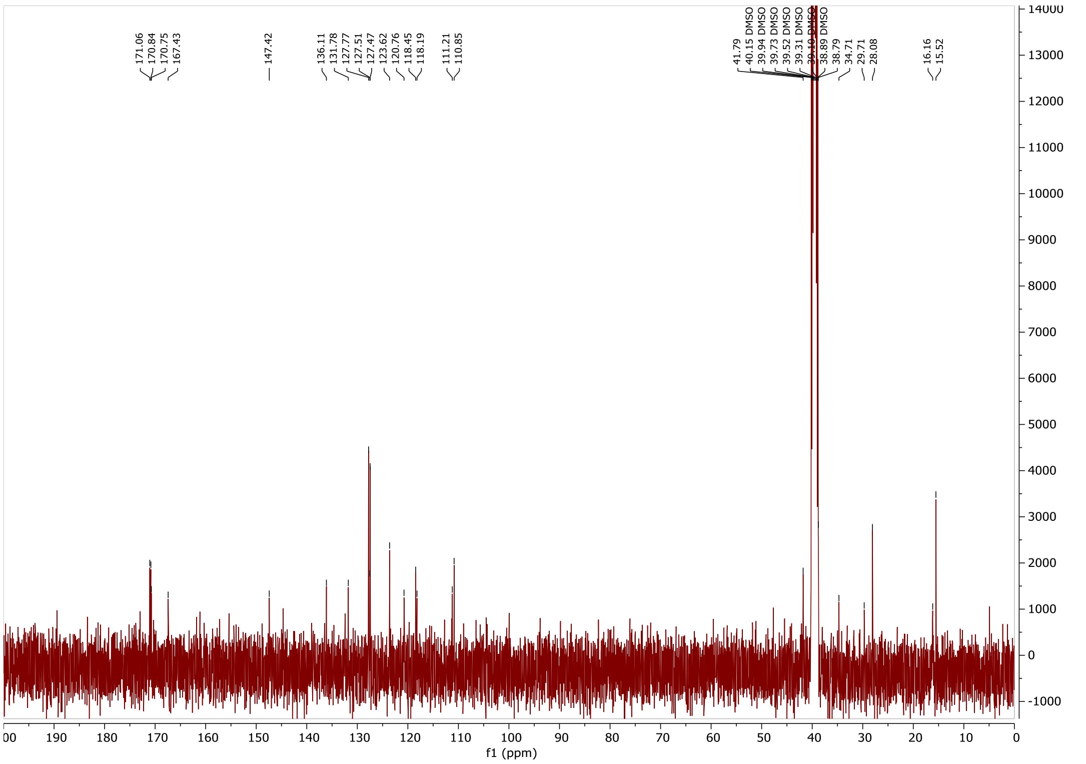

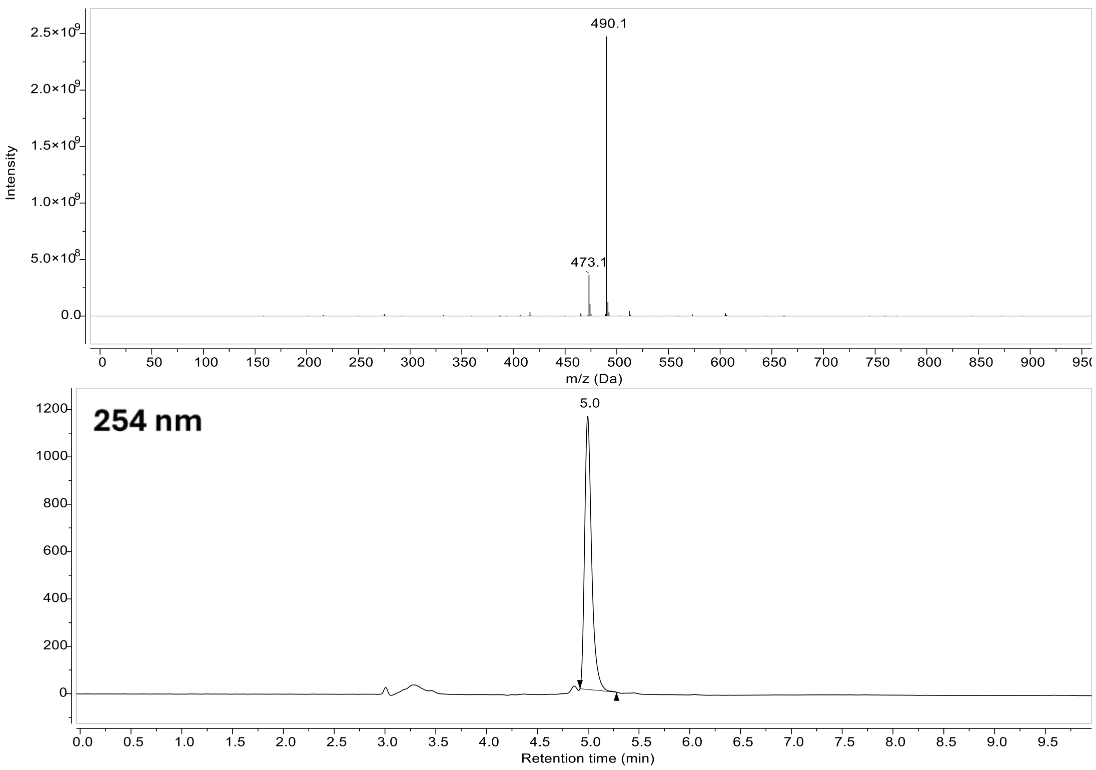
490.1
473.1
Intensity
m/z (Da)
254 nm
5.0
Retention time (min)

**Compound 27:**

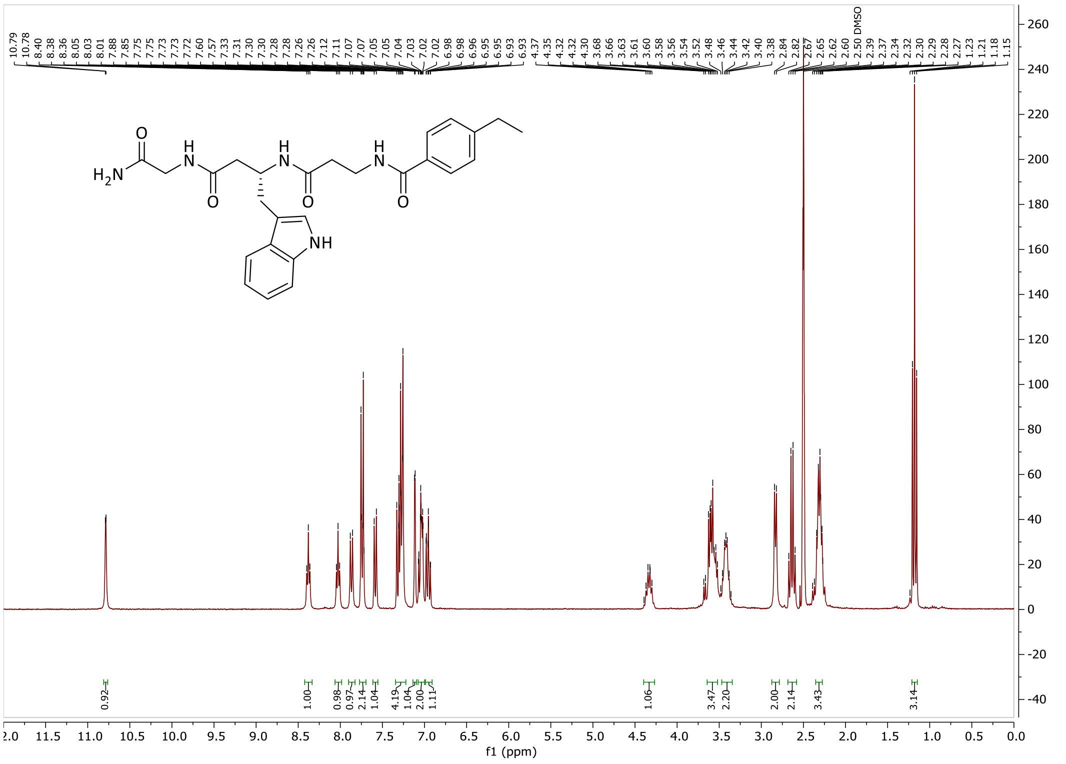

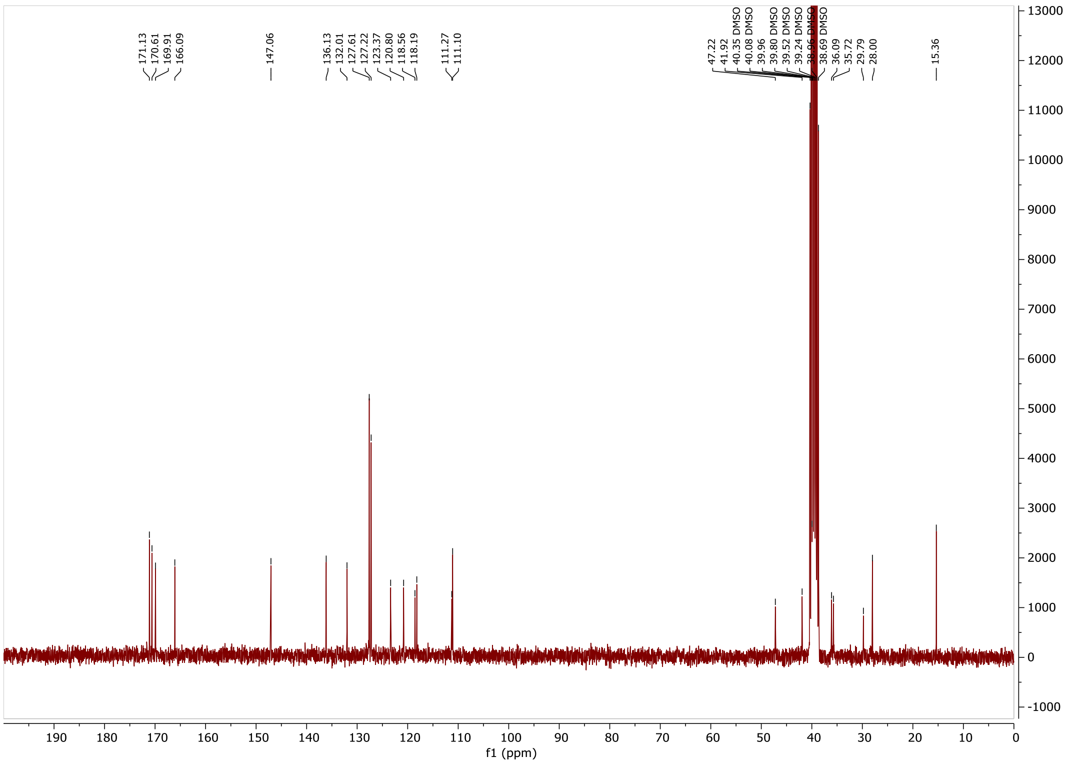

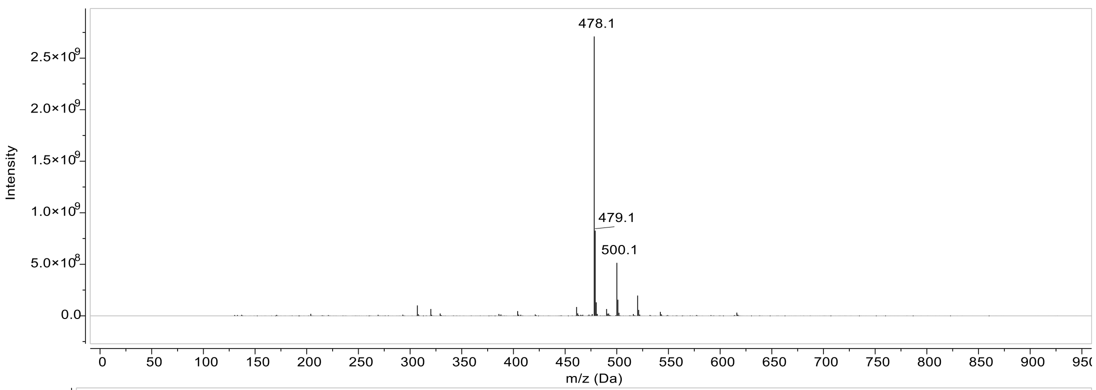
478.1
479.1
500.1
Intensity
2.5×10^9
2.0×10^9
1.5×10^9
1.0×10^9
5.0×10^8
0.0
0 50 100 150 200 250 300 350 400 450 500 550 600 650 700 750 800 850 900 950
m/z (Da)

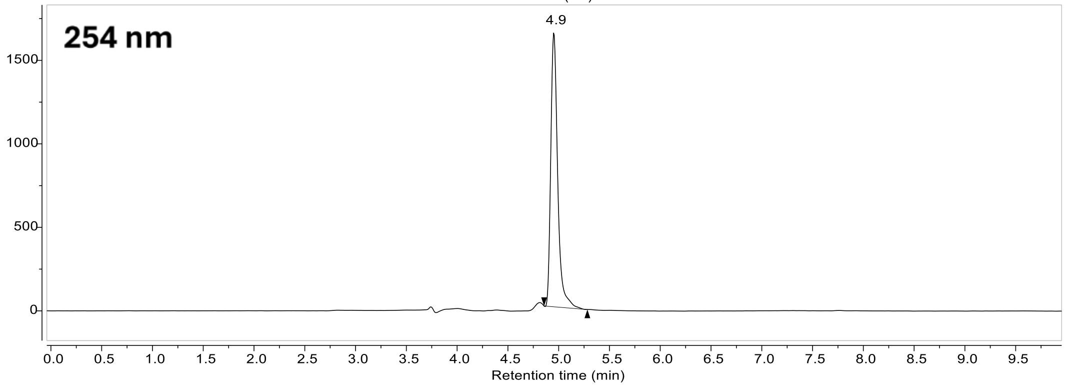
254 nm
4.9
1500
1000
500
0
0.0 0.5 1.0 1.5 2.0 2.5 3.0 3.5 4.0 4.5 5.0 5.5 6.0 6.5 7.0 7.5 8.0 8.5 9.0 9.5
Retention time (min)

**Compound 28:**

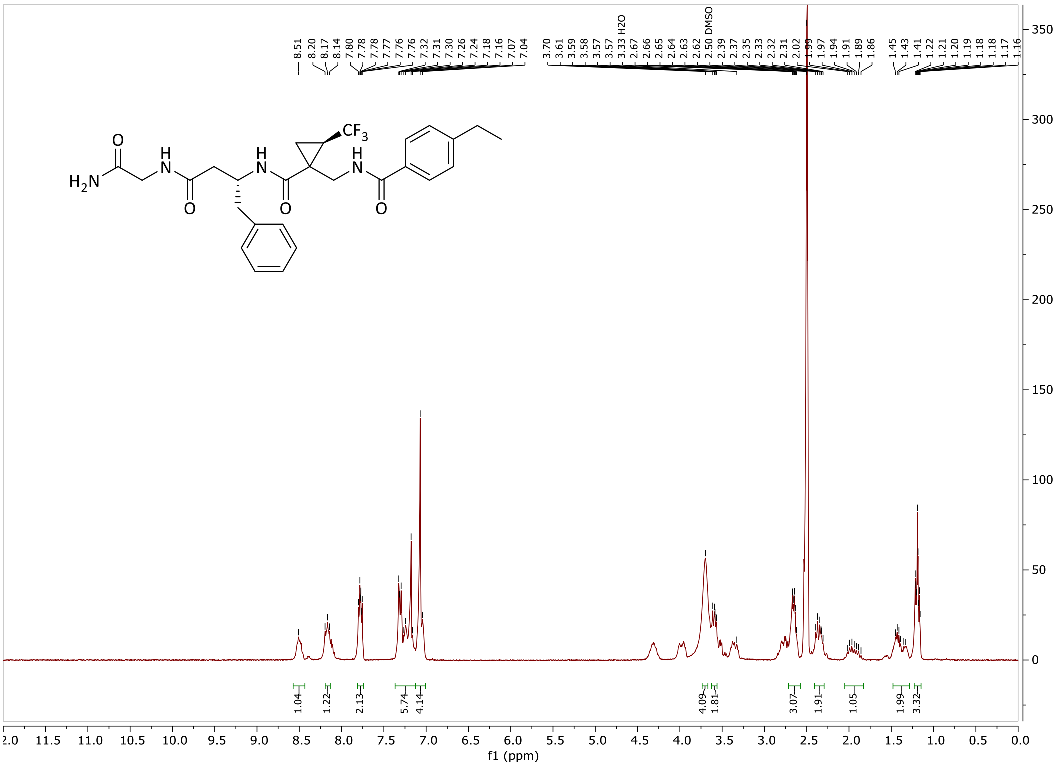

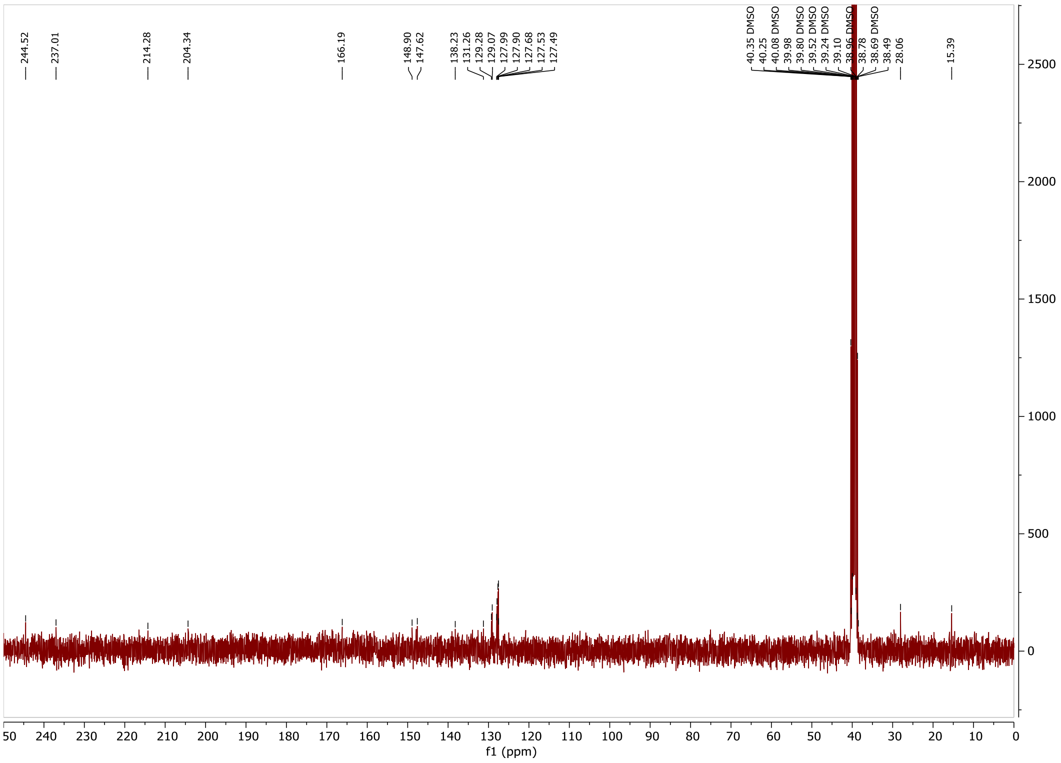

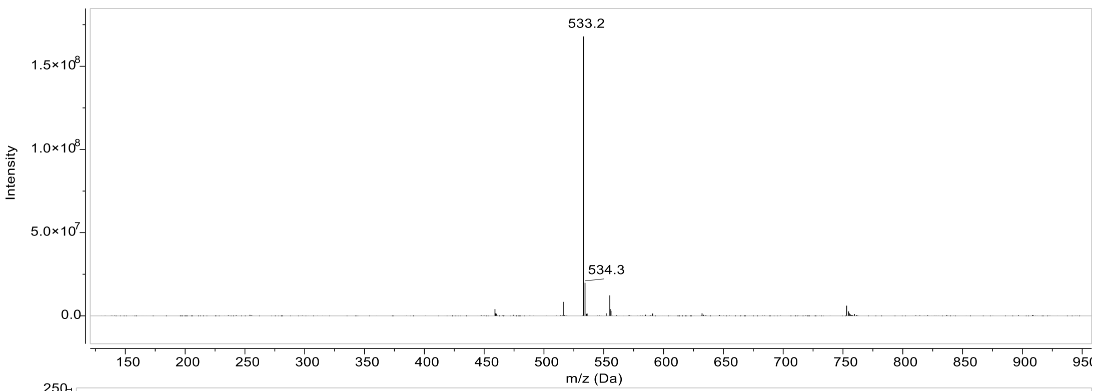

533.2
534.3
Intensity
m/z (Da)

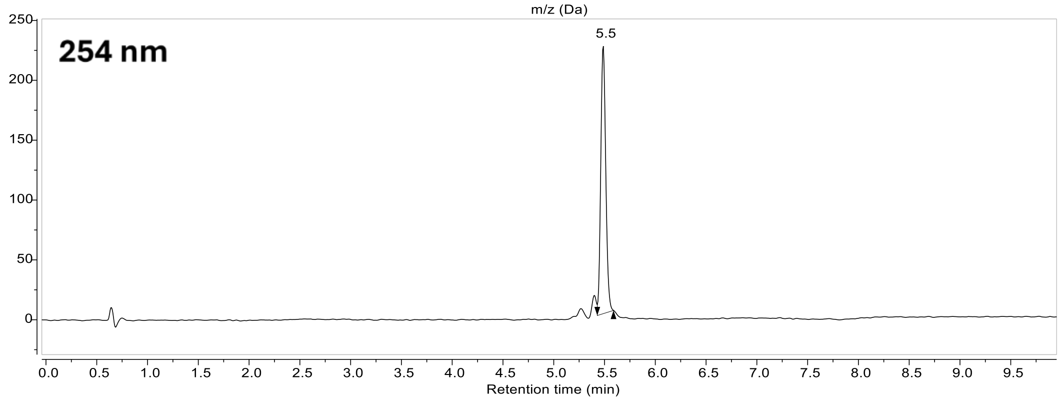

254 nm
5.5
Retention time (min)

#### Compound 29:

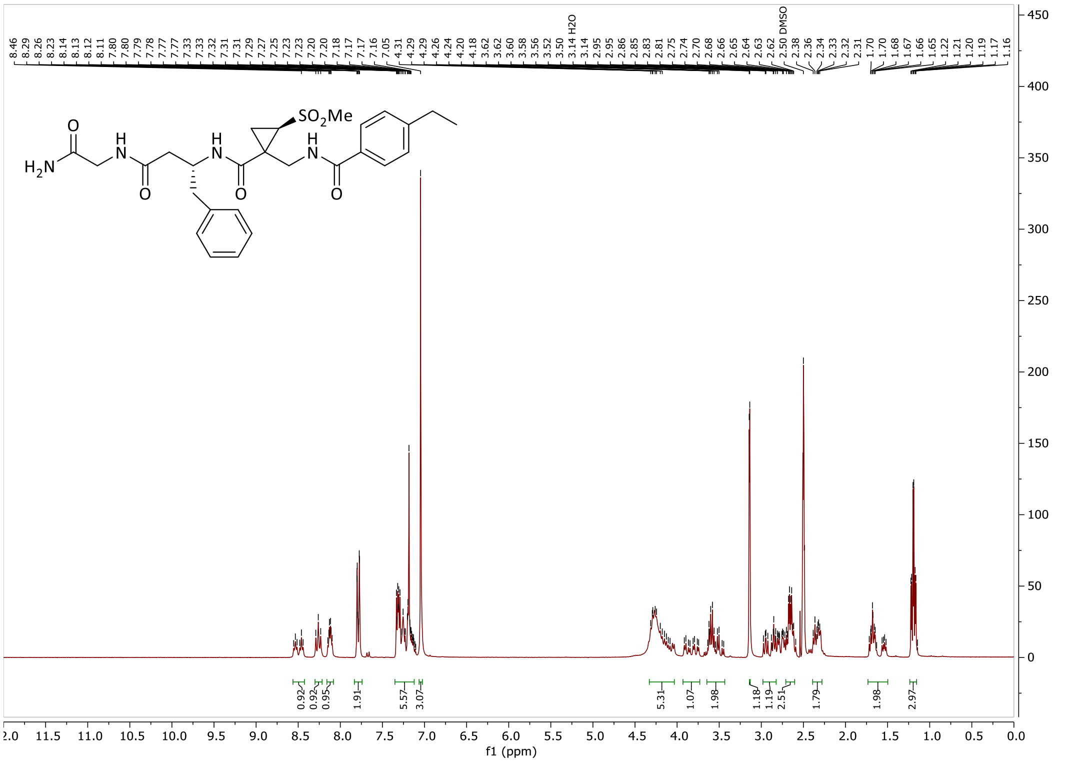

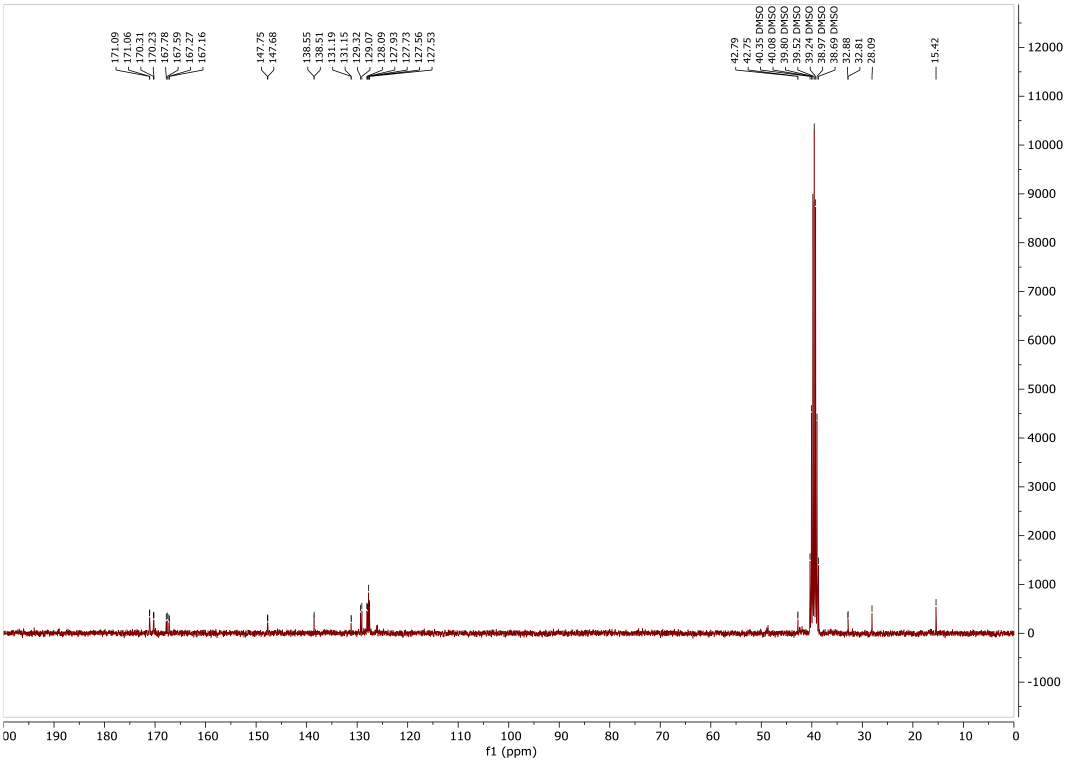

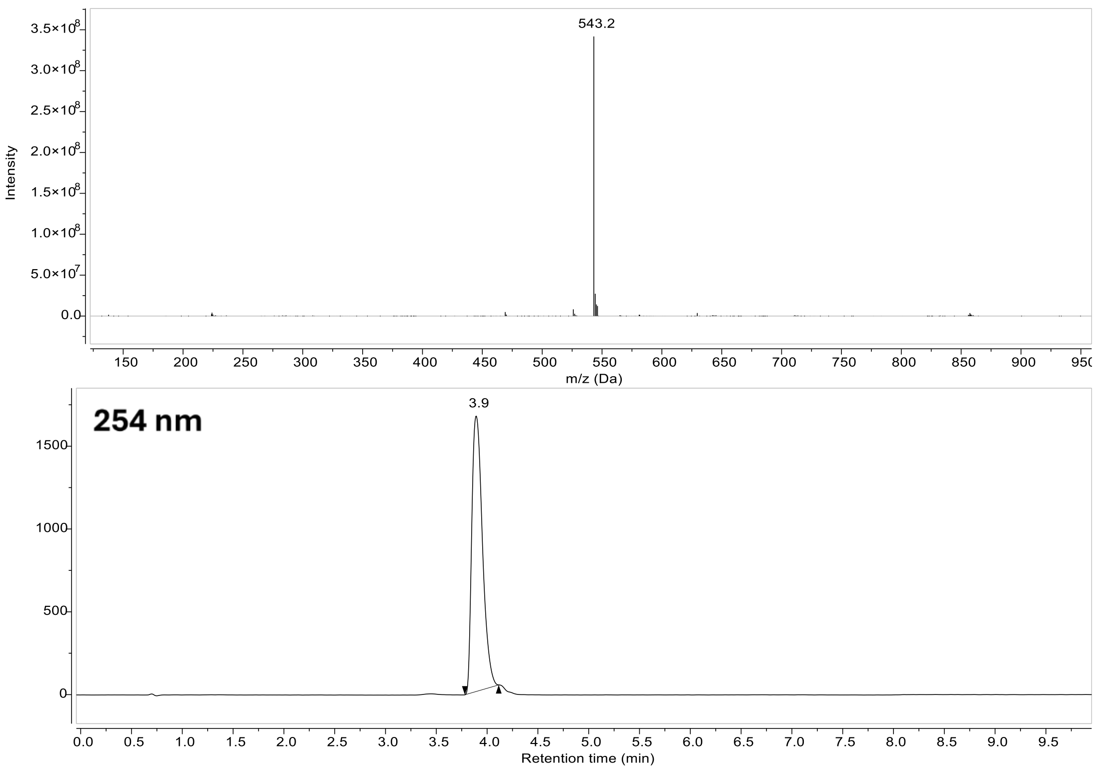

543.2
Intensity
m/z (Da)
254 nm
3.9
Retention time (min)

**Compound 30:**

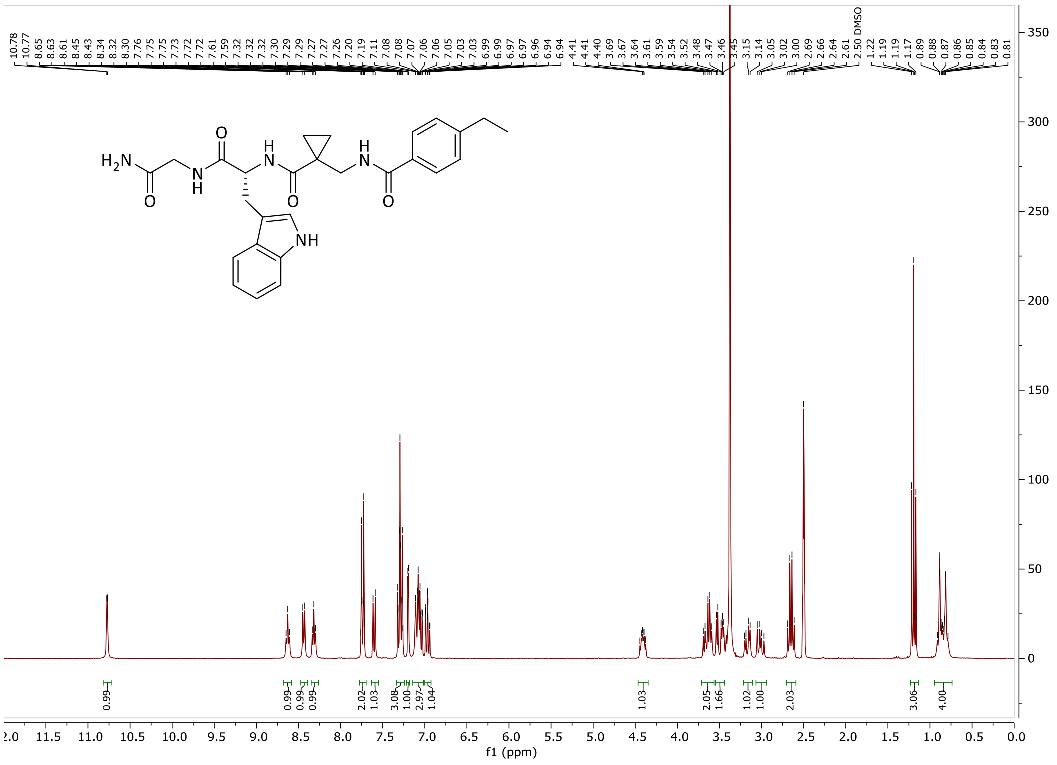

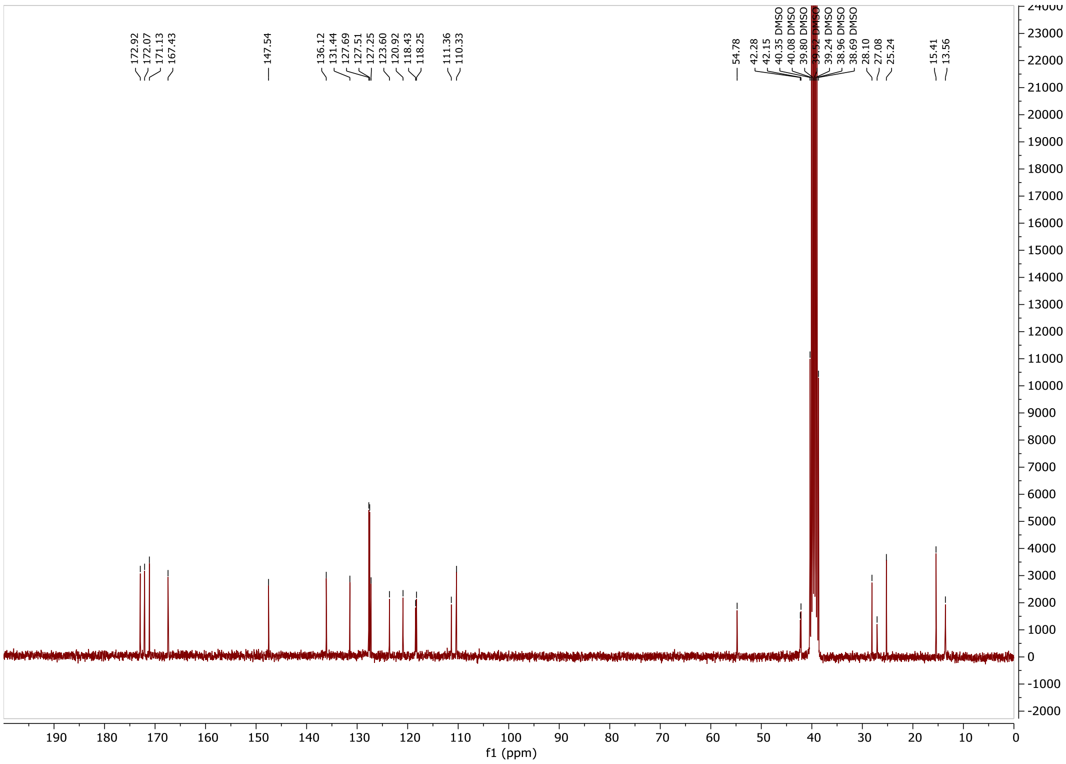

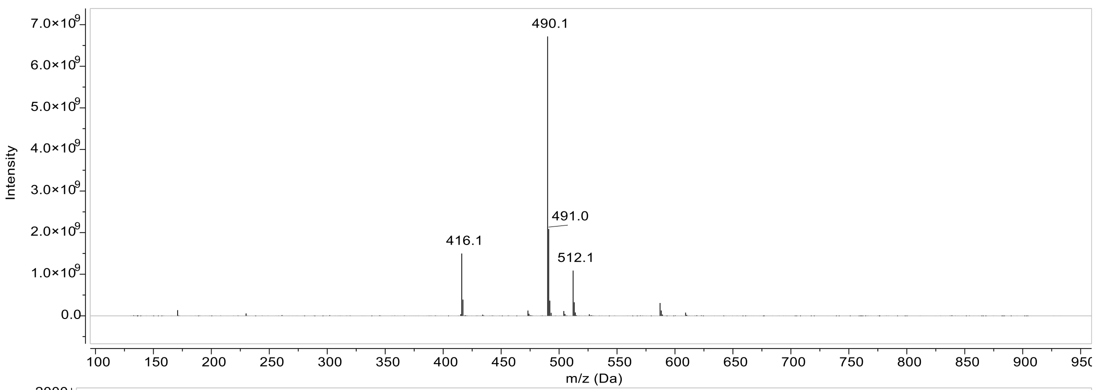

7.0×10^9
6.0×10^9
5.0×10^9
4.0×10^9
3.0×10^9
2.0×10^9
1.0×10^9
0.0
Intensity
490.1
491.0
416.1
512.1
100 150 200 250 300 350 400 450 500 550 600 650 700 750 800 850 900 950
m/z (Da)

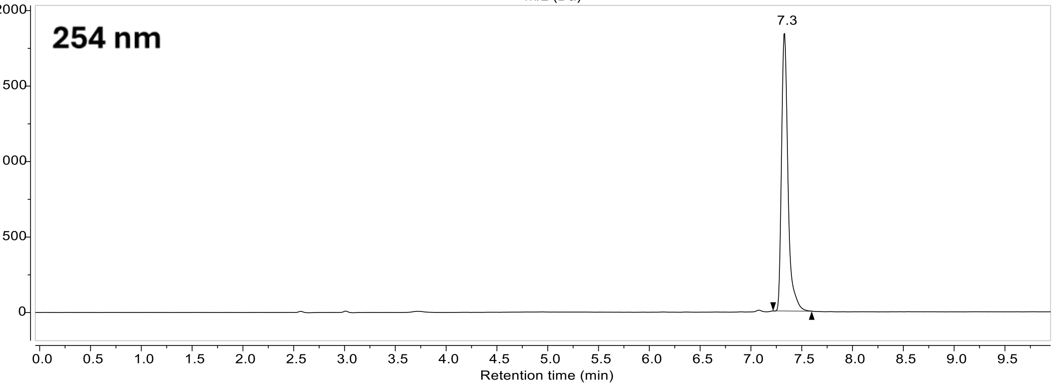

254 nm
2000
1500
1000
500
0
7.3
0.0 0.5 1.0 1.5 2.0 2.5 3.0 3.5 4.0 4.5 5.0 5.5 6.0 6.5 7.0 7.5 8.0 8.5 9.0 9.5
Retention time (min)

### Compound 31:

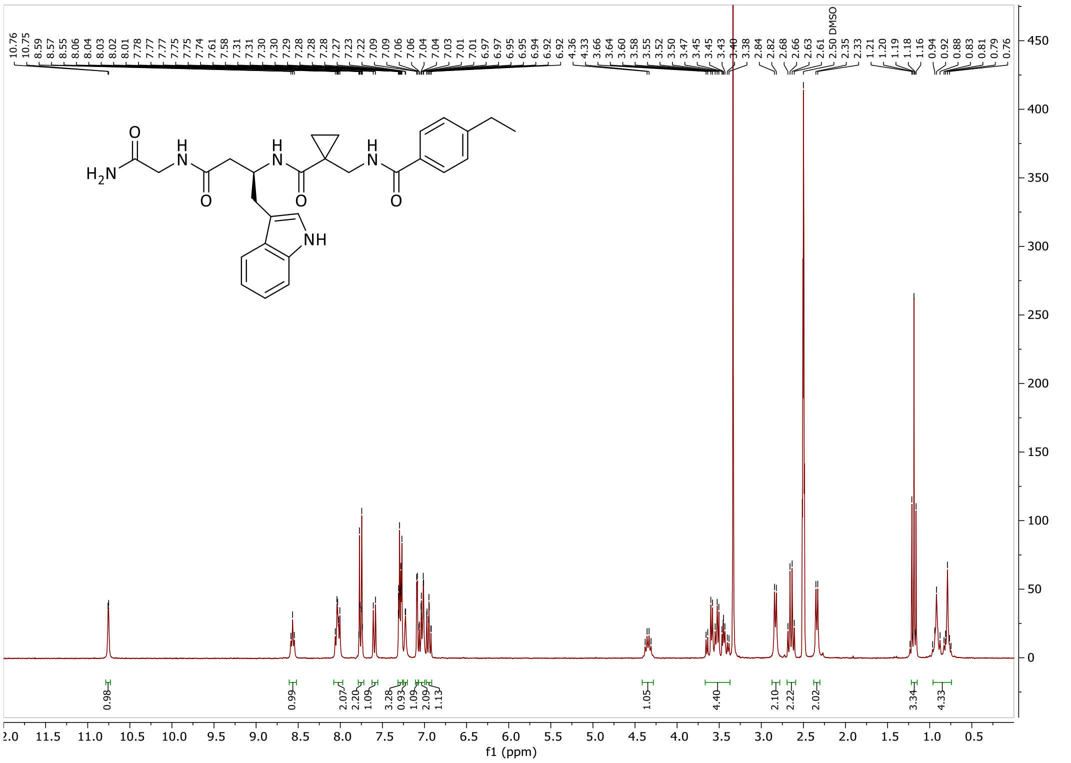

10.76 10.75 8.59 8.57 8.55 8.06 8.04 8.03 8.02 8.01 7.78 7.77 7.77 7.75 7.75 7.74 7.61 7.58 7.31 7.30 7.30 7.29 7.28 7.28 7.28 7.27 7.23 7.22 7.09 7.09 7.06 7.06 7.04 7.04 7.03 7.01 6.97 6.97 6.95 6.95 6.94 6.92 4.36 4.33 3.66 3.64 3.60 3.58 3.55 3.52 3.50 3.47 3.45 3.45 3.43 3.40 3.38 2.84 2.82 2.68 2.66 2.63 2.61 2.50 DMSO 2.35 2.33 1.21 1.20 1.19 1.18 1.16 0.94 0.92 0.88 0.83 0.81 0.79 0.76
H2N
NH
0.98 0.99 2.07 2.20 1.09 3.28 0.93 1.09 2.09 1.13 1.05 4.40 2.10 2.22 2.02 3.34 4.33
f1 (ppm)

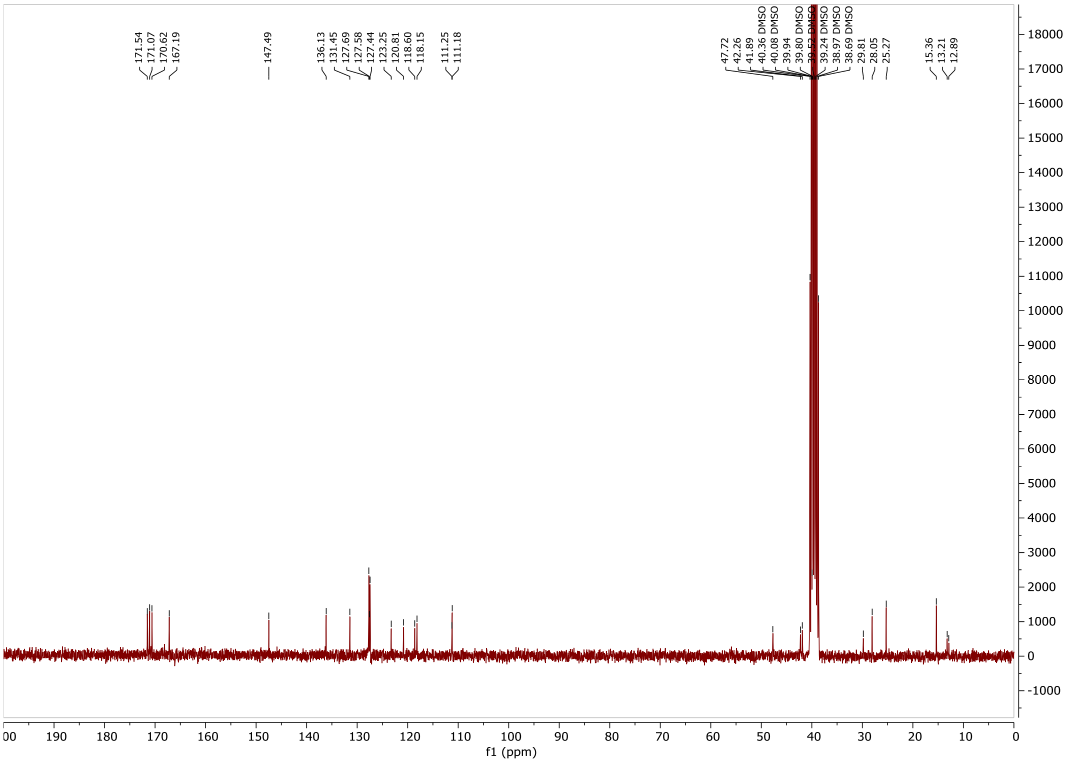

171.54 171.07 170.62 167.19 147.49 136.13 131.45 127.69 127.58 127.44 123.25 120.81 118.60 118.15 111.25 111.18 47.72 42.26 41.89 40.36 DMSO 40.08 DMSO 39.94 39.80 DMSO 39.52 DMSO 39.24 DMSO 38.97 DMSO 38.69 DMSO 29.81 28.05 25.27 15.36 13.21 12.89
f1 (ppm)

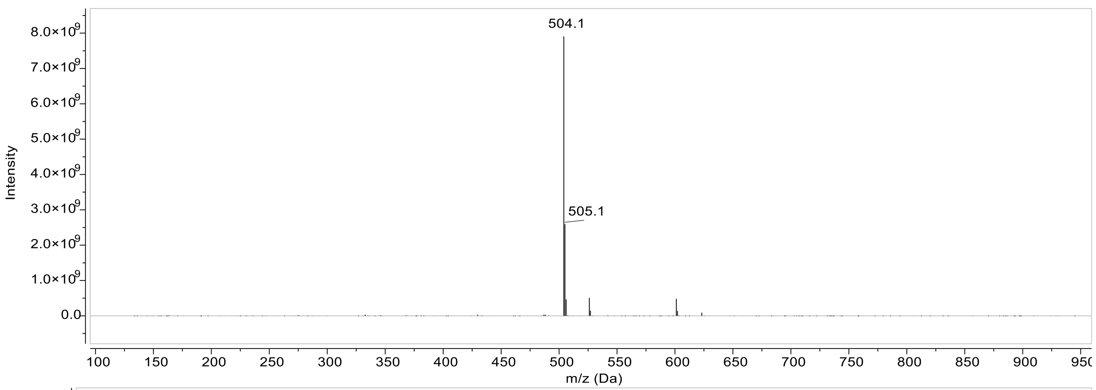

504.1
505.1
Intensity
m/z (Da)

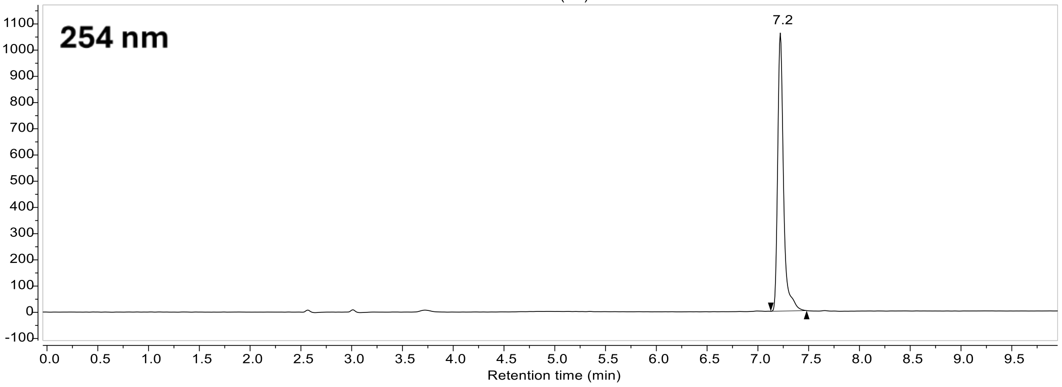

254 nm
7.2
Retention time (min)

#### Compound 32:

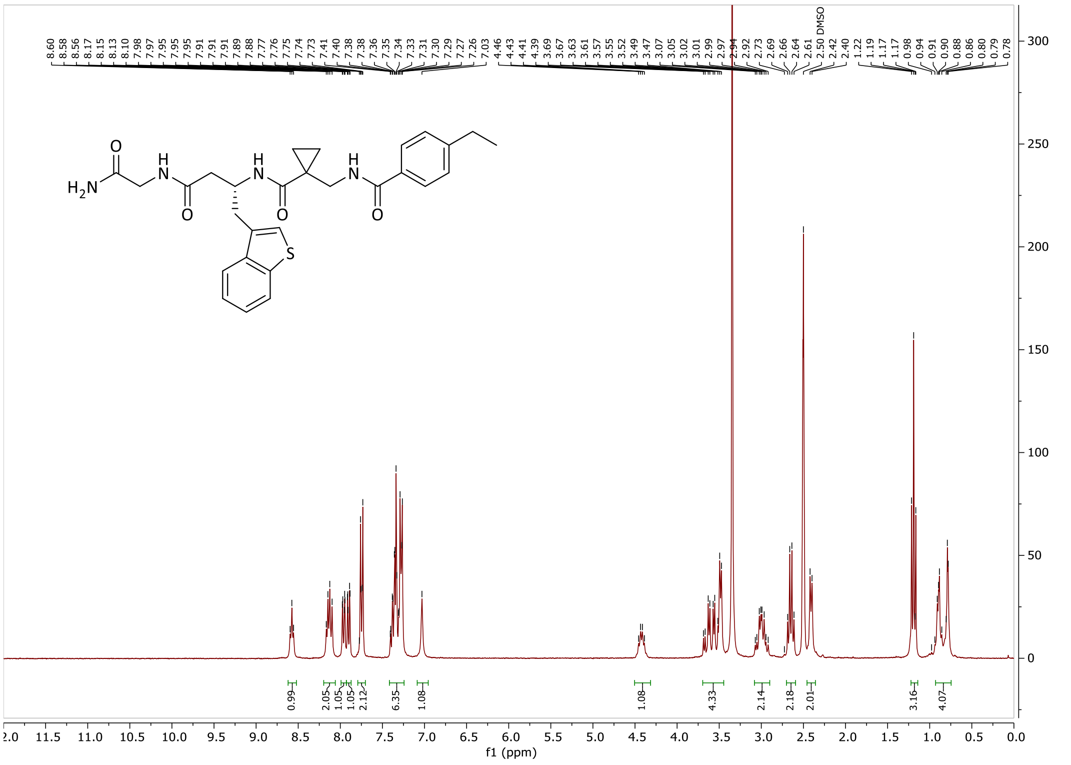

8.60 8.58 8.56 8.17 8.15 8.13 8.10 7.98 7.97 7.95 7.95 7.95 7.91 7.91 7.91 7.89 7.88 7.77 7.76 7.75 7.74 7.73 7.41 7.40 7.38 7.38 7.36 7.35 7.34 7.33 7.31 7.30 7.29 7.27 7.26 7.03
4.46 4.43 4.41 4.39 3.69 3.67 3.63 3.61 3.57 3.55 3.52 3.49 3.47 3.07 3.05 3.02 3.01 2.99 2.97 2.94 2.92 2.73 2.69 2.66 2.64 2.61 2.50 DMSO 2.42 2.40 1.22 1.19 1.17 1.17 0.98 0.94 0.91 0.90 0.88 0.86 0.80 0.79 0.78
0.99 2.05 1.05 1.05 2.12 6.35 1.08 1.08 4.33 2.14 2.18 2.01 3.16 4.07
f1 (ppm)
H2N O H N O H N O H N O S

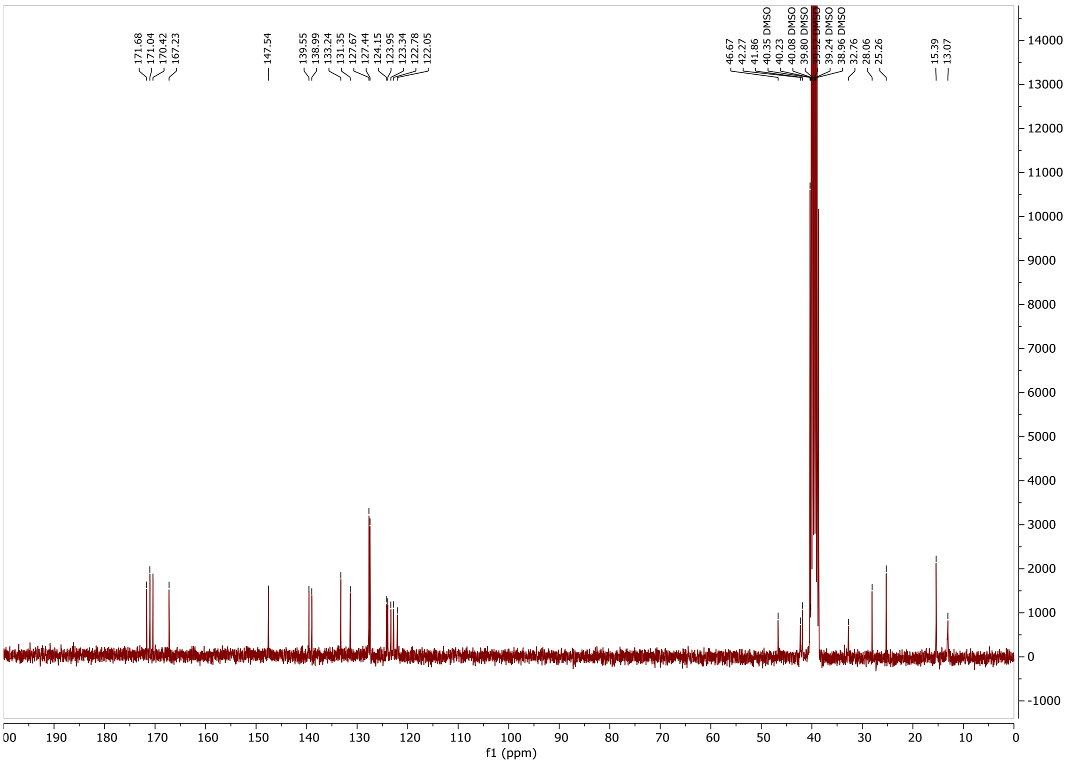

171.68 171.04 170.42 167.23 147.54 139.55 138.99 133.24 131.35 127.67 127.44 124.15 123.95 123.34 122.78 122.05
46.67 42.27 41.86 40.35 DMSO 40.23 40.08 DMSO 39.80 DMSO 39.52 DMSO 39.24 DMSO 38.96 DMSO 32.76 28.06 25.26 15.39 13.07
f1 (ppm)

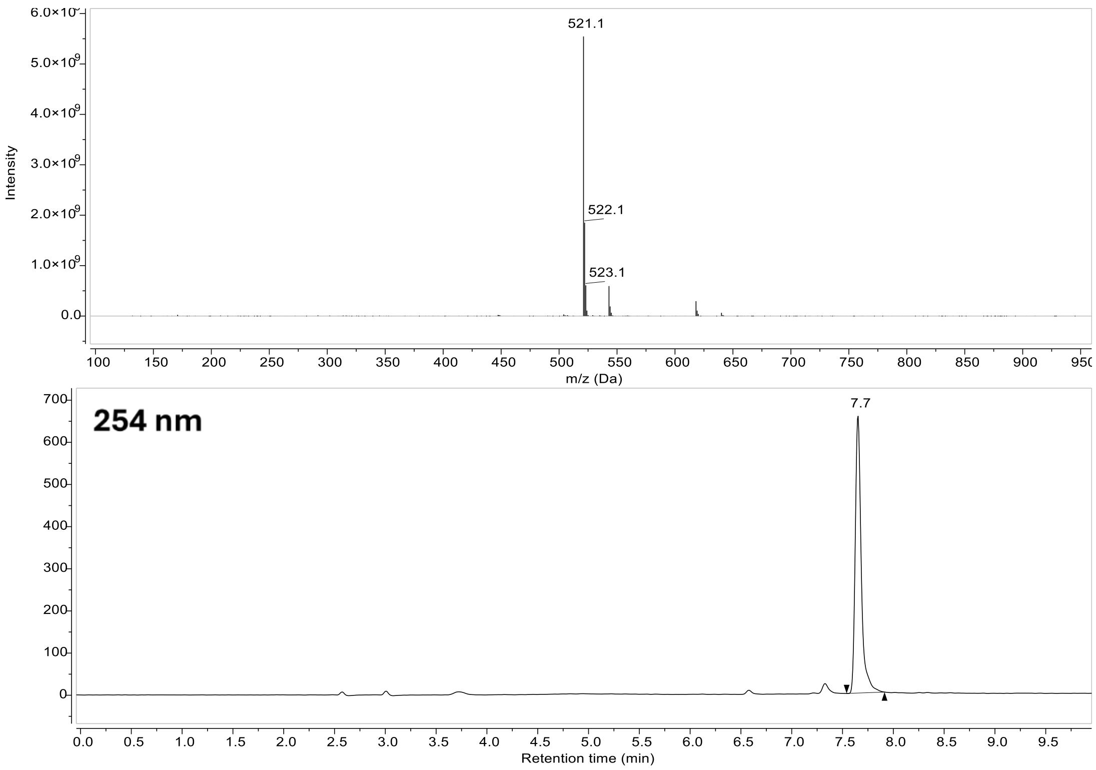

521.1
522.1
523.1
Intensity
m/z (Da)
254 nm
7.7
Retention time (min)

#### Compound 14:

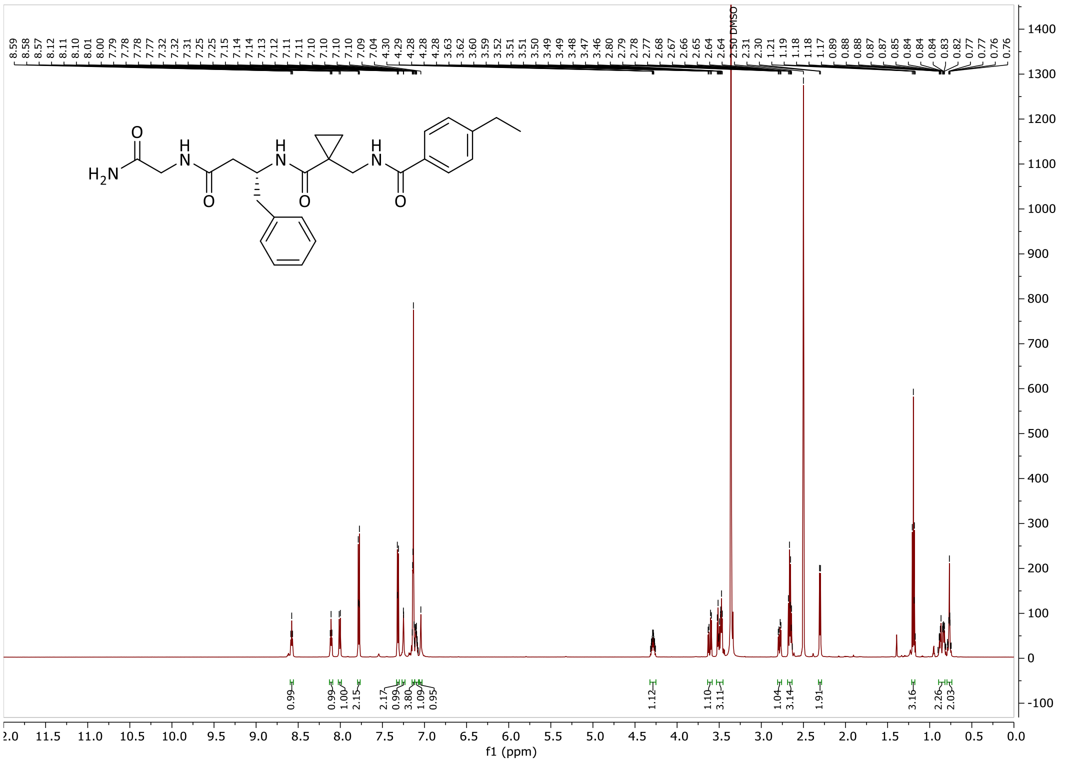

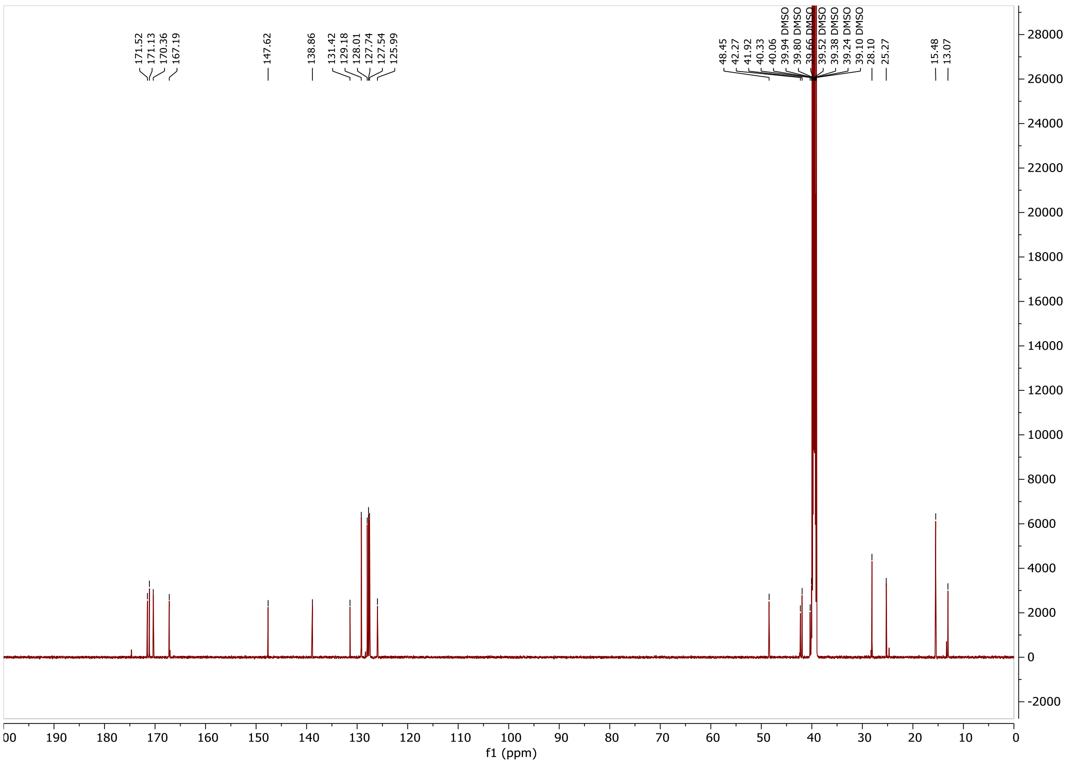

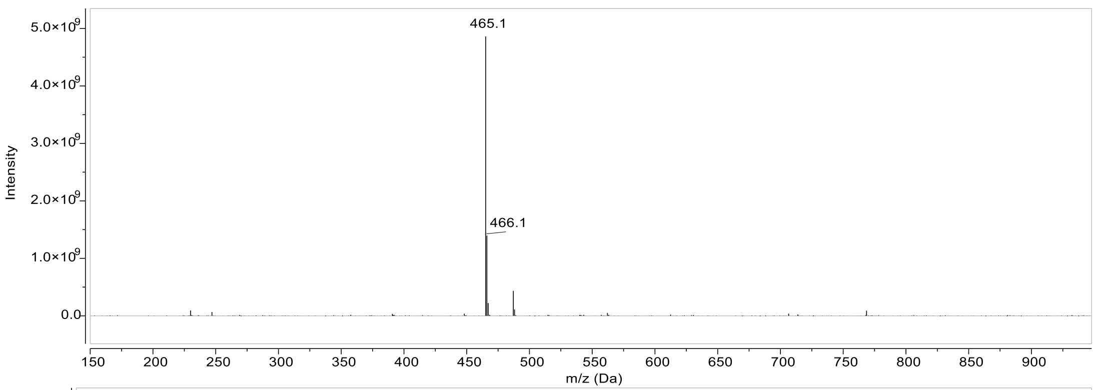

465.1
466.1
Intensity
m/z (Da)

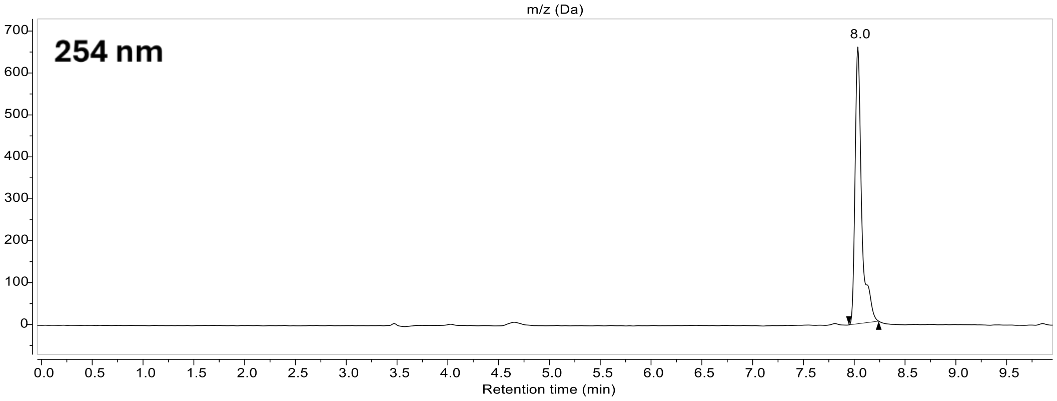

254 nm
8.0
Retention time (min)

#### Compound 15:

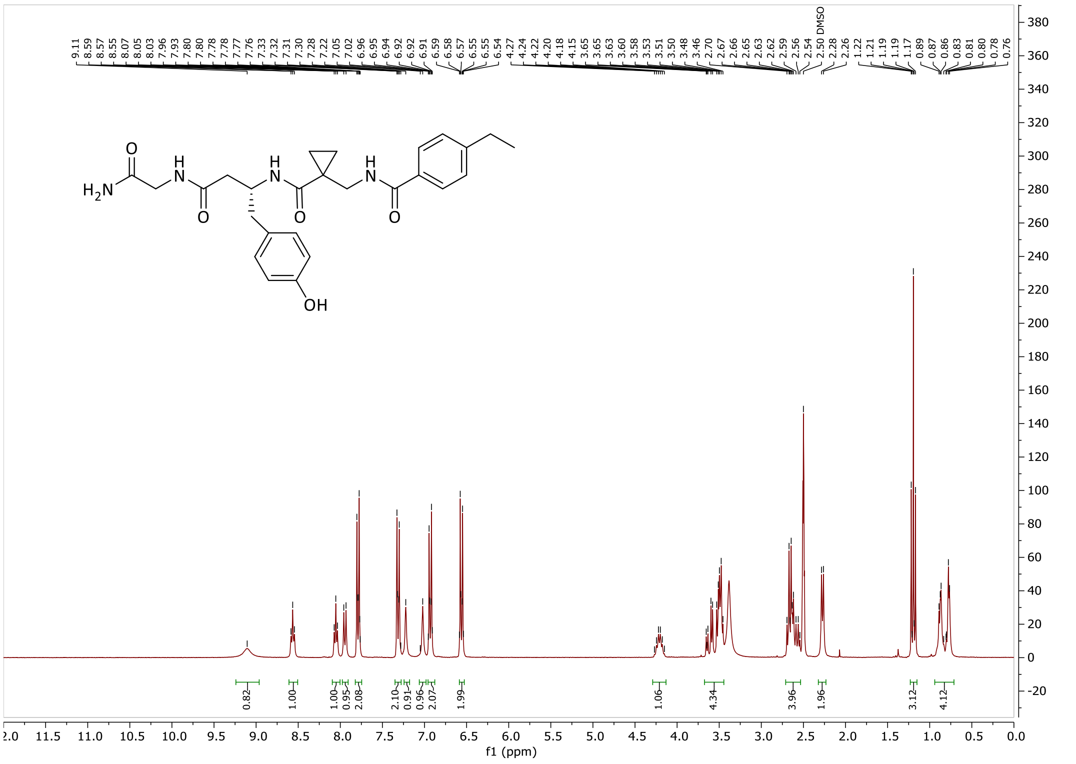

9.11 8.59 8.57 8.55 8.07 8.05 8.03 7.96 7.93 7.80 7.80 7.78 7.78 7.77 7.76 7.33 7.32 7.31 7.30 7.28 7.22 7.05 7.02 6.96 6.95 6.94 6.92 6.92 6.91 6.59 6.58 6.57 6.55 6.55 6.54 4.27 4.24 4.22 4.20 4.18 4.15 3.65 3.65 3.63 3.60 3.58 3.53 3.51 3.50 3.48 3.46 2.70 2.67 2.66 2.65 2.63 2.62 2.59 2.56 2.54 2.50 DMSO 2.28 2.26 1.22 1.21 1.19 1.19 1.17 0.89 0.87 0.86 0.83 0.81 0.80 0.78 0.76
H$_2$N
O
H
N
O
H
N
O
H
N
O
OH
0.82 1.00 1.00 0.95 2.08 2.10 0.91 0.96 2.07 1.99 1.06 4.34 3.96 1.96 3.12 4.12
f1 (ppm)

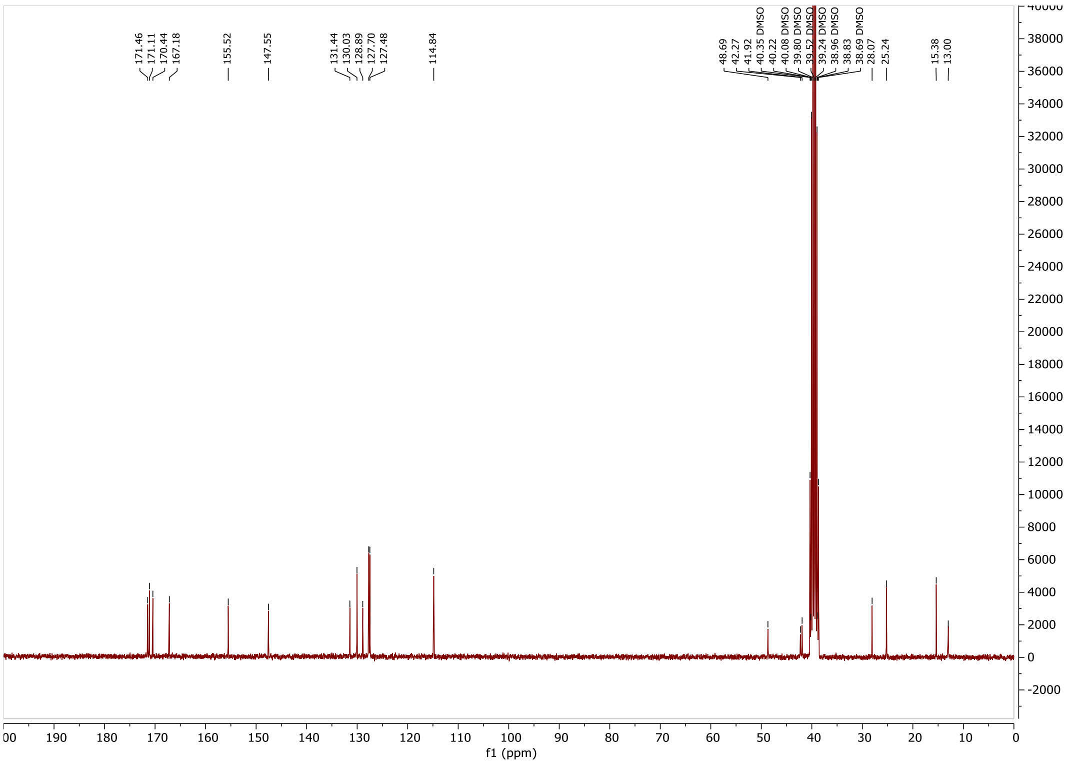

171.46 171.11 170.44 167.18 155.52 147.55 131.44 130.03 128.89 127.70 127.48 114.84 48.69 42.27 41.92 40.35 DMSO 40.22 40.08 DMSO 39.80 DMSO 39.52 DMSO 39.24 DMSO 38.96 DMSO 38.83 38.69 DMSO 28.07 25.24 15.38 13.00
f1 (ppm)

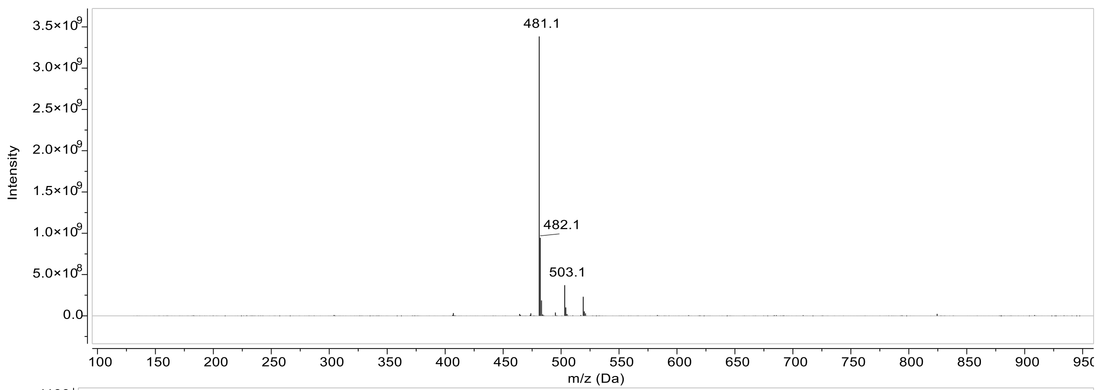
481.1
482.1
503.1
Intensity
m/z (Da)

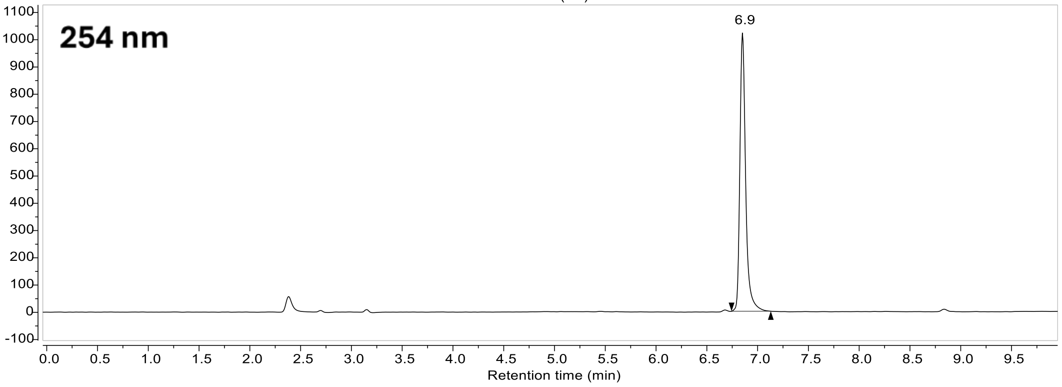
254 nm
6.9
Retention time (min)

#### Compound 35:

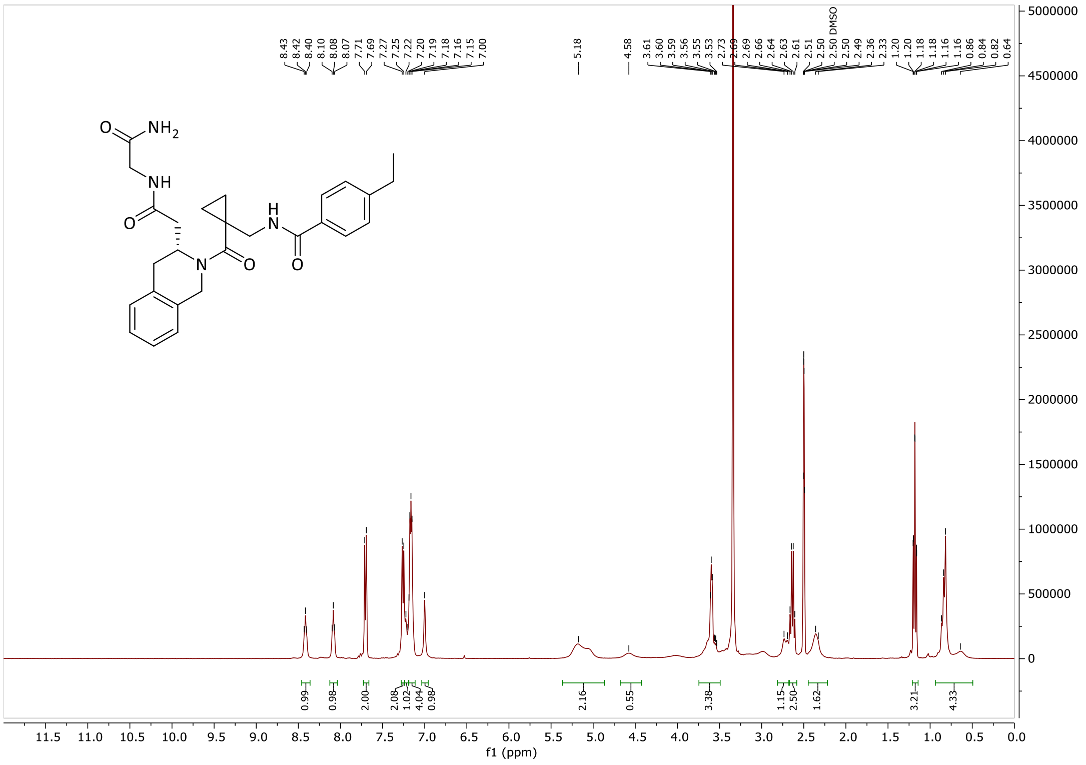

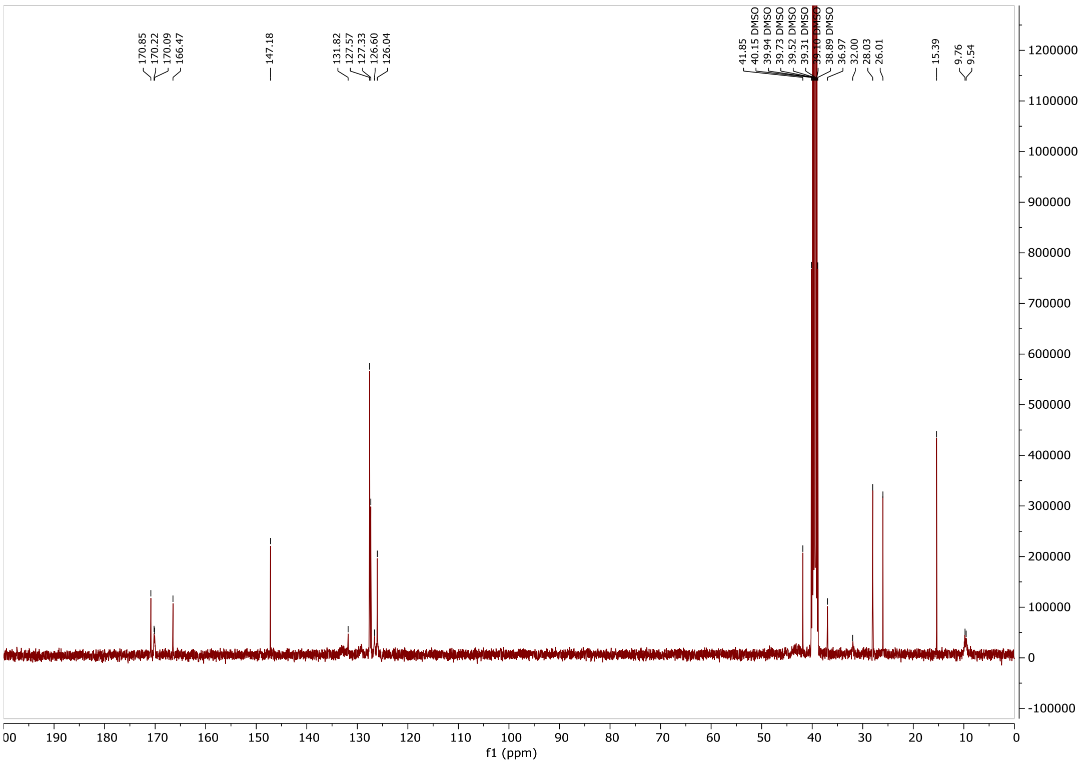

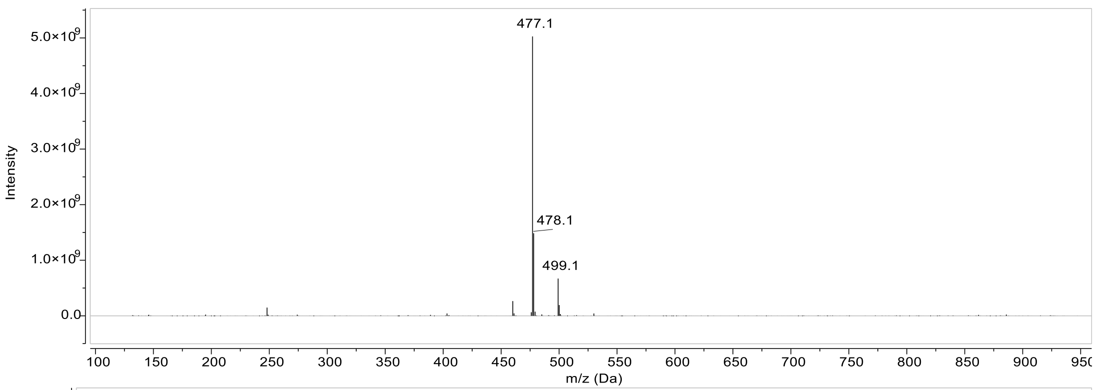
477.1
478.1
499.1
Intensity
m/z (Da)

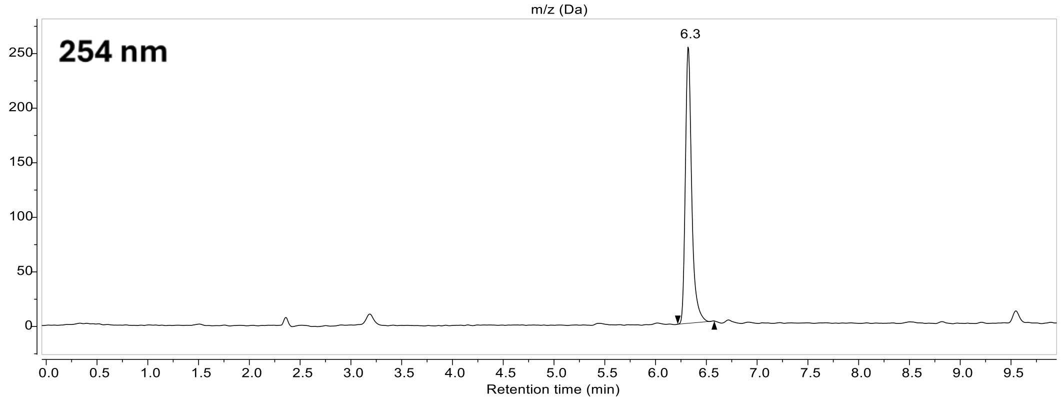
254 nm
6.3
Retention time (min)

**Compound 36:**

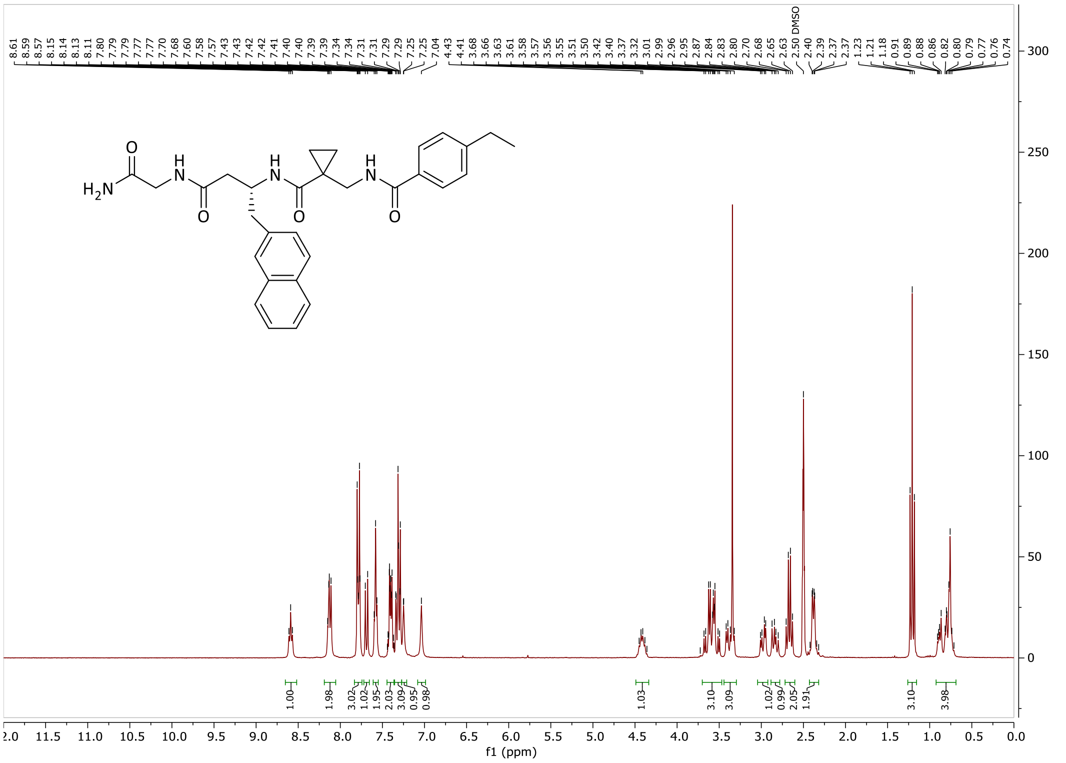

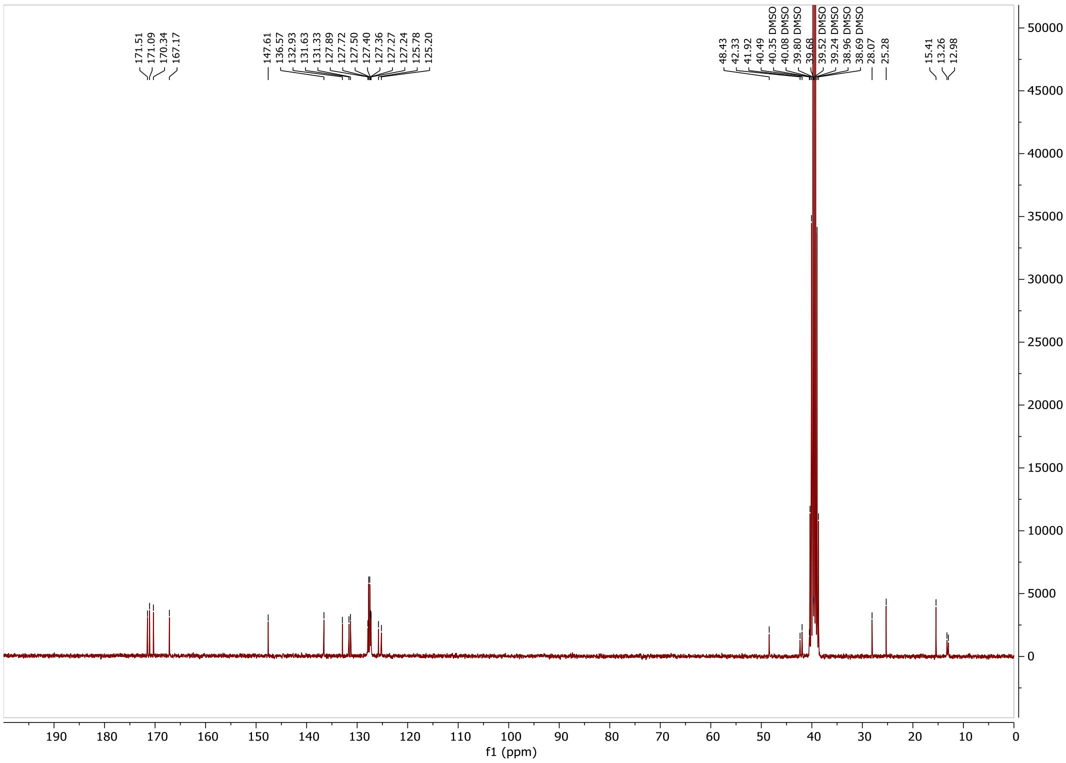

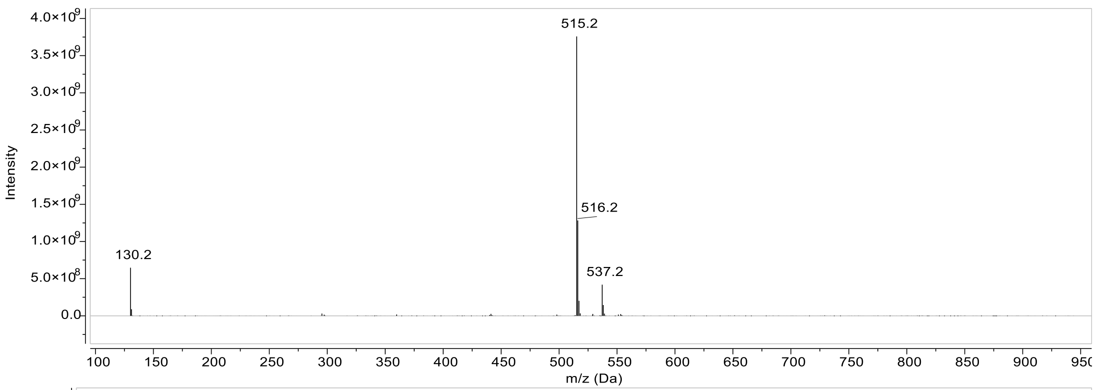

Intensity
4.0×10^9
3.5×10^9
3.0×10^9
2.5×10^9
2.0×10^9
1.5×10^9
1.0×10^9
5.0×10^8
0.0
515.2
516.2
537.2
130.2
m/z (Da)

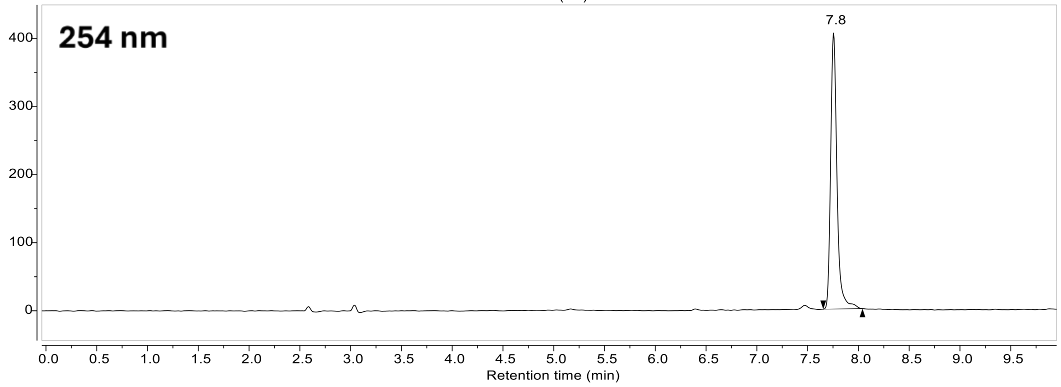

254 nm
7.8
Retention time (min)

### Compound 37:

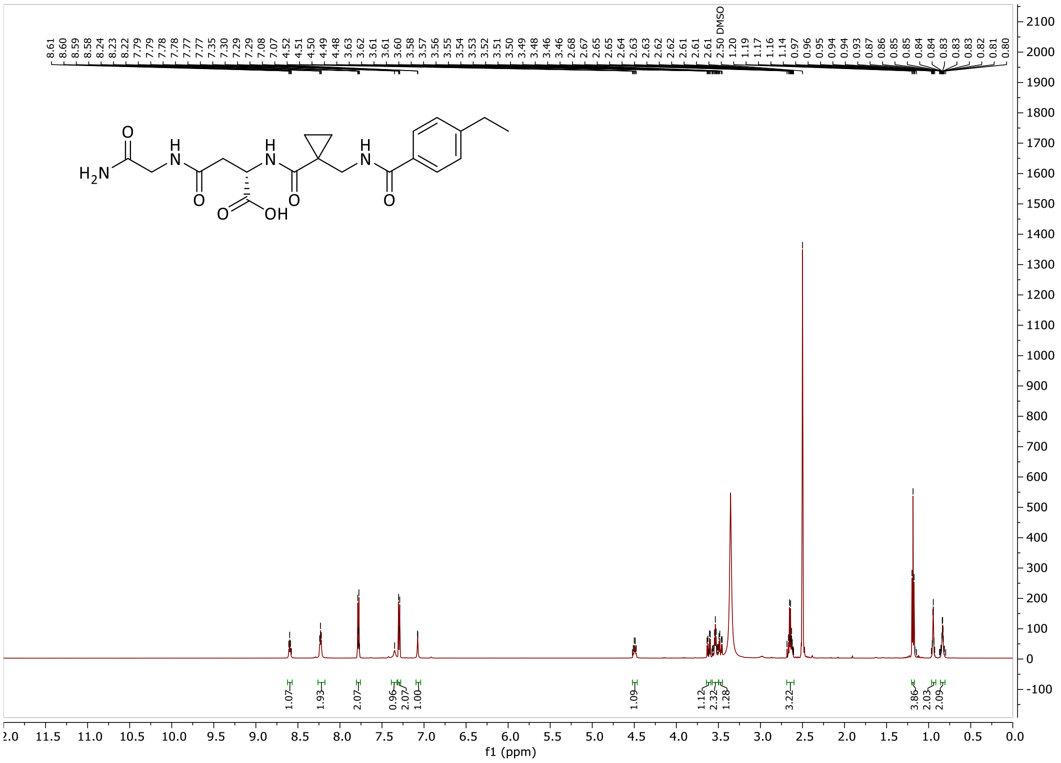

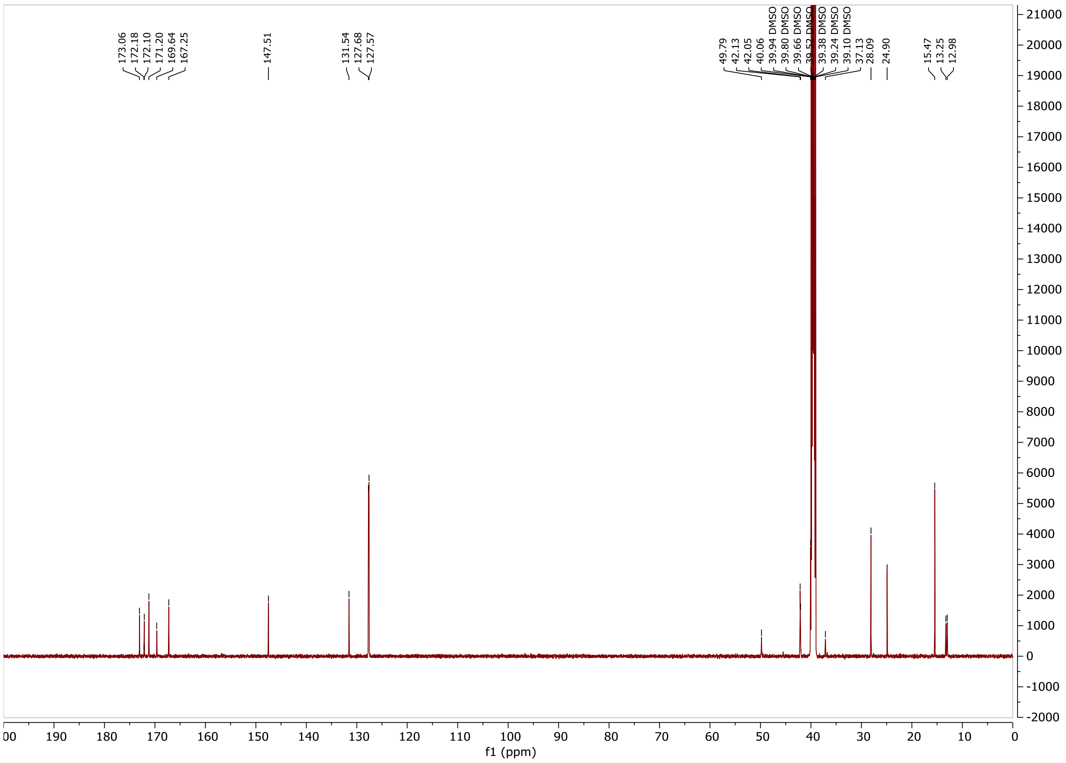

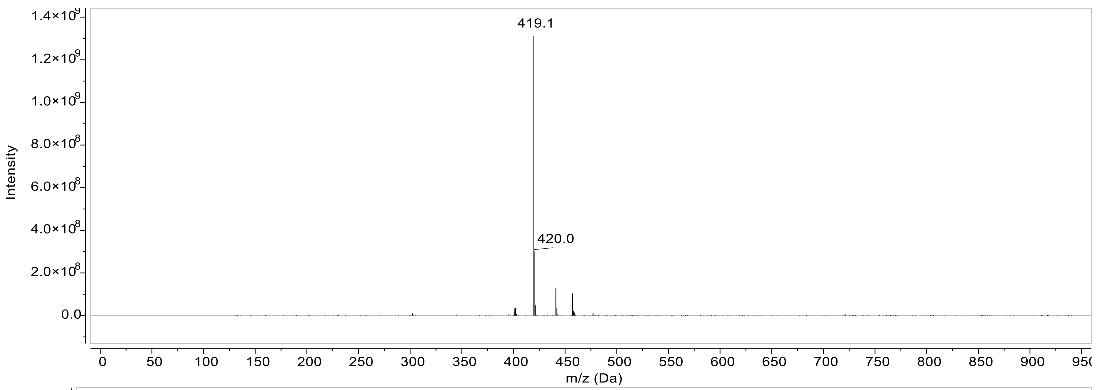
419.1
420.0
Intensity
m/z (Da)

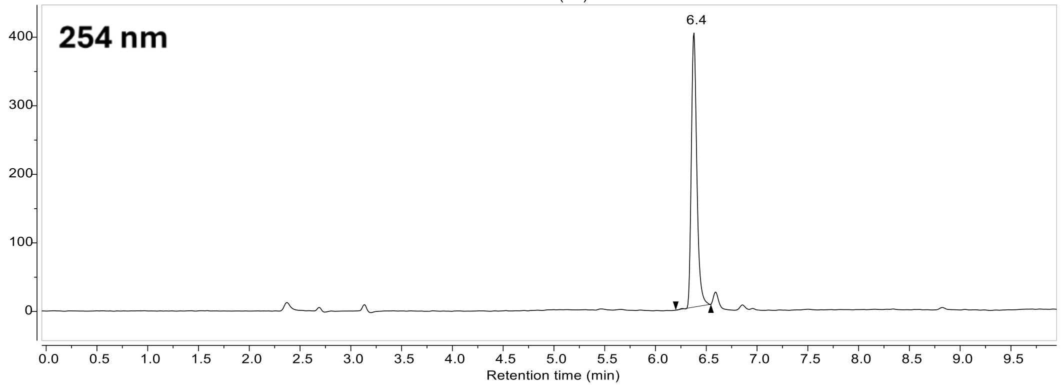
254 nm
6.4
Retention time (min)

#### Compound 38:

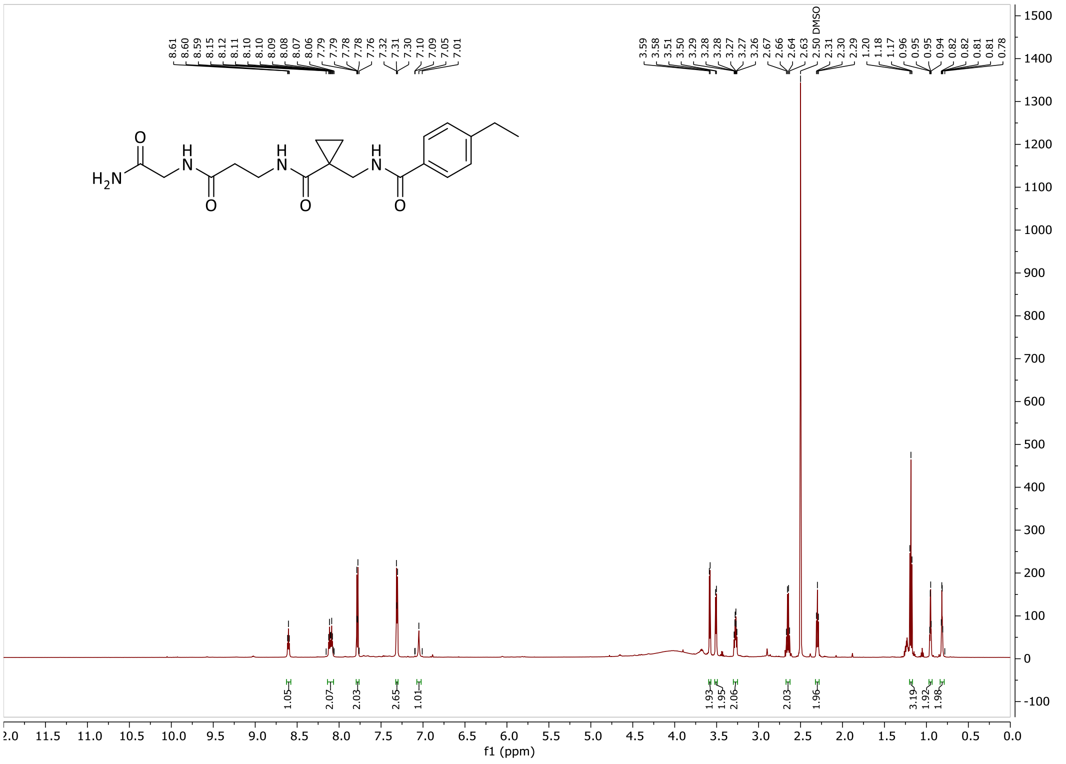

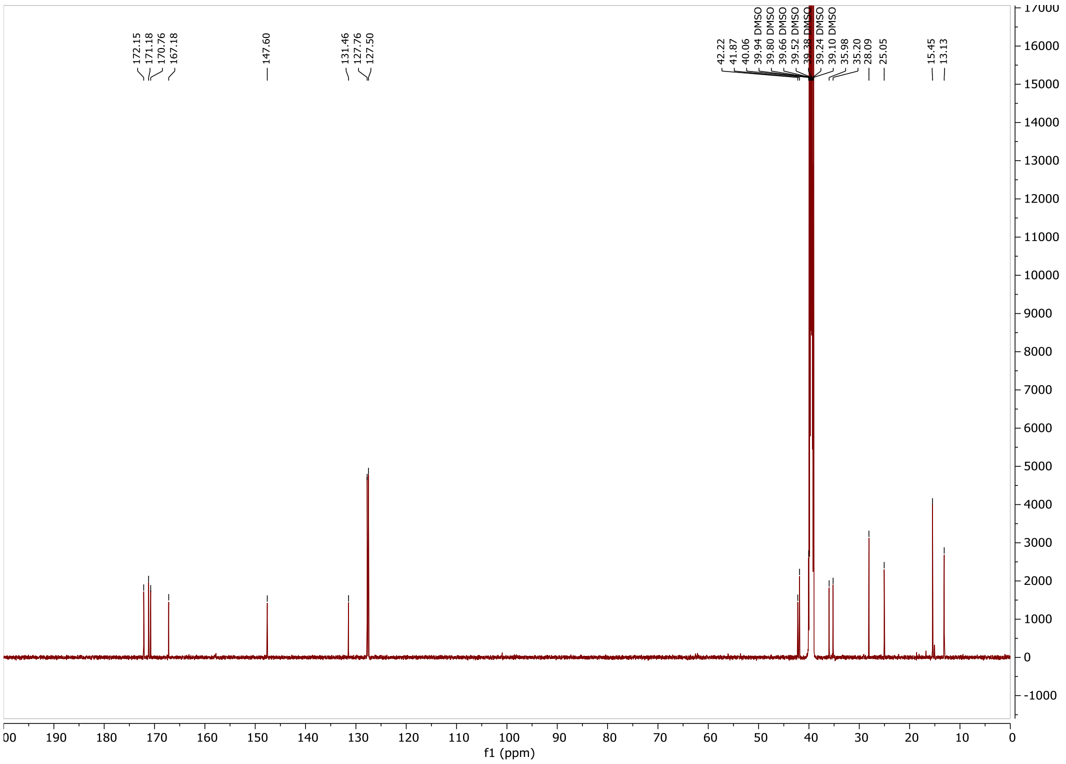

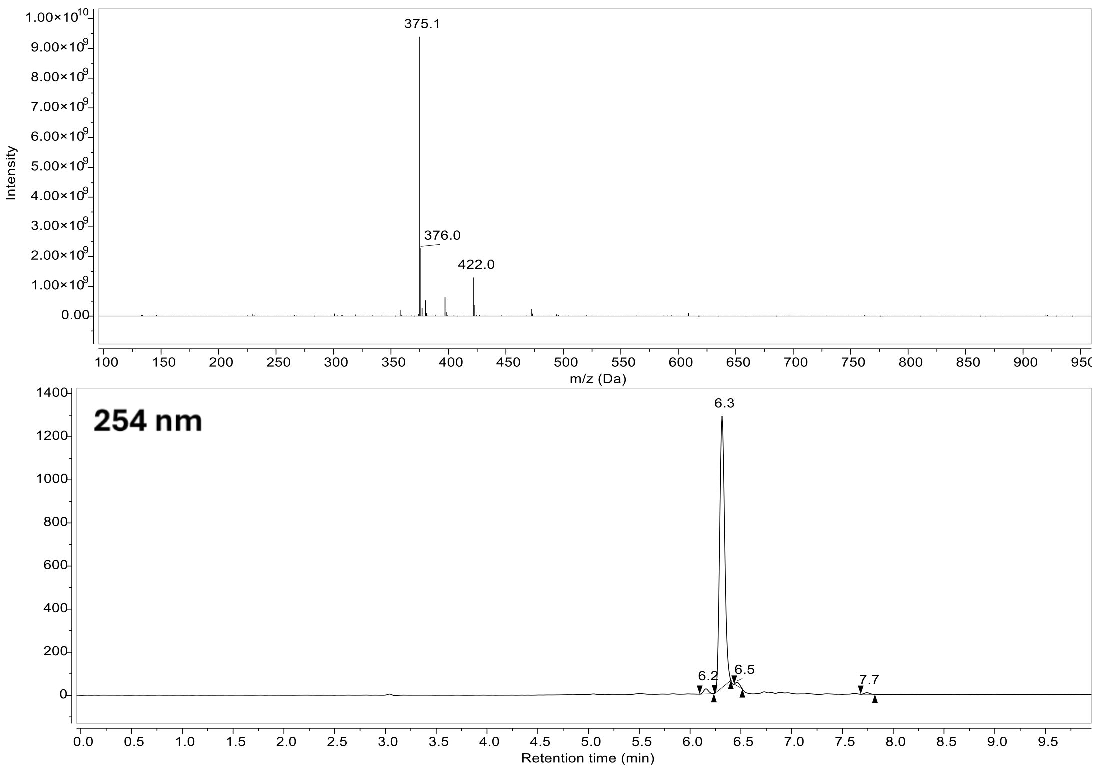

375.1
376.0
422.0
Intensity
m/z (Da)
254 nm
6.3
6.2
6.5
7.7
Retention time (min)

#### Compound 16:

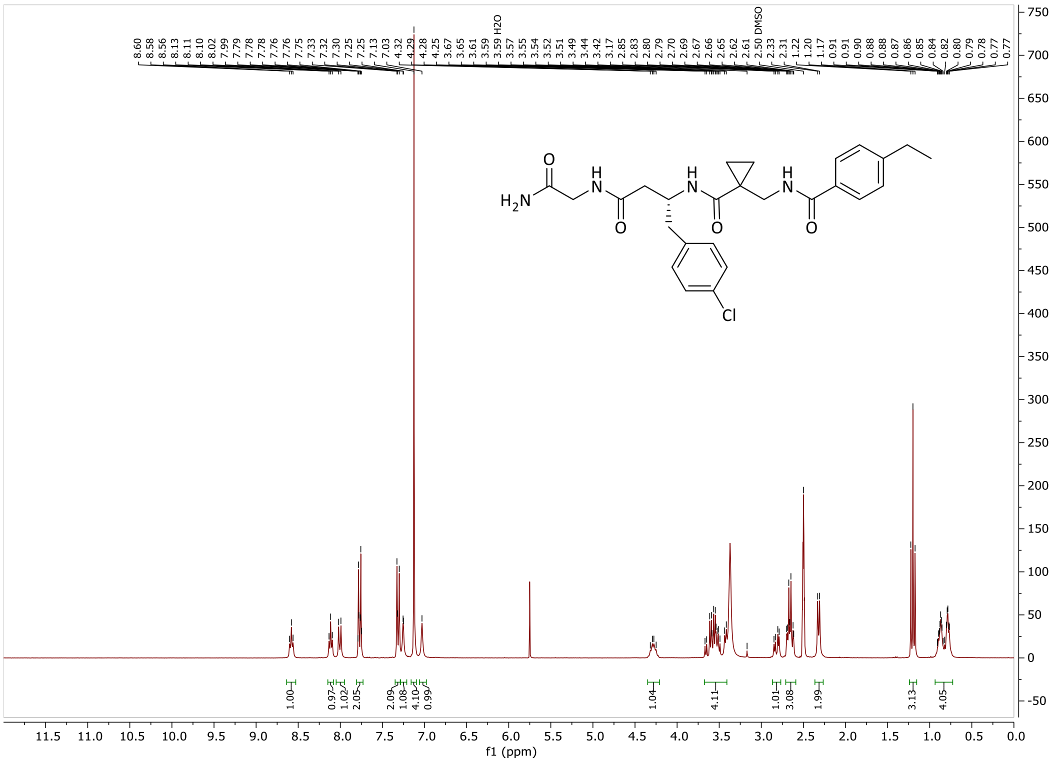

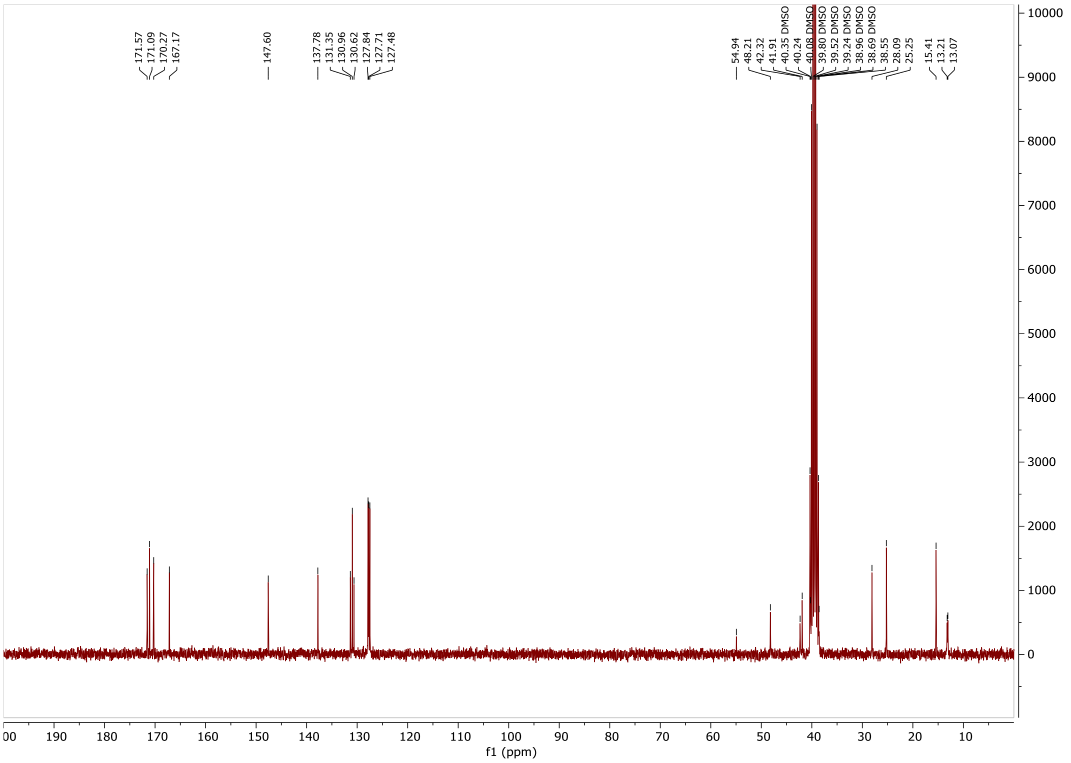

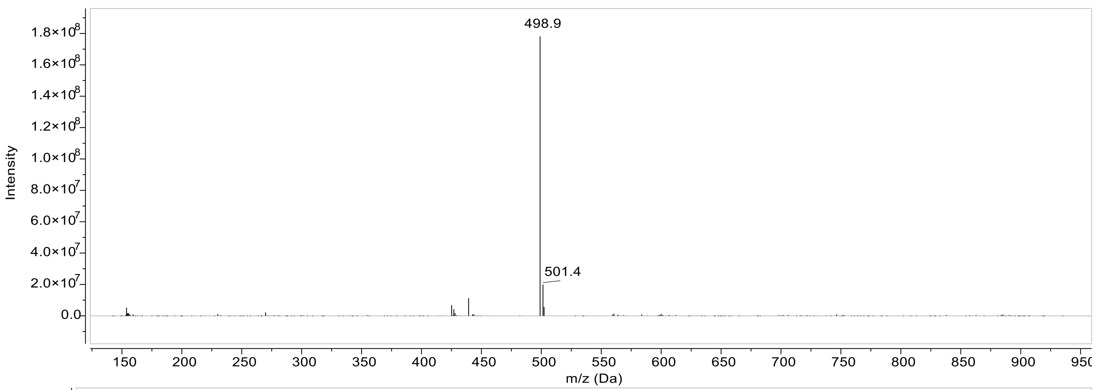

498.9
501.4
Intensity
m/z (Da)

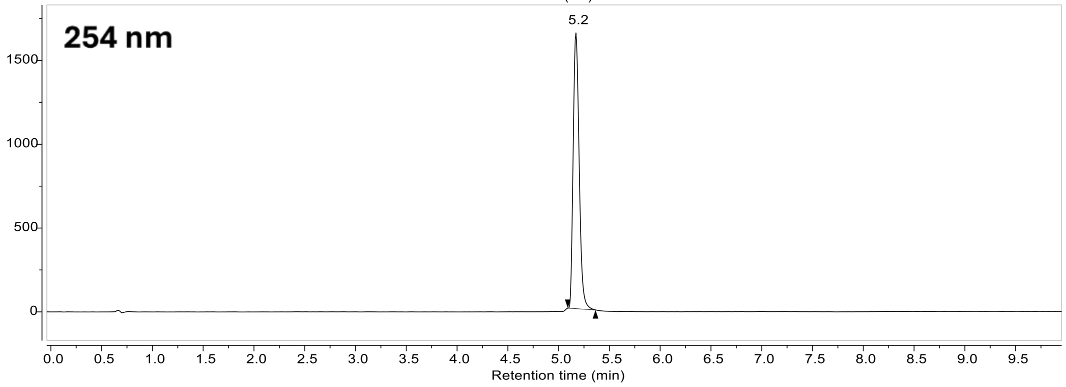

254 nm
5.2
Retention time (min)

#### Compound 17:

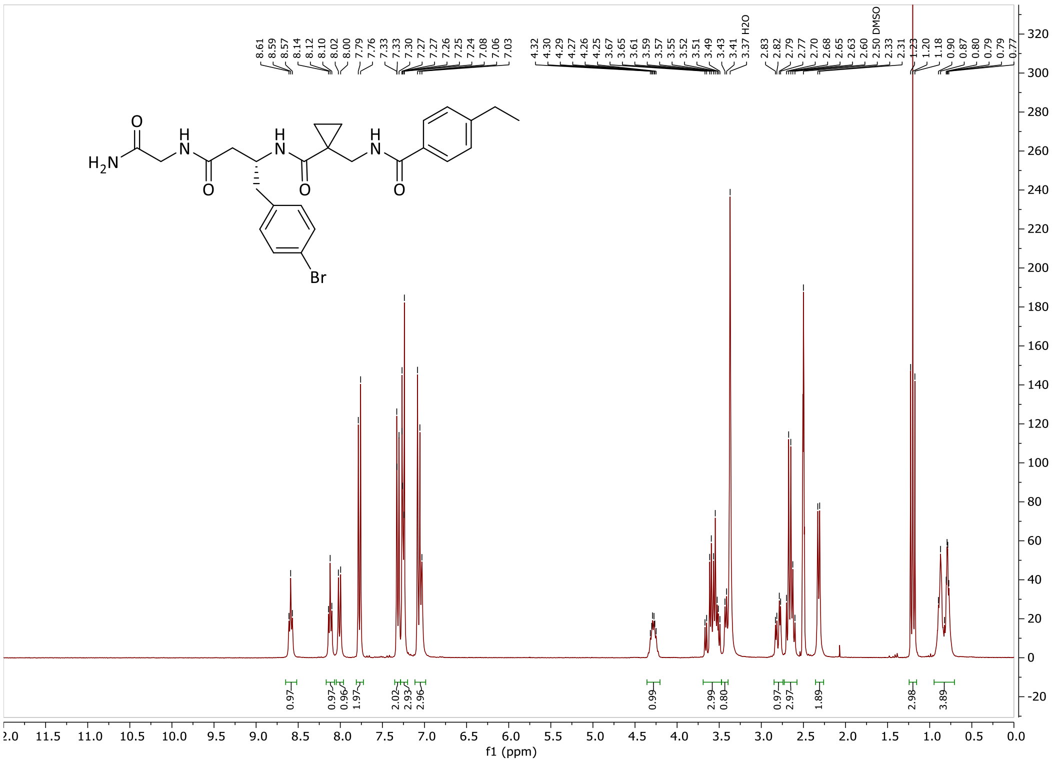
8.61 8.59 8.57 8.14 8.12 8.10 8.02 8.00 7.79 7.76 7.33 7.33 7.30 7.27 7.27 7.26 7.25 7.24 7.08 7.06 7.03
4.32 4.30 4.29 4.27 4.26 4.25 3.67 3.65 3.61 3.59 3.57 3.55 3.52 3.51 3.49 3.43 3.41 3.37 H2O
2.83 2.82 2.79 2.77 2.70 2.68 2.65 2.63 2.60 2.50 DMSO 2.33 2.31 1.23 1.20 1.18 0.90 0.87 0.80 0.79 0.77
0.97 0.97 0.96 1.97 2.02 2.93 2.96 0.99 2.99 0.80 0.97 2.97 1.89 2.98 3.89
H2N
O
H
N
O
H
N
O
H
N
O
Br
f1 (ppm)

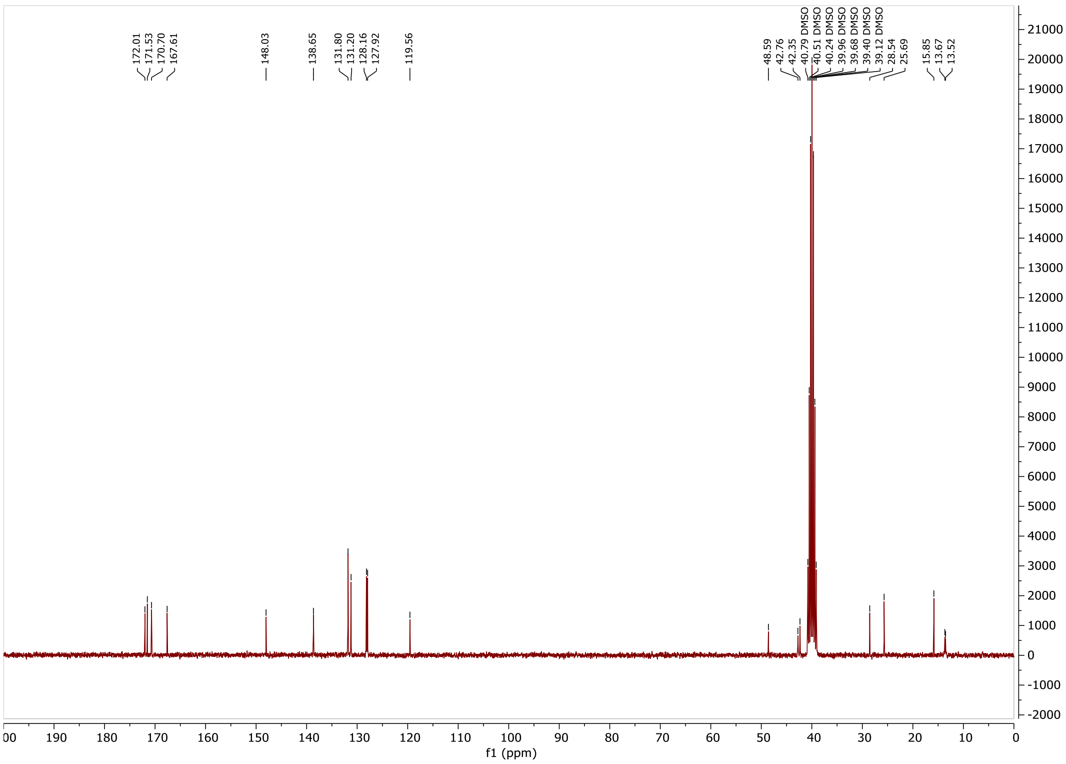
172.01 171.53 170.70 167.61
148.03
138.65
131.80 131.20 128.16 127.92
119.56
48.59 42.76 42.35 40.79 DMSO 40.51 DMSO 40.24 DMSO 39.96 DMSO 39.68 DMSO 39.40 DMSO 39.12 DMSO 28.54 25.69
15.85 13.67 13.52
f1 (ppm)

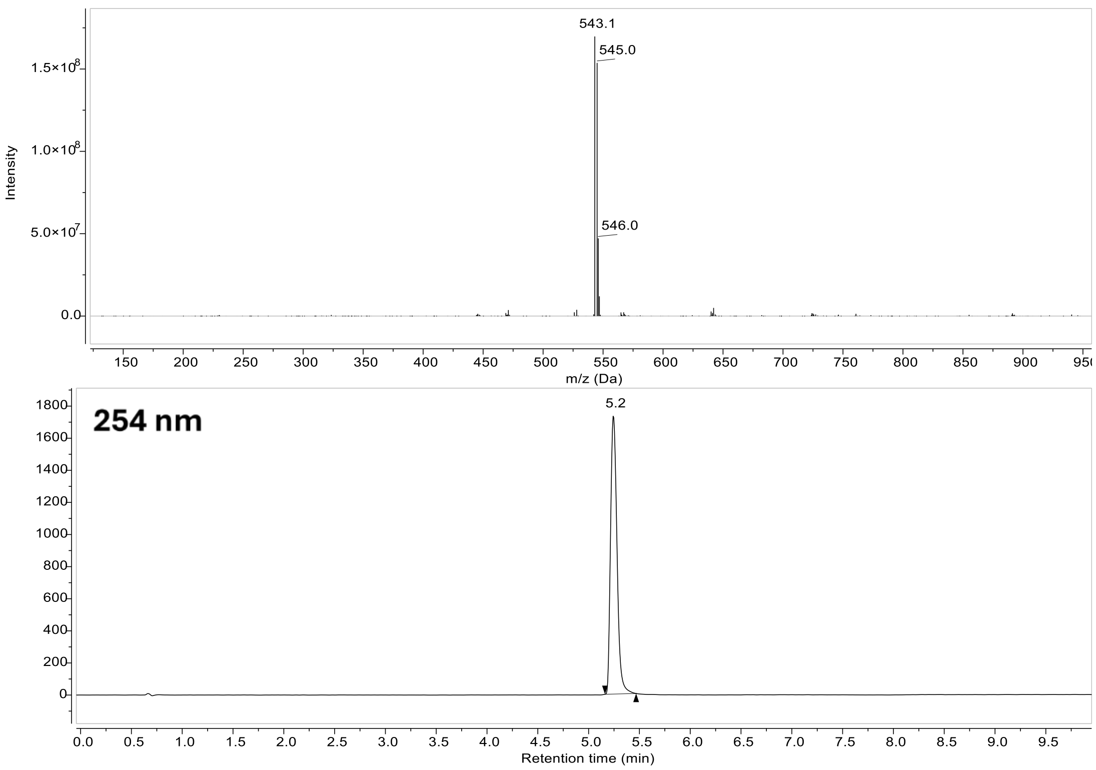

543.1
545.0
546.0
Intensity
1.5×10^8
1.0×10^8
5.0×10^7
0.0
150
200
250
300
350
400
450
500
550
600
650
700
750
800
850
900
950
m/z (Da)
254 nm
5.2
1800
1600
1400
1200
1000
800
600
400
200
0
0.0
0.5
1.0
1.5
2.0
2.5
3.0
3.5
4.0
4.5
5.0
5.5
6.0
6.5
7.0
7.5
8.0
8.5
9.0
9.5
Retention time (min)

#### Compound 33:

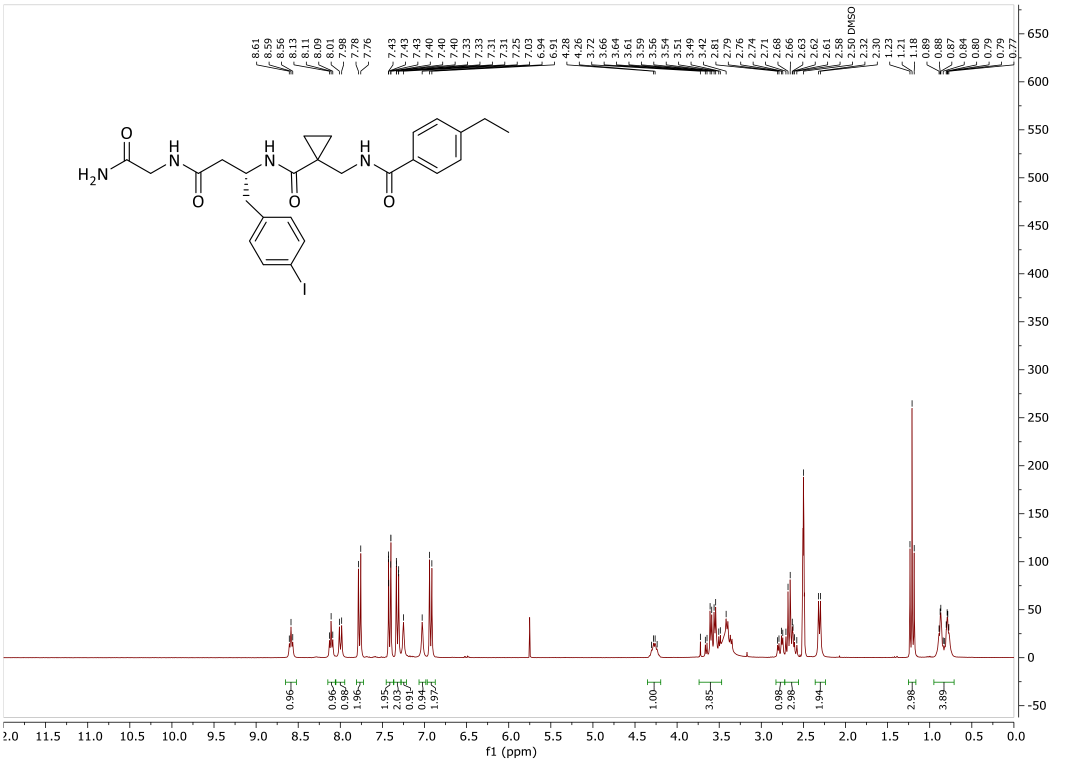

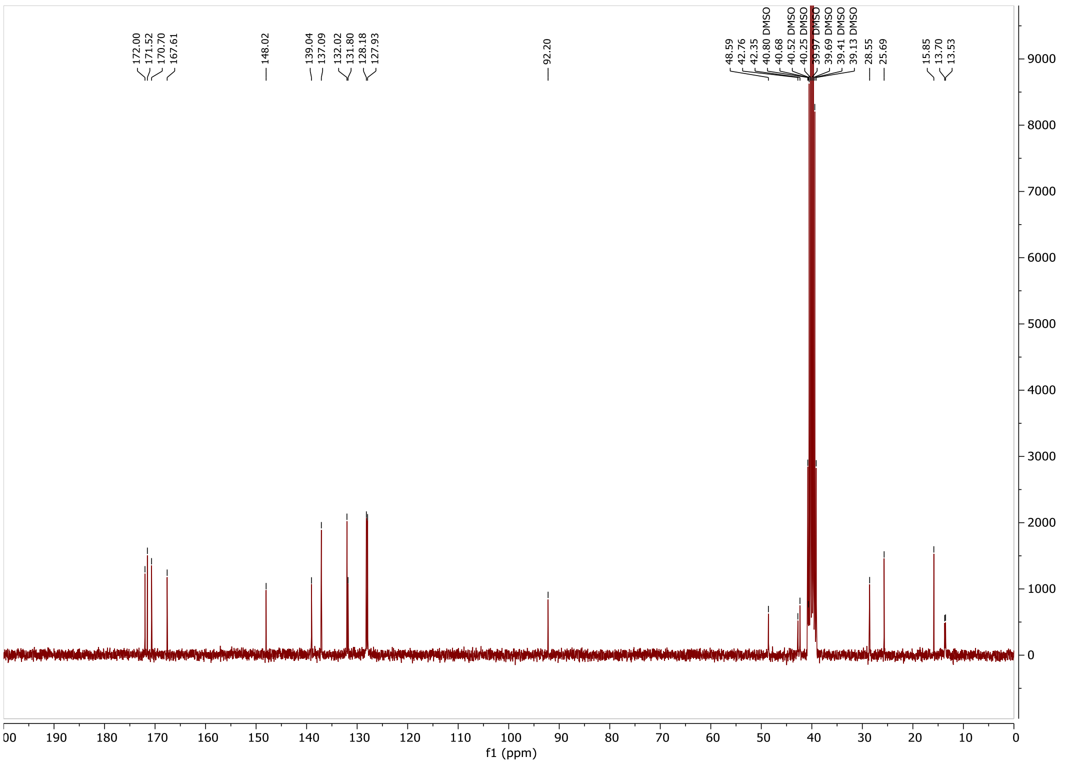

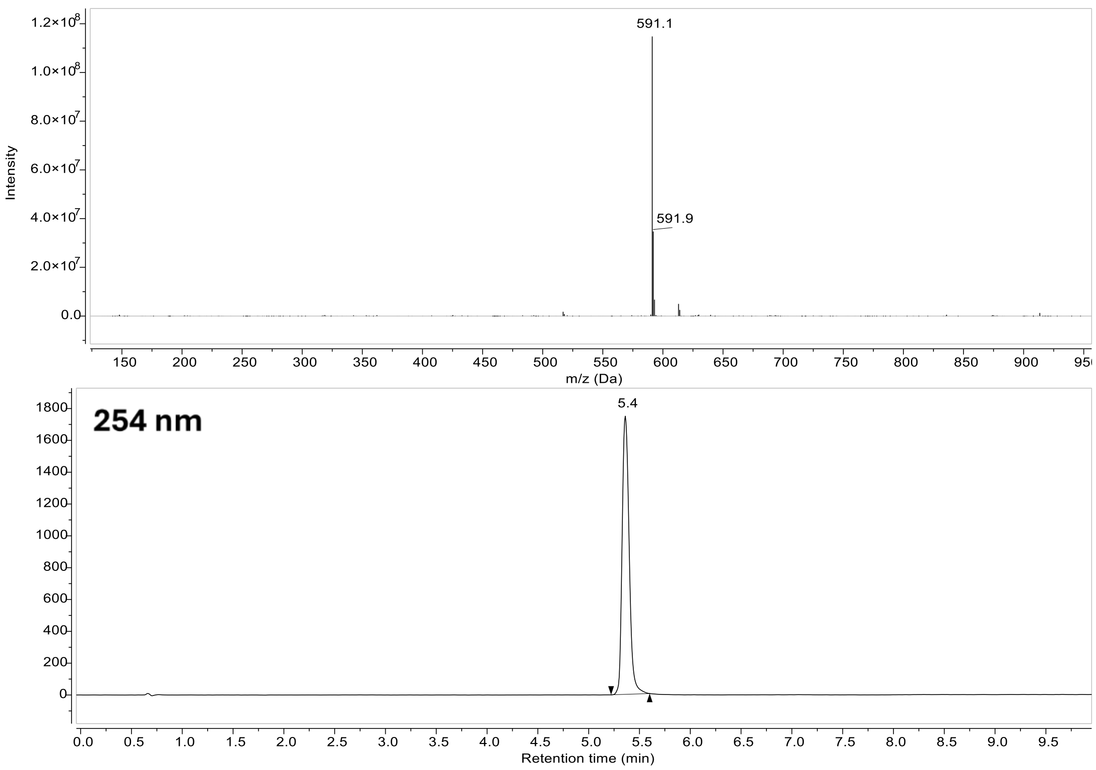
591.1
591.9
Intensity
m/z (Da)
254 nm
5.4
Retention time (min)

### Compound 34:

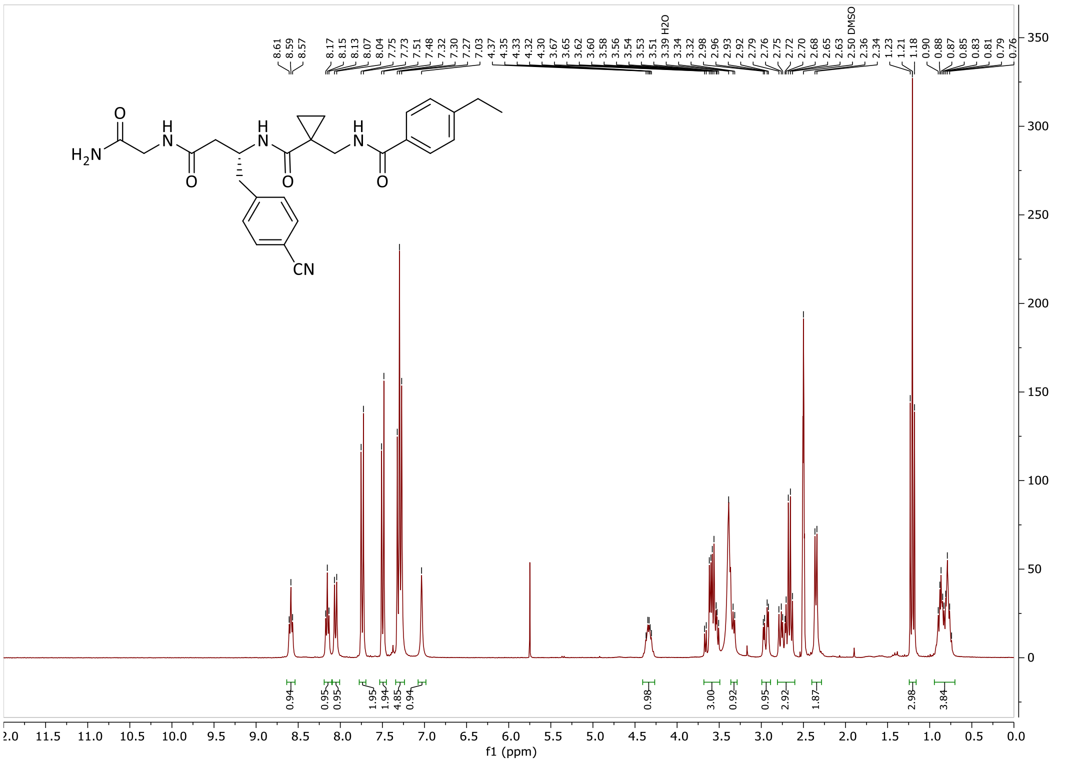

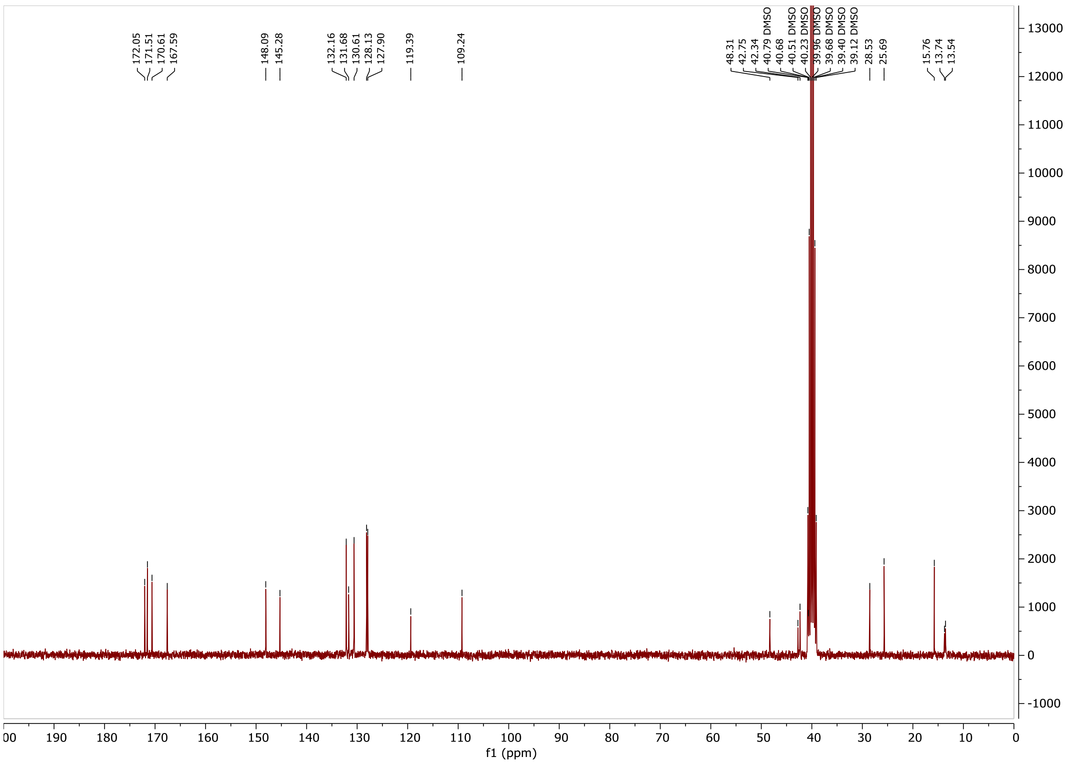

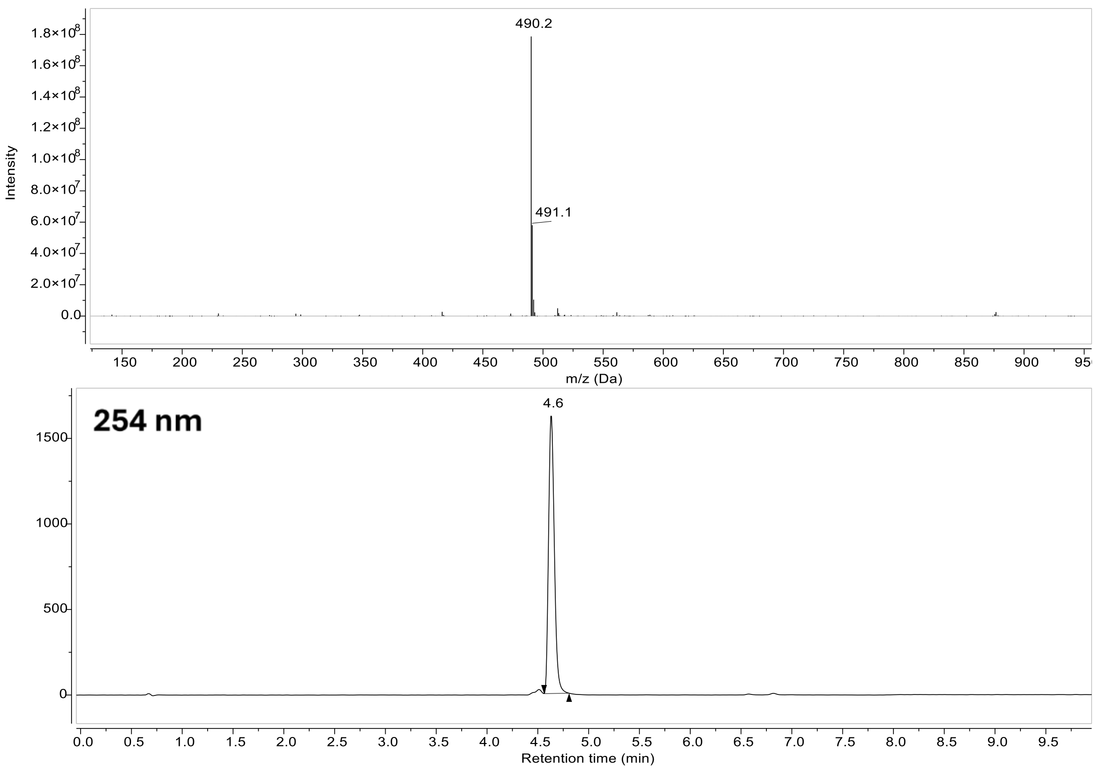

490.2
491.1
Intensity
m/z (Da)
254 nm
4.6
Retention time (min)

### Compound 39:

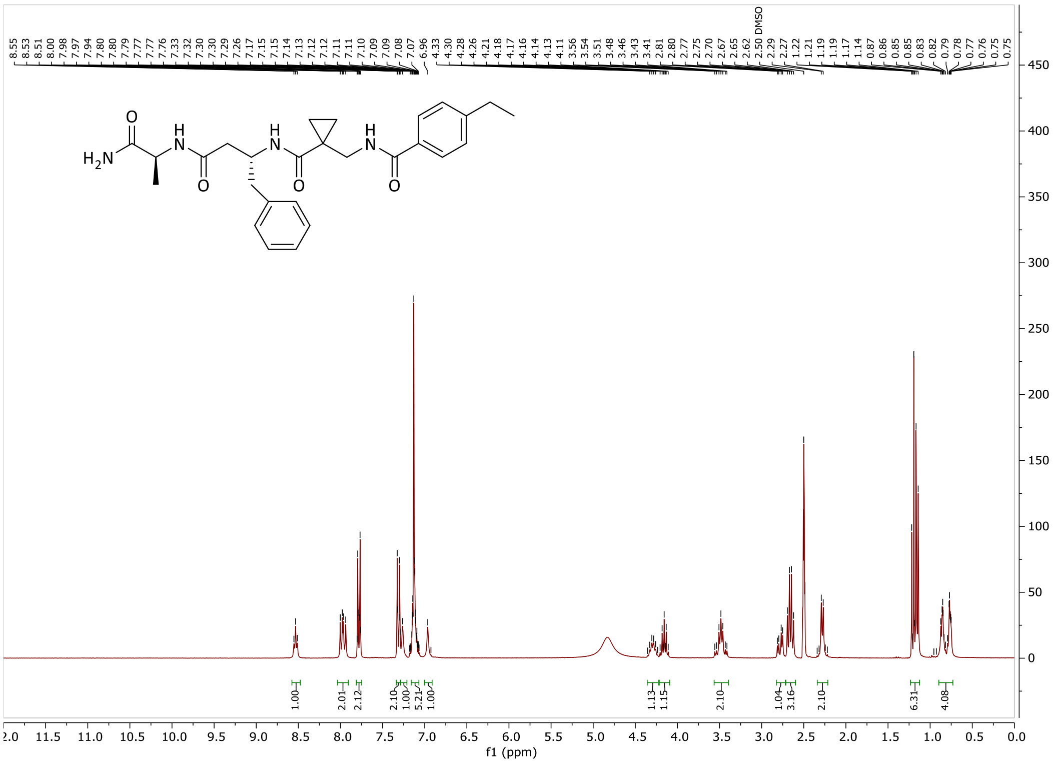

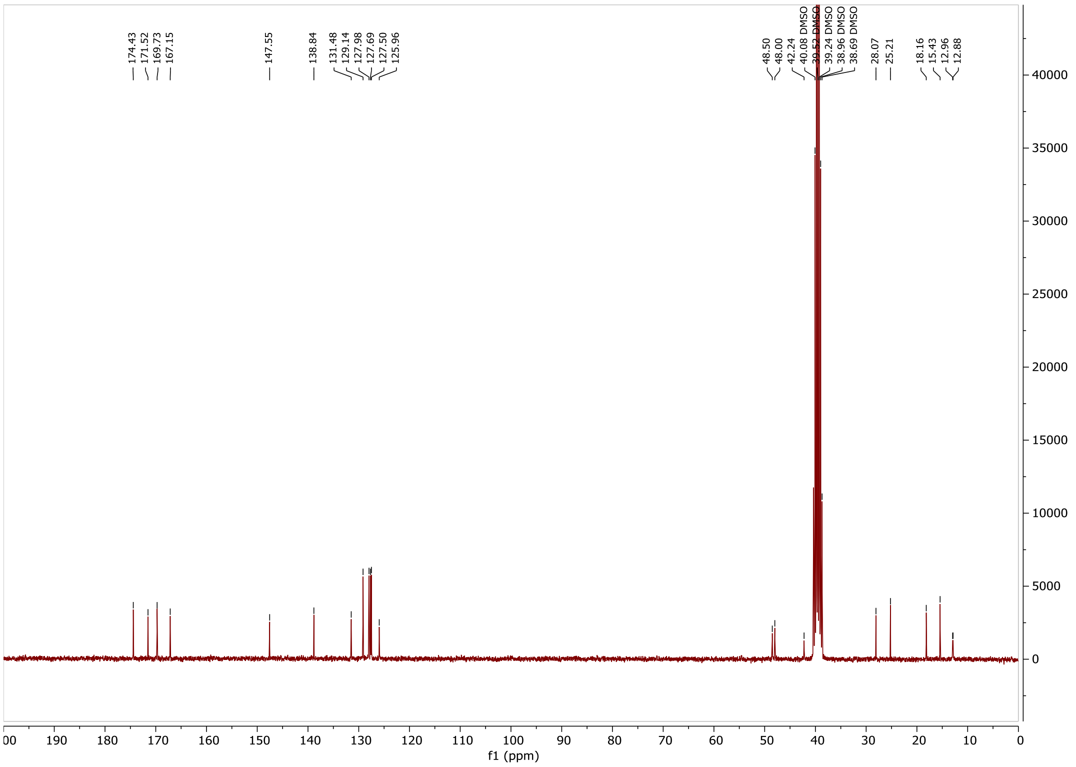

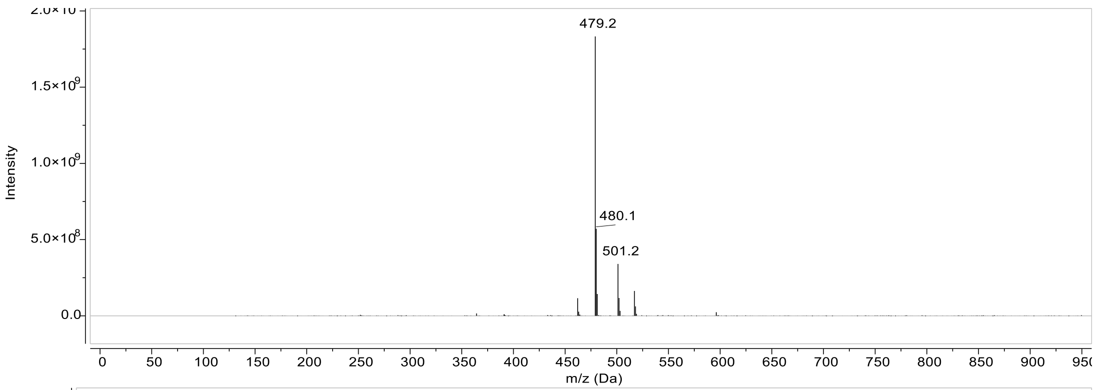
479.2
480.1
501.2
Intensity
m/z (Da)

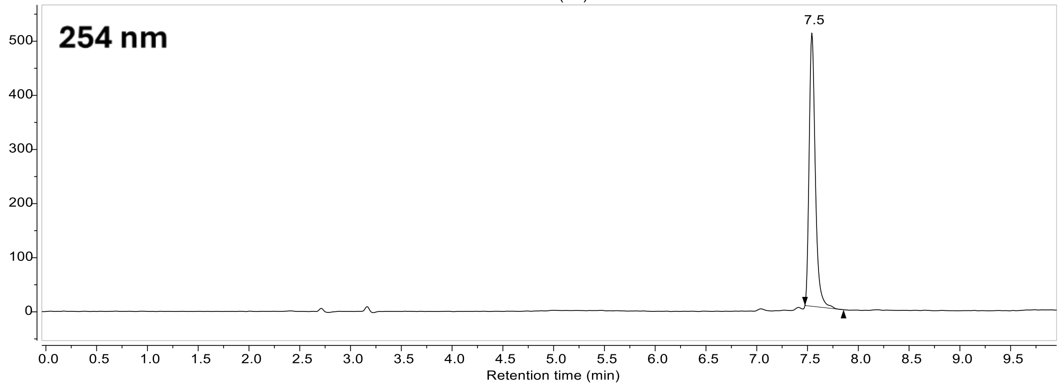
254 nm
7.5
Retention time (min)

### Compound 40:

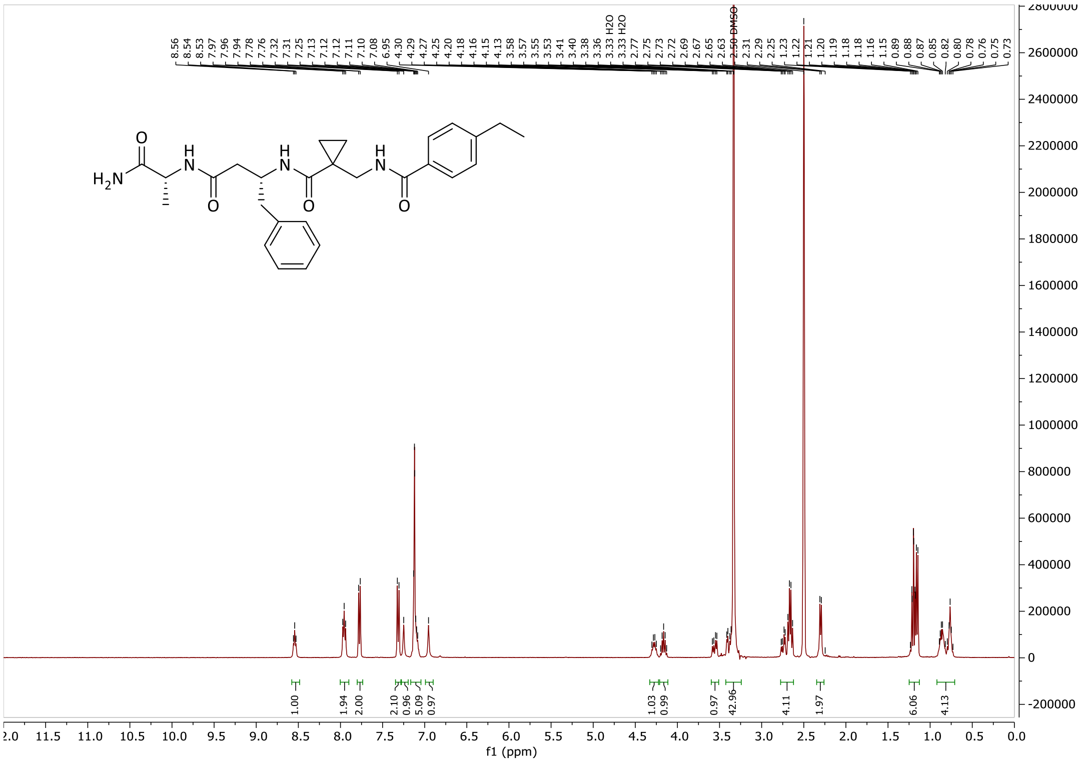
8.56
8.54
8.53
7.97
7.96
7.94
7.78
7.76
7.32
7.31
7.25
7.13
7.12
7.12
7.11
7.10
7.08
6.95
4.30
4.29
4.27
4.25
4.20
4.18
4.16
4.15
4.13
3.58
3.57
3.55
3.53
3.41
3.40
3.38
3.36
3.33 H2O
3.33 H2O
2.77
2.75
2.73
2.72
2.69
2.67
2.65
2.63
2.50 DMSO
2.31
2.29
2.25
2.23
1.22
1.21
1.20
1.19
1.18
1.18
1.16
1.15
0.89
0.88
0.87
0.85
0.82
0.80
0.78
0.76
0.75
0.73
H2N
O
H
N
O
H
N
O
H
N
O
1.00
1.94
2.00
2.10
0.96
5.09
0.97
1.03
0.99
0.97
42.96
4.11
1.97
6.06
4.13
f1 (ppm)

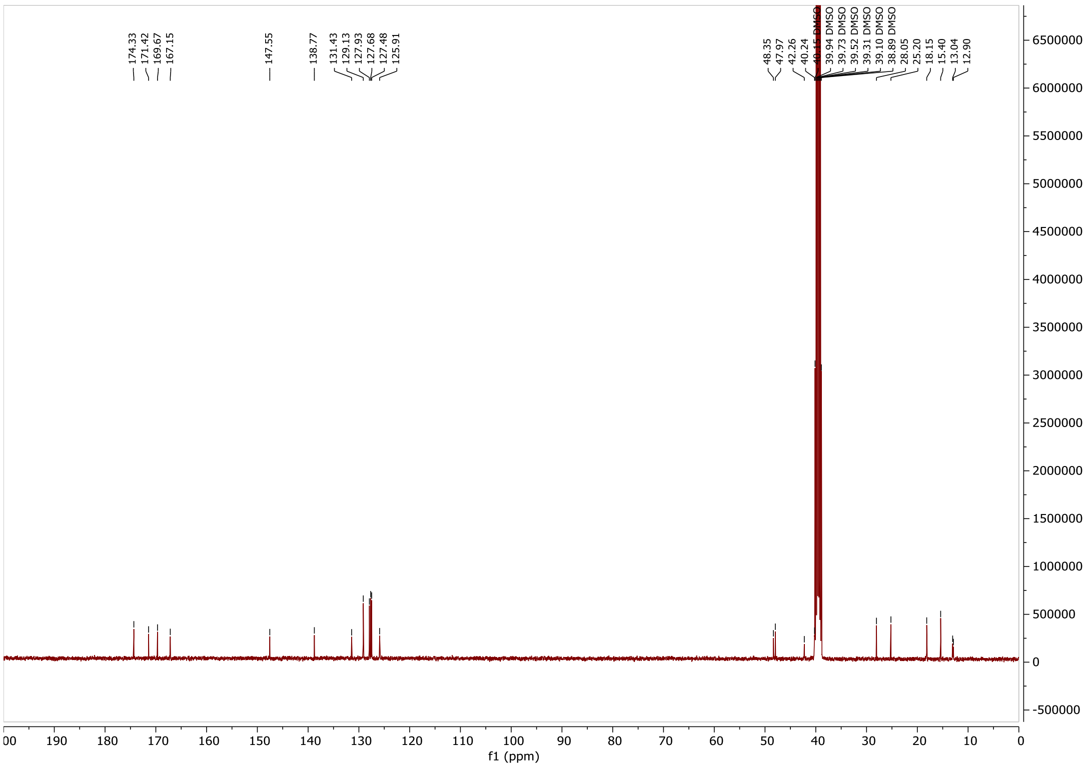
174.33
171.42
169.67
167.15
147.55
138.77
131.43
129.13
127.93
127.68
127.48
125.91
48.35
47.97
42.26
40.24
39.94 DMSO
39.73 DMSO
39.52 DMSO
39.31 DMSO
39.10 DMSO
38.89 DMSO
28.05
25.20
18.15
15.40
13.04
12.90
f1 (ppm)

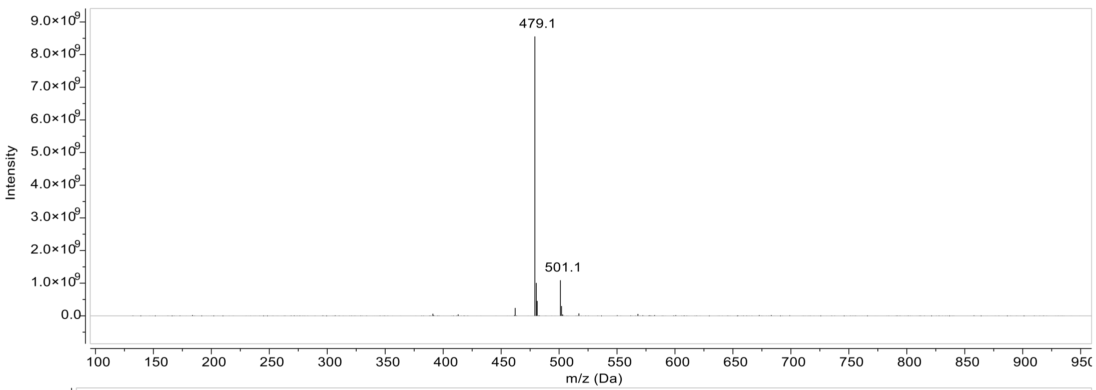
479.1
501.1
Intensity
m/z (Da)

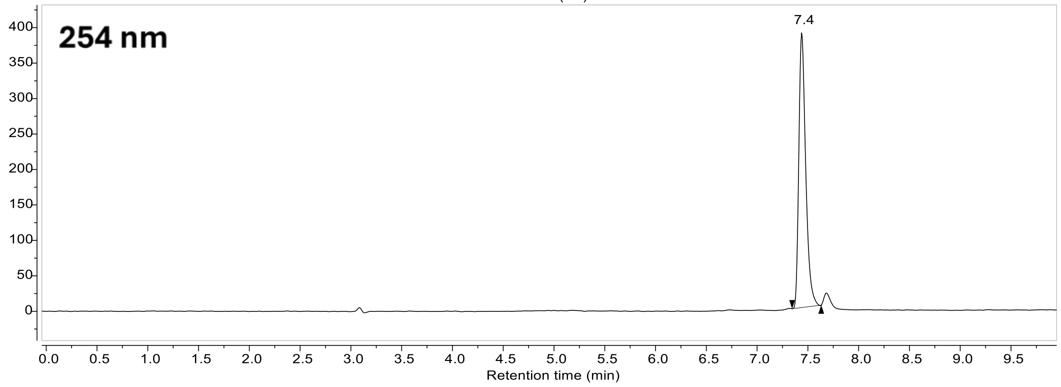
254 nm
7.4
Retention time (min)

**Compound 41:**

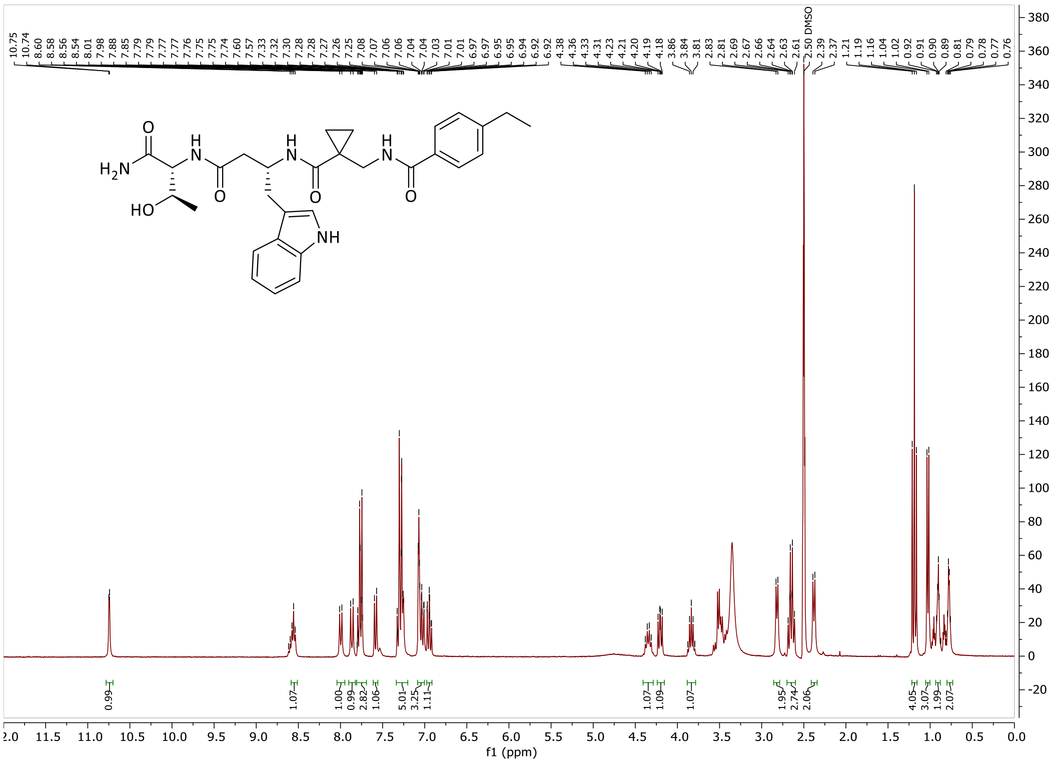

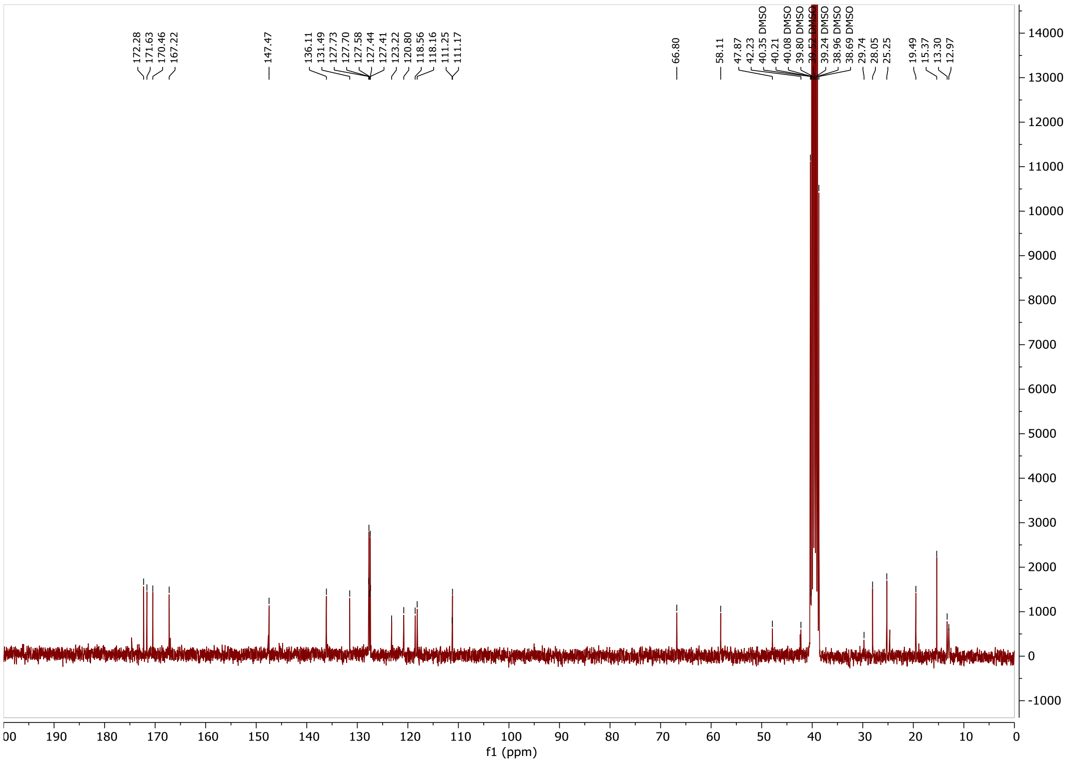

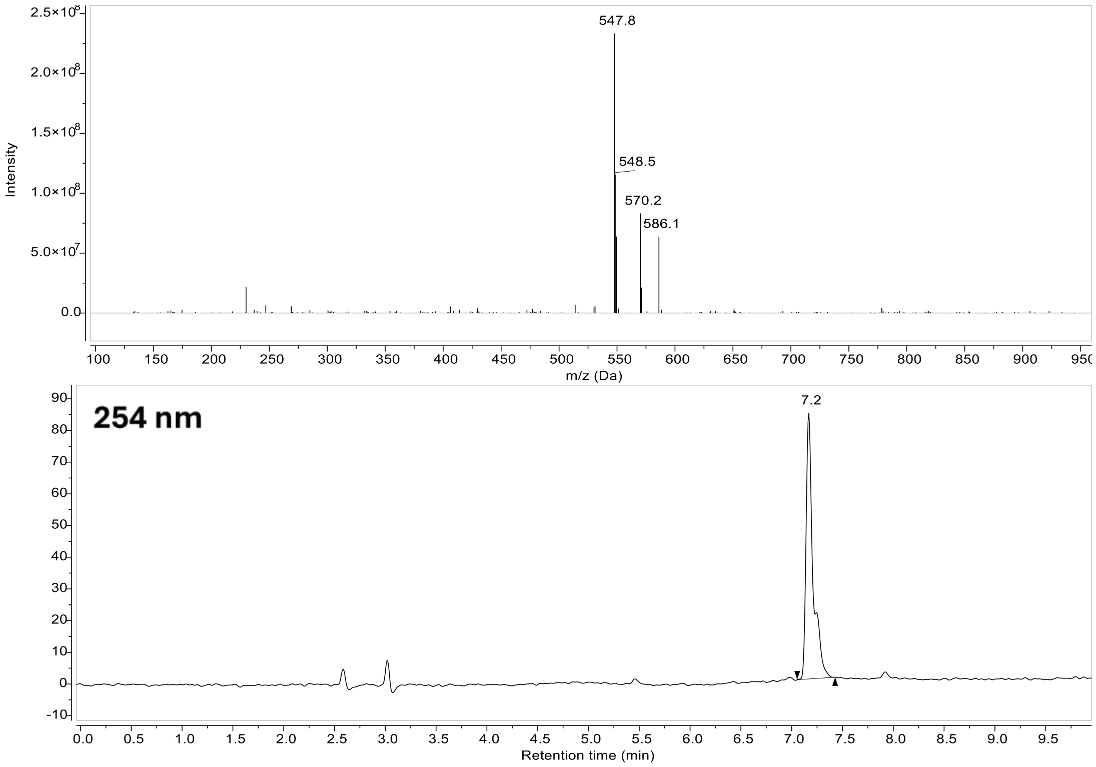

547.8
548.5
570.2
586.1
Intensity
m/z (Da)
254 nm
7.2
Retention time (min)

**Compound 42:**

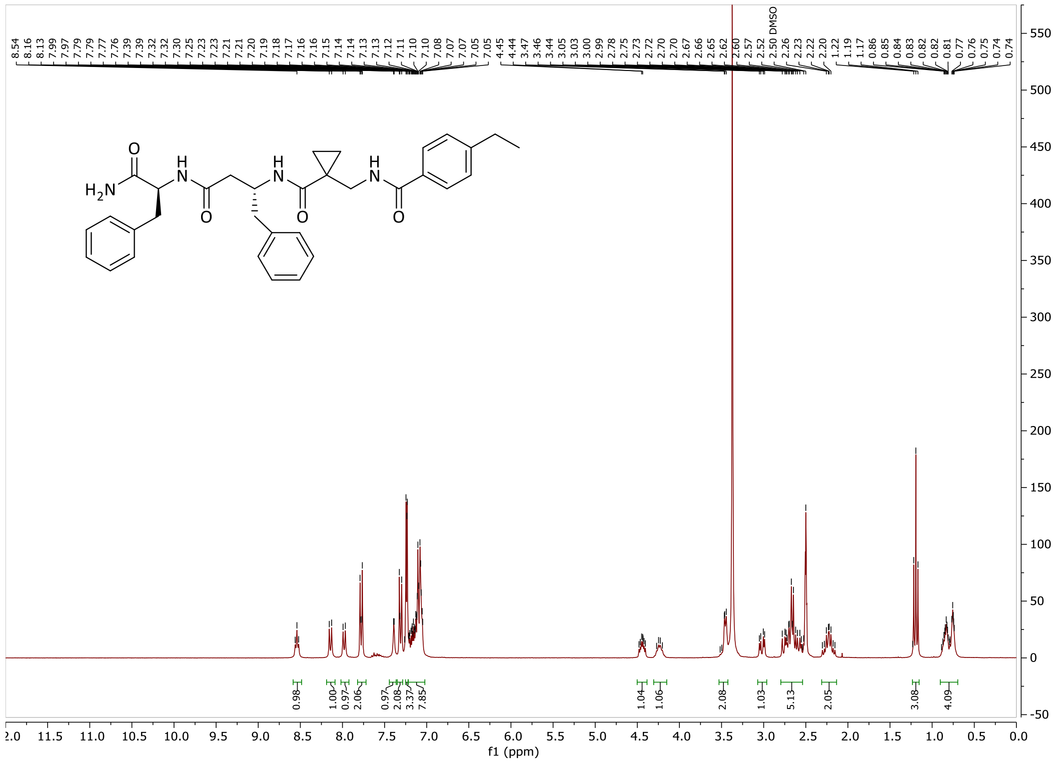

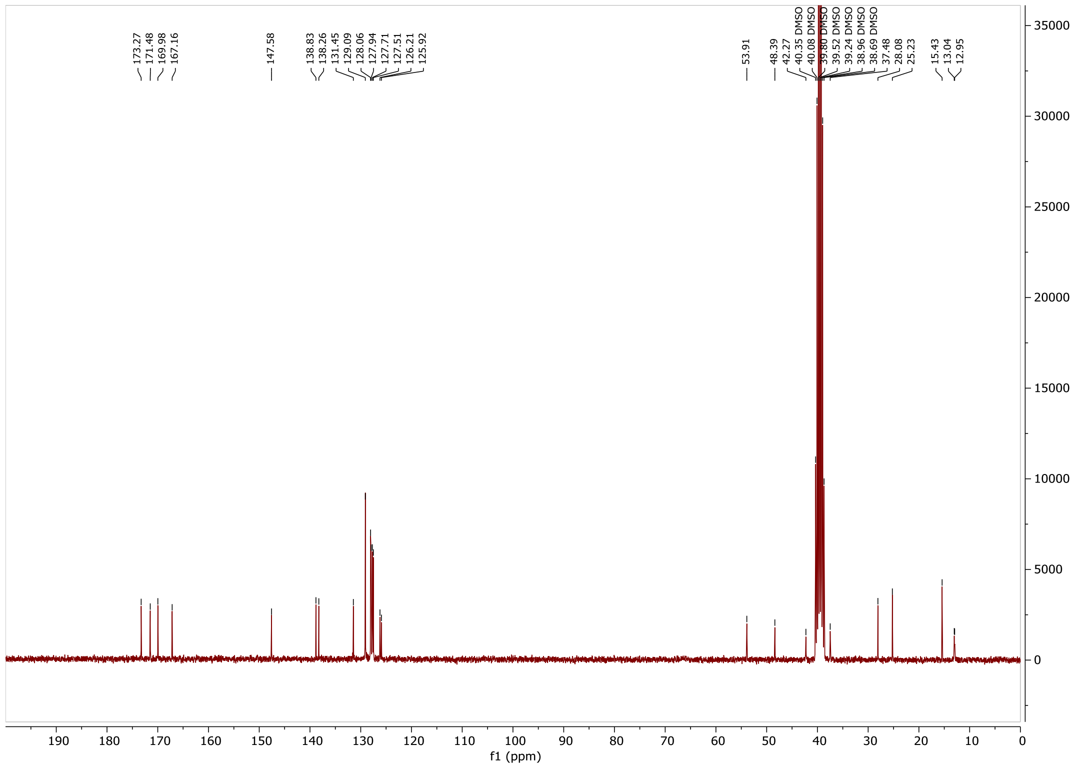

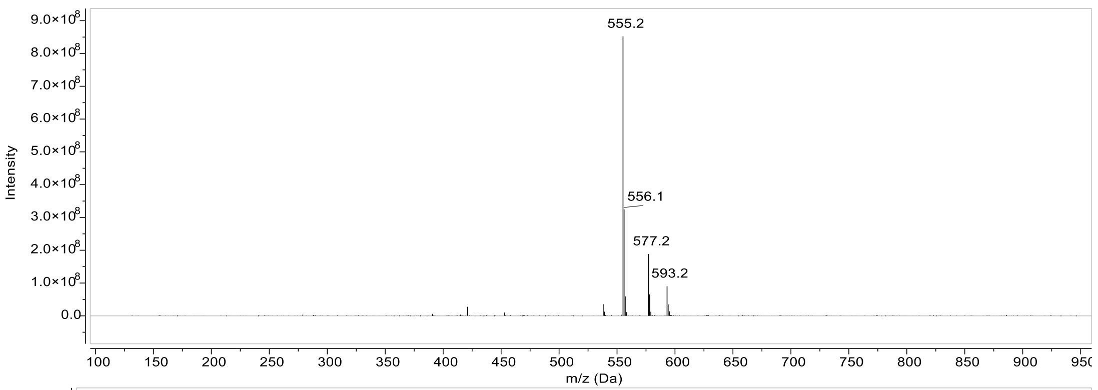
Intensity
m/z (Da)
555.2
556.1
577.2
593.2

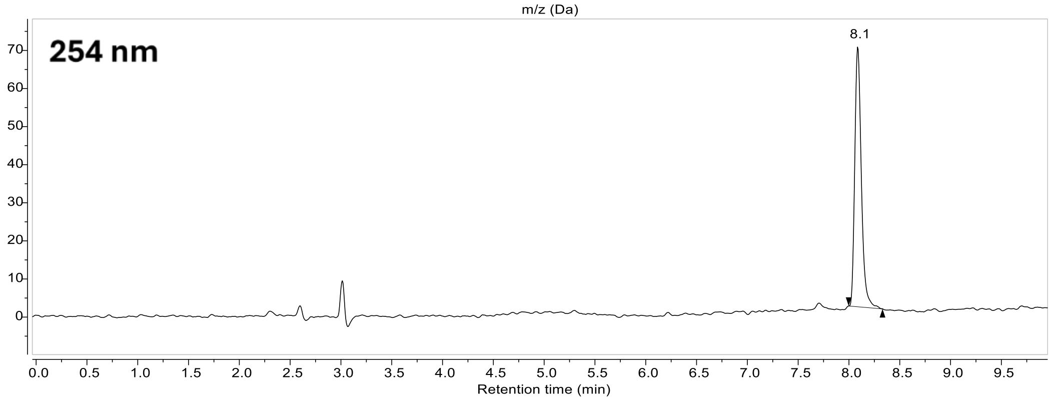
254 nm
8.1
Retention time (min)

**Compound 43:**

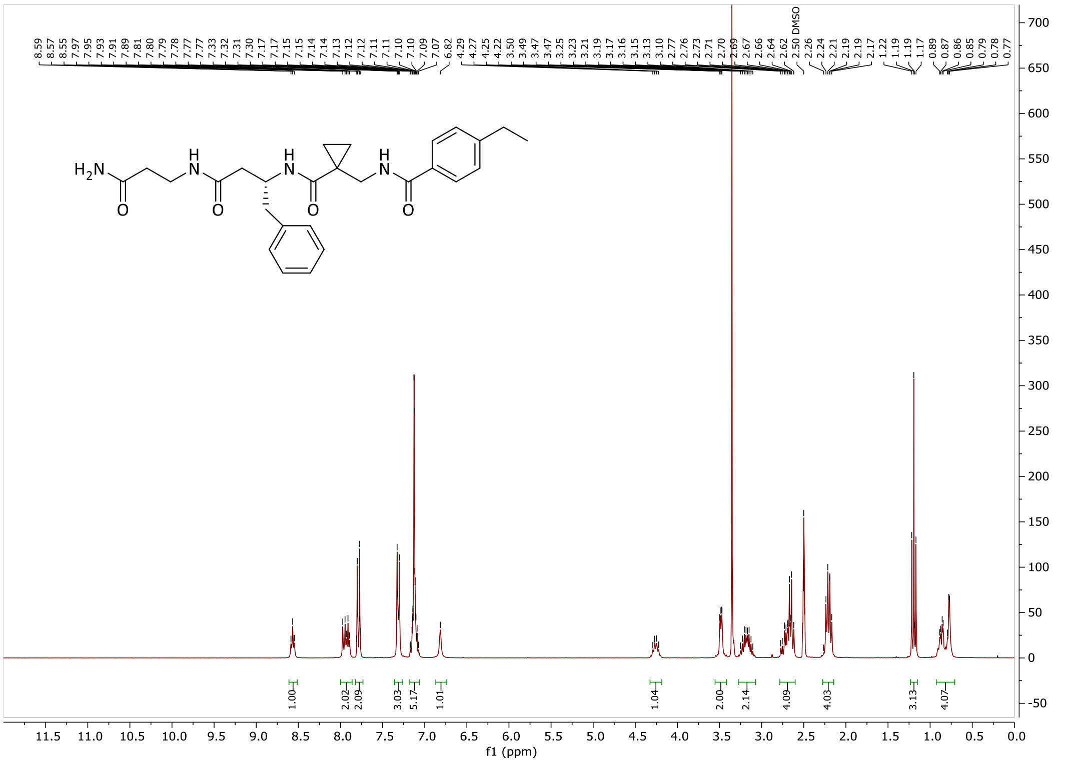

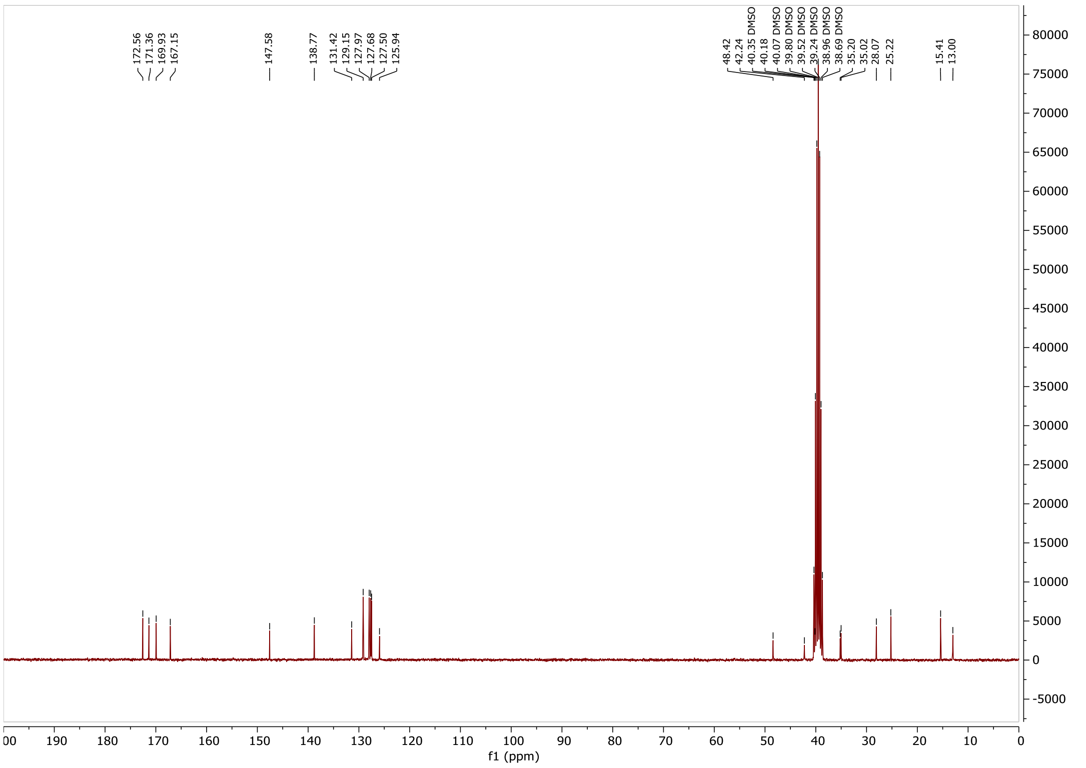

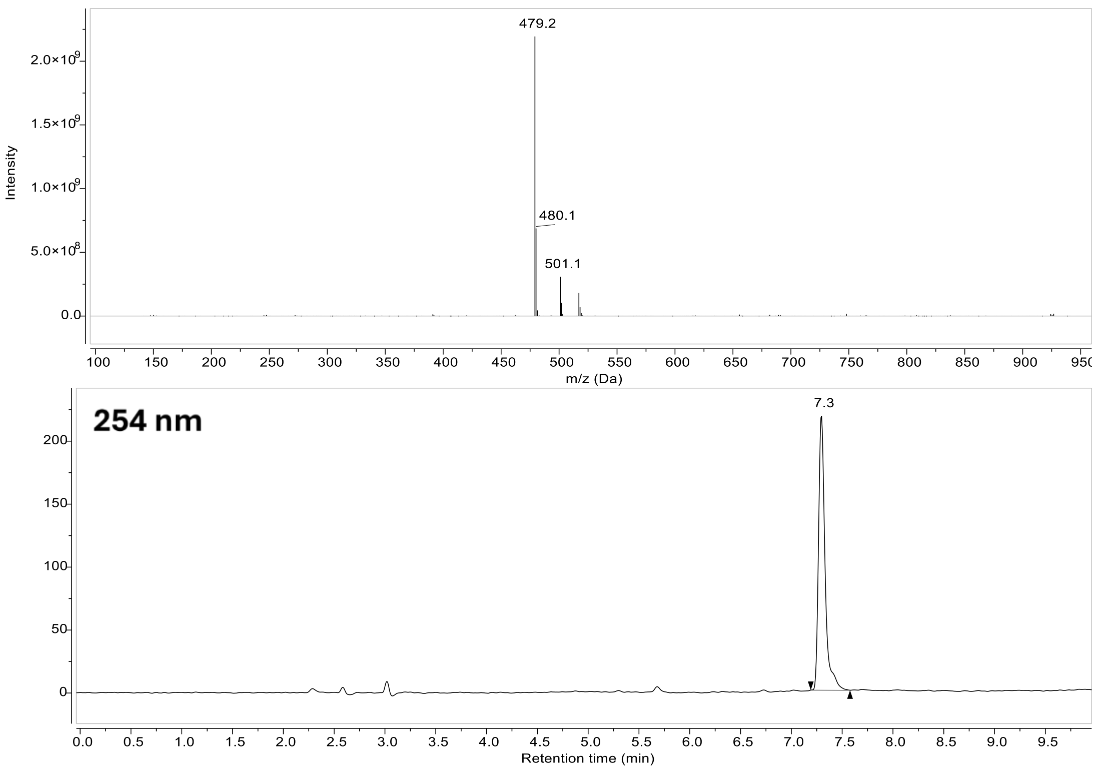
479.2
480.1
501.1
Intensity
2.0×10^9
1.5×10^9
1.0×10^9
5.0×10^8
0.0
100 150 200 250 300 350 400 450 500 550 600 650 700 750 800 850 900 950
m/z (Da)
254 nm
7.3
200
150
100
50
0
0.0 0.5 1.0 1.5 2.0 2.5 3.0 3.5 4.0 4.5 5.0 5.5 6.0 6.5 7.0 7.5 8.0 8.5 9.0 9.5
Retention time (min)

### Compound 45:

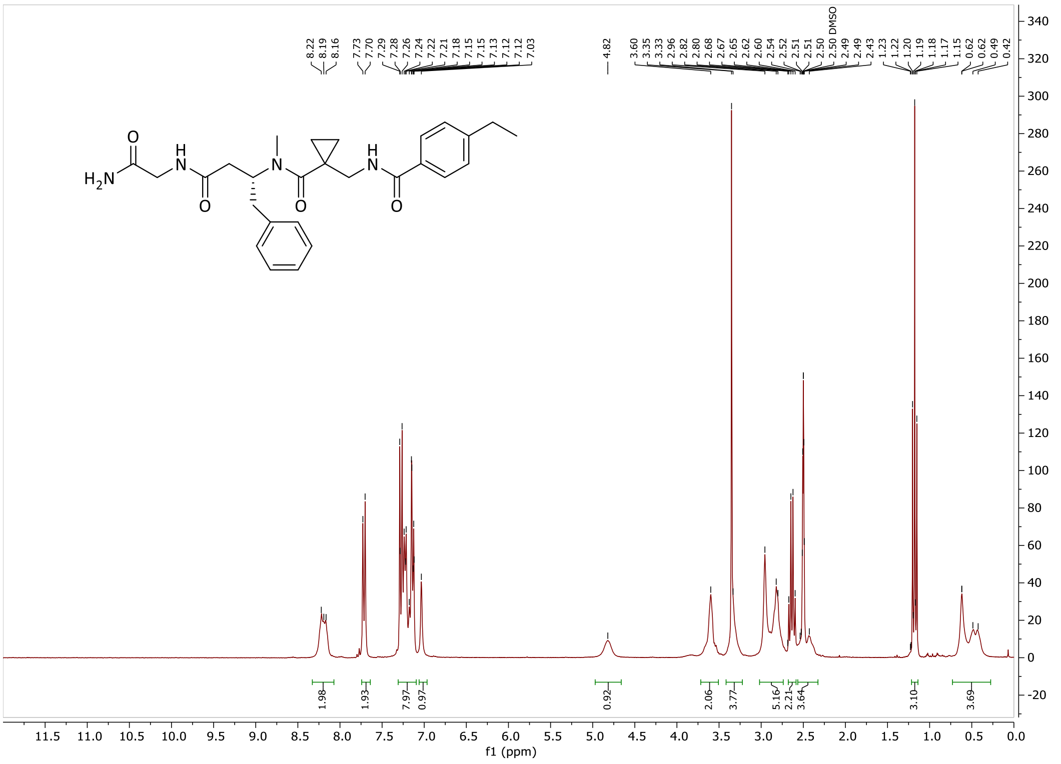

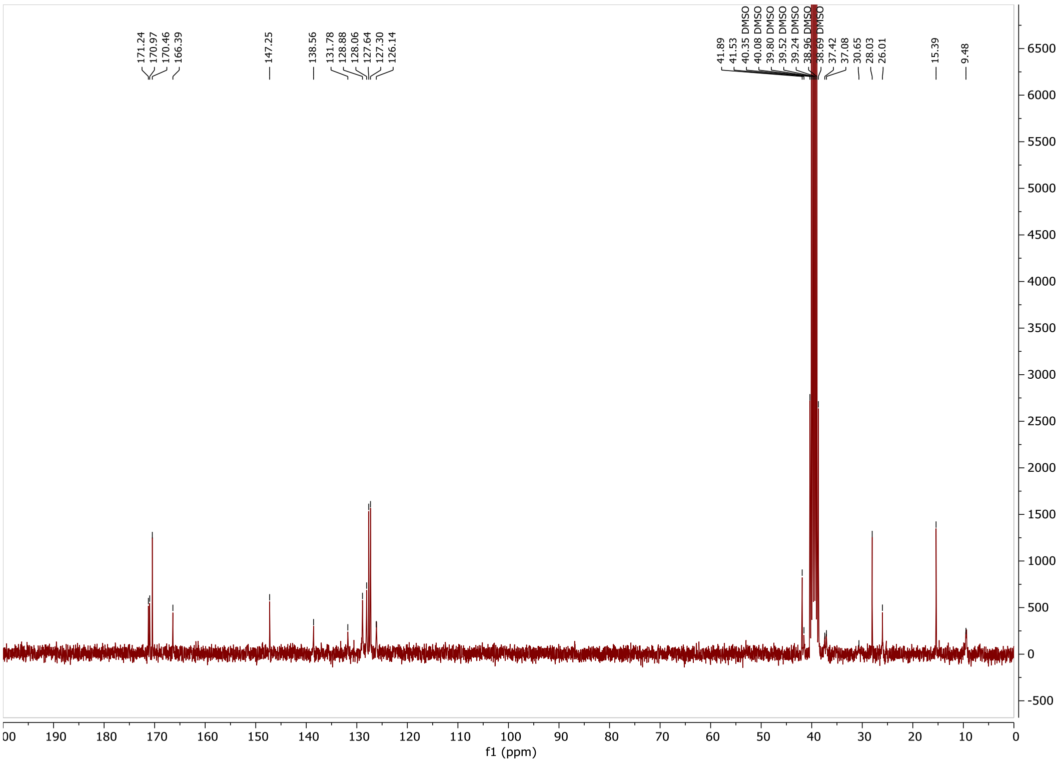

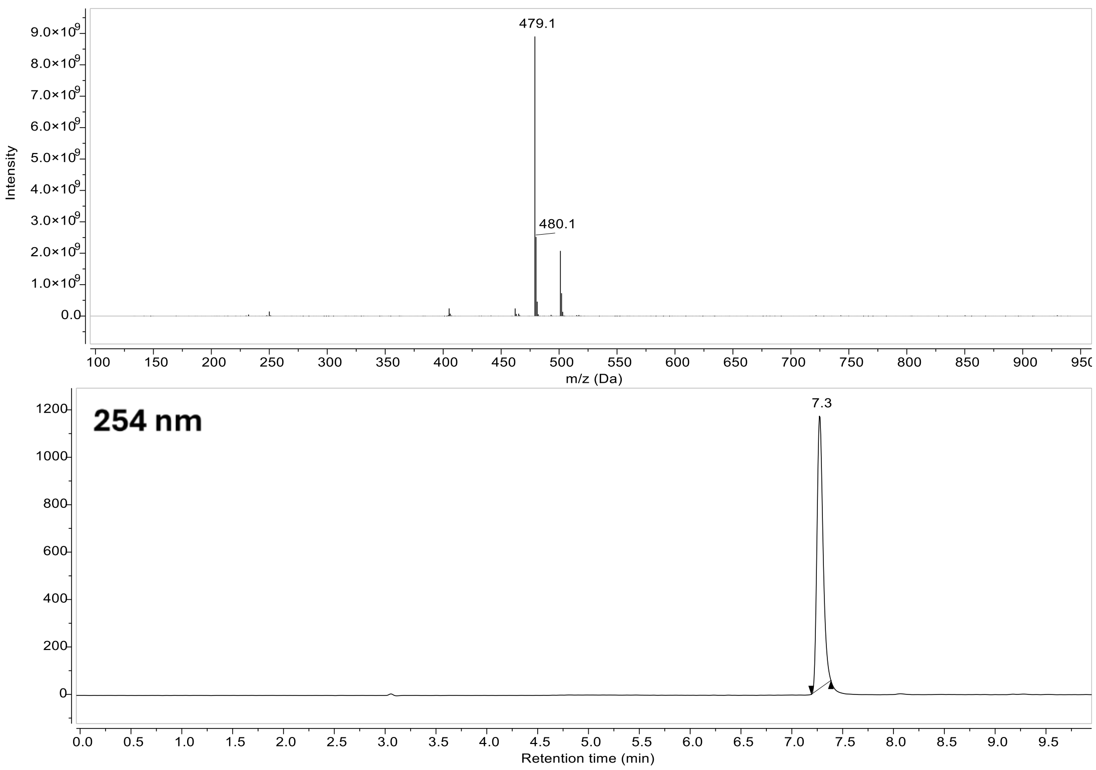

479.1
480.1
Intensity
m/z (Da)
254 nm
7.3
Retention time (min)

#### Compound 47:

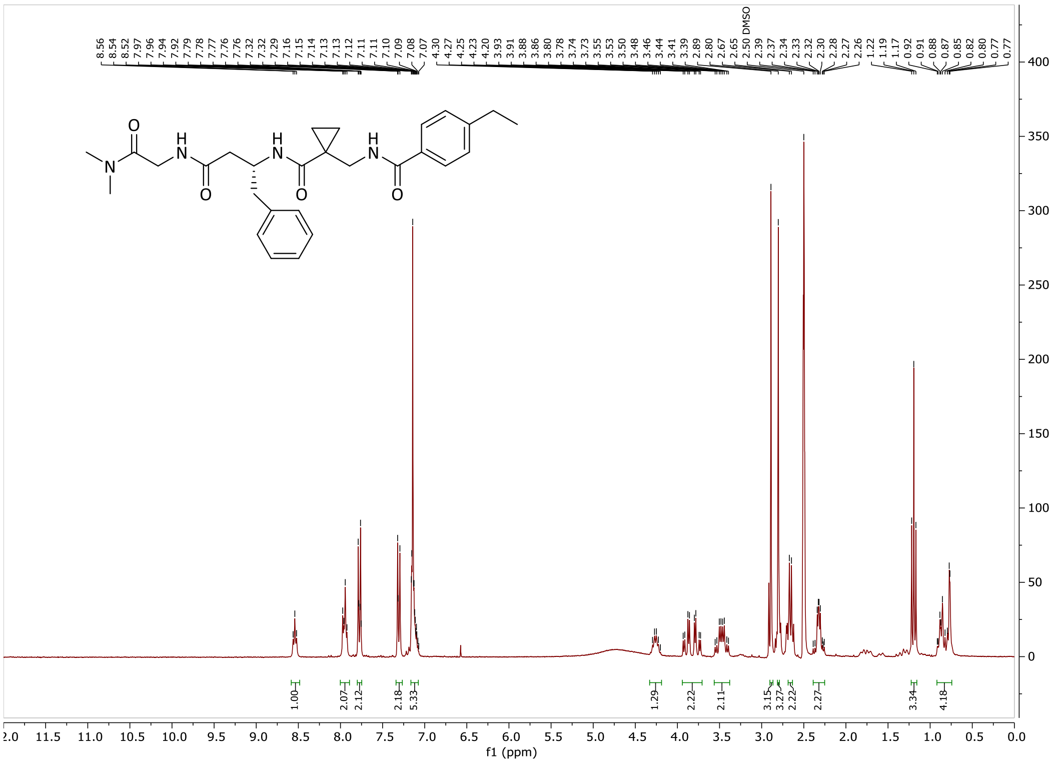

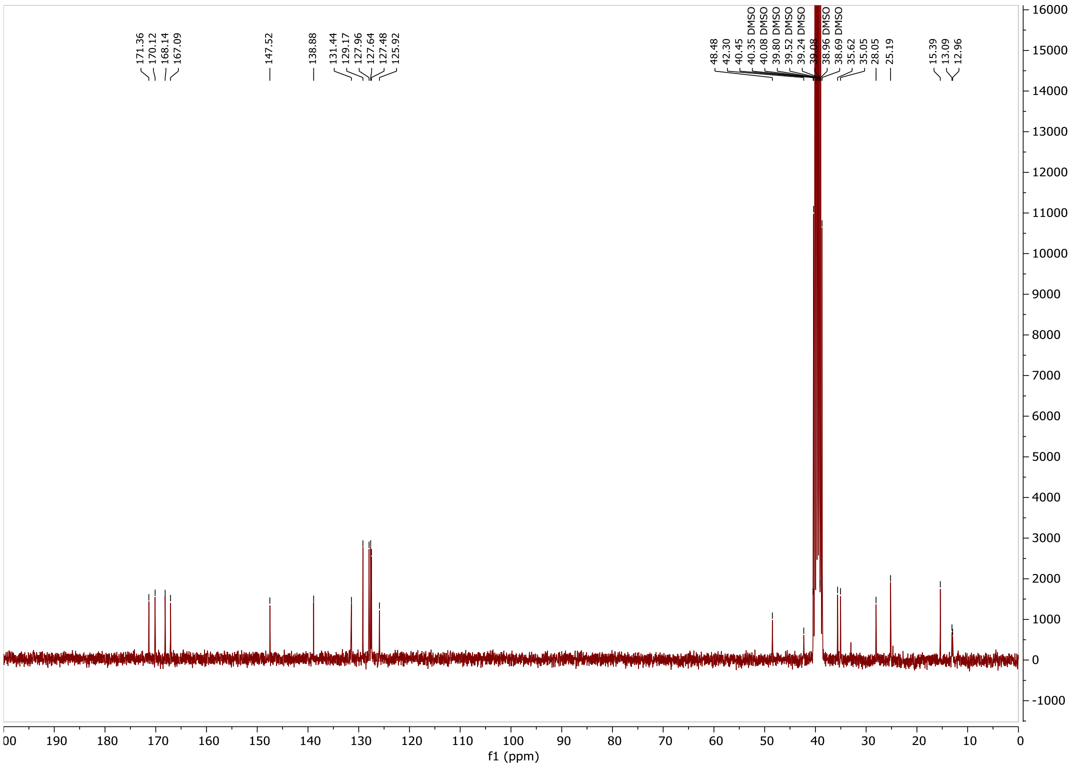

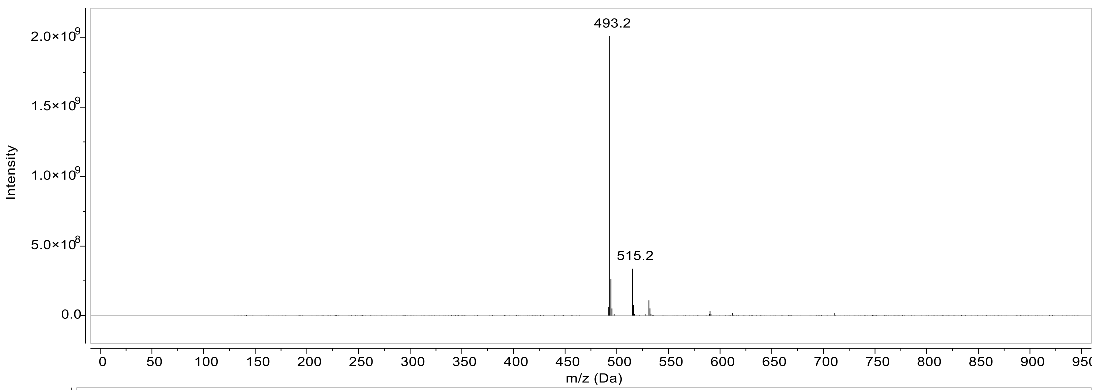
493.2
515.2
Intensity
2.0×10^9
1.5×10^9
1.0×10^9
5.0×10^8
0.0
m/z (Da)

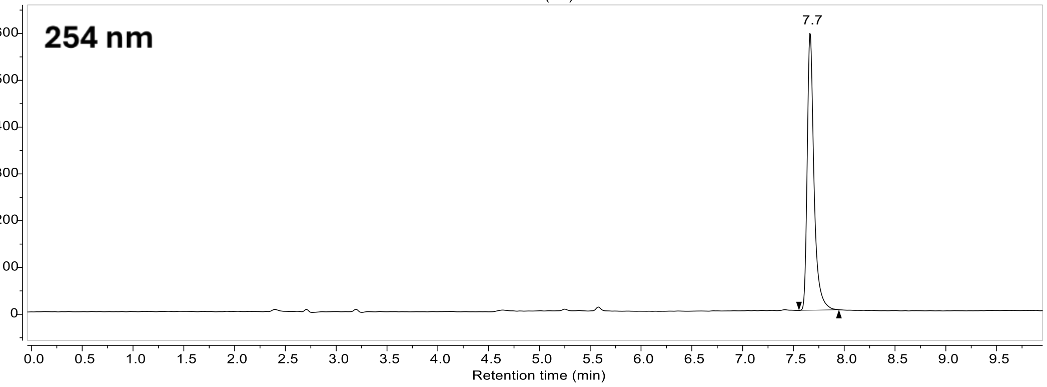
254 nm
7.7
Retention time (min)

#### Compound 46:

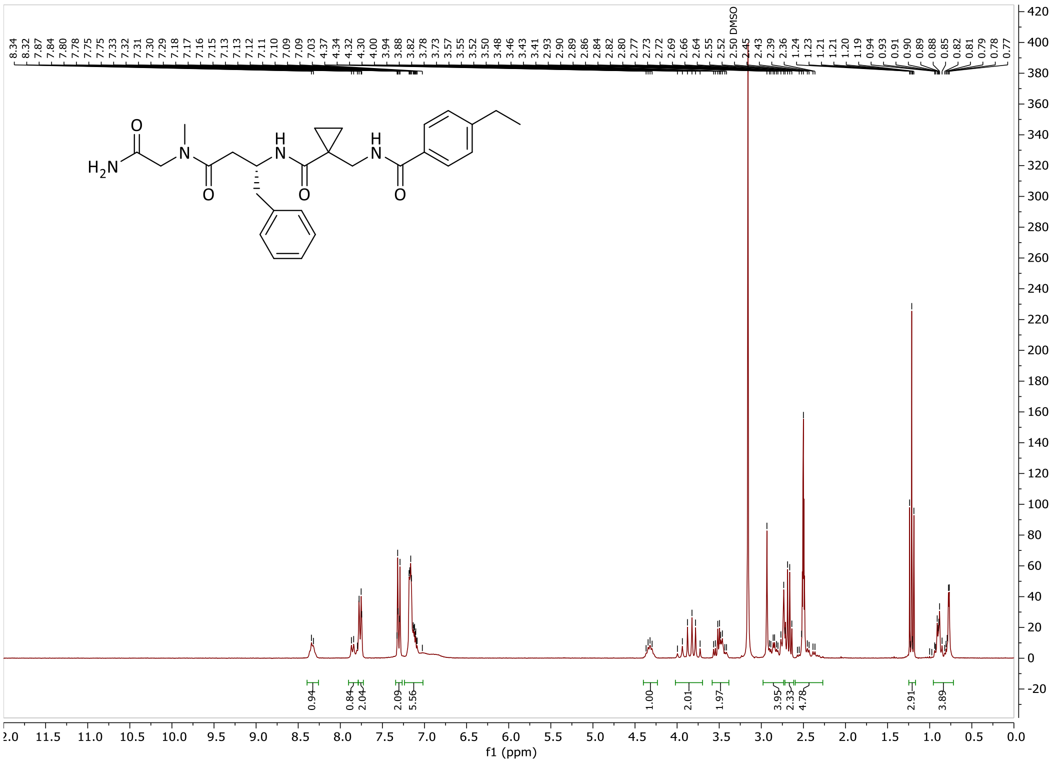

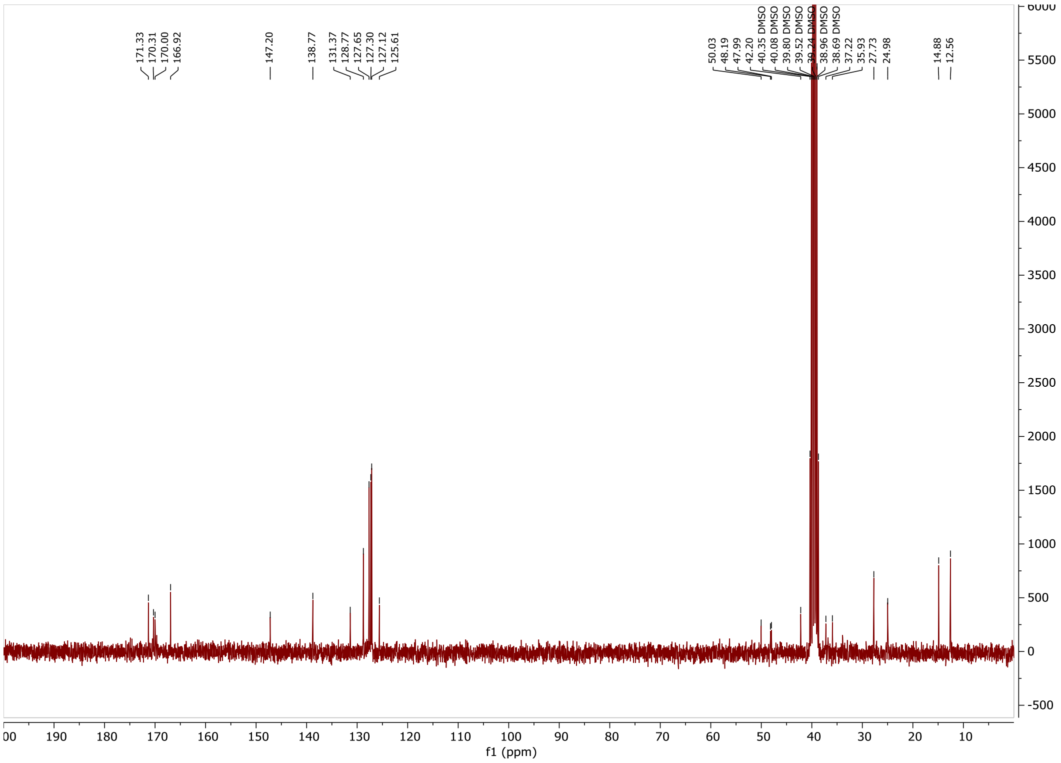

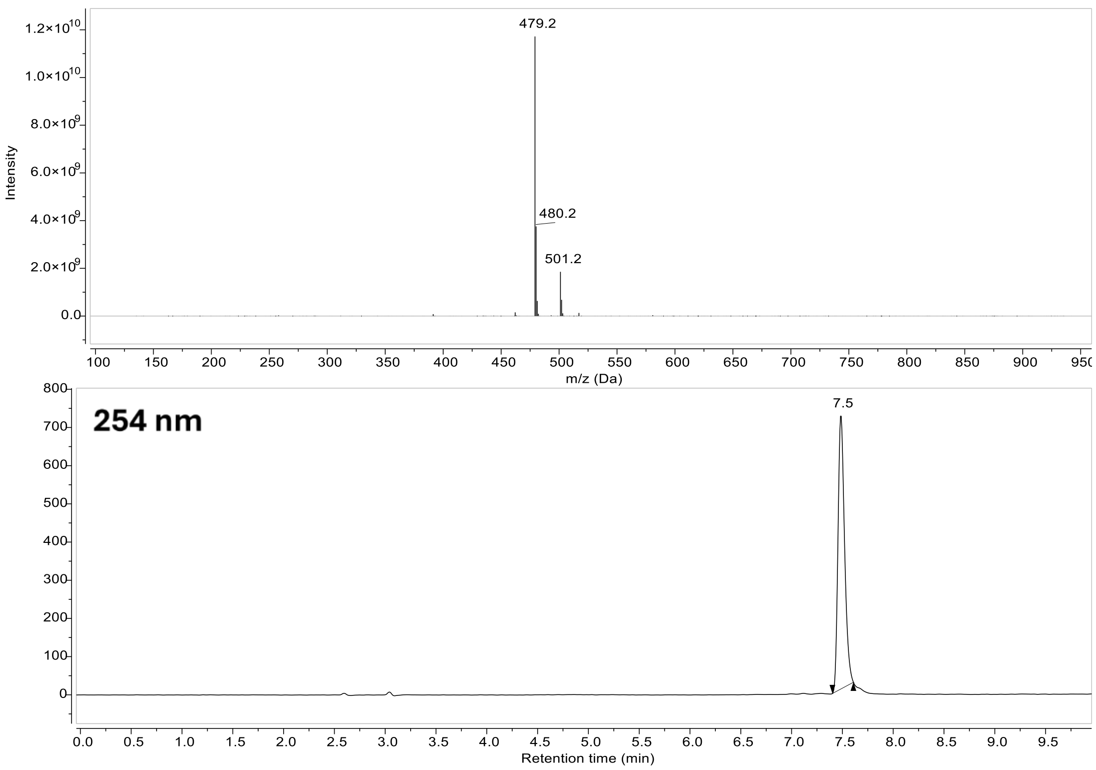

479.2
480.2
501.2
Intensity
m/z (Da)
254 nm
7.5
Retention time (min)

### Compound 44:

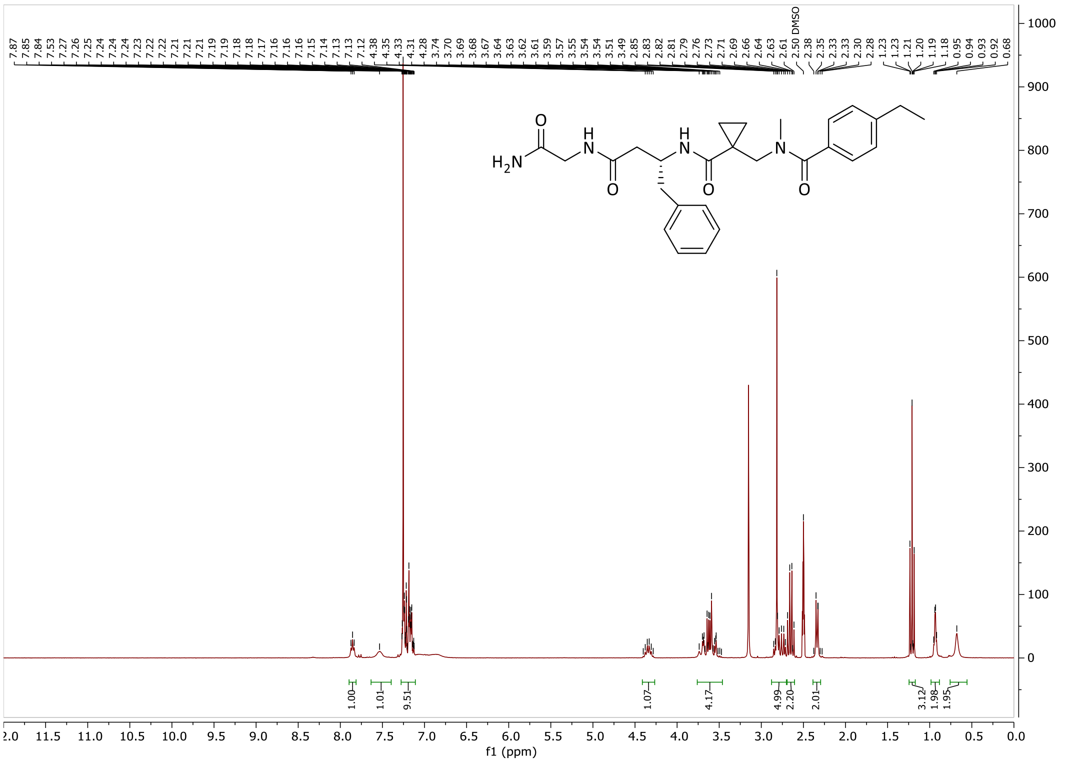
7.87 7.85 7.84 7.53 7.27 7.26 7.25 7.24 7.24 7.24 7.23 7.22 7.22 7.21 7.21 7.21 7.19 7.19 7.18 7.17 7.16 7.16 7.16 7.15 7.14 7.13 7.13 7.12 4.38 4.35 4.33 4.31 4.28 3.74 3.70 3.69 3.68 3.67 3.64 3.63 3.62 3.61 3.59 3.57 3.55 3.54 3.54 3.51 3.49 2.85 2.83 2.82 2.81 2.79 2.76 2.73 2.71 2.69 2.66 2.64 2.63 2.61 2.50 DMSO 2.38 2.35 2.33 2.33 2.30 2.28 1.23 1.23 1.21 1.20 1.19 1.18 0.95 0.94 0.93 0.92 0.68
H2N
H
N
O
1.00 1.01 9.51 1.07 4.17 4.99 2.20 2.01 3.12 1.98 1.95
f1 (ppm)

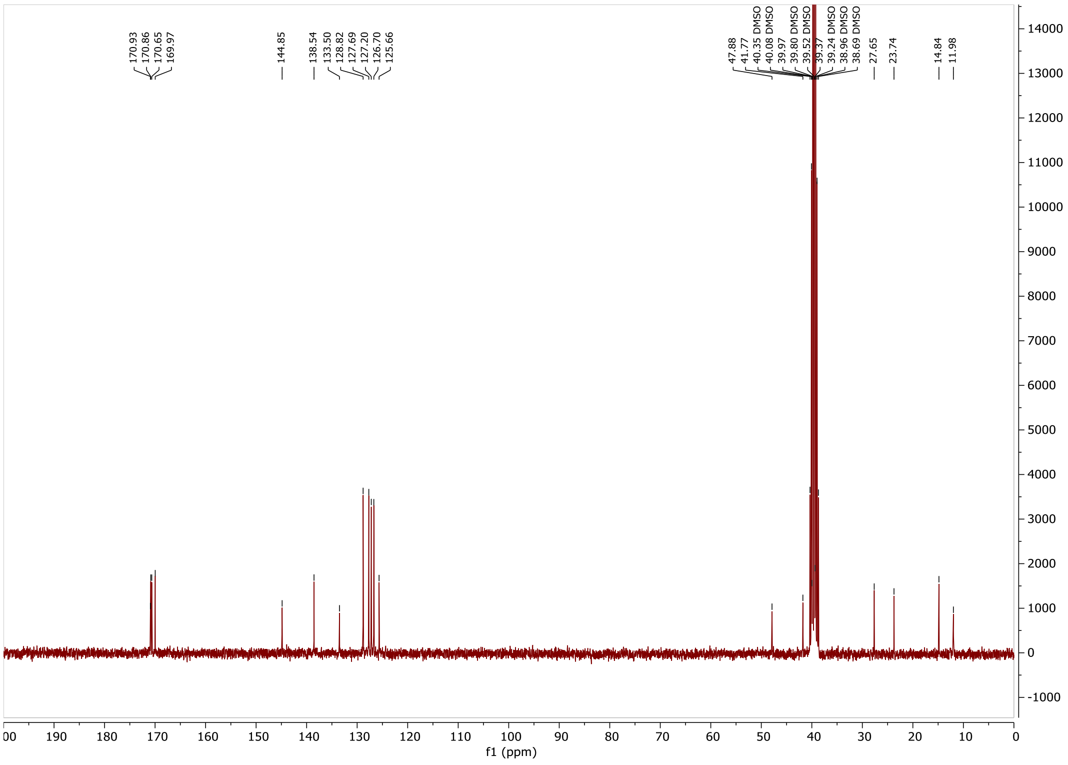
170.93 170.86 170.65 169.97 144.85 138.54 133.50 128.82 127.69 127.20 126.70 125.66
47.88 41.77 40.35 DMSO 40.08 DMSO 39.97 39.80 DMSO 39.52 DMSO 39.37 39.24 DMSO 38.96 DMSO 38.69 DMSO 27.65 23.74 14.84 11.98
f1 (ppm)

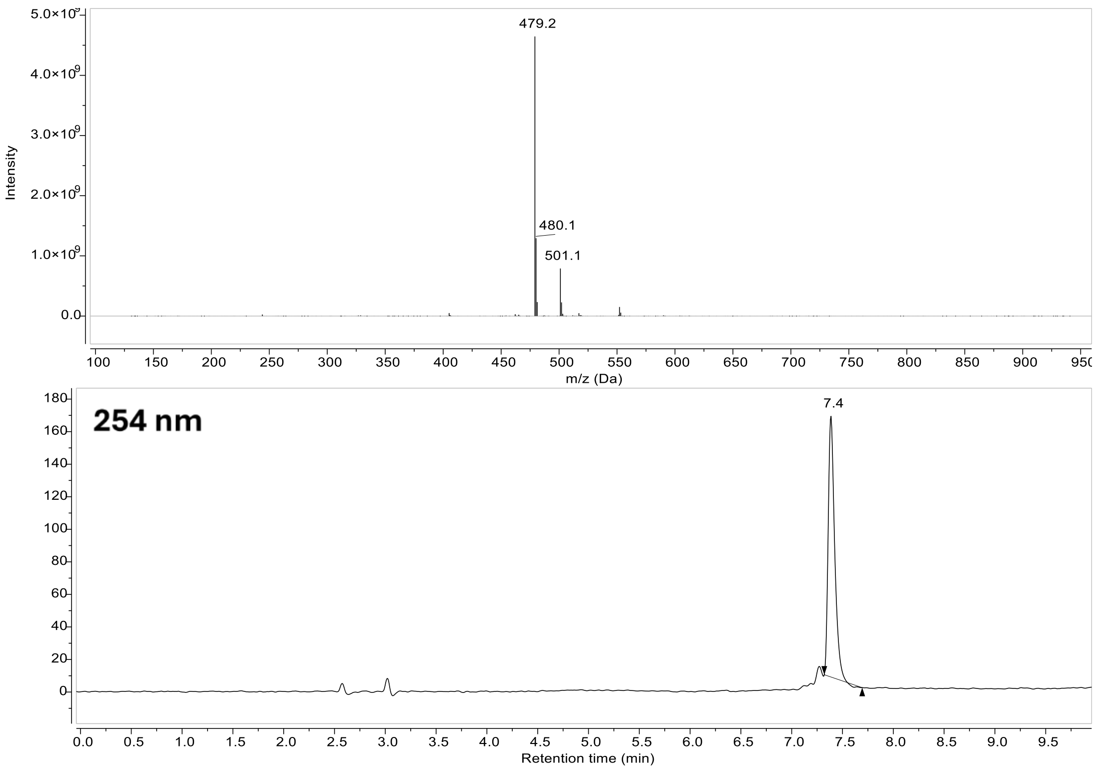

479.2
480.1
501.1
Intensity
m/z (Da)
254 nm
7.4
Retention time (min)

### Compound 9:

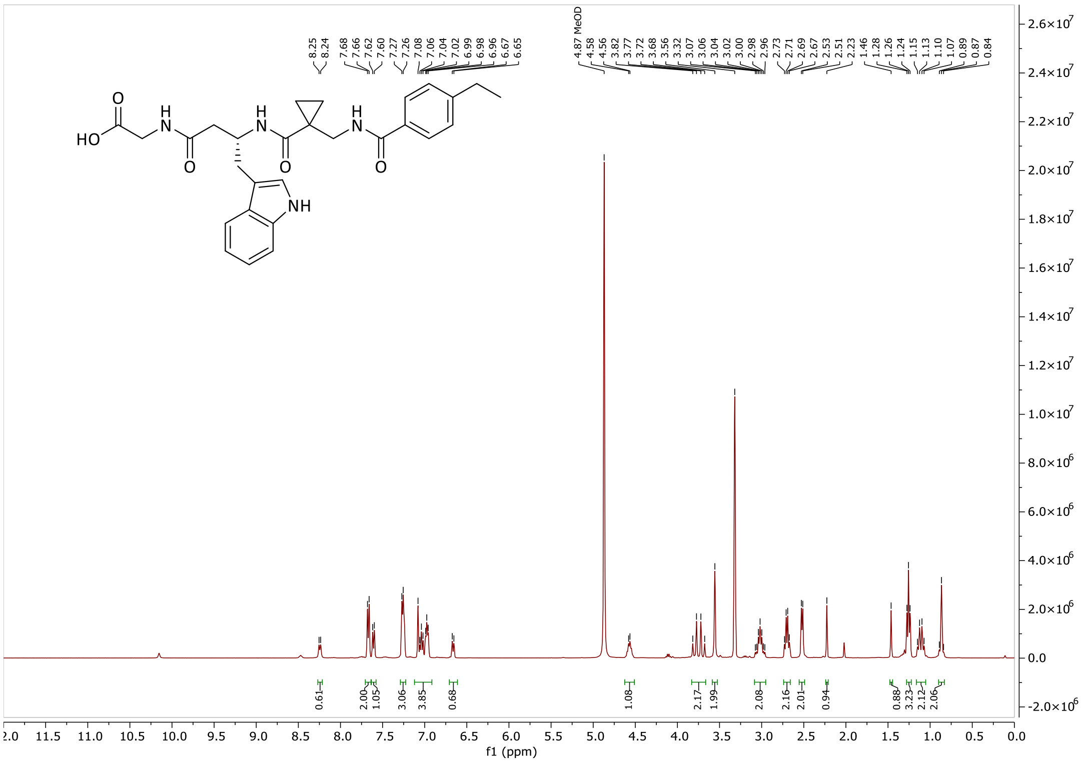

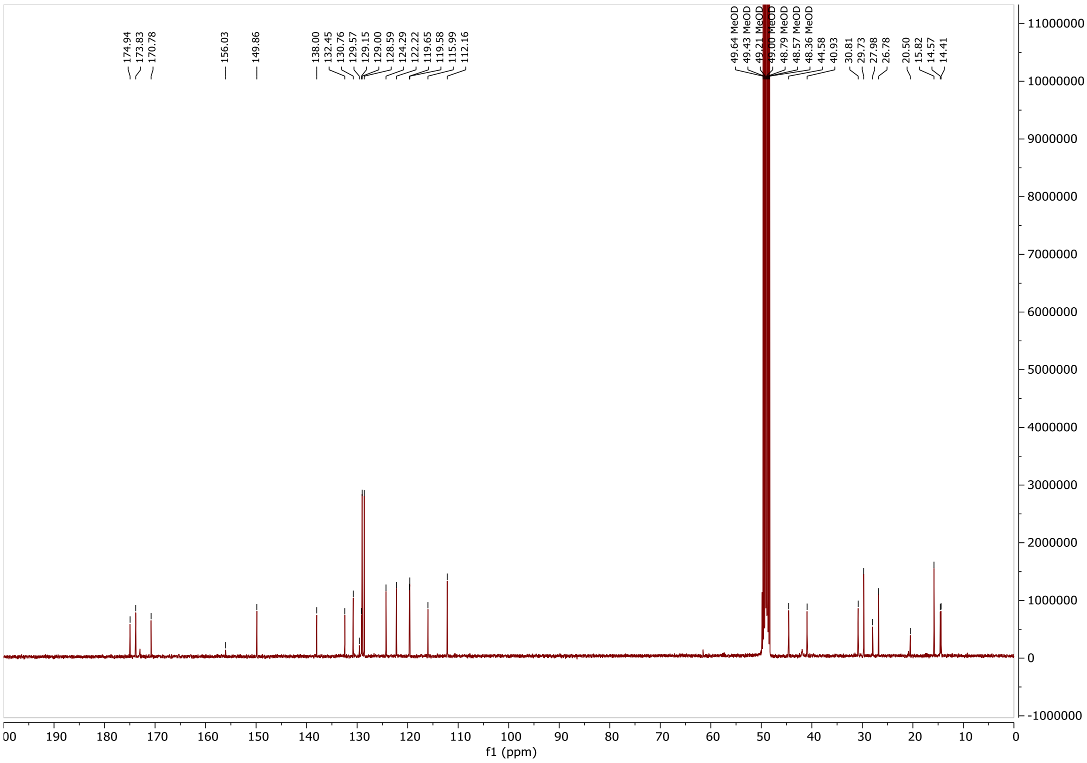

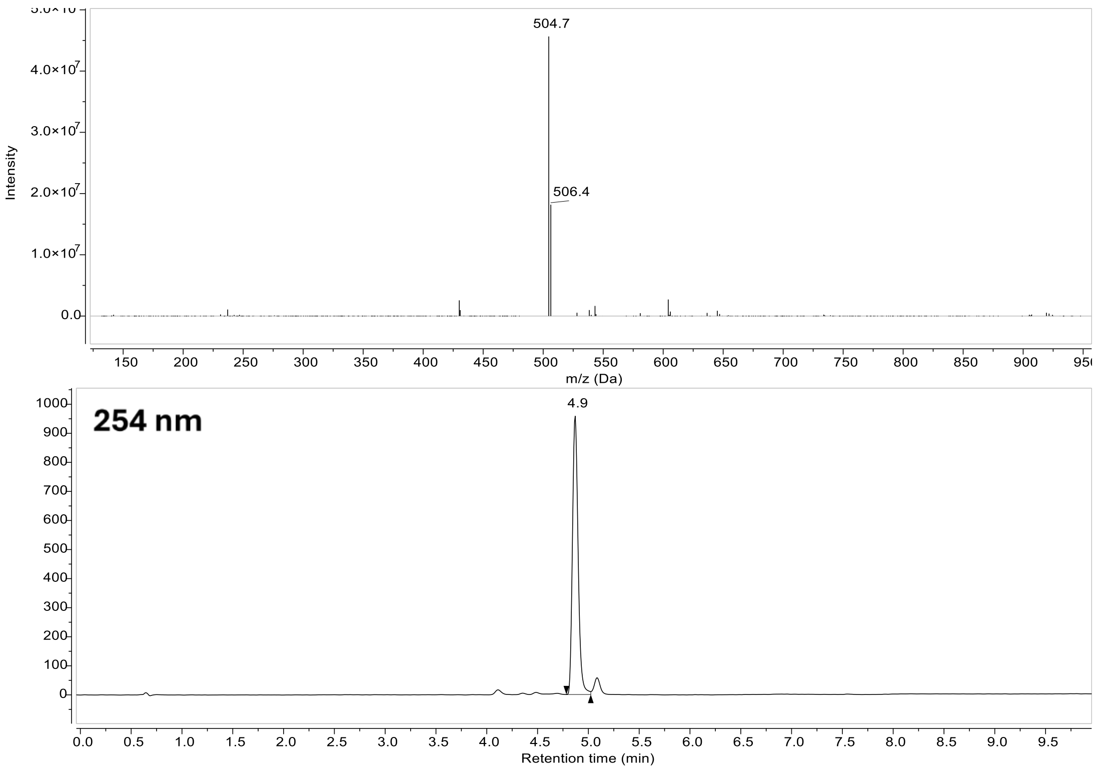

504.7
506.4
Intensity
m/z (Da)
254 nm
4.9
Retention time (min)

#### Compound 48:

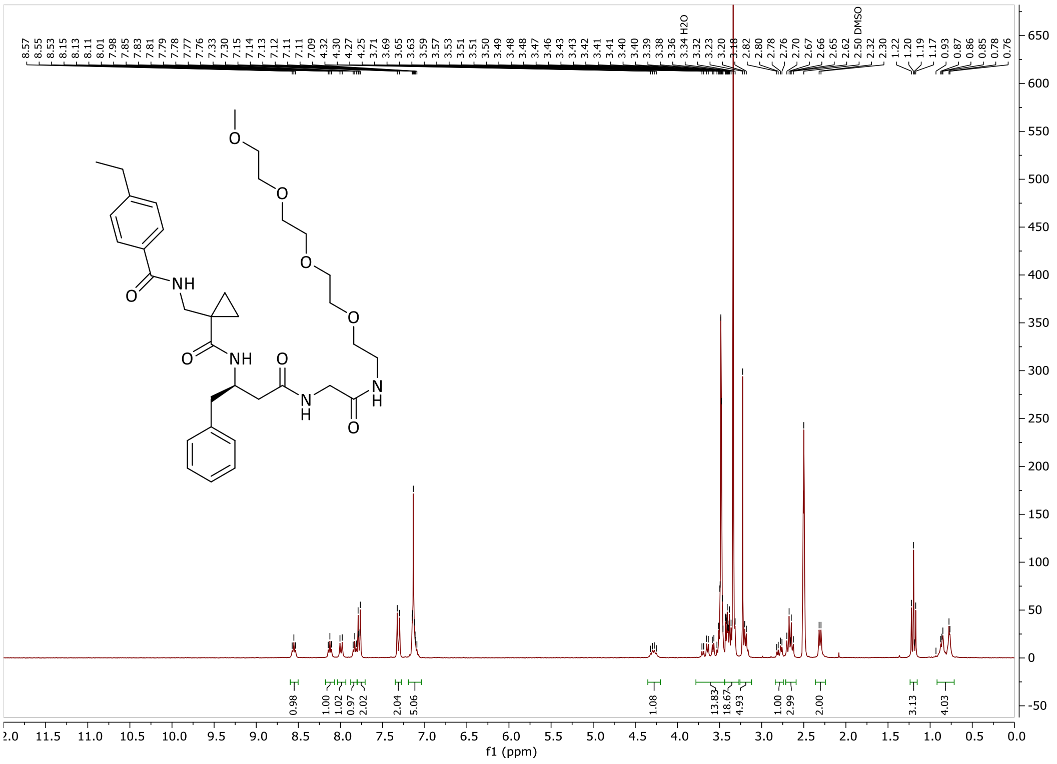
8.57 8.55 8.53 8.15 8.13 8.11 8.01 7.98 7.85 7.83 7.81 7.79 7.78 7.77 7.76 7.33 7.30 7.15 7.14 7.13 7.12 7.11 7.11 7.09 4.32 4.30 4.27 4.25 3.71 3.69 3.65 3.63 3.59 3.57 3.53 3.51 3.50 3.49 3.48 3.48 3.47 3.46 3.43 3.43 3.42 3.41 3.41 3.40 3.39 3.38 3.36 3.34 H2O 3.32 3.23 3.20 3.18 2.82 2.80 2.78 2.76 2.70 2.67 2.66 2.65 2.62 2.50 DMSO 2.32 2.30 1.22 1.20 1.19 1.17 0.93 0.87 0.86 0.85 0.78 0.76
NH
O
NH
O
O
N
H
N
H
O
O
O
O
O
0.98 1.00 1.02 0.97 2.02 2.04 5.06 1.08 13.83 18.67 4.93 1.00 2.99 2.00 3.13 4.03
f1 (ppm)

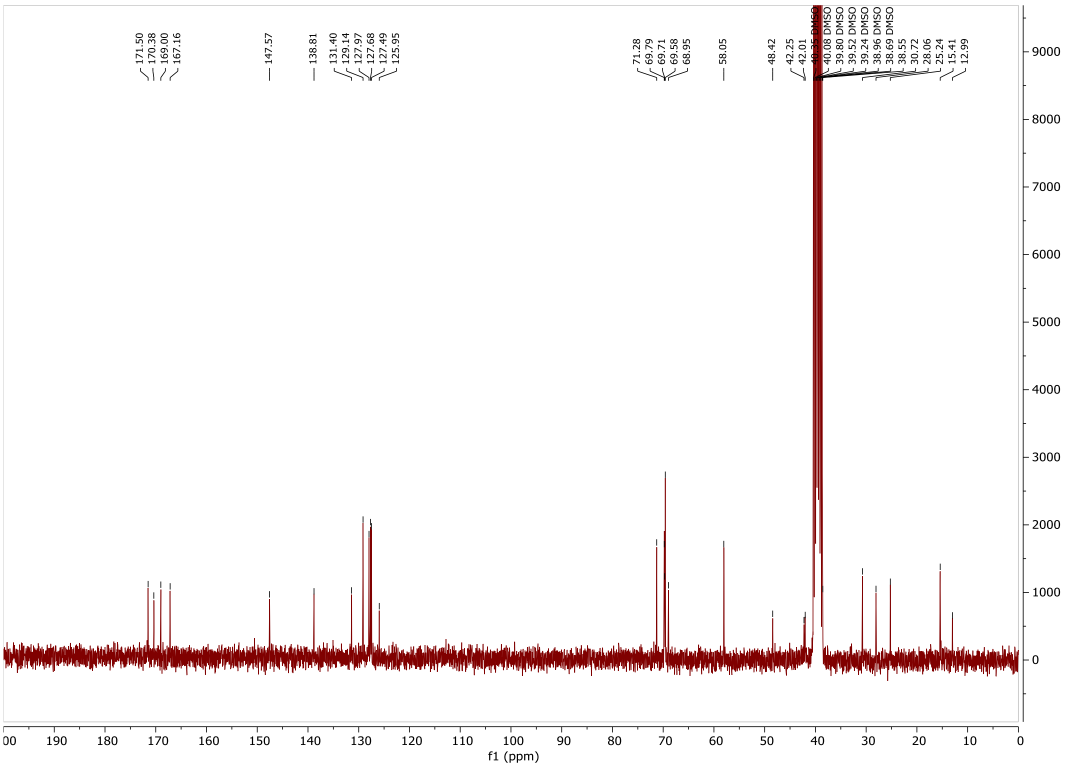
171.50 170.38 169.00 167.16 147.57 138.81 131.40 129.14 127.97 127.68 127.49 125.95 71.28 69.79 69.71 69.58 68.95 58.05 48.42 42.25 42.01
40.08 DMSO 39.80 DMSO 39.52 DMSO 39.24 DMSO 38.96 DMSO 38.69 DMSO 38.55 30.72 28.06 25.24 15.41 12.99
f1 (ppm)

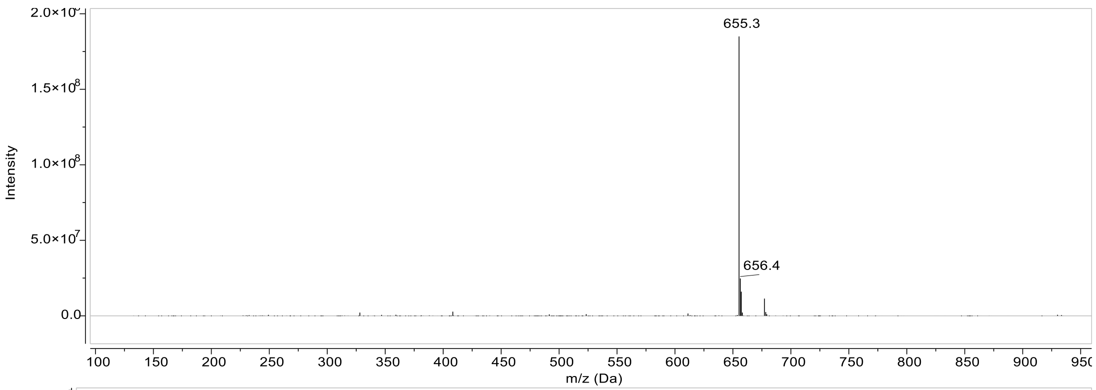
655.3
656.4
Intensity
m/z (Da)

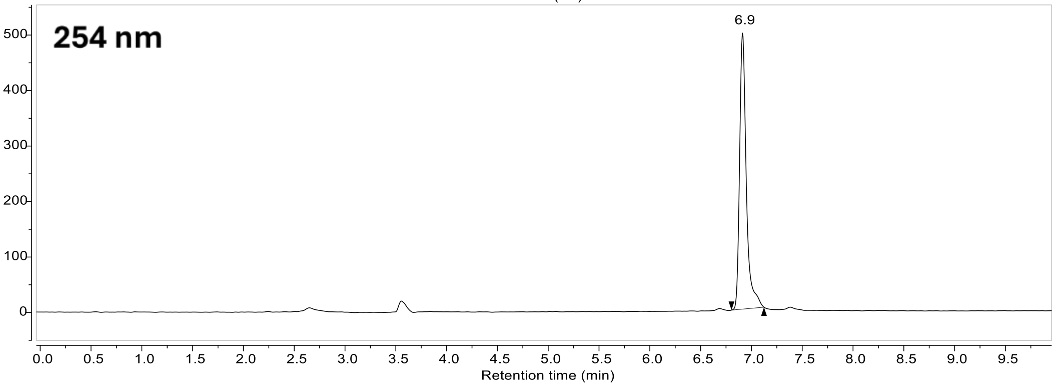
254 nm
6.9
Retention time (min)

#### Compound 7:

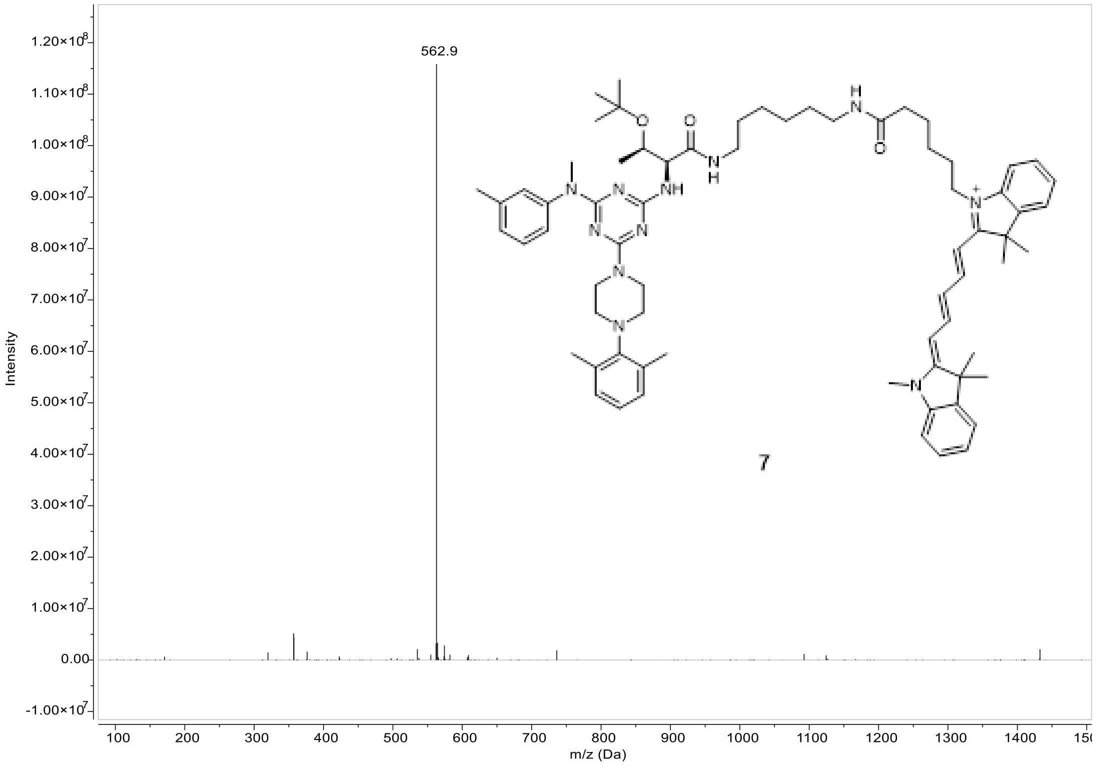

562.9
1.20×10^8
1.10×10^8
1.00×10^8
9.00×10^7
8.00×10^7
7.00×10^7
6.00×10^7
5.00×10^7
4.00×10^7
3.00×10^7
2.00×10^7
1.00×10^7
0.00
-1.00×10^7
Intensity
100
200
300
400
500
600
700
800
900
1000
1100
1200
1300
1400
m/z (Da)
7

#### Compound 6:

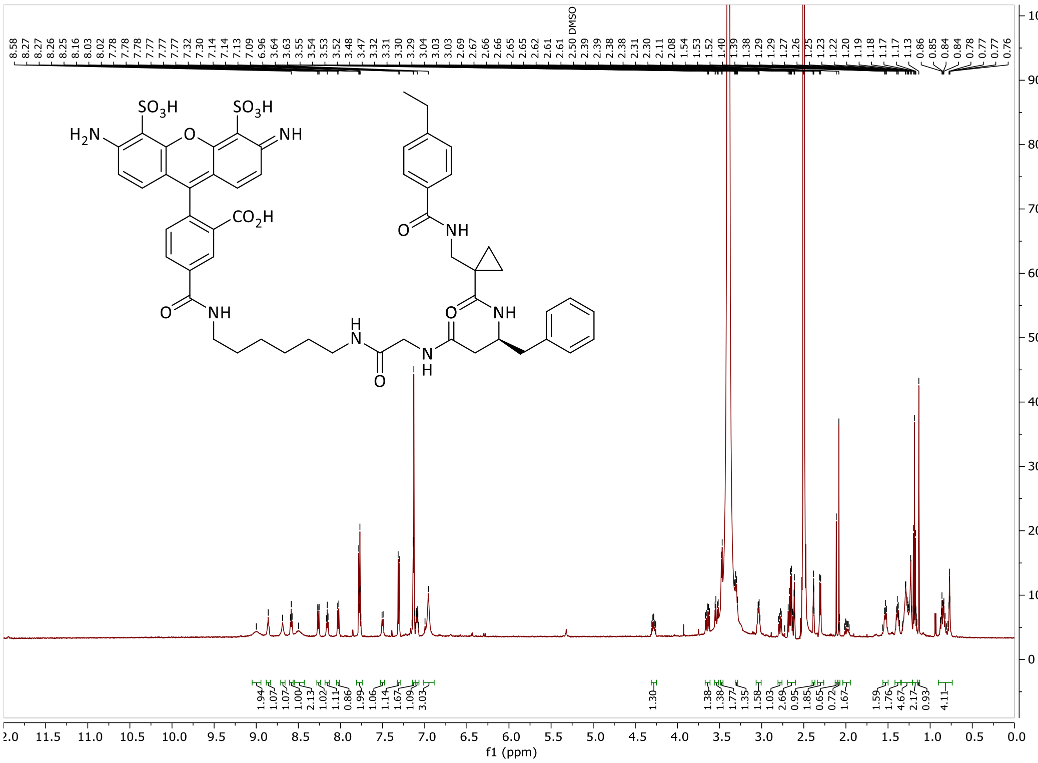
f1 (ppm)
2.50 DMSO

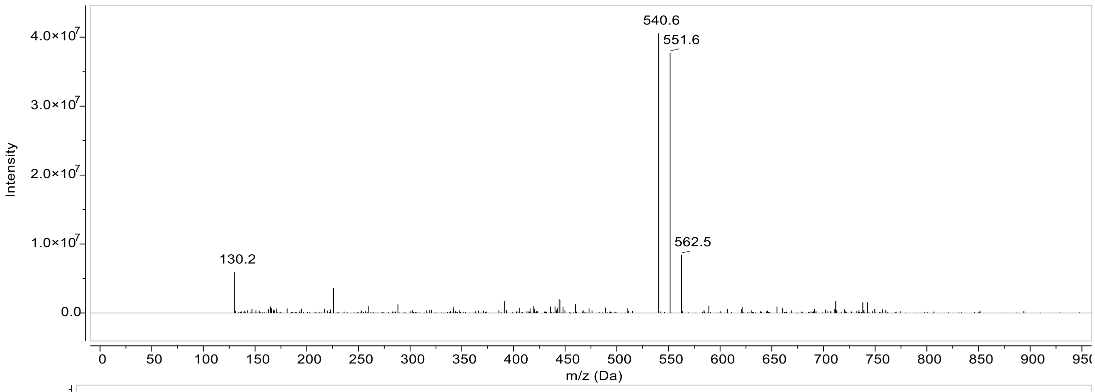
540.6
551.6
562.5
130.2
Intensity
m/z (Da)

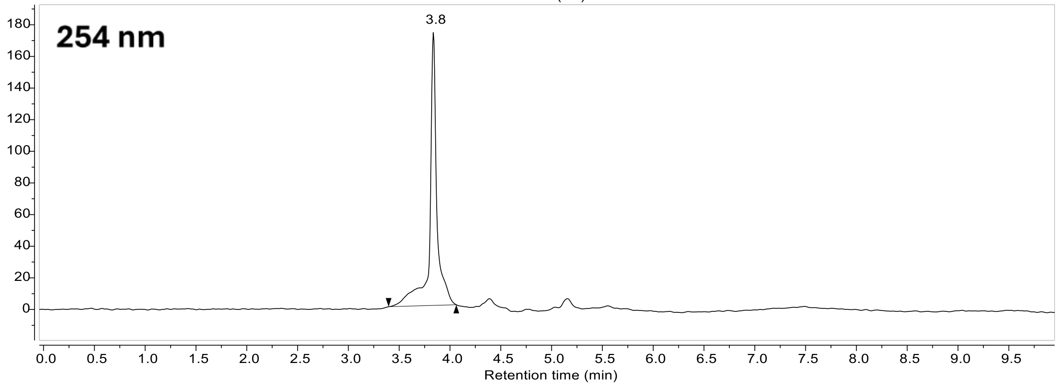
254 nm
3.8
Retention time (min)

#### Compound 8:

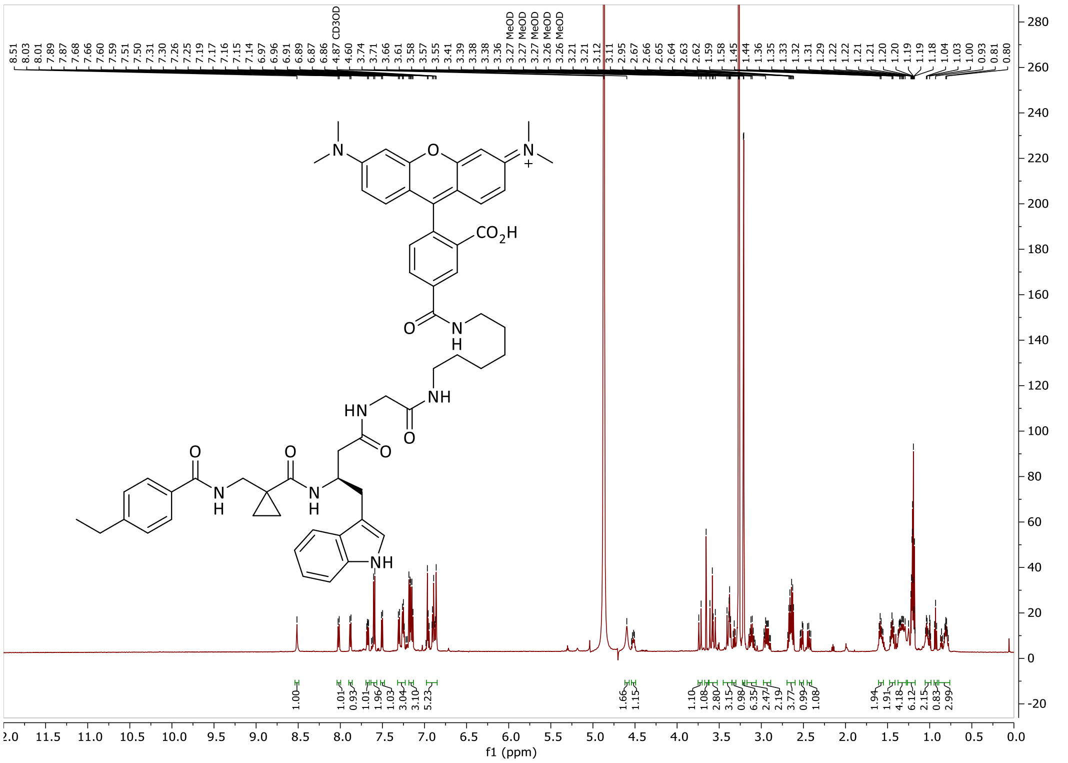

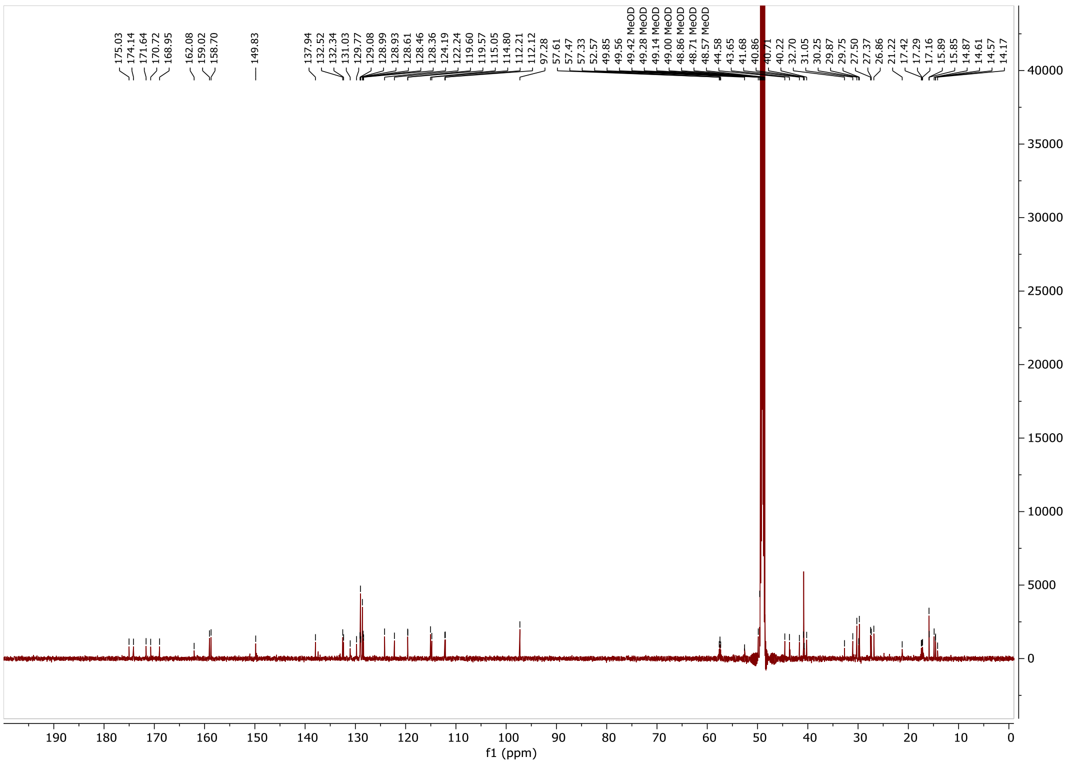

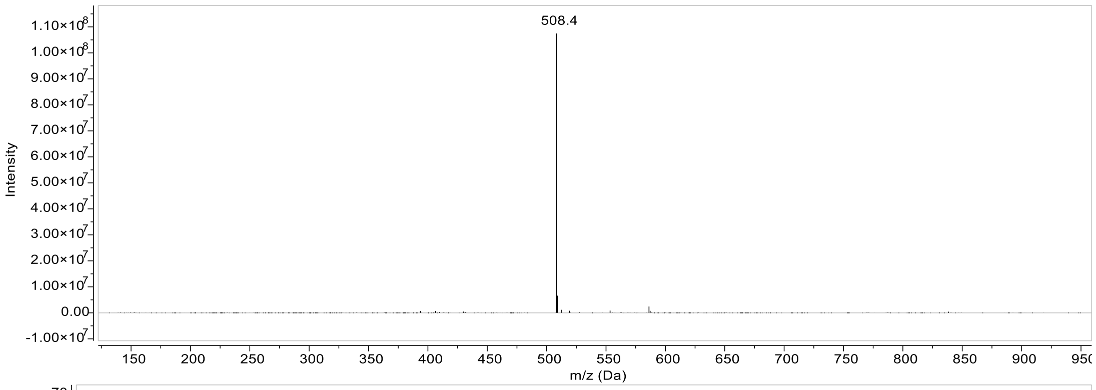

508.4
Intensity
m/z (Da)

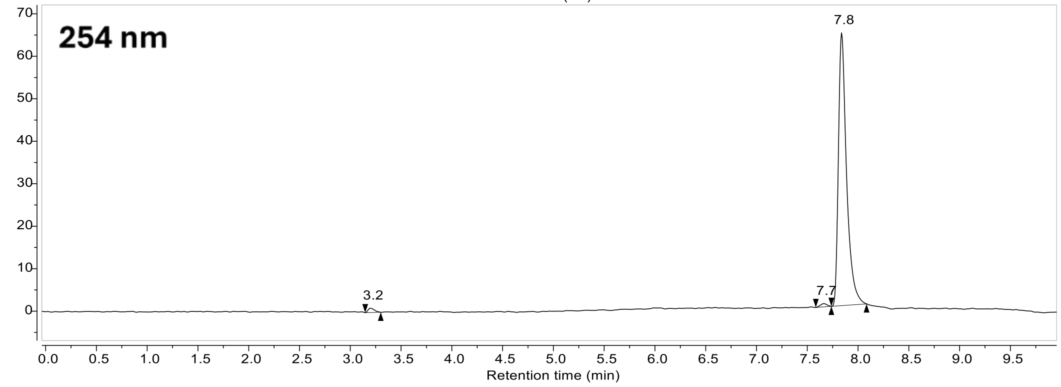

254 nm
7.8
7.7
3.2
Retention time (min)
